# A Comprehensive Benchmarking of Spatial Deconvolution and Domain Detection Methods across Diverse Tissues and Spatial Transcriptomic Technologies

**DOI:** 10.64898/2026.05.11.724248

**Authors:** Ajita Shree, V Aditya, Tanush Kumar, Hamim Zafar

**Author notes:** Authors contributed equally (V Aditya, Tanush Kumar).

## Abstract

Spatial transcriptomic technologies enable high-resolution characterization of gene expression patterns and reconstruction of cellular architecture within tissue contexts. Two key computational problems have emerged for analyzing these datasets: spatial deconvolution, for disentangling cell-type compositions at spatial locations, and spatial domain detection, for identifying spatially coherent regions within a tissue section. Although numerous methods have been developed for each task, a comprehensive and unified benchmarking study spanning diverse tissue types, spatial resolutions, and technological platforms remains lacking, hindering informed method selection by end users and impeding future methodological advancements. Here, we present spDDB (https://github.com/Zafar-Lab/spDDB), a comprehensive benchmarking framework for spatial deconvolution and domain detection methods across a large and diverse collection of datasets spanning multiple tissues, technologies, and biological conditions. We evaluated 21 deconvolution methods, including seven recently-developed approaches, across 37 datasets curated from brain, cancer, and organ tissues encompassing four distinct technologies. To enable rigorous evaluation, we introduced SynthST, a deep graph attention autoencoder-based simulator that generates realistic spatial cell-type distributions from spatial transcriptomic data, and employed a suite of spatial bivariate metrics including a novel bivariate Geary’s C metric, alongside rare cell-type, and cell-shape characterization metrics, for multidimensional performance assessment. While Cell2location, RCTD and SONAR emerged as top-performing methods for spatial deconvolution across tissue types, deconvolution performance varied substantially based on tissue architecture, spatial technology, dataset scale, and cell type diversity. For domain detection, we benchmarked 18 methods across 36 datasets spanning six spatial technologies, identifying PROST, BASS, and SpaceFlow as the leading approaches, while revealing notable limitations of existing methods in handling large-scale datasets. Finally, we provide practical guidelines to assist end users in selecting optimal methods for both tasks across diverse experimental settings.

## Introduction

Recent advances in spatial transcriptomic (ST) technologies enable high-throughput profiling of mRNA molecules while preserving their spatial locations within the tissue architecture, thereby providing complementary insights into the cellular microenvironment that are not captured by single-cell RNA sequencing (1; 2). By enabling the discovery of novel spatial transcriptional patterns, reconstruction of tissue-wide expression landscapes, and inference of interactions among neighboring cells and their microenvironments, ST technologies have offered new insights into the role of cellular heterogeneity and tissue architecture in development and diseases (3; 4; 5; 6). Current ST technologies can be broadly classified into two groups: imaging-based and sequencing-based techniques, each offering distinct yet complementary advantages (7). Imaging-based spatial methods, such as in situ sequencing and hybridization can capture mRNA at a subcellular resolution with high sensitivity, but are limited by low gene coverage and throughput (8; 9). In contrast, sequencing-based methods use spatially-barcoded oligonucleotides to capture mRNA, enabling unbiased, transcriptome-wide profiling of tissue sections at scale (10; 11). A key limitation of sequencing-based platforms, however, is their relatively low spatial resolution, where each capture area (or ‘spot’) typically samples multiple cells of distinct types. Although recent advances have reduced spot diameters from 100 µm to near-subcellular dimensions (12; 13), individual spots rarely correspond to single cells, necessitating computational deconvolution to infer their cell-type composition.

To tackle the limited resolution of sequencing-based techniques, several computational methods have been developed for cell-type deconvolution, which estimate the contribution of all cell types in each spot (14). These methods typically yield cell-type proportion estimates of all spots in the dataset enabling the reconstruction of the fine-grained cellular landscape of heterogeneous tissues. Given the availability of wide range of deconvolution methods employing diverse computational models, selecting the most suitable approach depends on various factors including the underlying spatial technology, experimental design, tissue type and specific study objectives. Although recent studies (15; 16; 17) have performed benchmarking of different deconvolution methods, their scope has been largely limited, as these evaluations focused primarily on a narrow range of tissue types, predominantly brain datasets. Moreover, prior benchmarking studies relied on simplified simulation strategies (e.g., uniform distribution of cell types across spots) which fail to recapitulate the realistic spatial cell-type distributions and biological variability observed across tissue types. These simulations have been largely confined to 10x Visium-like datasets, underscoring the need to extend benchmarking to higher-resolution platforms such as Slide-seq (18) and 10x Visium HD. Furthermore, the conclusions in previous studies have also been extracted based on non-spatial evaluation metrics, which do not adequately assess prediction performance within a spatial context. Another limitation of existing studies is that evaluations are typically performed globally across all cell types, overlooking accuracy at the level of individual cell types. Compounding these limitations, new deconvolution methods continue to emerge at a rapid pace, each claiming improved accuracy, underscoring the pressing need for a comprehensive, spatially informed benchmarking framework that systematically evaluates accuracy and robustness across realistic settings (Supplementary Figure 1a).

Another core computational problem in spatial transcriptomics is the detection of spatial domains for delineating tissue microenvironments and characterizing functional roles of different tissue regions (19). A fundamental step in this process is the clustering of spatial spots to identify spatially coherent regions, leveraging both gene expression profiles and positional information of spots. This requirement has fueled the development of numerous statistical and graph-based deep learning methods for the clustering of spots into spatial domains. Subsequently, some studies (20; 21; 22) performed benchmarking of existing domain detection methods, yet remain limited in scope and rigor. These benchmarks predominantly relied on a limited diversity of datasets (primarily brain samples), which do not represent the full spectrum of spatial domain organization present across diverse tissue types. Additionally, cross-platform robustness and scalability across resolutions and technologies (e.g., 10x Visium vs. Stereo-seq) are rarely evaluated. Meanwhile, the steady emergence of new domain detection methods, each claiming superior performance over earlier approaches, further motivates the need for a comprehensive, biologically grounded benchmarking framework.

To address these gaps, we present spDDB (spatial <u>D</u>econvolution and <u>D</u>omain detection <u>B</u>enchmarking), a comprehensive benchmarking study encompassing 21 cell-type deconvolution methods and 18 domain detection methods. Our deconvolution benchmarking assembles an extensive dataset repository spanning 21 distinct tissue types curated from three major tissue categories - Brain, Cancer and Organs. Notably, our cancer datasets encompass 13 cancer subtypes,, compared to only one to two in prior benchmarks (Supplementary Figure 1b). A major focus of our study is on sequencing-based spatial transcriptomics datasets, which provide full transcriptomic profiles and are thus more suitable for evaluating deconvolution performance. To overcome the longstanding challenge of obtaining ground-truth cell-type proportions, we introduce SynthST (<u>Synth</u>etic <u>S</u>patial Transcriptomics <u>T</u>issue), a novel graph attention autoencoder-based simulator that generates cell type proportions and gene expression profiles with realistic spatial patterns for sequencing-based ST. The multi-view training strategy of SynthST ensures that dominant spatial structures are effectively captured for all cell types, including rare populations. Our benchmarking further includes datasets generated by recently introduced higher-resolution platforms such as Visium HD, enabling performance assessment across a wide range of spatial resolutions.

For evaluation, we introduce a suite of spatial bivariate metrics, including a novel bivariate Geary’s C, that more effectively assess prediction performance within a spatial context. We further incorporate cell shape characterization metrics (curl, elongation, linearity), and rare cell type metrics (rare score, regional rare score) to provide a multidimensional assessment of the method’s performance. In addition, we perform cell-type–specific analyses to identify major cell type groups that are more accurately predicted. Our benchmarking incorporates seven recently developed methods, and we conclude with practical guidelines for end users, highlighting the strengths of different approaches across diverse tissues, technologies, and experimental scenarios.

In the second part of spDDB, we present a comprehensive domain detection benchmarking framework that evaluates seven recently developed methods alongside the top-performing methods from prior studies. Our benchmark spans a diverse collection of datasets covering multiple tissue types and technologies, providing a broader and more representative assessment of current domain detection methods. While prior benchmarks were were restricted to six to seven tissue types, our study spans 13 tissue types and includes five cancer samples compared to only one to two in earlier studies. We also analyzed three large-scale datasets, providing critical insights into the computational scalability and biological fidelity of methods at the scale of emerging spatial atlases, and systematically assess cross-platform and cross-tissue robustness. Finally, we propose practical guidelines for end users, highlighting the strengths of different methods across various experimental scenarios.

## Results

### Overview of spatial cell-type deconvolution benchmarking

For a comprehensive evaluation of spatial deconvolution methods, we identified 21 state-of-the-art methods employing different computational techniques - linear model-based methods: Stereoscope (23), RCTD (24), Autogenes (25), and Redeconve (26); non-negative matrix factorization (NMF)- based methods: SpotLight (27), SpatialDWLS (28), CARD (29), and SpiceMix (30); graph-based methods: SD (31); deep Learning-based methods: Tangram (32), DestVI (33), GraphST (3), CellDART (34), and POLARIS (35); anchor-based methods: Seurat V3 (36), and Bayesian probabilistic methods: STRIDE (37), SpatialDecon (38), Cell2location (39), STDeconvolve (40), SONAR (41), and CelloScope (42) (Figure 1a, Supplementary Table 1). Out of 21 methods, two methods Spicemix and STDeconvolve did not require any external single-cell RNA-seq reference datasets and inferred spatial distribution of cell type proportions solely from the spatial coordinates and gene expression profiles of the spots in spatial transcriptomic datasets. The remaining 19 methods relied on reference scRNA-seq data derived from the same tissue as the spatial transcriptomics dataset.

**Figure 1:**
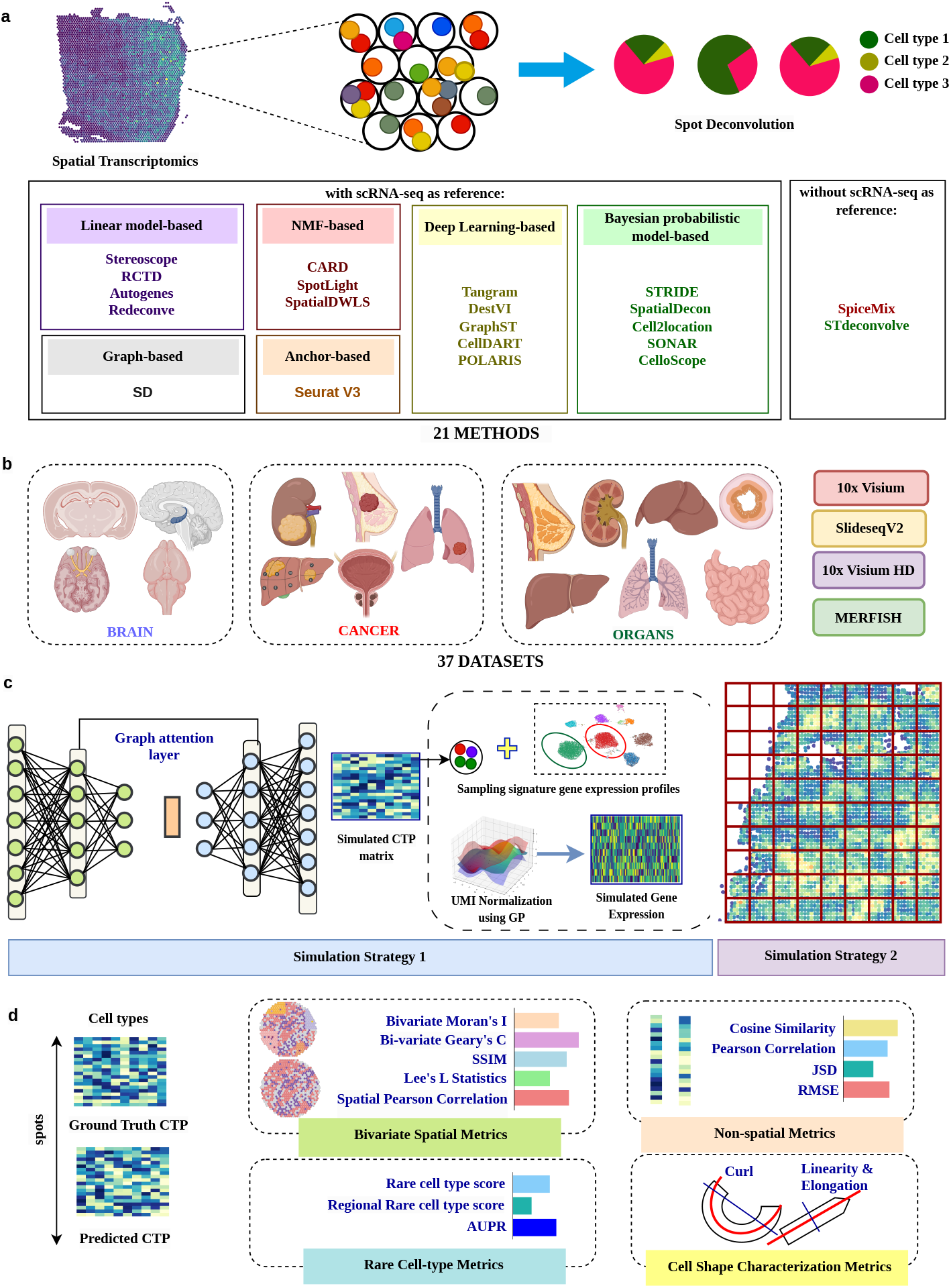
Overview of the benchmarking workflow for cell-type deconvolution. (a) Schematic representation of the benchmarking workflow for evaluating the performance of 21 cell-type deconvolution methods (i.e. Autogenes, CARD, Cell2location, CellDART, CelloScope, GraphST, Polaris, RCTD, Redeconve, Seurat, SONAR, SpatialDecon, SpatialDWLS, Spotlight, STDeconvolve, Stereoscope, STRIDE, Tangram, DestVI, SD, SpiceMix), categorized by their data requirements and computational strategies, using paired spatial transcriptomics and scRNA-seq datasets. (b) The benchmarking was performed on 37 datasets, spanning cancer, brain, and organ tissues, encompassing different spatial technologies including 10x Visium, Slide-seq V2, 10x Visium HD, and image-based sequencing techniques. c) Overview of simulation strategy 1 and simulation strategy 2 for generation of simulated spatial gene expression and cell type proportion datasets. (d) The benchmarking results were evaluated using a set of five bi-variate spatial and four non-spatial evaluation metrics, five shape characterization metrics and three rare cell-type metrics. Created in [BioRender](https://biorender.com/).

For the benchmarking of spatial deconvolution methods, we collected 37 real datasets generated using different spatial technologies including 10x Visium (43), Slide-seq V2 (44), 10x Visium HD (45), and MERFISH (Figure 1b) spanning three different tissue categories - brain, cancer and organs. For each spatial dataset, the corresponding scRNA-seq dataset was also acquired as a complementary reference resource. For the brain deconvolution tasks, we utilized 12 datasets derived from six different brain regions from human and mouse: Visium DLPFC (46; 47), Visium mouse brain (39), Slide-seq V2 hippocampus (24), Slide-seq V2 cerebellum (24), Visium HD mouse brain (45) (39), and MERFISH preoptic hypothalamic brain region (48)(Supplementary Figure 2a-f). For the cancer deconvolution tasks, we utilized 13 datasets derived from seven distinct cancer types: Visium kidney cancer (49), Visium breast cancer (50), Visium liver cancer (51; 52), Slide-seq V2 prostate cancer (53), Visium HD lung cancer (45; 54), MERFISH breast cancer (55), and MERFISH lung cancer (55) (Supplementary Figure 2g-m). For the organs deconvolution tasks, we collected 12 datasets across eight distinct tissue types spanning three species (human, mouse, and chicken): Visium lung development (56), Visium chicken heart (57), Visium human breast (50), Visium intestine (58), Visium mouse liver (59), Visium human liver (59), Slide-seq V2 kidney (60), and MERFISH ileum (61)(Supplementary Figure 2n-u, Supplementary Table 2). The real datasets were used for generating synthetic datasets with ground-truth realistic cell type proportion distributions based on which we performed our benchmarking. Overall, the deconvolution tasks presented distinct challenges, including variations in tissue type, variations in spatial resolution, species variety, variations in cancer subtypes, variations in cell type annotation resolutions and variations in tissue regions within the same tissue (Supplementary Table 1).

We introduced two simulation strategies for generating synthetic datasets to comprehensively evaluate deconvolution methods (Figure 1c). Extended Figure 1 shows the overview of SynthST, a novel simulator for the generation of datasets using simulation strategy 1. SynthST employs a deep graph attention (GAT) network to generate a synthetic cell type proportion matrix based on the cell type proportions inferred from a real spatial dataset. For inferring the cell type proportions from the real dataset, we employed top-performing deconvolution methods identified in existing benchmarking studies (16; 15; 17) and the cell type proportion matrices obtained from these methods were used for training the GAT to achieve multi-view learning, allowing each cell type to be modeled against the same graph topology (see Methods). This approach ensured that, for every cell type, its most dominant spatial distribution pattern was captured in the simulated data. Moreover, as different spatial patterns were captured by different subsets of methods, the multi-view strategy helped preserve spatial structures for all the cell types. Furthermore, the weighted neighborhood aggregation operation of the graph attention network enabled SynthST to generate smooth spatial patterns for cell type proportions (62). The simulated cell-type proportion matrix was then used to generate spatial gene expression profiles by weighted aggregation of sampled gene expression vectors from the cell type-specific gene signature matrix derived from the reference single-cell data. We employed Gaussian processes to model the target UMI distribution for the simulated spatial gene expression dataset, enabling appropriate downsampling while preserving global structural patterns present in the original spatial gene expression dataset (see Methods). Our simulation strategy 2 uses a binning approach on single-cell–resolution imaging-based datasets (e.g., MERFISH) to generate simulated gene expression and ground-truth cell type proportions for spots. Such binning approaches have been employed in prior benchmarking studies (15; 16); however, these studies used datasets with limited gene coverage or small numbers of spots. In contrast, we leveraged datasets with higher number of captured genes and ensured greater diversity in both the number and composition of cell types represented across deconvolution tasks (Supplementary Figure 1b).

We conducted a comprehensive evaluation of 21 deconvolution methods by comparing their predicted cell type proportion (CTP) matrices to the simulated ground truth CTP matrix using four sets of evaluation metrics: (1) Bivariate Spatial Metrics, (2) Non-spatial Metrics, (3) Rare Cell-type Metrics, and (4) Cell Shape Characterization Metrics. We employed five bivariate spatial evaluation metrics including four existing metrics: bivariate Moran’s I (63), SSIM (16), Lee’s L Statistics (64), and spatial Pearson correlation metric (65; 66) and a novel bivariate Geary’s C metric as these metrics are particularly useful for assessing whether inferred cell type proportions preserve expected spatial co-localization and exclusion patterns across neighboring locations. We developed the novel bivariate Geary’s C metric by extending the univariate Geary’s C metric to a bivariate setting to quantify spatial correlation between cell-type proportions while explicitly capturing local similarity, in contrary to existing bivariate metrics which primarily emphasize global similarity (63; 65; 64). Univariate Geary’s C measures local spatial similarity by summing the squared differences between neighboring nodes, with smaller values indicating stronger spatial autocorrelation. Extending this concept, we defined a bivariate Geary’s C in which, for a given cell type, squared differences are computed between each node in the ground-truth proportion vector and its neighboring nodes in the predicted proportion vector. When the two spatial patterns are well aligned, these differences remain small, reflecting strong local spatial agreement (see Methods). Non-spatial evaluation metrics include Cosine Similarity, Pearson Correlation, Jensen Shannon Divergence (JSD), and Root Mean Square Error (RMSE). Rare cell type evaluation metrics include composite score for rare cell types, composite score for regional rare cell types and area under precision and recall (AUPR) metrics and these metrics evaluate the performance of a deconvolution method in capturing the cell type abundances of rare cell types. We defined regional rare cell types as rare cell types that exhibit bimodal or multimodal spatial distributions, characterized by their presence in multiple, distinct spatial regions within the tissue. Finally, cell shape characterization metrics evaluate the ability of a deconvolution method in capturing the spatial abundance pattern for cell types exhibiting high curl (twisted or curved), high and low elongation (measures the ratio of the major and minor axes) and high and low linearity (measures the degree of linearity of the shape of cell type distribution) (see Methods) (Figure 1d). We also evaluated the methods based on usability metrics, including the frequency of top ranking, code quality, ease of installation, and computational resource requirements (time and memory) for both small-scale datasets (e.g., Visium) and large-scale datasets (e.g., Slide-seq V2 and Visium HD).

### Cell2location, RCTD and SONAR achieve the highest accuracy in cell-type de-convolution

To assess the overall performance of cell-type deconvolution methods, we computed four composite scores for each dataset: the Composite Bivariate Spatial Score, Composite Non-Spatial Score, Composite Shape Characterization Score, and Composite Rare Celltypes Score, each aggregating the corresponding set of performance metrics within its category. Further, across our extensive experiments spanning a wide range of datasets, we observed that some methods were unable to complete execution on particular datasets and therefore produced no results (Supplementary Figure 3a). To ensure that the methods are penalized for such incomplete executions, while computing the composite score, we employed a slope-based penalization approach that applies a gradually increasing penalty proportional to the number of datasets for which a method failed to execute (see Methods). Finally an Overall Composite Score was calculated for each method by averaging the four composite scores. Figure 2a summarizes the cell-type deconvolution performance of 21 methods across all 37 datasets, highlighting the composite scores, bivariate spatial metrics, non-spatial metrics, cell shape characterization metrics, and the rare cell-type metrics. Figure 2a ranks the methods according to their overall composite scores averaged over all 37 datasets.

**Figure 2:**
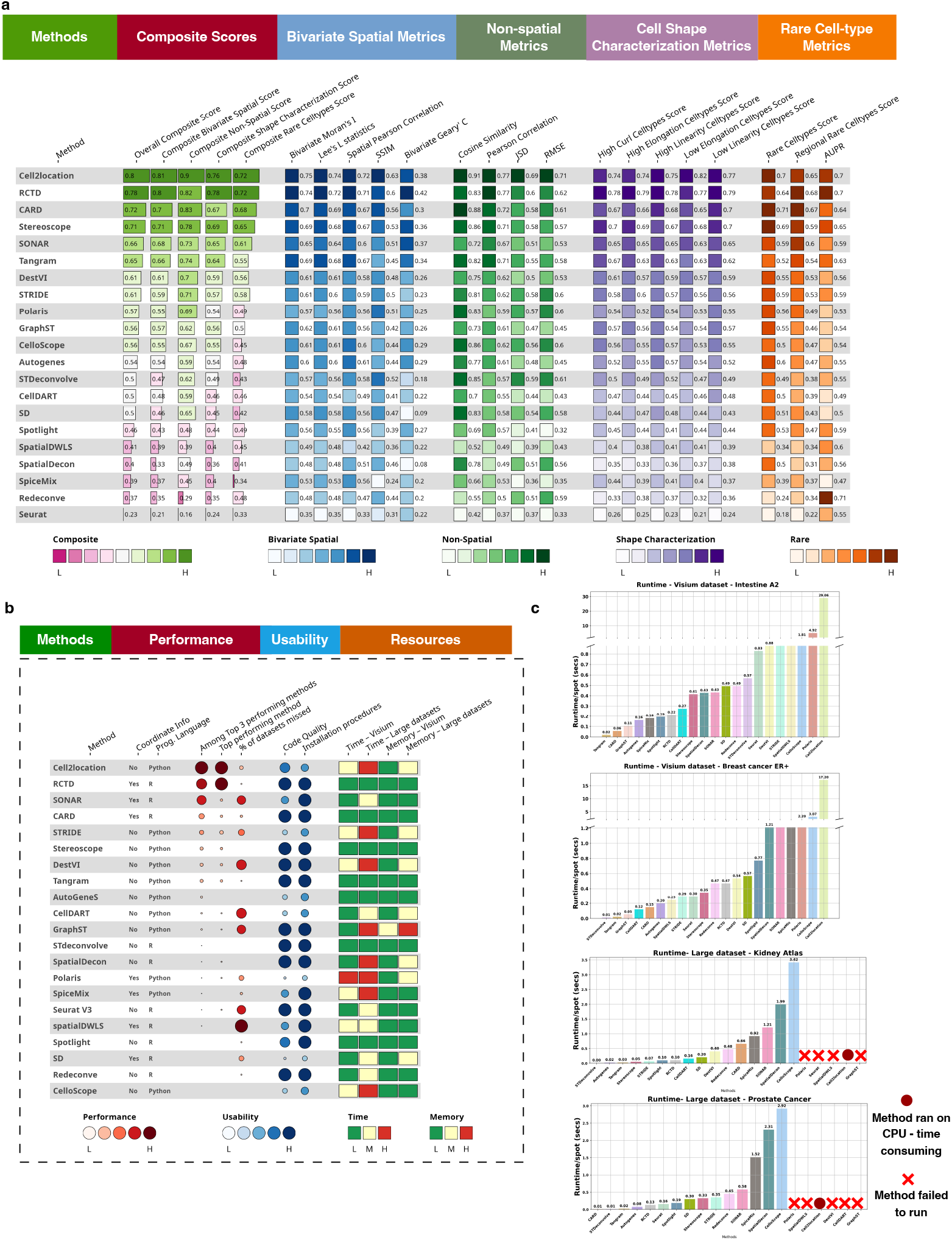
Comparison of overall performance of all spatial cell type deconvolution methods. a) Visualization of the spatial cell type deconvolution methods’ quantitative performance in terms of Composite scores (red), Bi-variate Spatial metrics (purple), Non-Spatial metrics (blue), Shape Characterization metrics (Orange) and Rare Cell-type metrics (Green). b) Qualitative evaluation of the spatial cell type deconvolution methods in terms of Performance(red), Usability (blue) and Resources (Orange). c) Runtime comparison of spatial cell type deconvolution methods for representative 10x Visium and Slide-seq V2 datasets.

Cell2location, RCTD, CARD, Stereoscope, and SONAR emerged as the top-performing methods based on the Overall Composite Score. The performance of the methods in terms of individual composite scores was consistent with that of the Overall Composite Score. Considering individual metrics, we observed that bivariate spatial metrics were more rigorous as compared to the non-spatial metrics. Particularly, within the bivariate spatial metrics, Bivariate Geary’s C and SSIM were more stringent compared to the other three metrics, with Bivariate Geary’s C being the most rigorous. RCTD, Cell2location, and SONAR ranked as the top three methods based on Bivariate Geary’s C. Across all cell shape characterization metrics, RCTD achieved the highest performance, both for individual cell-type shape patterns and for the Composite Shape Characterization Score. In terms of rare cell type metrics, Cell2location and CARD achieved the best Rare Cell types Score, while RCTD achieved the best Regional Rare Celltypes Score. We also observed that the AUPR metric had a different trend compared to the other two rare cell type metrics where some lower-ranked methods (e.g., Redeconve, SpatialDWLS) had disproportionately higher AUPR scores indicating that the lower-ranked methods were able to correctly predict the presence or absence of rare cell types but failed to estimate their correct proportion. Supplementary Figure 3b summarizes the performance of all methods without penalty, identifying Cell2location, SONAR, RCTD, DestVI, and CARD as the top performers. In this setting, SONAR ranked highly because no penalty was applied for datasets on which it failed to run, indicating that when SONAR executed successfully, it consistently performed among the top methods. Notably, Cell2location, RCTD, and SONAR most frequently ranked among the top three methods, doing so in 23, 19, and 17 datasets, respectively (Supplementary Figure 4a).

We next evaluated the performance of deconvolution methods in a tissue-specific manner, focusing first on brain datasets, where across 12 datasets, Cell2location, RCTD, and Tangram emerged as the top-performing methods for most composite scores, while CARD performed the best in detecting rare and regionally rare cell types (Supplementary Figure 5a). In the unpenalized analysis (Supplementary Figure 5b), Cell2location, DestVI, and CellDART ranked highest, indicating that DestVI and CellDART’s overall performances were negatively impacted by the penalty specifically due to their failure in execution for large datasets. Across brain datasets, CARD and Tangram most frequently appeared among the top three methods (6 and 5 of 12 datasets, respectively), whereas Cell2location consistently ranked first whenever it appeared in the top three (Supplementary Figure 4b), explaining its strong overall performance. Supplementary Figure 6a-f demonstrates the composite scores as well as the individual metrics for each brain dataset.

Across 13 cancer datasets, Cell2location, RCTD, and Stereoscope emerged as the top-performing methods. Performance across most metrics was consistent with the Overall Composite Score, except for Bivariate Geary’s C, for which RCTD, Cell2location, and SONAR achieved the highest values (Supplementary Figure 5c). In the unpenalized analysis (Supplementary Figure 5d), SONAR, Cell2location and RCTD emerged as the top performing methods highlighting SONAR’s strong performance in the datasets where it successfully ran. Across cancer datasets, Cell2location, RCTD, and SONAR most frequently appeared among the top three methods (10, 6 and 6 of 13 datasets, respectively) (Supplementary Figure 4c). Notably, STRIDE ranked among the top three methods in five datasets, outperforming RCTD and SONAR, with SONAR failing to run on four liver cancer datasets (Supplementary Figure 4c). Supplementary Figure 7a-g demonstrates the composite scores as well as the individual metrics for each cancer dataset.

Finally, for the datasets encompassing different organs other than brain (*n* = 12), Cell2location, RCTD and CARD emerged as the top-performing methods (Supplementary Figure 5e). In terms of Bivariate Geary’s C metric, RCTD, SONAR, and Stereoscope achieved the best performance. In the unpenalized analysis (Supplementary Figure 5f), SONAR, RCTD, and Cell2location emerged as the top performing methods which indicated SONAR’s superior deconvolution performance for the datasets where it successfully executed. Across organ datasets, Cell2location, RCTD, SONAR most frequently appeared among the top three methods (9, 9 and 7 out of 12 datasets, respectively) (Supplementary Figure 4d). Supplementary Figure 8a-h demonstrates the composite scores as well as the individual metrics for each organ dataset.

Considering performance across tissue types, Cell2location and RCTD consistently demonstrated strong and robust performance across diverse biological contexts, while SONAR achieved highly competitive accuracy when it successfully executed, particularly for the cancer and organ datasets. CARD and Tangram were effective for brain datasets, particularly for rare cell-type detection. Overall, Cell2location, RCTD, and SONAR emerged as the leading computational methods with high cell-type deconvolution accuracy.

We next evaluated the methods based on usability metrics, including the frequency of a method appearing in the top three, code quality, ease of installation, and computational resource requirements (Figure 2b). Usability was assessed based on code quality and ease of installation, with RCTD, SONAR, and CARD emerging as the top-ranked methods. Several other methods including Stereoscope, DestVI and Tangram also demonstrated good performance in terms of usability metrics. We further performed a qualitative assessment of each method’s time and memory requirements using three categories - low (L), medium (M), and high (H) - where L denotes runtimes under 30 minutes and CPU RAM usage under 50 GB; M corresponds to a runtime between 30 minutes and 3.5 hours with RAM usage between 50–200 GB; and H corresponds to runtimes exceeding 3.5 hours and memory usage above 200 GB.

We also conducted runtime experiments on two Visium datasets (Intestine A2 and ER+ Breast Cancer) and two Slide-seq V2 datasets (Prostate Cancer and Kidney Atlas) (Figure 2c). Among the four most accurate methods, Cell2location, RCTD, SONAR, and CARD, Cell2location exhibited the longest runtimes, whereas RCTD and CARD were the fastest. RCTD and CARD achieve high computational efficiency through their linear and NMF-based formulations, respectively. Cell2location, while computationally more demanding due to its hierarchical Bayesian modeling with variational inference, leverages this richer framework to deliver strong performance across multiple evaluation criteria. SONAR offers an intermediate trade-off, achieving lower runtimes than Cell2location with its Poisson–Gamma regression model, which is simpler to optimize. For large-scale datasets, many methods failed to run due to memory limitations or lack of convergence. The relative ordering of runtime remained consistent for large datasets, with RCTD and CARD being the fastest, followed by SONAR. Cell2location runtimes on large datasets are not directly comparable, as GPU memory constraints required CPU execution while other methods ran on GPU. Notably, although total runtime increases for large datasets (Figure 2c, Supplementary Figure 8i), per-spot runtime is effectively lower on large, higher-resolution datasets, which contain fewer cells per spot. For example, RCTD required 0.10-0.13 sec per spot on large datasets, compared to 0.20-0.47 sec on Visium datasets. This trend was observed across multiple methods, which exhibited near-zero per-spot runtimes for the large datasets.

### Technical variability and cell type diversity within datasets can impact deconvolution performance

We next evaluated if the deconvolution performance of the methods vary across the different spatial technologies. For this, we divided the datasets into three categories depending on the spatial technology. The first category (Visium) contained simulated Visium datasets, whereas the second category (large) contained simulated Slide-seq V2 and Visium HD datasets. The third category contained the datasets simulated using simulation strategy 2 using imaging-based datasets. These categories differed substantially in spatial resolution, sequencing depth, and scale, thereby providing complementary perspectives for evaluating the robustness and generalizability of deconvolution methods. Figure 3a demonstrates the deconvolution performance of the methods across datasets generated using three technology categories. The Visium category summarizes composite scores across 26 Visium datasets, the large-scale category across five Slide-seq V2 and two Visium HD datasets, and the image-based category across four simulated datasets generated based on imaging-based spatial datasets.

**Figure 3:**
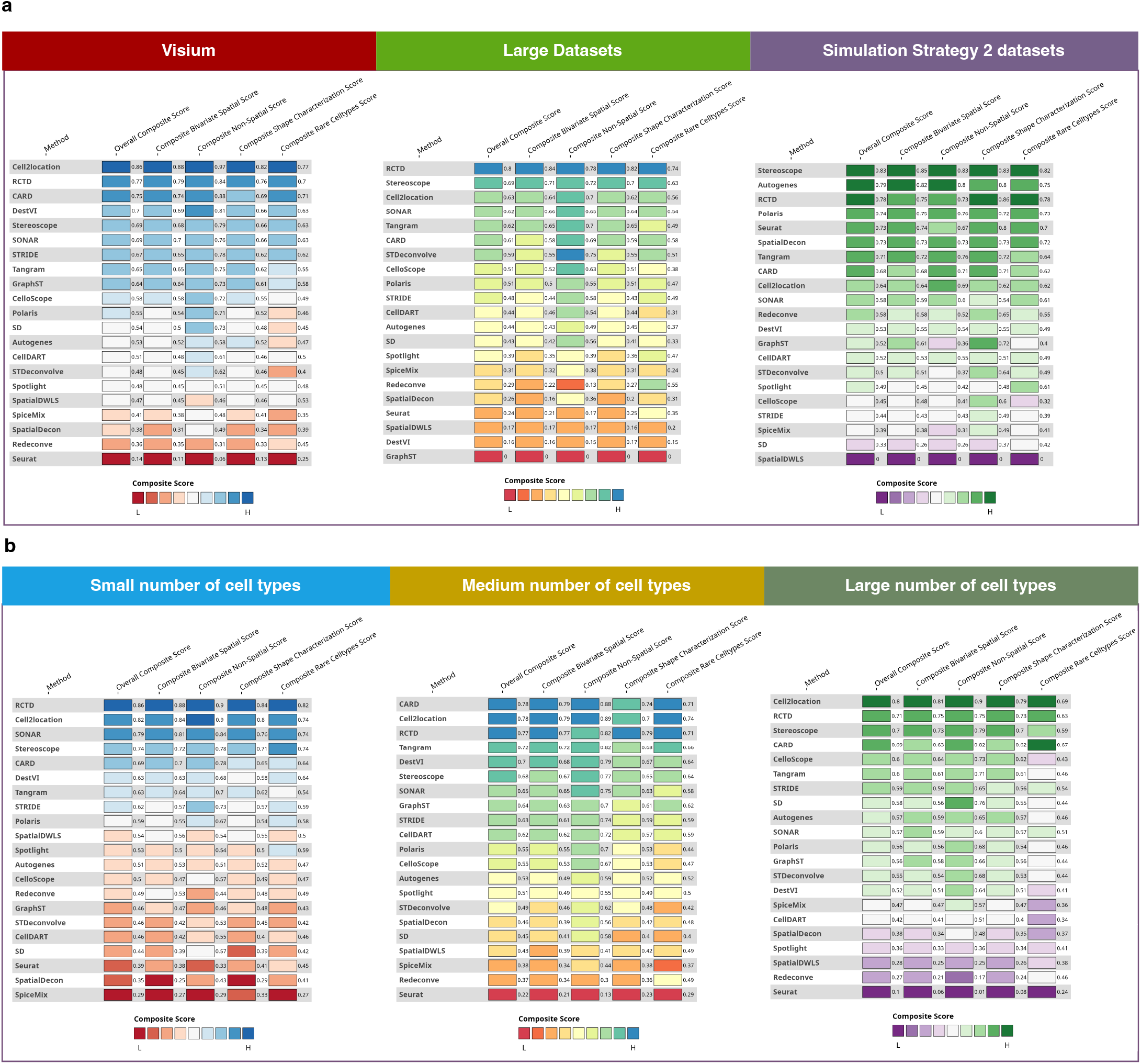
Comparison of performance of all spatial cell type deconvolution methods across spatial technologies and cell type diversity. a) Quantitative comparison of deconvolution methods in terms of Composite scores for 10x Visium datasets (red), Large datasets including Slide-seq V2 and Visium HD (green), and datasets generated using binning strategy on imaging-based datasets (purple) b) Quantitative comparison of deconvolution methods in terms of Composite scores for deconvolution tasks involving *<*= 12 cell types (blue), *>* 12 *<*= 21 cell types (yellow) and *>* 21 cell types (green).

For Visium datasets, we observed that Cell2location, RCTD, and CARD emerged as the top-performing methods based on Overall Composite Score. However, when usability was assessed by the frequency of appearance among the top three performers, Cell2location, RCTD, and SONAR ranked highest, appearing in the top three for 19, 14, and 12 datasets, respectively (Supplementary Figure 4a, 9a) mirroring the overall performance as described in the previous section. SONAR’s reduced Overall Composite Score reflects penalties incurred from failed executions on a subset of datasets, despite being the top-performing method in the majority of datasets where it ran successfully. Similarly, although CARD appeared less frequently among the top three performers for individual datasets, its consistently strong performance across all datasets placed it among the top three in the final aggregated ranking.

For large-scale datasets, RCTD, Stereoscope, and Cell2location emerged as the top three methods according to the Overall Composite Score, and performance across individual composite scores was consistent with the overall ranking. However, based on the performance for individual datasets, RCTD, SONAR and Cell2location ranked in top three for 5, 4, 4 datasets, respectively (Supplementary Figures 9b). Notably, several methods failed to execute for different large datasets, including Cell2location and SONAR failing to execute for 3 and 2 datasets respectively, which negatively affected their Overall Composite Scores. In contrast, Stereoscope, Tangram, and RCTD successfully ran on nearly all large datasets, incurred no penalties, and consequently ranked among the top-performing methods in this category. Notably, we observed an overall reduction in composite scores for top-performing methods on large-scale datasets compared with Visium datasets, highlighting the increased difficulty of accurate deconvolution in large-scale spatial transcriptomics datasets (Supplementary Figures 9c).

Finally, for the imaging-based datasets simulated using simulation strategy 2, Stereoscope, Autogenes, and RCTD emerged as the top-performing methods based on Overall Composite Score, with Autogenes and Stereoscope ranking among the top three methods for three and two datasets, respectively (Supplementary Figs. 9d), while RCTD achieved a top ranking for Overall Composite Score owing to its stable performance across all datasets. In addition, SONAR and Polaris demonstrated strong performance in the datasets on which they successfully executed (Supplementary Figures 6f, 8h). Given that a distinct set of methods emerged as top performers for image-based datasets compared with Visium and large-scale datasets, we further characterized the deconvolution task in image-based data by analyzing the distribution of contributing cell types per spot. We observed that in datasets generated using simulation strategy 1, a large number of cell types contributed to individual spots (Supplementary Figure 10a). In contrast, in datasets generated using simulation strategy 2 from image-based datasets, the majority of spots were composed of contributions from only one to two cell types, despite the presence of seven to ten cell types within the broader tissue region (Supplementary Figure 10b). These differences indicate a fundamentally distinct deconvolution regime characterized by one or two dominant cell types per spot, under which simpler models such as Stereoscope and Autogenes perform favorably, whereas more complex probabilistic methods, such as Cell2location, which tend to assign low but non-zero contributions to multiple cell types, are disadvantaged in this setting.

We next examined whether the performance of the methods varies with the number of cell types harbored in the dataset by stratifying datasets into three groups: Small (0–12 cell types), Medium (13–21 cell types), and Large (≥22 cell types) (Supplementary Figure 11a). Figure 3b summarizes composite scores for datasets with Small (11 datasets), Medium (12 datasets), and Large number of cell types (14 datasets). The overall top performing methods, RCTD and Cell2location exhibited consistent performance across all three categories, indicating robustness to increasing cell-type diversity (Supplementary Figure 11b-d). In contrast, while SONAR performed strongly on datasets with a small number of cell types, execution failures on three medium and five large cell-type datasets adversely affected its Overall Composite Score. CARD and Stereoscope showed stable performance across all complexity levels (Supplementary Figure 11b, e), with CARD achieving the highest Overall Composite Score in the medium cell-type category, largely driven by its strong performance in DLPFC datasets that dominate this group. Overall, we observed a gradual decline in composite scores with increasing cell-type complexity, reflecting the greater difficulty of deconvolution in more heterogeneous settings. Among the top-performing methods, CARD and Cell2location exhibited greater robustness to increasing cell-type diversity, whereas the performance of RCTD, SONAR, and Stereoscope declined as cell-type diversity increased (Supplementary Figure 11e).

### Deconvolution performance in detecting rare cell types and recovering spatial abundance patterns across tissues

Accurate detection of rare cell types is a critical aspect of cell-type deconvolution because these populations often play functional roles in tissue organization, disease progression, and therapeutic response. In cancer and other complex tissues, rare immune subsets or transient malignant states can drive invasion, immune evasion, and treatment resistance, and their occurrence is frequently leveraged to construct prognostic models of patient outcomes (67; 68). Consequently, we next focused on evaluating the deconvolution methods based on their ability to accurately recover rare cell types. In our analyses, we considered two definitions of rare cell types: (i) globally rare cell types, defined as those with low overall abundance (mean cell-type proportion values in bottom 10th percentile), and (ii) regional rare cell types, defined as those with mean cell-type proportion values in the bottom 30th percentile and exhibiting bimodal or multimodal spatial distributions, indicative of abundance being restricted to specific spatial regions. To evaluate the performance in handling these cell types, we introduced two scores, Rare Cell types Score and Regional Rare Cell Types Score, which summarizes the bivariate spatial metrics for globally rare and regional rare cell types respectively (see Methods for details). Figure 4a illustrates the identification of globally rare (microglia) and regional rare cell types (OPC, microglia, INSV2C, and L5 6 CC) from the DLPFC 151508 dataset. Supplementary Figure 12 demonstrates the density plots of regional rare cell types identified for each dataset (Supplementary Table 3).

**Figure 4:**
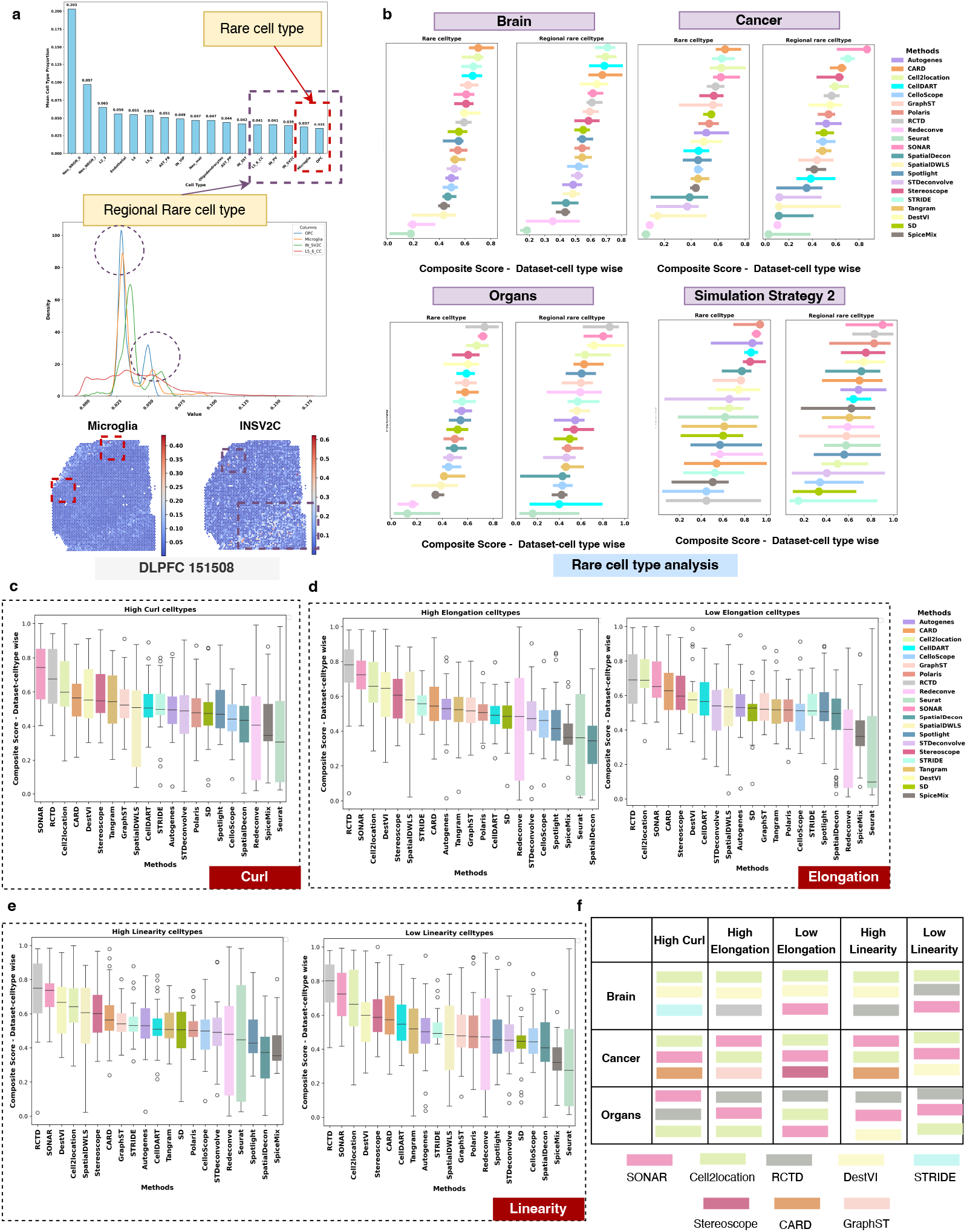
Performance comparison of deconvolution methods in detecting rare cell types and recovering spatial abundance patterns across tissues. a) Schematic illustration defining globally rare and regionally rare cell types for a representative dataset (DLPFC 151508): (top) bar plot of mean cell-type proportions across all cell types, with the rare cell types highlighted; (center) density plot illustrating regionally rare cell types with bimodal distributions; and (bottom) spatial cell-type abundance plots for rare cell types. b) Comparison of methods for rare and regionally rare cell type recovery across brain, cancer, organ tissues and simulation strategy 2 datasets. Each point represents the median value of bivariate composite score for the target cell types, and horizontal bars correspond to the 95% confidence intervals. Comparison of methods based on bivariate composite scores for (c) high curl cell types, (d) high and low elongation cell types, and (e) high and low linearity cell types. f) Summary table highlighting the top three performing methods for high curl, high elongation, low elongation, high linearity, and low linearity cell types across tissue categories, i.e. Brain, Cancer, and Organs. Each box-and-whisker plot presents the distribution of scores across datasets and cell types within a category. The box represents the interquartile range (IQR), spanning from the 25th to the 75th percentile, with the median indicated inside. The whiskers extend to the minimum and maximum values within 1.5 times the IQR, while outliers are shown as black circles.

Figure 4b summarizes the performance of all methods for the deconvolution of globally rare and regional rare cell types across brain, cancer, and organs datasets as well as datasets simulated using simulation strategy 2. For brain datasets, STRIDE, Cell2location, CellDART, and CARD achieved the best performance for both globally rare and regionally rare cell types. For cancer datasets, SONAR, STRIDE, and CARD achieved the best performance, whereas RCTD, SONAR, and DestVI performed best on organ datasets. In datasets generated using simulation strategy 2, SONAR, Polaris, and Stereoscope demonstrated strong performance. Among the overall top-performing methods, SONAR consistently performed well in detecting both rare and regionally rare cell types across tissues and technologies. STRIDE also emerged as a strong performer, particularly in brain and cancer tissues. CARD and DestVI additionally demonstrated robust performance for rare cell-type prediction across tissue categories. Moreover, the composite bivariate score heatmaps for the datasets generated using simulation strategy 2 demonstrate that POLARIS exhibited less variation in performance across cell types, including rare cell types, explaining its strong performance for datasets in this category.

Next, we assessed the performance of each method using cell shape characterization metrics, providing information on each method’s ability to recover cell types exhibiting prominent spatial abundance patterns, as quantified by curl, elongation, and linearity shape metrics (69) (see Methods for details). Accurate recovery of such spatial abundance patterns is critical, as many important cell types, including invasive tumor fronts, vascular-associated cells, and immune niches, exhibit distinct and characteristic spatial organization (70). Based on these metrics, we defined five categories to characterize cell types according to the geometry of their spatial abundance patterns. Cell types with high curl exhibit round or curvilinear structures, whereas those with high linearity consist of large, well-defined linear patterns. In contrast, low linearity corresponds to cell types with small or less prominent linear shapes. High elongation denotes extended spatial patterns with a large distance between the two extremities of the structure, while low elongation characterizes compact or weakly structured patterns.

Figures 4c-e summarize the composite scores of all methods across all datasets for cell types exhibiting high curl, high elongation, high linearity, low elongation, and low linearity Supplementary Table 4-6. Across all prominent cell-type shape patterns, the top-performing methods, RCTD, SONAR, and Cell2location, consistently demonstrated strong performance. We further examined performance stratified by tissue type (Supplementary Figure 13), with Figure 4f summarizing the top-performing methods for each tissue category. For brain datasets, Cell2location and DestVI consistently performed best across all cell-type shape patterns, with the exception of low-linearity cell types (Supplementary Figure 13a). In cancer datasets, SONAR and Cell2location alternated as leading methods (Supplementary Figure 13b). For organ datasets, RCTD emerged as the best performer in most cases, followed by SONAR and Cell2location. Notably, DestVI also performed well in organ datasets, ranking just below these top methods, highlighting its strength in recovering well-defined spatial patterns (Supplementary Figure 13c).

### Evaluating deconvolution accuracy for major cell types across tissue types

Tissue-specific architectural constraints, including laminar organization in the brain or heterogeneous tumor microenvironments in cancer, can differentially influence the deconvolution of certain cell types from spatial transcriptomic datasets (46; 70). Next, we compared the deconvolution accuracy for different cell types within the same tissue to identify the biological determinants of accurate cell type deconvolution. For that, we analyzed the composite scores of top-performing methods across major cell types present in multiple datasets of the same tissue type to identify the most accurately predicted and the most challenging cell types (Supplementary Table 7-9). In brain tissues, oligodendrocytes and inhibitory neurons achieved the highest composite scores, whereas macrophages were the most challenging to predict (Figure 5a, Supplementary Figure 14a). To determine whether deconvolution accuracy of a cell type was associated with cell-type abundance or prominent spatial structure, we calculated the average cell-type proportion per spot and Moran’s I statistic for each major cell type across datasets. We observed that among cell types exceeding a minimum abundance threshold, deconvolution accuracy was strongly correlated with the degree of spatial localization, as observed for oligodendrocytes, inhibitory neurons, and astrocytes (Supplementary Figure 14b-d). Notably, although excitatory neurons exhibited higher average abundance per spot, the presence of multiple transcriptionally similar subtypes reduced their separability, thereby limiting deconvolution precision (Supplementary Figure 14e). We further examined whether the number of single cells per cell type in the reference dataset influenced deconvolution accuracy, using DLPFC 151673 as a representative case. Interestingly, OPCs, despite having a large number of single cells (~ 5000), ranked among the lowest-performing cell types, likely due to their low spatial abundance ~ 0.05). In contrast, endothelial cells, with fewer single cells ~ 1, 500) but higher spatial abundance ~ 0.2), exhibited better deconvolution performance (Supplementary Figure 14f).

**Figure 5:**
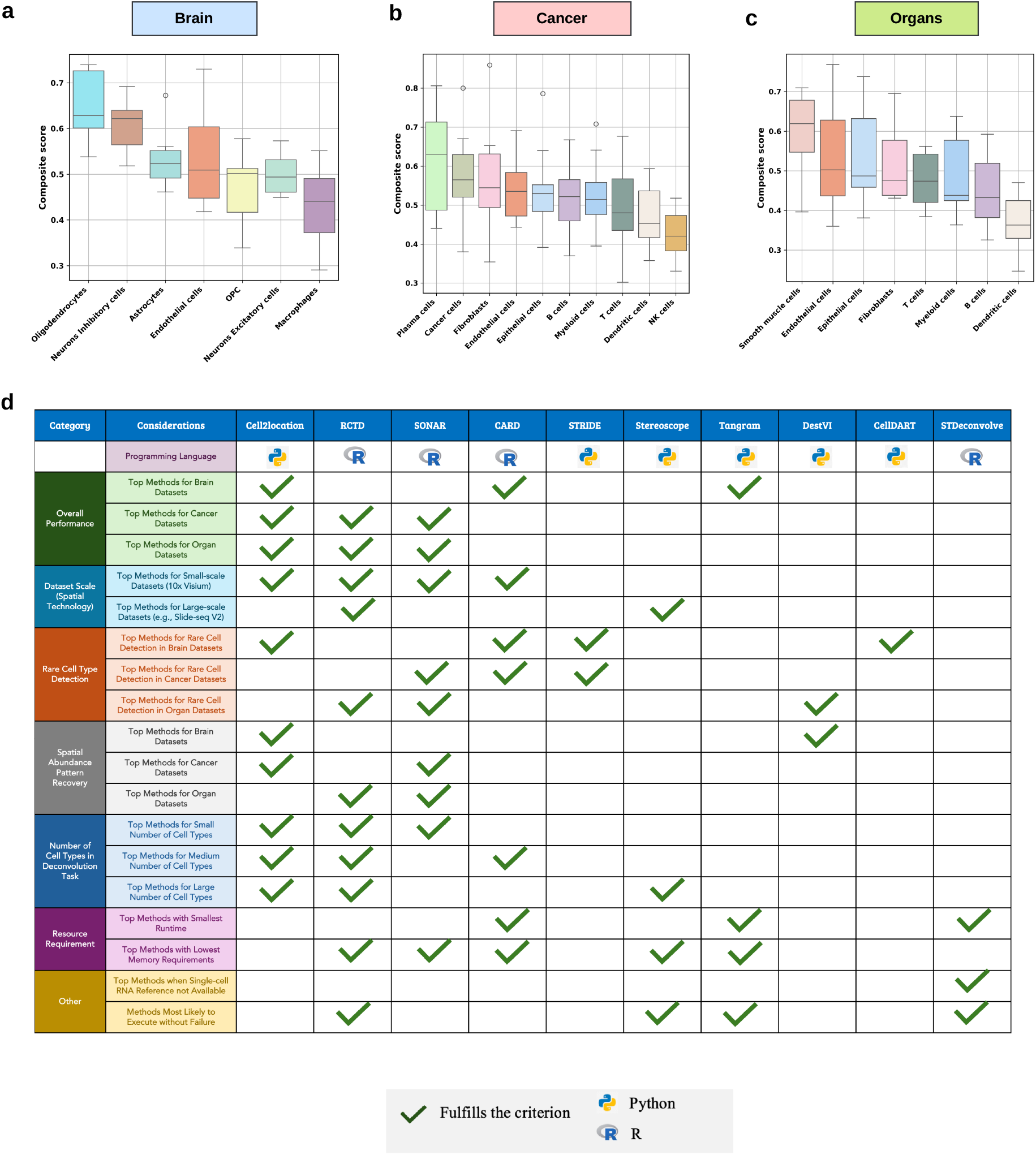
Deconvolution accuracy for major cell types across tissue types and guidelines for selecting cell-type deconvolution methods. Comparison of deconvolution accuracy across major cell types in a) brain (oligodendrocytes, inhibitory neurons, astrocytes, endothelial cells, OPCs, excitatory neurons, and macrophages), b) cancer (plasma cells, cancer cells, fibroblasts, endothelial cells, B cells, myeloid cells, T cells, dendritic cells, and NK cells), and c) organ (smooth muscle cells, endothelial cells, epithelial cells, fibroblasts, T cells, myeloid cells, B cells, and dendritic cells) datasets. Each box-and-whisker plot illustrates the distribution of bi-variate composite scores across datasets within a category for a specific cell type. The box represents the interquartile range (IQR), spanning from the 25th to the 75th percentile, with the median indicated inside. The whiskers extend to the minimum and maximum values within 1.5 times the IQR, while outliers are shown as black circles. d) Table of criteria for selecting cell-type deconvolution methods and the corresponding methods that satisfy each criterion. Considerations are divided into the seven broad categories - Overall Performance, Dataset Scale (Spatial Technology), Rare Cell Type Detection, Spatial Abundance Pattern Recovery, Number of Cell Types in Deconvolution Task, Resource Requirements and Other. Python and R symbols indicate the primary language in which the method is programmed and used. A green tick indicates which methods fulfill each criterion.

In cancer datasets, top performing methods achieved the highest composite score for plasma cells, cancer cells and fibroblasts, whereas dendritic cells and NK cells were among the most challenging to predict accurately (Figure 5b, Supplementary Figure 14g). We observed a strong association between deconvolution performance and spatial structure, the cell types with higher Moran’s I values demonstrated good deconvolution performance (Supplementary Figure 14h-i). Plasma cells exhibited strong performance due to their high regional prevalence, despite having a low mean cell-type proportion (Supplementary Figure 14h-j). Using NK cells in a kidney cancer dataset, we further showed that spatial abundance, rather than the number of cells in the reference data, primarily determines deconvolution performance (Supplementary Figure 14k). In contrast, cancer cells exhibited strong performance due to their high regional prevalence and overall proportion within the tissue (Supplementary Figure 14k). Finally, in organ datasets, smooth muscle cells, endothelial cells, epithelial cells and fibroblasts achieved the highest composite scores, whereas B cells and dendritic cells were among the most difficult to predict, consistent with earlier observations (Figure 5c, Supplementary Figure 14l). For organs datasets too, higher deconvolution performance of a cell type was associated with higher spatial abundance and strong spatial structure (Supplementary Figure 14m-n). For example, mesothelial cells in the Intestine dataset, despite having a relatively low average proportion (0.08), achieved the highest composite score due to their broad and coherent spatial distribution across the tissue section. In contrast, moderately performing cell types, such as certain immune populations, were reasonably well captured when their contributions were concentrated with high magnitude in localized regions (Supplementary Figure 14o).

### Practical guidelines for selecting spatial deconvolution methods

To facilitate method selection across diverse experimental settings, we summarized our benchmarking results into a guideline framework that maps key analytical considerations to top-performing methods (Figure 5d). Overall, Cell2location, RCTD, and SONAR consistently emerged as top performers across tissue types, with CARD and Tangram showing strong performance for brain datasets. Method performance varied substantially with dataset scale and technology. For standard 10x Visium datasets, Cell2location, RCTD, SONAR, and CARD performed best, whereas for large-scale datasets (e.g., Slide-seq V2, Visium HD), RCTD and Stereoscope were more reliable due to their scalability. In predicting rare cell type proportions, SONAR, STRIDE, and CARD demonstrated the most consistent performance across tissue types, with SONAR performing best for cancer and organ datasets, while CARD and STRIDE showed strong performance for brain and cancer datasets. Notably, top overall performers such as Cell2location were less effective at accurately capturing rare cell types. For recovering cell types with prominent spatial abundance patterns, Cell2location and DestVI performed best for brain datasets, Cell2location and SONAR for cancer datasets, and RCTD and SONAR for organ datasets. Performance was also influenced by cell-type complexity: Cell2location and RCTD were robust across varying numbers of cell types, while Stereoscope performed well in high-complexity settings. From a computational perspective, CARD, Tangram, and STDeconvolve exhibited the shortest runtimes, whereas RCTD, SONAR, CARD, Stereoscope, and Tangram had the lowest memory usage. Finally, for scenarios where single-cell references are unavailable, STDeconvolve remains a key reference-free option, while RCTD, Stereoscope, Tangram, and STDeconvolve showed the highest reliability in terms of successful execution across datasets. Collectively, these guidelines highlight that no single method is universally optimal, and that method selection should be tailored to the biological question, dataset scale, and computational constraints.

### Benchmarking of spatial domain detection methods

Finally, we performed a comprehensive evaluation of 18 state-of-the-art spatial domain detection methods employing different computational techniques - Graph deep learning-based methods: SpaSRL (71), SCANIT (72), and SpatialPCA (73); GCN and contrastive learning-based methods: DeepST (74), GraphST (3), SpaceFlow (75) and CCST (76); Spatial kernel-based method: Banksy (77); hidden Markov random field (HMRF)-based method: Giotto (78), DR-SC (79), and iSCMEB (80); Bayesian model-based method: BayesSpace (81), PRECAST (82), BayesCafe (83), and BASS (84); graph attention network-based methods: STAGATE (85) and PROST (86); and single-cell reference-based method: IRIS (87) (Figure 6a, Supplementary Table 10).

**Figure 6:**
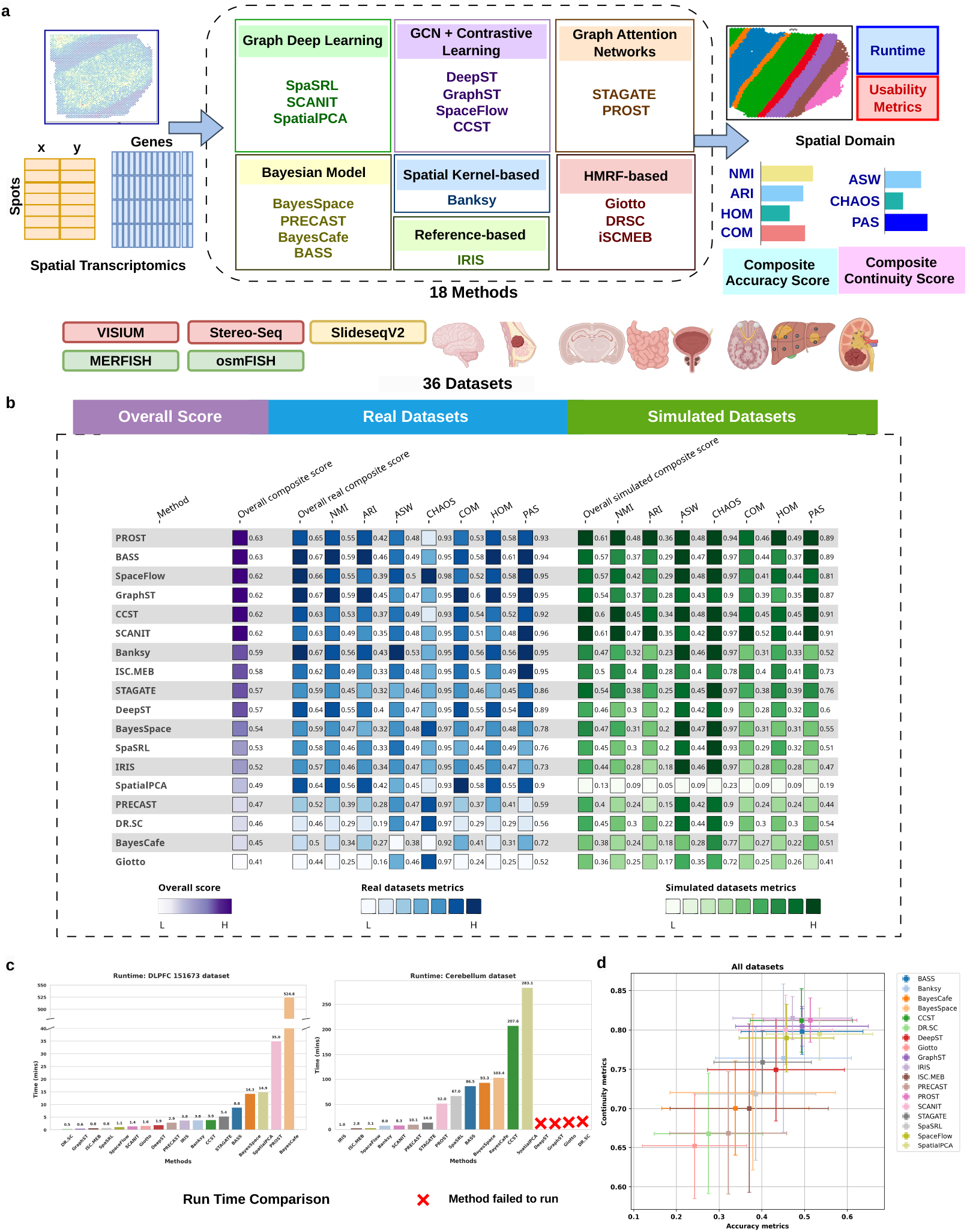
Overview of the workflow and results for domain detection benchmarking. a) Schematic representation of the benchmarking workflow used for evaluating the performance of 18 domain detection methods (i.e. Banksy, BayesCafe, BayesSpace, BASS, CCST, DeepST, DR.SC, Giotto, GraphST, IRIS, ISC.MEB, PRECAST, PROST, SCANIT, SpaSRL, SpaceFlow, SpatialPCA and STAGATE), categorized by their data requirements and computational strategies. The benchmarking was performed on 36 datasets, spanning cancer, brain, and organ tissues, and generated using various spatial technologies, including 10x 10x Visium, Slide-seq V2, Stereo-seq and imaging-based techniques (MERFISH, osmFISH). The benchmarking results were evaluated using a set of four accuracy metrics and three continuity metrics, along with their respective composite scores. b) Visualization of the methods’ quantitative performance in terms of Composite scores and individual metrics for real and simulated datasets. c) Comparison of methods in terms of run time for Visium and Slide-seq V2 datasets. d) Quantitative assessment of 18 spatial domain detection methods in terms of Composite continuity score and Composite accuracy score for 36 datasets. Data are presented as mean values ± standard deviation.

We benchmarked 18 spatial domain detection methods across 36 datasets, comprising 21 real datasets and 15 simulated datasets generated from 5 different spatial technologies: 10x Visium, Slide-seq V2, MERFISH, osmFISH, and Stereo-seq, also providing a comprehensive coverage across diverse tissue types. The real datasets used in our benchmarking include 12 10x Visium DLPFC datasets (46; 47), Stereo-seq mouse embryo datasets (E9.5 and E14.5) (88; 89), 10x Visium mouse breast cancer dataset (3; 90), 5 MERFISH mouse hypothalamus datasets (48), and osmFISH mouse somatosensory cortex dataset (91; 92). The simulated datasets used in our benchmarking include 10x Visium kidney cancer dataset (49), 10x Visium breast cancer dataset (93), 10x Visium liver cancer dataset (51; 52), 10x Visium human breast dataset (50), 10x Visium chicken heart dataset (57), 10x Visium intestine dataset (58), Slideseq V2 prostate cancer dataset (53) and Slideseq V2 cerebellum dataset (24)(Figure 6a, Supplementary Table 11). Thus, in addition to tissue-type diversity, our dataset repository captures a broad spectrum of variation, encompassing 23 distinct spatial patterns, three large datasets, five cancer samples, and considerable heterogeneity in the number of spatial domains across samples (Figure 6a, Supplementary Figure 15). We comprehensively evaluated the methods by comparing their predicted spatial domains to the ground truth domain annotations using two categories of evaluation metrics: (1) accuracy metrics and (2) continuity metrics. The accuracy metrics included Normalized Mutual Information (NMI) (94), Adjusted Rand Index (ARI) (95), Homogeneity Score (HOM) (96), and Completeness Score (COM) (96). The continuity metrics comprised Average Silhouette Width (ASW) (97), Clustering Homogeneity Across Ordered Samples (CHAOS) (98; 99), and Percentage of Abnormal Spots (PAS) (100). These metrics evaluate the extent to which predicted spatial domains exhibit spatial coherence and clear inter-domain boundaries. Continuity metrics also assess the quality of unsupervised clustering by measuring the preservation of spatial organization. An overall composite score was computed by averaging the scores from both metric categories (20) (Figure 6a).

Figure 6b summarizes the overall performance of the 18 domain detection methods based on three composite scores: (1) Overall composite score, (2) Overall real composite score, and (3) Overall simulated composite score, which respectively represent performance across all datasets, real datasets, and simulated datasets. As in the cell type-deconvolution benchmarking, we applied a slope-based penalization strategy to account for incomplete method execution across datasets. Specifically, the composite score was adjusted using a gradually increasing penalty proportional to the number of datasets on which a method failed to execute (see Methods) (Supplementary Figure 16). According to the Overall composite score, PROST, BASS, SpaceFlow, GraphST, CCST, and SCANIT were identified as the top-performing methods, with only marginal differences observed among them (Fig. 6b, Supplementary Figure 17b). For real datasets, the top-performing methods were BASS, GraphST, Banksy, SpaceFlow, and PROST. In contrast, PROST, SCANIT, CCST, BASS, and SpaceFlow achieved the highest rankings on simulated datasets. Performance across individual metrics largely aligned with Overall Composite Score rankings, except for CHAOS and PAS (Supplementary Figure 17c-i). Supplementary Figure 17a presents usability metrics for each method, including programming language, required input parameters (cluster number or resolution), frequency of top-three rankings overall and across Visium, image-based, and large datasets, the number of datasets on which the method failed to run, and runtime category (Low ≥ 30 min; Medium 30 − 100 min; High *>* 100 min). We found that PROST, GraphST, and CCST consistently ranked among the top three methods across most datasets, a pattern that was also observed for Visium datasets. In contrast, for image-based datasets, Banksy, BASS, and SpaceFlow emerged as the top-performing methods. For large datasets, SpaceFlow ranked highest, followed by PROST and SCANIT. Most methods showed low (L) runtime, whereas PROST, BASS, BayesSpace, and spaSRL exhibited medium (M) runtime, and CCST, SpatialPCA, and BayesCafe demonstrated high (H) runtime. Figure 6c presents runtime comparisons for smaller Visium datasets (DLPFC 151673) and larger datasets such as Cerebellum (*>* 30, 000 cells). DRSC, GraphST, and iSCMEB were the fastest on Visium datasets, while IRIS, iSCMEB, and SpaceFlow performed most efficiently on Slide-seq datasets.

Figure 6d compares the performance of all methods across accuracy and continuity, along with their variability represented by the error bar. PROST, CCST, and GraphST ranked highest across both metrics. Notably, these top-performing methods also exhibited smaller error bars than lower-ranked methods, indicating more consistent performance. Supplementary Figure 18a further shows that PROST, BANKSY, and GraphST performed best on real datasets, whereas IRIS, PROST, and CCST achieved the highest performance on simulated datasets. Although SpatialPCA performed well on real datasets, it failed to execute on many simulated and large datasets; consequently, its simulated-data performance appears substantially lower, resulting in reduced composite scores under that setting. Supplementary Figure 18b demonstrates the performance of domain detection methods on brain, cancer, and organ datasets. Across brain datasets, many methods performed well, reflecting their ability to capture the structured, layered architecture characteristic of brain tissue. In contrast, performance was generally reduced for cancer datasets, where spatial organization is irregular due to tumor heterogeneity; however, CCST, PROST, and IRIS remained top-performing methods. This difference likely reflects modeling assumptions in certain approaches, for instance, methods such as BASS that rely on spatial homogeneity assumptions may be less compatible with heterogeneous tumor architecture. For organ datasets, IRIS and PROST achieved the highest overall performance. Supplementary Figure 18c summarizes performance across technology types, Visium, large datasets, and image-based datasets. PROST, IRIS, CCST, SpaceFlow, and GraphST achieved the highest performance on Visium datasets. Evaluation on large datasets was limited due to memory constraints; however, PROST consistently ranked highest, followed by SpaceFlow. For image-based datasets, BASS and Banksy performed best, with SpaceFlow and PROST ranking next.

Qualitative analyses largely mirrored composite score rankings (Supplementary Figure 19). GraphST, DeepST, and BASS performed best for discontinuous brain regions, whereas PROST, GraphST, CCST, and DeepST were most accurate overall. For complex embryo datasets, STA-GATE, IRIS, PRECAST, BayesSpace, and DeepST detected fine structures most reliably, while SpaceFlow performed best on large samples. In MERFISH data, BASS, Banksy, and PROST showed the strongest layer delineation, whereas GraphST and STAGATE underperformed. For simulated datasets, PROST, IRIS, GraphST, CCST, SCANIT, and SpaceFlow inferred spatial domains with high accuracy across cancer datasets. PROST and GraphST most accurately captured complex organ spatial structures, while SpaceFlow, PROST, and IRIS best resolved intricate patterns in prostate cancer and cerebellum samples.

## Discussion

Spatial transcriptomics has transformed our ability to study gene expression within its native tissue context, yet the rapid development of computational methods for cell-type deconvolution and spatial domain detection has outpaced efforts to systematically evaluate their performance (14). Here, we presented spDDB, a comprehensive and robust benchmarking framework that addresses critical gaps in prior evaluations by substantially expanding the diversity of datasets, methods, evaluation metrics, and biological contexts examined. Our deconvolution benchmarking offers several key advances: a curated repository of 37 datasets spanning brain, cancer, and organ tissues across multiple species and spatial technologies; a novel graph attention autoencoder-based simulator (SynthST) for generating realistic synthetic spatial transcriptomics data; a suite of spatially informed evaluation metrics including a novel bivariate Geary’s C metric; and systematic evaluation of 21 deconvolution methods with practical guidelines for end users.

Our extensive benchmarking of spatial cell type deconvolution methods identified Cell2location, RCTD, and SONAR as the top-performing methods across diverse tissue types, spatial technologies, and evaluation criteria. Cell2location’s hierarchical Bayesian framework delivered strong performance across most contexts despite higher computational cost, while RCTD achieved robust accuracy with the fastest runtimes, making it well suited for large-scale datasets, and SONAR showed excellent predictive accuracy when it ran successfully, although execution failures on multiple datasets reduced its overall reliability. Our technology-stratified analyses showed that deconvolution performance depends strongly on platform characteristics: Bayesian methods such as Cell2location and CARD performed best on Visium datasets, linear models like RCTD and Stereoscope scaled better to large Slide-seq V2 and Visium HD datasets, and simpler models such as Stereoscope and Autogenes were favored in imaging-based datasets where spots were dominated by one or two cell types, underscoring the need for platform-aware method selection.

A methodological contribution of this study is the development of SynthST, a graph attention autoencoder–based simulation framework designed to generate realistic spatial transcriptomic datasets. Unlike previous simulation approaches that rely on simplified or uniform spatial distributions, SynthST captures complex spatial structures by integrating multiple deconvolution outputs through a multi-view learning strategy. This approach enables the generation of synthetic datasets that better reflect the spatial heterogeneity observed in real tissues. In addition, we introduced spatially informed evaluation metrics that extend beyond conventional global measures such as Pearson correlation and RMSE. Our bivariate spatial metrics including the novel bivariate Geary’s C enable stringent assessment of local spatial agreement, while shape characterization and rare cell-type metrics provide a multidimensional evaluation of method performance, capturing aspects of deconvolution quality, such as the recovery of spatially structured rare populations and morphologically distinct abundance patterns, that are biologically meaningful but have been overlooked in prior benchmarks.

Our analyses revealed a discrepancy between overall deconvolution accuracy and rare cell-type recovery: although Cell2location achieved the highest overall scores, SONAR, STRIDE, and CARD more effectively captured rare and regionally rare populations, suggesting that methods optimized for global accuracy may dilute low-abundance signals and studies focused on rare cell-type identification may benefit from different approaches than those optimizing for global accuracy. Given the biological importance of rare cellular states, improving their detection and quantification will remain a critical area for future methodological development. Our cell-type-specific analyses across tissue categories revealed that deconvolution accuracy is primarily determined by the spatial abundance and degree of spatial localization of a given cell type, rather than by its representation in the reference single-cell dataset. Cell types exhibiting strong spatial autocorrelation (e.g., oligodendrocytes in brain or Plasma cells in cancer) were consistently predicted with higher accuracy. Conversely, transcriptionally similar subtypes, such as excitatory neuron subtypes in brain datasets, were more challenging to deconvolve, reflecting the difficulty of separating closely related cell identities from bulk spot-level expression profiles.

Based on our cell-type deconvolution benchmarking results, we identify several directions for future method development. First, there is a clear need for more reference-free deconvolution methods, as only a few currently exist despite their importance in settings lacking high-quality single-cell references. Second, dedicated approaches are required to improve the detection and quantification of spatially rare cell types, which remain challenging for most existing tools. Integrating spatial transcriptomics with complementary modalities, such as histological images, chromatin accessibility, or protein measurements, represents a promising direction for improving deconvolution accuracy and resolving finer cellular heterogeneity. Furthermore, as spatial transcriptomics datasets grow in scale to encompass entire organs or longitudinal time series, the scalability and efficiency of deconvolution methods will become increasingly critical. The development of methods that can adaptively adjust their complexity based on local tissue architecture, applying simpler models in homogeneous regions and more sophisticated approaches in complex microenvironments, could address the observation that no single method excels universally.

In domain detection benchmarking, PROST, BASS, SpaceFlow, GraphST, and CCST emerged as the top-performing methods, though with relatively small margins separating them. Performance patterns varied considerably across tissue types and technologies. For brain datasets, which exhibit well-defined laminar organization, most methods performed comparably well, reflecting the relative ease of detecting structured, layered architectures. In contrast, cancer datasets posed greater challenges due to the irregular and heterogeneous nature of tumor spatial organization; here, CCST, PROST, and IRIS demonstrated more robust performance, suggesting that these methods are better equipped to handle spatial heterogeneity. For large-scale datasets, scalability emerged as a critical differentiator, with SpaceFlow and PROST maintaining strong performance where many other methods failed to execute. Based on these findings, we provide a practical guideline summarizing the recommended domain detection methods across diverse experimental considerations, including tissue type, spatial technology, dataset scale, and computational efficiency (Extended Figure 2), to assist users in selecting the most suitable approach for their specific analytical context.

Our domain detection benchmarking reveals several priorities for future methodological development. A key unmet need is the development of methods capable of handling tissues with irregular or poorly defined spatial organization (e.g., cancer): approaches relying on spatial homogeneity priors perform well in structured tissues but tend to impose artificial domains in heterogeneous cancers, highlighting the need for models that can adaptively modulate spatial regularization, enforcing coherence where appropriate while allowing sharp, irregular, or diffuse boundaries where required by the underlying biology. Future methods can also focus on the estimation of number of domains as a data-driven model selection problem, using hierarchical, multi-resolution, or nonparametric frameworks, rather than requiring users to pre-specify the number of domains or resolution parameter despite its inherent biological ambiguity and scale dependence. Scalability remains a critical challenge, as many methods fail or become prohibitively slow on datasets with large number of spots due to memory, intensive graph constructions, highlighting the need for algorithmic and engineering advances, such as sparse and hierarchical graph strategies, mini-batching, and GPU-enabled implementations. The strong performance of IRIS, the sole reference-based method in our benchmarking, highlights a promising and underexplored direction: integrating external single-cell and multi-omic data, such as spatial ATAC-seq, proteomics, and histology, to provide complementary evidence for domain boundaries and improve spatial domain identification beyond transcriptomics alone. Finally, our benchmarking reveals a key gap in current methods, where each spot is assigned to a single discrete domain despite the presence of gradual biological transitions, underscoring the need for approaches that provide probabilistic assignments or uncertainty estimates to better capture boundary ambiguity.

Taken together, spDDB provides a comprehensive and spatially informed benchmarking resource for evaluating spatial deconvolution and domain detection algorithms. By systematically comparing methods across diverse tissues, technologies, and spatial complexity regimes, our study offers practical guidance for selecting computational approaches and highlights key challenges that remain in spatial transcriptomics analysis. We anticipate that this framework will support the development of next-generation algorithms and facilitate more accurate interpretation of spatial transcriptomic data in studies of development, physiology, and disease.

## Methods

### Overview of SynthST

Extended Figure 1 shows the overview of SynthST, a novel simulator that employs a deep graph attention (GAT) network to generate a synthetic cell type proportion matrix based on the cell type proportions inferred from a real spatial dataset, along with a paired synthetic spatial transcriptomics dataset. SynthST consists of two modules: the first generates a simulated cell-type proportion matrix, and the second uses this matrix to generate the corresponding spatial gene expression data. The first module uses a deep GAT-based architecture comprising three multi-layer neural networks: an encoder *E*, a decoder *D*, and an inner product decoder 𝒟. The encoder *E* takes as input the celltype proportion matrix *x* and the adjacency matrix *A* = [*a*_*ij*_]_*n*×*n*_, and learns a lower-dimensional embedding *z*. The decoder *D* reconstructs the simulated cell-type proportion matrix 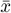 from *z*, while the inner product decoder 𝒟 reconstructs the adjacency matrix *Ā* = [*ā*_*ij*_]_*n*×*n*_ from the same embedding. We train the network using two loss functions. The root mean squared error (RMSE) loss minimizes the difference between the input cell-type proportion matrix *x* and the reconstructed matrix 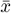. The binary cross-entropy loss measures the discrepancy between the original adjacency matrix *A* and its reconstruction *Ā*. In addition, we incorporate a weight decay term, defined as the squared Euclidean norm of the network weights, which acts as an *L*_2_ regularizer to prevent overfitting and avoid excessive optimization toward a single loss component (see Supplementary Note 1 for details).

For inferring the cell type proportions from the real dataset, we employed top-performing deconvolution methods identified in existing benchmarking studies (16; 15; 17) and the cell type proportion matrices obtained from these methods were used for training the GAT to achieve multiview learning, allowing each cell type to be modeled against the same graph topology. This approach ensured that, for every cell type, its most dominant spatial distribution pattern was captured in the simulated data. Moreover, as different spatial patterns were captured by different subsets of methods, the multi-view strategy helped preserve spatial structures for all the cell types. Furthermore, the weighted neighborhood aggregation operation of the graph attention network enabled SynthST to generate smooth spatial patterns for cell type proportions (62). During inference, the average of the cell-type proportion matrices from the top-performing deconvolution methods is used to generate the simulated cell-type proportion matrix, which is treated as ground truth (Supplementary Note 1).

In the second module of SynthST, we used the simulated cell-type proportion matrix 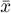 as ground truth to generate a spatial gene expression dataset 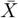 by aggregating the gene expression signature of each cell type weighted by the number of cells in each spot. We employed Cell2location to estimate the gene signature matrix used in this formulation and to obtain the cell-type abundance matrix, scaling abundances to reflect the number of cells per spot according to the technology. To capture the overall spatial structure of the original spatial gene expression *X* and transfer it to the simulated spatial gene expression dataset, we employed Gaussian Processes to model UMI counts, defining the mean as the UMI counts of the original data and using an RBF kernel-based covariance function. Finally, we scaled the simulated spatial gene expression according to the modeled UMI counts (see Supplementary Note 2 for details).

### Simulation of spatial domains

For the spatial domain benchmarking, we used a subset of datasets simulated using SynthST. SynthST’s graph attention network is used for obtaining latent space representations *z* for each cell type proportion matrix of the multiview used for training the network as well as the averaged cell type proportion matrix used in inference. To determine the appropriate number of spatial domains, we applied Leiden clustering (101) across a range of resolutions (0.2–1.0) on the latent embeddings and computed ASW for each setting. As resolution increases, cluster numbers rise and ASW typically decreases; we selected the cluster number corresponding to the first resolution value where ASW begins to improve again. This procedure was performed for the latent embeddings of all training cell type proportion matrices, and the final number of spatial domains was selected via majority voting across these estimates.

### Simulation of datasets using strategy 2

Our simulation strategy 2 employs a binning approach on single-cell–resolution imaging-based datasets (e.g., MERFISH) to generate simulated gene expression and ground-truth cell type proportions for spots following prior benchmarking studies (15; 16). Starting from a real single-cell resolution spatial dataset, we partitioned the two-dimensional spatial coordinates into 50 *µ*m × 50 *µ*m square grids to mimic Visium like splots containing multiple cells with known cell type compositions. We then aggregated the gene expression values of all cells within each grid to form a simulated spot and assigned the center of each grid as its spatial location. The proportion of each cell type within each spot was then computed to obtain the ground-truth cell-type proportion matrix. Since most deconvolution methods require a reference scRNA-seq dataset, we used matched single-cell data from the same study as reference whenever available. Otherwise, we used single-cell resolution spatial datasets with annotated cell types as references for deconvolution.

### Evaluation metrics for cell-type deconvolution

#### Bivariate evaluation metrics

For the computation of bivariate spatial metrics, we employed a weight matrix, *w*_*ij*_, to account for spatial proximity between spots, where *n* is the number of spots, *d*_*ij*_ is the Euclidean distance between spots *i* and *j*. The weights were defined using a Gaussian kernel with bandwidth parameter *l*, chosen such that the highest weights are assigned to one-hop neighbors (Supplementary Table 12):

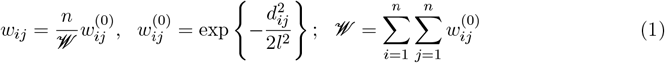

For each cell type *k*, let *x* and *y* denote the vectors of predicted and ground-truth proportions, respectively, with 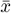 and 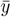 representing their mean values. Metrics were computed independently for each cell type, yielding a score *M*^*k*^, and the final metric value was obtained by averaging across all *K* cell types as:

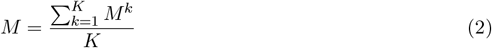

- **Bivariate Moran’s I**: Bivariate Moran’s I is a spatial correlation metric (63) that extends Moran’s I to assess how well two vectors correlate in a spatial context, capturing the preservation of spatial patterns between predicted and ground-truth cell-type distributions. As a global measure of similarity, it evaluates the cross-product of deviations from the mean and ranges from −1 to 1, where 1 indicates strong positive correlation and −1 indicates strong negative correlation. For interpretability, we linearly transformed this range to [0, 1], where 0 denotes negative correlation, 0.5 denotes no correlation, and 1 denotes positive correlation (Supplementary Figure 21).

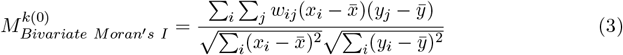

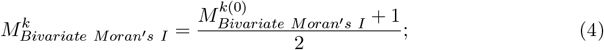
- **Bivariate Geary’s C**: We developed Bivariate Geary’s C as a novel extension of Geary’s C for assessing multivariate spatial association (102). Unlike bivariate Moran’s I, which captures global spatial correlation, bivariate Geary’s C emphasizes local spatial similarity by quantifying squared differences between neighboring locations. Hence, it functions as a deviation-based metric that ranges from 0 to ∞, where values closer to 0 indicate high similarity, and higher metric values indicate greater dissimilarity. For consistency with other metrics, we applied a non-linear transformation to map this range to [0, 1], such that higher values correspond to better agreement between predicted and ground-truth spatial distributions of cell type proportions. Bivariate Geary’s C is defined as:

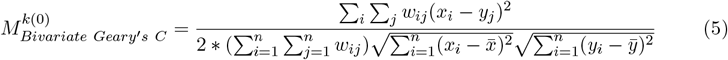

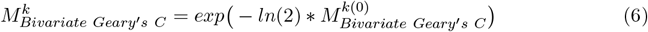
- **Structural Similarity Index Measure (SSIM)**: SSIM quantifies structural similarity between the predicted and ground-truth cell type proportions and is defined as:

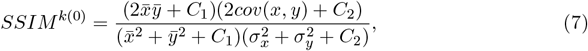

where *C*_1_, *C*_2_ are scalars with values 0.01 and 0.03, respectively, and *σ*_*x*_ and *σ*_*y*_ are the standard deviations of *x* and *y* (15). The metric attains higher values when the two vectors exhibit similar means and variance patterns. Since SSIM ranges from −1 to 1, with 1 indicating perfect structural similarity, we applied a linear transformation to rescale it to [0, 1] for consistency with other bivariate metrics.
- **Lee’s L Statistics**: Lee’s L statistics is a bivariate spatial association metric that integrates Pearson correlation and Moran’s I to assess global spatial relationships between two vectors (64). Unlike bivariate Moran’s I, which relies on the weighted product of deviations from the mean, Lee’s L statistics computes the correlation between spatially aggregated deviations (64). Lee’s L statistic is defined as:

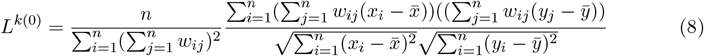

and its value ranges from [−1, 1], with 1 indicating strong positive spatial autocorrelation. We linearly transformed it to [0, 1] for consistency with other metrics.
- **Spatial Pearson Correlation**: Spatial Pearson Correlation is an extension of Pearson correlation that quantifies linear association between vectors while accounting for the spatial relationships (66; 65). It is defined as follows and ranges between [-1, 1], where 1 indicates strong spatial autocorrelation. We linearly transformed it to [0, 1] for consistency with other metrics.

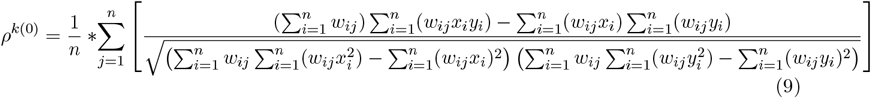

#### Non-spatial evaluation metrics

- **Cosine Similarity**: Cosine similarity measures the cosine of the angle between the predicted and ground-truth variables and is defined as:

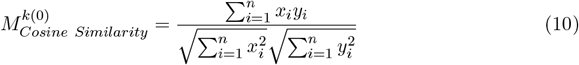

The metric ranges from −1 to 1, where 1 indicates a strong similarity, 0 indicates orthogonality (no similarity), and –1 indicates complete opposition. To align it with the interpretation of other metrics, we linearly transformed its range to [0, 1].
- **Pearson Correlation**: Pearson correlation measures linear association between predicted and ground truth variables and is defined as:

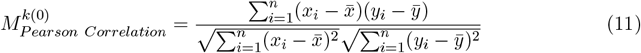

The metric ranges from −1 to 1, where 1 indicates a strong association. We applied a linear transformation similar to the metrics above to map the range to [0, 1].

#### Spot level evaluation metrics

We utilized two other non-spatial metrics which are defined below. In the definitions below, let *X* and *Y* denote the ground truth and predicted cell type proportions matrices, respectively, and let *x*′ and *y*′ denote the cell type proportion vectors for a given spot *s*, with *K* denoting the number of cell types in the deconvolution task. For consistency with other metrics, we applied a linear transformation on the raw metric *M* ^(0)^(*X, Y*) as:

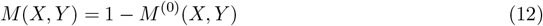

- **Jensen Shannon Divergence (JSD)**: JSD is the symmetric variant of KL divergence, used to measure the similarity between two probability distributions. In the context of cell type deconvolution, each row in the cell type proportion matrix sums to 1, representing a probability distribution. Therefore, we compute JSD at a spot level as: The range of the metric is [0, 1].

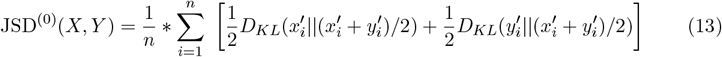

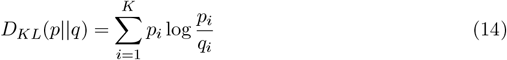
- **Root Mean Squared Error (RMSE)**: We defined RMSE for evaluating cell-type proportion matrices by normalizing the squared error with respect to the number of spots, enabling better discrimination between methods. Under this formulation, RMSE can slightly exceed 1 in extreme cases with many cell types and large deviations between *Y* and *X*; therefore, we cap the metric at a maximum value of 1.

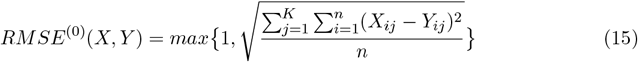

#### Composite score metrics for cell-type deconvolution

- **Composite scores for each dataset:** We used five composite scores to summarise performance for each dataset: 1) Overall Composite Score, 2) Composite Bivariate Spatial Score, 3) Composite Non-Spatial Score, 4) Composite Shape Characterization Score, and 5) Composite Rare Celltypes Score. Before computing these composite scores, we applied min-max scaling to each individual metric, i.e. *M*_*Bivariate Moran*_*′*_*s I*_, *M*_*Bivariate Geary*_*′*_*s C*_, *SSIM, L, ρ, M*_*Cosine Similarity*_, *M*_*P earson Correlation*_, *JSD*(*X, Y*), and *RMSE*(*X, Y*) across all deconvolution methods and their respective cell types. We then applied a second round of min–max scaling to the averaged metric values to ensure that all metrics were comparable and contributed equally to the composite score.
  1. **Composite Bivariate Spatial Score**, *CS*_*Bivariate*_: The Composite Bivariate Spatial Score is defined as the average of all bivariate spatial evaluation metrics:

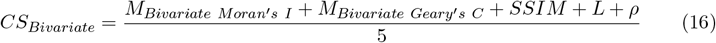
  2. **Composite Non-Spatial Score**, *CS*_*Non*−*Spatial*_: The Composite Non-Spatial Score is defined as the average of all non-spatial evaluation metrics:

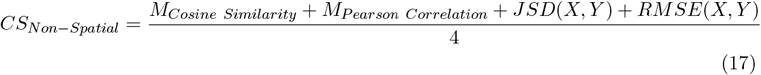
  3. **Composite Shape Characterization Score**, *CS*_*Shape Characterization*_: The Composite Shape Characterization Score is defined as the average of *CS*_*HC*_, *CS*_*HE*_, *CS*_*LE*_, *CS*_*HL*_, *CS*_*LL*_, representing high curl, high elongation, high linearity, low elongation, and low linearity cell shape characterization scores, respectively.

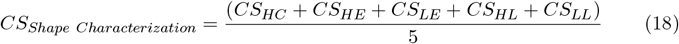

The cell shape characterization scores as used in equation (18) are defined for a set of cell types which satisfy celltype patterns criteria i.e. high curl (HC), high elongation (HE), high linearity (HL), low elongation (LE), and low linearity (LL) respectively. For a specific cell shape pattern (denoted as *pattern*) the score, *CS*_*pattern*_ is computed as the average of all bivariate spatial evaluation metrics for the cell types that harbor the cell shape pattern (*K*′ denotes the set of such cell types):

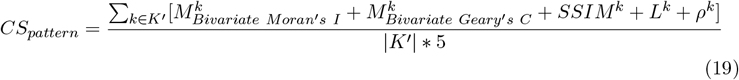

We introduced shape characterization metrics to evaluate performance on cell types exhibiting prominent spatial patterns. Using functions from CellCharter (69; 103), we quantified geometric properties of cell-type distributions, specifically *curl, elongation* and *linearity*, and followed the steps below to identify cell types exhibiting these spatial patterns.
    a. For a given cell type *k*, we converted the cell type proportion vector into a binary vector by assigning a value of 1 (presence) if the proportion was greater than the median proportion value, and 0 (absence) otherwise. Using these binarized ground-truth proportions, we employed CellCharter (69) to identify connected components and compute their geometric properties. CellCharter utilized the concept of alpha shapes, a family of geometric structures parameterized by *α*, to infer the shape of the cell-type distribution (103). For each cell type, a polygon representing its spatial presence was constructed. The final shape was defined using the minimum value of *α* that yielded a single connected polygon. In cases where multiple components were obtained, we considered the component with the largest number of spots.
    b. The prominent spatial shape patterns were defined as follows: high curl represents large curved or round structures; high and low linearity capture large and small linear patterns, respectively; high elongation corresponds to large unstructured shapes, while low elongation reflects small unstructured patterns, including compact curves.
    c. For each dataset, we then defined high curl, high elongation, and high linearity cell types as the cell types falling within the top 25% based on their respective curl, elongation and linearity metrics. Similarly, we defined low elongation and low linearity cell types as the cell types falling in the bottom 25% in terms of elongation and linearity metric. We did not consider low-curl cell types from this analysis because the majority of cell types exhibited zero curl (Supplementary Table 4-6).
  4. **Composite Rare Celltypes Score**, *CS*_*Rare*_: The Composite Rare Celltypes Score is defined as the average of rare celltypes score *CS*_*R*1_, regional rare cell types score *CS*_*R*2_ and *AUPR*, where *R*1 and *R*2 denote the set of rare and regionally rare cell types, respectively.

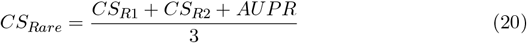

*CS*_*R*1_ and *CS*_*R*2_ are computed as the average of all the bivariate spatial evaluation metrics considering the cell types in *R*1 and *R*2, respectively.
    a. **Rare cell types**, *R*1: We defined rare cell types as the cell types that are present in the bottom 10% based on their mean proportion value across all the spots. Supplementary Table 3 summarizes the rare cell types identified for all the datasets.
    b. **Regional rare cell type definition**, *R*2: We defined regional rare cell types as rare cell types which are rare as well as regionally restricted. We followed the steps below to identify regional rare cell types.
      i. We selected cell types in the bottom 30% based on the mean proportion value across spots. For these cell types, we then performed Hartigan Dip Test to assess the unimodality of the distribution of their proportions across spatial locations (104). A low p-value (*<* 0.05) indicates significant evidence of multimodality, suggesting regional presence of rare cell types. We also confirmed the bimodality using kernel density estimation plots (KDE).
      ii. If the number of regional rare cell types was *>*=5, we selected the bottom 20% of cell types based on mean proportion as our starting set. If no regionally rare cell types are detected, we increased the threshold up to the bottom 50 − 60% until at least one regionally rare cell type was identified. A few datasets did not contain any regional rare cell types. Supplementary Table 3 summarizes the regional rare cell types identified for all the datasets.
    c. **Area under Precision/Recall curve (***AUPR*): AUPR is a classification metric used to evaluate agreement between ground truth and predicted datasets in detecting the presence of rare cell types. Let *x* and *y* denote the column vectors representing the cell type proportion for a given rare cell type *r* in the ground-truth and predicted cell type proportion matrices, respectively. We binarized these vectors by applying a threshold on the ground-truth proportions: for unimodal distributions, the median was used, while for multimodal distributions, the threshold was set at the onset of the second mode. A cell type was considered to be present if its proportion exceeded the threshold, and absent otherwise. Precision, recall and AUPR were then computed using binarized *x* and *y*

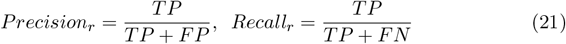

Let *R* be the set of all rare cell types. The AUPR score for a dataset is computed as the mean AUPR across all the rare cell types in *R*.

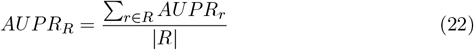
  5. **Overall Composite Score**, *CS*_*Overall*_: The Overall Composite Score is defined as the average of all composite scores, as follows.

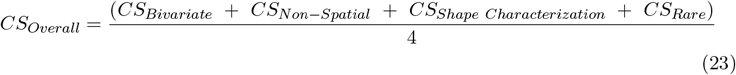
- **Composite scores for tissue category:** While computing the composite and individual metric scores across a set of datasets within a tissue category, we devised a penalization approach to penalize the methods that failed to execute on some datasets. Let *T* denote the total number of datasets in a tissue category, and *N*_*datasets*_(=*< T*) denote the number of datasets on which a method successfully ran. We used a slope-based penalty to adjust the scores accordingly:

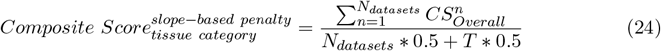

The penalization in equation (24) for computing the average composite score for a tissue category ensures that methods failing on some datasets incur a penalty. Supplementary figure 21d shows the plot for computing penalized denominator based on *N*_*datasets*_ and *T*. Additionally, we computed composite scores without penalizing failed executions to assess the absolute performance of each method on the datasets where it successfully ran:

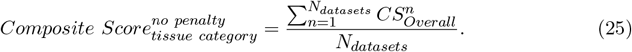

### Evaluation of deconvolution methods not requiring a scRNA-seq reference

Reference-free methods such as STDeconvolve and SpiceMix require the number of cell types as input for deconvolution. These methods output a cell-type proportion matrix (spots × cell types) along with inferred gene expression profiles (cell types × genes). To assign biological annotations to the inferred cell types, we mapped each inferred cell type to a known reference by first aggregating scRNA-seq data into a cell-type–by–gene matrix (summing expression across cells of the same type), then computing the Pearson correlation coefficient (PCC) between each reference profile and the inferred profiles, and finally assigning each inferred cell type the label of the reference cell type with the highest correlation.

### Evaluation metrics for domain detection

To evaluate predicted spatial domains from domain detection methods against ground-truth annotations, we utilized metrics from two broad categories: 1) accuracy metrics, and 2) continuity metrics. Accuracy metrics included Normalized Mutual Information (NMI) (94), Adjusted Rand Index (ARI) (95), Homogeneity (HOM) and Completeness Score (COM) (96). Continuity metrics included Average Silhouette Width (ASW) (97), CHAOS (98; 99) and PAS (100). Let *C*^*True*^ denote the true cluster labels, *C*^*Pred*^ the predicted cluster labels, and *n* the number of spots (20). The metrics are defined as follows:

- **Normalized Mutual Information (NMI):** NMI compares the overlap between two clustering assignments, where the overlap is measured in terms of mutual information. The NMI value is scaled in the range 0 to 1 based on the mean entropy for cluster assignments and cell-type labels.

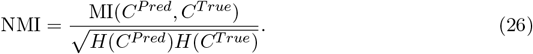
- **Adjusted Rand Index (ARI):** Rand index compares two clustering assignments considering all pairs of points and finding agreement between the clustering assignments (95). The Adjusted Rand Index (ARI) ranges from 0 to 1, where 0 indicates random clustering and 1 represents a perfect agreement. It is defined based on the following components: TP (true positives) represents the number of spot pairs that are grouped in the same cluster in both the true and predicted labels; TN (true negatives) denotes the number of spot pairs assigned to different clusters in both true and predicted labels; FN (false negatives) counts the pairs grouped in the true labels but separated in the predicted clustering; and FP (false positives) indicates the pairs separated in the true labels but grouped in the predictions. The ARI is calculated as follows, where *E* is the expected value of the index under random clustering

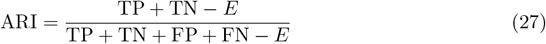

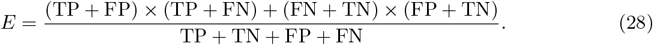
- **Completeness Score (COM)**: The COM score is a metric that evaluates the completeness of a cluster labeling with respect to the ground truth. A clustering result is considered complete if all data points that belong to a certain class are grouped into the same cluster. The COM score ranges from 0 to 1, with a value of 1 indicating a perfect and complete labeling and is defined as:

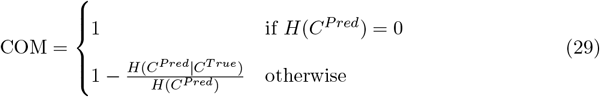

COM scores were computed using functions implemented in the scikit-learn package (105).
- **Homogeneity Score (HOM)**: The Homogeneity (HOM) score quantifies the homogeneity of a cluster labeling when compared to a known ground truth. A clustering is perfectly homogeneous if every cluster consists exclusively of one class. The HOM score ranges from 0 to 1, with 1 representing a perfectly homogeneous labeling and is defined as:

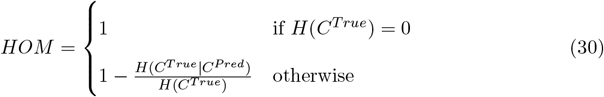

HOM scores were computed using functions implemented in the scikit-learn package (106).
- **Cell type average silhouette width (ASW):** The silhouette width for spot *i* is defined as follows, where *a*(*i*) is the average distance from spot *i* to other spots in the same cluster and *b*(*i*) is the average distance of spot *i* from the spots in the nearest neighboring cluster. Average silhouette width (ASW) is calculated by averaging over all spots, and it ranges from –1 to 1, where −1 or 0 indicate overlapping clusters and values near 1 indicate to well-separated dense clusters. We linearly scaled ASW to the range [0, 1] for consistency with other metrics.

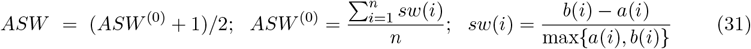
- **CHAOS**: The CHAOS is an unsupervised evaluation metric for measuring the spatial continuity of predicted spatial domains in spatial transcriptomics. For each domain *k, ω*_*kij*_ is computed for all spots present in the domain as follows. Suppose *n*_*k*_ is the number of cells in the *k*-th domain, *d*_*ij*_ is the euclidean distance between spot *i* and *j* and *β* is a hyperparameter in the transformation function. The range of the metric is 0 − ∞, hence we scaled it to the interval [0, 1].

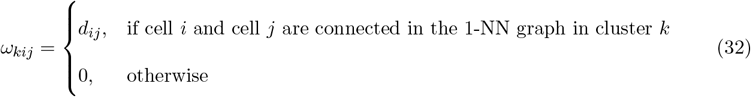

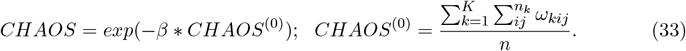
- **PAS**: The PAS score measures the spatial continuity of clusters (domains) by calculating the proportion of cells whose assigned spatial domain label differs from at least six of their ten nearest neighbors. The original metric ranges from 0 to 1, where a lower value indicates better spatial continuity. To align the interpretation with clustering performance, we transform the metric so that a score closer to 1 reflects improved clustering performance.

### Competing methods

We performed benchmarking of 21 state-of-the-art cell-type deconvolution methods: Stereoscope (23), RCTD (24), Autogenes (25), Redeconve (26), SpotLight (27), SpatialDWLS (28), CARD (29), SpiceMix (30), SD (31), Tangram (32), DestVI (33), GraphST (3), CellDART (34), POLARIS (35), Seurat V3 (36), STRIDE (37), SpatialDecon (38), Cell2location (39), STDeconvolve (40), SONAR (41) and CelloScope (42). To ensure a fair evaluation, preprocessing was performed according to each method’s recommended guidelines. Details of the run configurations for the cell-type deconvolution methods are provided in Supplementary Table 1.

For the benchmarking of domain detection methods, we selected 18 state-of-the-art methods: SpaSRL (71), SCANIT (72), SpatialPCA (73), DeepST (74), GraphST (3), SpaceFlow (75), CCST (76), Banksy (77), Giotto (78), DR-SC (79), iSCMEB (80), BayesSpace (81), PRECAST (82), BayesCafe (83), and BASS (84), STAGATE (85), PROST (86), IRIS (87). To ensure a fair evaluation, preprocessing was performed according to each method’s recommended guidelines. The details of the run configurations for the domain detection methods are provided in Supplementary Table 10.

## Data availability

All datasets used in the study are publicly available. The simulated spatial gene expression datasets generated for spatial deconvolution and domain detection benchmarking have been deposited on figshare and are accessible through the project website: https://zafar-lab.github.io/spDDB_datasets.github.io/. Detailed information about all datasets used in this study is provided in Supplementary Tables 2 and 11.

For the spatial cell-type deconvolution task, the spatial and single cell gene expression datasets used to generate simulated spatial gene expression datasets are as follows. The spatial and scRNA dataset for DLPFC are available at ref (46) and (47) respectively, Mouse Brain at ref (39), Hippocampus at ref (24), Cerebellum at ref (24), Visium HD Mouse brain at ref (45) (https://www.10xgenomics.com/datasets, scRNA data at ref (39)), Kidney Cancer at ref (49), Breast Cancer at ref (50), Liver Cancer at ref (51) and (52) respectively, Prostate Cancer at ref (53), Visium HD Lung Cancer at ref (45) (https://www.10xgenomics.com/datasets)and (54), Developmental Lung at ref (56), Human Breast Atlas at ref (50), Intestine at ref (58), Mouse Liver Atlas at ref (59), Human Liver Atlas at ref (59), Kidney Atlas at ref (60) and Chicken heart at ref (57). For the deconvolution task on the simulation strategy 2 datasets, MERFISH preoptic hypothalamic brain dataset is available at ref (48), MERFISH Ileum is available at ref (61), MERFISH Breast Cancer at ref (55), and MERFISH Lung Cancer at ref (55).

For the domain detection task, Tthe spatial and single cell gene expression datasets for DLPFC is available at ref(46) (spatial:https://research.libd.org/spatialLIBD/ (107)) and (47) respectively, Mouse embryo at ref (88)https://db.cngb.org/stomics/mosta/ and (89) respectively, Mouse breast cancer at ref (3) and (90) respectively, osmFISH Mouse cortex at ref (91) (http://sdmbench.drai.cn/ (20)), and (92) respectively, and MERFISH brain at ref (48) (http://sdmbench.drai.cn/ (20)). All other data supporting the findings of this study are available within the article and its supplementary files. Any additional requests for information can be directed to, and will be fulfilled by, the corresponding author.

## Code availability

Our benchmarking pipeline for both the tasks is provided as reproducible pipeline at https://github.com/Zafar-Lab/spDDB. The simulator code is available at https://github.com/Zafar-Lab/spDDB/tree/main/SynthST and code for all the evaluation experiments at https://github.com/Zafar-Lab/spDDB/tree/main/Experiments.

## Acknowledgements

This work was partially funded by the DBT/Wellcome Trust India Alliance Early Career Fellowship [grant IA/E/21/1/506298], IIT Kanpur initiation grant [IITK /CS /2019236], and Science and Engineering Research Board (SERB), Government of India Startup Research Grant [SRG/2020/001333] to H.Z.

## Author contributions

H.Z. designed the study. A.S. and H.Z. developed the simulator algorithm. A.S. implemented the simulator, curated the dataset repository, and generated the simulated datasets. A.S., A.V., and T.K. performed the installation setup, ran the benchmarking methods, and implemented the bivariate metrics. A.S. conducted the evaluation experiments and generated all the figures. A.S. and H.Z. wrote the manuscript; all authors approved the final manuscript.

## Competing interests

The authors declare no competing interests.

---

**Extended Figure 1:**
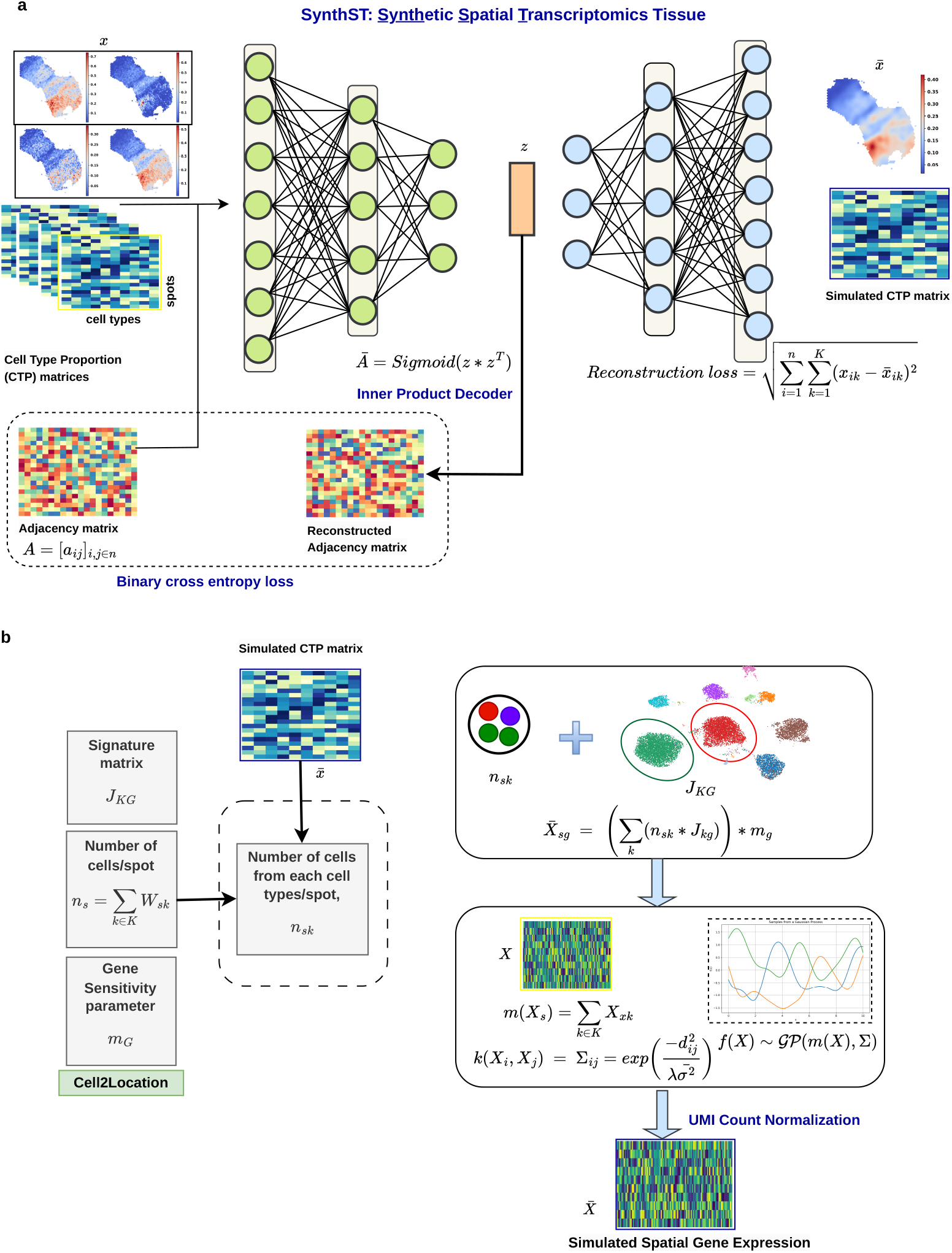
Overview of SynthST: a) SynthST generates a synthetic cell type proportion matrix based on the cell type proportions inferred from a real spatial dataset. SynthST consists of a graph attention autoencoder and an inner-product decoder. The graph attention autoencoder reconstructs the cell-type proportion matrix and is trained using mean squared error loss, while the inner-product decoder reconstructs the adjacency matrix and is trained using a binary cross-entropy loss. The output of this stage is a simulated cell-type proportion matrix, which is treated as ground truth. b) In the second stage of SynthST, the simulated cell-type proportion matrix is combined with cell-type gene signatures, abundance estimates, and gene sensitivity parameters derived from reference single-cell data to generate a simulated spatial gene expression matrix, with UMI counts modeled and normalized via Gaussian processes.

**Extended Figure 2:**
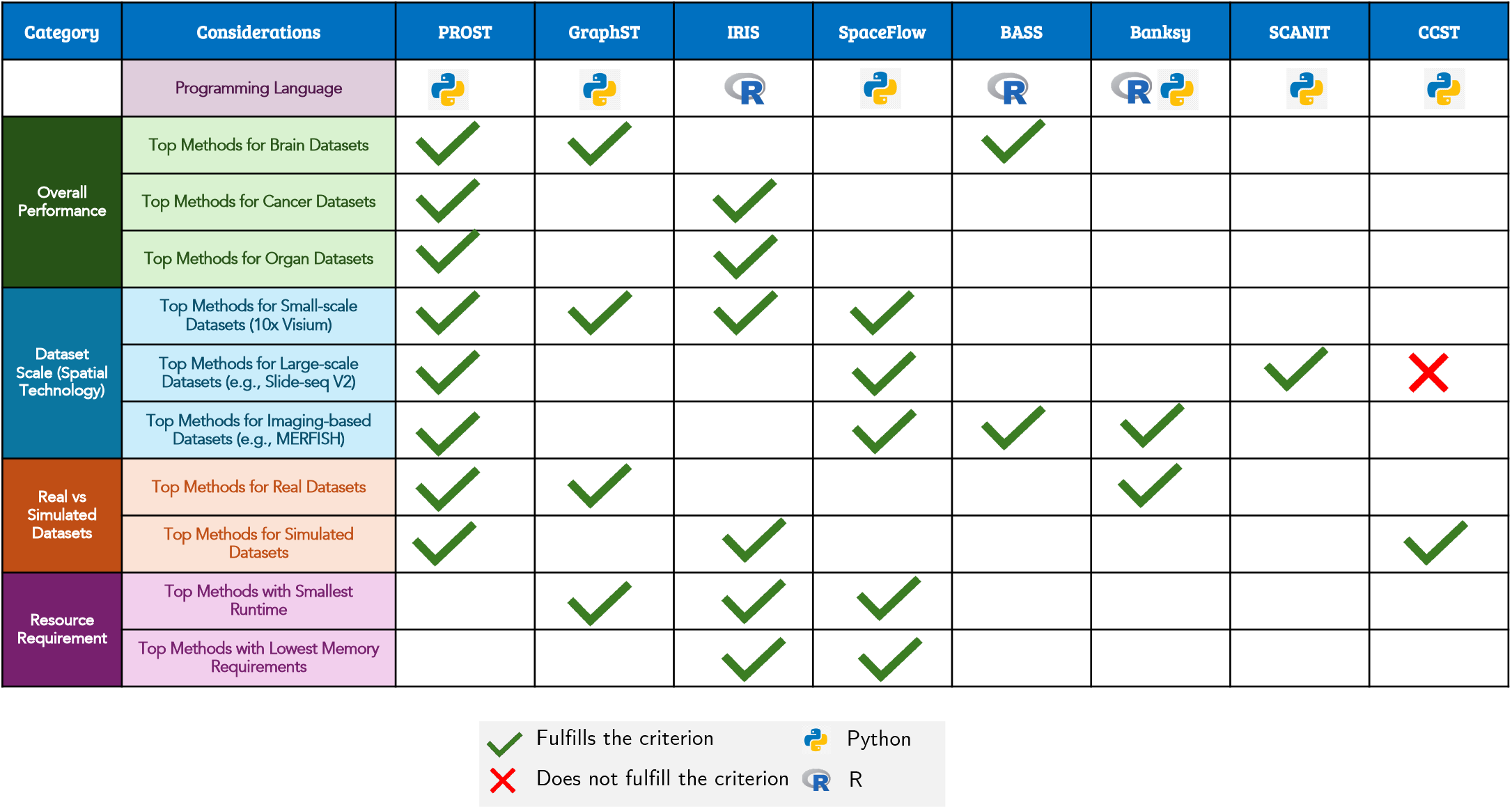
Guidelines for selecting domain detection methods. Table of criteria for selecting a domain detection method, and the corresponding methods that satisfy each criterion. Considerations are divided into the four broad categories - Overall Performance, Dataset Scale (Spatial Technology), Real vs Simulated Datasets and Resource Requirements. Python and R symbols indicate the primary language in which the method is programmed and used. A green tick indicates if a method fulfills the criterion, and a red cross indicates that the method does not fulfill the criterion.

## Notes

### Competing Interest Statement

The authors have declared no competing interest.

