## Supplementary Material for "A Comprehensive Benchmarking of Spatial Deconvolution and Domain Detection Methods across Diverse Tissues and Spatial Transcriptomic Technologies"

May 11, 2026

#### Contents

**Supplementary Figures** **2**

**Supplementary Tables** **55**

**Supplementary Note** **67**

**References** **80**

### Supplementary Figures

a

| Li et al., 2022 (Nat Methods) | Li et al., 2023 (Nat Comm.) | Sang-aram et al., 2024 (eLife) | spDDB | Methods |
|---|---|---|---|---|
| | | | ✓ | Autogenes |
| ✓ | | | ✓ | CARD |
| ✓ | ✓ | ✓ | ✓ | Cell2location |
| | | | ✓ | CellDART |
| | | | ✓ | CelloScope |
| ✓ | ✓ | ✓ | ✓ | DestVI |
| | | | ✓ | GraphST |
| | | | ✓ | Polaris |
| ✓ | ✓ | ✓ | ✓ | RCTD |
| | | | ✓ | Redeconve |
| | ✓ | | ✓ | SD |
| ✓ | | ✓ | ✓ | Seurat |
| | ✓ | | ✓ | SpatialDecon |
| ✓ | ✓ | ✓ | ✓ | SpatialDWLS |
| | | | ✓ | SONAR |
| ✓ | ✓ | ✓ | ✓ | Spotlight |
| | ✓ | | ✓ | Spicemix |
| | ✓ | | ✓ | STDeconvolve |
| ✓ | ✓ | ✓ | ✓ | Stereoscope |
| ✓ | ✓ | ✓ | ✓ | STRIDE |
| ✓ | ✓ | ✓ | ✓ | Tangram |

b

| Li et al., 2023 (Nat Comm.) | Sang-aram et al., 2024 (eLife) | spDDB | Criteria |
|---|---|---|---|
| 2 | 3 | 4 | Tissue Types in Image based datasets |
| 5, 10, 15, 20, 50, 100 um | 50um | 50um | Binning size |
| 45 | 3 | 4 | Number of binned datasets |
| 12 * (~256 * 135), 6<br>12 * (~1100 * 135), 6<br>12 * (4500 * 135), 6<br>2300 * {26k, 18k, 9k}, 22<br>660 * {26k, 18k, 9k}, 22<br>299 * {26k, 18k, 9k}, 22 | 108 * 996,<br>63 * 10000,<br>63 * 10000 | ~7000 * 27998, 10<br>2445 * 242, 16<br>33000 * 500, 9<br>13000 * 488, 13 | Dimensions of binned datasets (Number of cell types in the deconvolution task) |
| 5 Spatial datasets | 6 scRNA datasets + 2 Spatial data | 17 (paired scRNA and Spatial datasets) | Tissue types in sequencing based datasets |
| 523 * 10000, 6<br>316 * 19738, 20<br>2576 * 31053, 56<br>2276 * 29570, 18<br>53208 * 23265, 11 | 9 * (750 * 17538, 18<br>750 * 11906, 8<br>750 * 12704, 6<br>750 * 10684, 12<br>750 * 9612, 16<br>750 * 25750, 15<br>750 * 11857, 14) | 6 * (~4000 * 11403), 17, 2 * (2328 * 12152), 59<br>41798 * 11702, 22, 27196 * 11808, 11<br>98569 * 10130, 7, 3 * (~2500 * 9648), 12<br>2 * (~2000 * 7093), 22, 4 * (~4500 * 7093), 29<br>23945 * 24728, 30, 150589 * 17797, 24<br>2 * (~2500 * 32664), 10, 2647 * 33514, 9<br>2315 * 13541, 9, 1915 * 3688, 14<br>3044 * 20374, 27, 2 * (~1700 * 31017), 17<br>1759 * 32643, 17, 20852 * 16584, 26<br>30296 * 16799, 26 | Number of samples, Dimensions of sequencing based datasets (Number of cell types in the deconvolution task)<br><br>Cancer datasets highlighted |

Supplementary Figure 1: a) Summary table of existing benchmarking studies and computational methods for cell-type deconvolution. A green tick indicates that the method was included in the respective benchmarking study. A yellow box indicates that the method is newly introduced in our benchmarking framework. b) Summary of benchmarking studies comparing cell-type deconvolution methods based on dataset characteristics, including tissue types, spot size, and dataset dimensions.

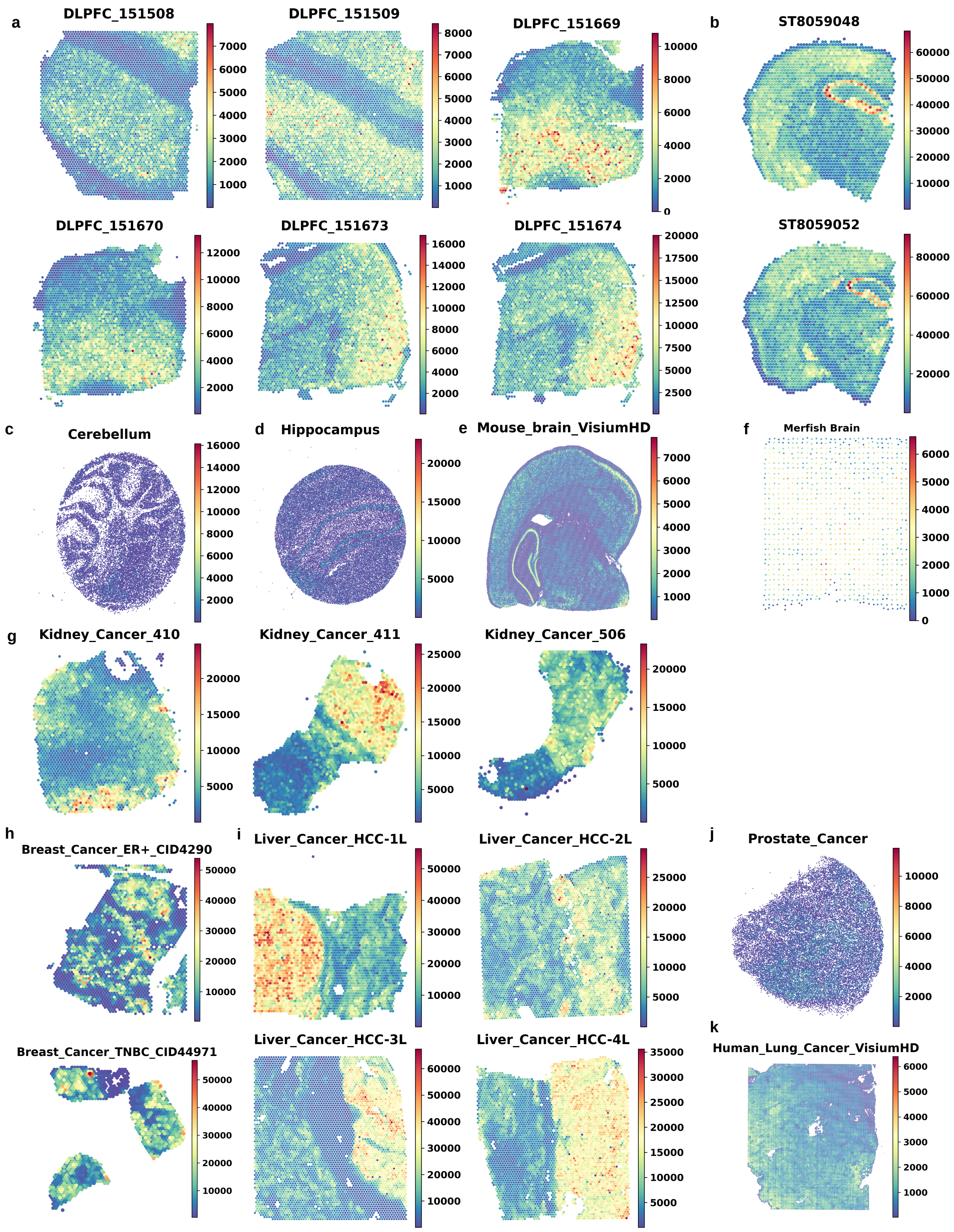
a
DLPFC_151508
DLPFC_151509
DLPFC_151669
b
ST8059048
DLPFC_151670
DLPFC_151673
DLPFC_151674
ST8059052
c
Cerebellum
d
Hippocampus
e
Mouse_brain_VisiumHD
f
Merfish Brain
g
Kidney_Cancer_410
Kidney_Cancer_411
Kidney_Cancer_506
h
Breast_Cancer_ER+_CID4290
i
Liver_Cancer_HCC-1L
Liver_Cancer_HCC-2L
j
Prostate_Cancer
Breast_Cancer_TNBC_CID44971
Liver_Cancer_HCC-3L
Liver_Cancer_HCC-4L
k
Human_Lung_Cancer_VisiumHD

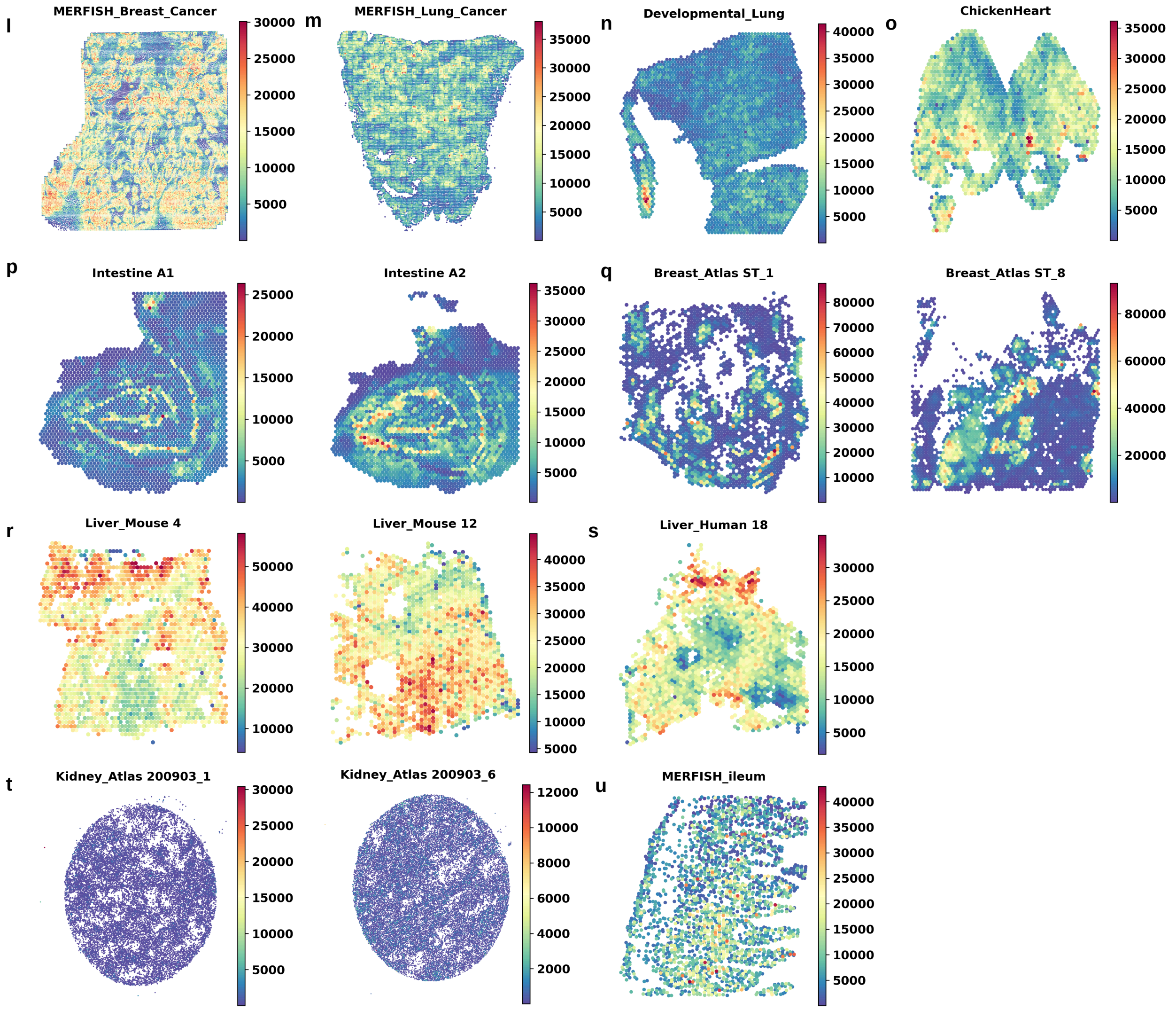


Supplementary Figure 2: Spatial pattern plot representing total UMI counts for Brain, Cancer and Organs datasets, including a) 10X Visium DLPFC 151508, DLPFC 151509, DLPFC 151669, DLPFC 151670, DLPFC 151673, DLPFC 151674, b) 10X Visium Mouse Brain ST8059048(ST48), Mouse Brain ST8059052 (ST52), c) SlideseqV2 Cerebellum, d) SlideseqV2 Hippocampus, e) Merfish brain, f) 10X Visium Kidney Cancer 410, Kidney Cancer 411, Kidney Cancer 506, g) 10X Visium Breast Cancer ER+ CID4290 (ER+), Breast Cancer TNBC CID44971(TNBC), h) 10X Visium Liver Cancer HCC-1L (1L), Liver Cancer HCC-2L (2L), Liver Cancer HCC-3L (3L), Liver Cancer HCC-4L (4L) j) SlideseqV2 Prostate Cancer, k) Visium HD Human Lung Cancer, l) Merfish Breast Cancer, m) Merfish Lung Cancer, n) 10X Visium Developmental Lung, o) 10X Visium Chicken Heart, p) 10X Visium Intestine A1, Intestine A2, q) 10X Visium Human Breast $ST\_1$, Human Breast $ST\_8$, r) 10X Visium Mouse Liver 4, Mouse Liver - 12, s) Human Liver - 18, t) SlideseqV2 Kidney 200903_1 (1), Kidney 200903_6 (6), and u) Merfish ileum

**a**

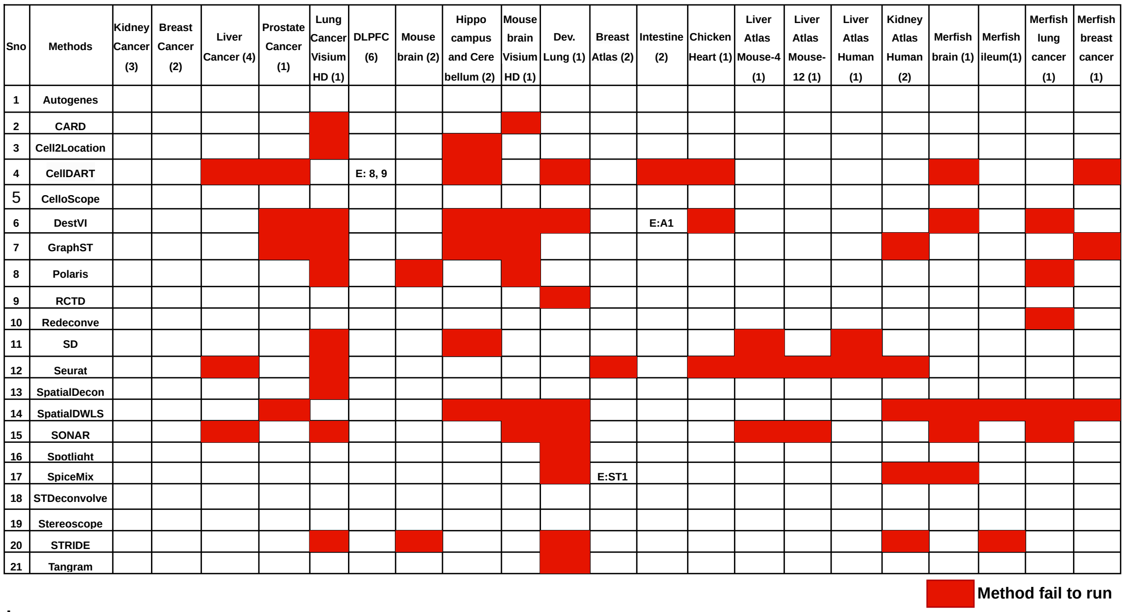


**b**

| Method | Overall Composite Score | Composite Bivariate Spatial Score | Composite Non-Spatial Score | Composite Shape Characterization Score | Composite Rare Celltypes Score | Bivariate Moran's I | Lee's L statistics | Spatial Pearson Correlation | SSIM | Bivariate Geary' C | Cosine Similarity | Pearson Correlation | JSD | RMSE | High Curl Celltypes Score | High Elongation Celltypes Score | High Linearity Celltypes Score | Low Elongation Celltypes Score | Low Linearity Celltypes Score | Rare Celltypes Score | Regional Rare Celltypes Score | AUPR |
|---|---|---|---|---|---|---|---|---|---|---|---|---|---|---|---|---|---|---|---|---|---|---|
| Cell2location | 0.83 | 0.85 | 0.94 | 0.8 | 0.75 | 0.78 | 0.77 | 0.75 | 0.66 | 0.39 | 0.94 | 0.8 | 0.72 | 0.74 | 0.77 | 0.77 | 0.79 | 0.86 | 0.8 | 0.73 | 0.74 | 0.73 |
| SONAR | 0.8 | 0.82 | 0.88 | 0.78 | 0.74 | 0.78 | 0.77 | 0.73 | 0.62 | 0.45 | 0.88 | 0.81 | 0.62 | 0.64 | 0.78 | 0.79 | 0.79 | 0.77 | 0.79 | 0.72 | 0.79 | 0.72 |
| RCTD | 0.79 | 0.81 | 0.83 | 0.79 | 0.73 | 0.75 | 0.73 | 0.72 | 0.61 | 0.42 | 0.84 | 0.78 | 0.61 | 0.63 | 0.79 | 0.8 | 0.79 | 0.78 | 0.8 | 0.65 | 0.78 | 0.71 |
| DestVI | 0.74 | 0.74 | 0.85 | 0.71 | 0.67 | 0.74 | 0.73 | 0.7 | 0.59 | 0.31 | 0.9 | 0.75 | 0.61 | 0.65 | 0.74 | 0.69 | 0.72 | 0.69 | 0.71 | 0.67 | 0.73 | 0.68 |
| CARD | 0.74 | 0.72 | 0.85 | 0.69 | 0.69 | 0.72 | 0.71 | 0.68 | 0.58 | 0.3 | 0.9 | 0.74 | 0.59 | 0.62 | 0.69 | 0.68 | 0.7 | 0.67 | 0.72 | 0.73 | 0.75 | 0.66 |
| Stereoscope | 0.71 | 0.71 | 0.78 | 0.69 | 0.65 | 0.69 | 0.68 | 0.67 | 0.53 | 0.36 | 0.86 | 0.71 | 0.58 | 0.57 | 0.7 | 0.69 | 0.67 | 0.68 | 0.7 | 0.69 | 0.67 | 0.65 |
| STRIDE | 0.68 | 0.66 | 0.79 | 0.64 | 0.64 | 0.68 | 0.67 | 0.65 | 0.56 | 0.25 | 0.9 | 0.7 | 0.65 | 0.67 | 0.65 | 0.68 | 0.64 | 0.63 | 0.62 | 0.66 | 0.66 | 0.65 |
| Tangram | 0.66 | 0.67 | 0.76 | 0.64 | 0.56 | 0.7 | 0.69 | 0.68 | 0.45 | 0.35 | 0.84 | 0.72 | 0.56 | 0.58 | 0.68 | 0.64 | 0.64 | 0.64 | 0.63 | 0.53 | 0.62 | 0.64 |
| CellDART | 0.65 | 0.63 | 0.77 | 0.6 | 0.61 | 0.71 | 0.7 | 0.65 | 0.54 | 0.29 | 0.91 | 0.71 | 0.57 | 0.56 | 0.61 | 0.57 | 0.58 | 0.6 | 0.62 | 0.65 | 0.61 | 0.64 |
| GraphST | 0.64 | 0.65 | 0.7 | 0.63 | 0.57 | 0.7 | 0.7 | 0.63 | 0.48 | 0.3 | 0.83 | 0.69 | 0.53 | 0.52 | 0.65 | 0.63 | 0.62 | 0.61 | 0.65 | 0.62 | 0.58 | 0.61 |
| Polaris | 0.62 | 0.61 | 0.75 | 0.59 | 0.54 | 0.62 | 0.62 | 0.62 | 0.56 | 0.28 | 0.91 | 0.64 | 0.63 | 0.65 | 0.59 | 0.58 | 0.59 | 0.58 | 0.6 | 0.61 | 0.6 | 0.58 |
| CelloScope | 0.56 | 0.55 | 0.67 | 0.55 | 0.45 | 0.61 | 0.61 | 0.6 | 0.44 | 0.29 | 0.86 | 0.62 | 0.56 | 0.6 | 0.56 | 0.55 | 0.57 | 0.53 | 0.55 | 0.5 | 0.53 | 0.54 |
| Autogenes | 0.54 | 0.54 | 0.59 | 0.54 | 0.48 | 0.6 | 0.6 | 0.61 | 0.44 | 0.29 | 0.77 | 0.61 | 0.48 | 0.45 | 0.52 | 0.55 | 0.57 | 0.54 | 0.52 | 0.6 | 0.53 | 0.55 |
| SD | 0.53 | 0.49 | 0.7 | 0.49 | 0.46 | 0.63 | 0.62 | 0.6 | 0.51 | 0.1 | 0.89 | 0.63 | 0.58 | 0.63 | 0.48 | 0.51 | 0.52 | 0.48 | 0.46 | 0.55 | 0.48 | 0.54 |
| STDeconvolve | 0.5 | 0.47 | 0.62 | 0.49 | 0.43 | 0.57 | 0.56 | 0.58 | 0.52 | 0.18 | 0.85 | 0.57 | 0.59 | 0.61 | 0.5 | 0.49 | 0.52 | 0.47 | 0.49 | 0.49 | 0.43 | 0.55 |
| SpatialDWLS | 0.49 | 0.47 | 0.47 | 0.48 | 0.55 | 0.59 | 0.58 | 0.51 | 0.44 | 0.27 | 0.63 | 0.59 | 0.47 | 0.52 | 0.48 | 0.47 | 0.49 | 0.49 | 0.47 | 0.41 | 0.48 | 0.72 |
| Spotlight | 0.47 | 0.44 | 0.48 | 0.44 | 0.5 | 0.57 | 0.56 | 0.56 | 0.4 | 0.27 | 0.7 | 0.58 | 0.42 | 0.33 | 0.46 | 0.44 | 0.41 | 0.45 | 0.45 | 0.54 | 0.54 | 0.6 |
| SpiceMix | 0.42 | 0.4 | 0.48 | 0.43 | 0.37 | 0.57 | 0.57 | 0.6 | 0.26 | 0.21 | 0.71 | 0.58 | 0.39 | 0.38 | 0.48 | 0.42 | 0.44 | 0.44 | 0.4 | 0.42 | 0.44 | 0.51 |
| SpatialDecon | 0.4 | 0.33 | 0.5 | 0.36 | 0.42 | 0.49 | 0.48 | 0.51 | 0.46 | 0.08 | 0.79 | 0.49 | 0.52 | 0.56 | 0.35 | 0.34 | 0.36 | 0.37 | 0.38 | 0.5 | 0.34 | 0.57 |
| Redeconve | 0.37 | 0.35 | 0.3 | 0.36 | 0.49 | 0.49 | 0.49 | 0.48 | 0.45 | 0.21 | 0.56 | 0.51 | 0.52 | 0.6 | 0.33 | 0.39 | 0.38 | 0.33 | 0.36 | 0.24 | 0.39 | 0.72 |
| Seurat | 0.3 | 0.27 | 0.2 | 0.3 | 0.41 | 0.45 | 0.44 | 0.42 | 0.4 | 0.27 | 0.53 | 0.48 | 0.47 | 0.42 | 0.34 | 0.32 | 0.29 | 0.27 | 0.3 | 0.22 | 0.28 | 0.7 |

Composite: L – H
Bivariate Spatial: L – H
Non-Spatial: L – H
Shape Characterization: L – H
Rare: L – H

**All datasets: No Penalty**

Supplementary Figure 3: a)Summary table of deconvolution methods (rows) and datasets (columns). Red boxes indicate methods that encountered errors on specific datasets. b) Summary Table of the composite score performance of all the methods without penalty. Visualization of the method's quantitative performance in terms of Composite scores (red), Bi-variate Spatial metrics (purple), Non-Spatial metrics (blue), Shape Characterization metrics (Orange) and Rare Cell-type metrics (Green).

**a**

| Sno | Methods | # in Top 3 | # Top | # Datasets-Run failed | # in Top 3 - Brain Tissue | # in Top 3 - Cancer Tissue | # in Top 3 - Organs Tissue | # in Top 3 - Visium datasets | # in Top 3 - Large datasets | # in Top 3 - Image-based datasets |
|---|---|---|---|---|---|---|---|---|---|---|
| 1 | Cell2Location | 23 | 10 | 3 | 4 | 10 | 9 | 19 | 4 | 0 |
| 2 | RCTD | 19 | 10 | 1 | 4 | 6 | 9 | 14 | 5 | 0 |
| 3 | SONAR | 17 | 2 | 7 | 4 | 6 | 7 | 12 | 4 | 1 |
| 4 | CARD | 10 | 2 | 2 | 6 | 1 | 3 | 6 | 3 | 1 |
| 5 | STRIDE | 8 | 3 | 5 | 3 | 5 | 0 | 5 | 3 | 0 |
| 6 | Stereoscope | 6 | 2 | 0 | 3 | 2 | 1 | 5 | 0 | 1 |
| 7 | DestVI | 6 | 2 | 8 | 3 | 1 | 2 | 5 | 0 | 1 |
| 8 | Tangram | 6 | 2 | 1 | 5 | 1 | 0 | 5 | 1 | 0 |
| 9 | Autogenes | 4 | 0 | 0 | 0 | 2 | 2 | 1 | 0 | 3 |
| 10 | CellDART | 3 | 1 | 8 | 2 | 0 | 1 | 2 | 1 | 0 |
| 11 | GraphST | 3 | 1 | 7 | 0 | 2 | 1 | 3 | 0 | 0 |
| 12 | STDeconvolve | 1 | 0 | 0 | 1 | 0 | 0 | 0 | 0 | 1 |
| 13 | SpatialDecon | 1 | 1 | 0 | 0 | 1 | 0 | 0 | 0 | 1 |
| 14 | Polaris | 1 | 1 | 4 | 1 | 0 | 0 | 0 | 0 | 1 |
| 15 | SpiceMix | 1 | 0 | 2 | 0 | 1 | 0 | 1 | 0 | 0 |
| 16 | Seurat V3 | 1 | 1 | 7 | 0 | 0 | 1 | 0 | 0 | 1 |
| 17 | spatialDWLS | 1 | 0 | 10 | 0 | 1 | 0 | 0 | 1 | 0 |
| 18 | Spotlight | 0 | 0 | 1 | 0 | 0 | 0 | 0 | 0 | 0 |
| 19 | SD | 0 | 0 | 4 | 0 | 0 | 0 | 0 | 0 | 0 |
| 20 | Redeconve | 0 | 0 | 1 | 0 | 0 | 0 | 0 | 0 | 0 |
| 21 | Celloscope | 0 | 0 | 0 | 0 | 0 | 0 | 0 | 0 | 0 |

**b**

| Sno | Brain Datasets | CARD | Tangram | Cell2 location | SONAR | RCTD | DestVI | Stereo scope | STRIDE | Cell DART | STDecon volve | Polaris |
|---|---|---|---|---|---|---|---|---|---|---|---|---|
| 1 | DLPFC 151508 | I | III | | | | | II | | | | |
| 2 | DLPFC 151509 | III | I | | | | II | | | | | |
| 3 | DLPFC 151669 | | II | | | III | | | | I | | |
| 4 | DLPFC 151670 | | I | | | | II | III | | | | |
| 5 | DLPFC 151673 | II | | I | | | | | III | | | |
| 6 | DLPFC 151674 | III | II | | | | I | | | | | |
| 7 | Mouse Brain ST 48 | | | I | II | III | | | | | | |
| 8 | Mouse Brain ST 52 | | | I | II | III | | | | | | |
| 9 | Cerebellum | II | | | III | I | | | | | | |
| 10 | Hippocampus | III | | | II | | | | I | | | |
| 11 | Mouse Brain Visium HD | | | I | | | | | II | III | | |
| 12 | MERFISH Brain | | | | | | | II | | | III | I |

c

| Sno | Cancer Datasets | Cell2 location | RCTD | SONAR | STRIDE | Graph ST | Spice mix | Stereo scope | CARD | Tangram | Spatial DWLS | Auto genes | Dest VI | Spatial Decon |
|---|---|---|---|---|---|---|---|---|---|---|---|---|---|---|
| 1 | Kidney Cancer 410 | III | I | II | | | | | | | | | | |
| 2 | Kidney Cancer 411 | II | I | III | | | | | | | | | | |
| 3 | Kidney Cancer 506 | II | I | III | | | | | | | | | | |
| 4 | Breast Cancer ER+ | I | III | II | | | | | | | | | | |
| 5 | Breast Cancer TNBC | I | | III | II | | | | | | | | | |
| 6 | Liver Cancer 1L | III | | | I | | II | | | | | | | |
| 7 | Liver Cancer 2L | II | | | III | | | I | | | | | | |
| 8 | Liver Cancer 3L | III | | | II | I | | | | | | | | |
| 9 | Liver Cancer 4L | II | | | I | III | | | | | | | | |
| 10 | Prostate Cancer | I | III | | | | | | II | | | | | |
| 11 | Lung Cancer Visium HD | | I | | | | | | | II | III | | | |
| 12 | MERFISH Breast Cancer | | | II | | | | | | | | III | I | |
| 13 | MERFISH Lung Cancer | | | | | | | I | | | | II | | III |

d

| Sno | Organ Datasets | Cell2 location | SONAR | RCTD | CARD | DestVI | Autogenes | GraphST | Cell DART | Seurat | Stereoscope |
|---|---|---|---|---|---|---|---|---|---|---|---|
| 1 | Developmental Lung | I | | | | | III | II | | | |
| 2 | Chicken Heart | I | II | | | | | | | | III |
| 3 | Breast Atlas ST1 | | I | II | | III | | | | | |
| 4 | Breast Atlas ST8 | III | | II | I | | | | | | |
| 5 | Intestine A1 | III | I | II | | | | | | | |
| 6 | Intestine A2 | | II | I | | III | | | | | |
| 7 | Liver Atlas Mouse - 4 | II | | I | | | | | III | | |
| 8 | Liver Atlas Mouse - 12 | II | | i | III | | | | | | |
| 9 | Liver Atlas Human | I | III | II | | | | | | | |
| 10 | Kidney Atlas - 1 | III | II | I | | | | | | | |
| 11 | Kidney Atlas - 6 | II | III | I | | | | | | | |
| 12 | MERFISH Ileum | | | | II | | III | | | I | |

Supplementary Figure 4: a) Summary table of deconvolution methods (rows) evaluated across different experimental criteria (columns). b–d) Summary tables showing datasets (rows) and deconvolution methods (columns) that ranked among the top three performers classified by tissue type: (b) brain, (c) cancer, and (d) organ datasets. Green boxes indicate top-performing methods, whereas red boxes indicate methods that failed to run.

a

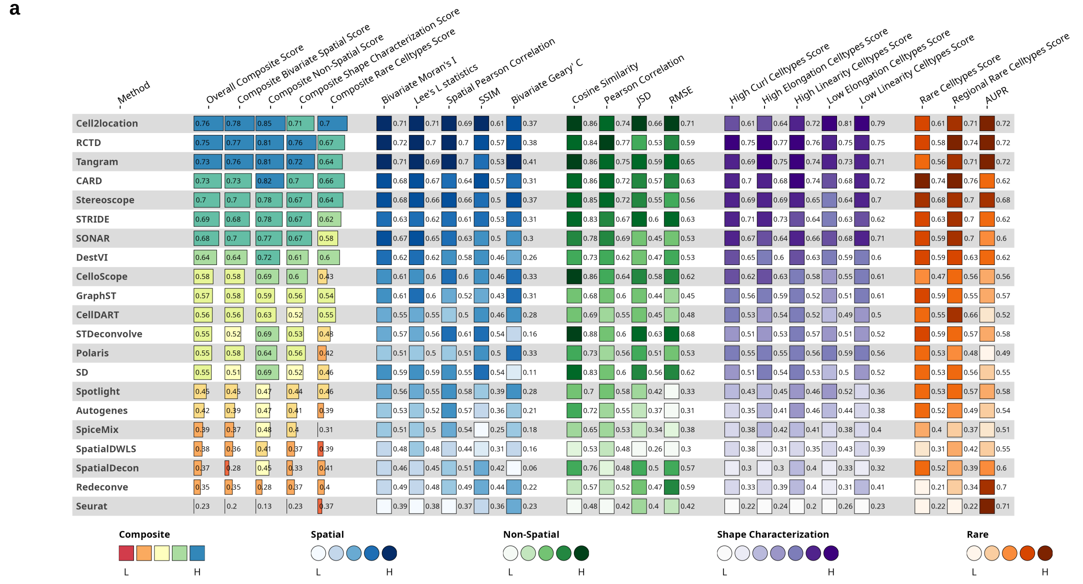


**Brain: Slope-based Penalty**

b

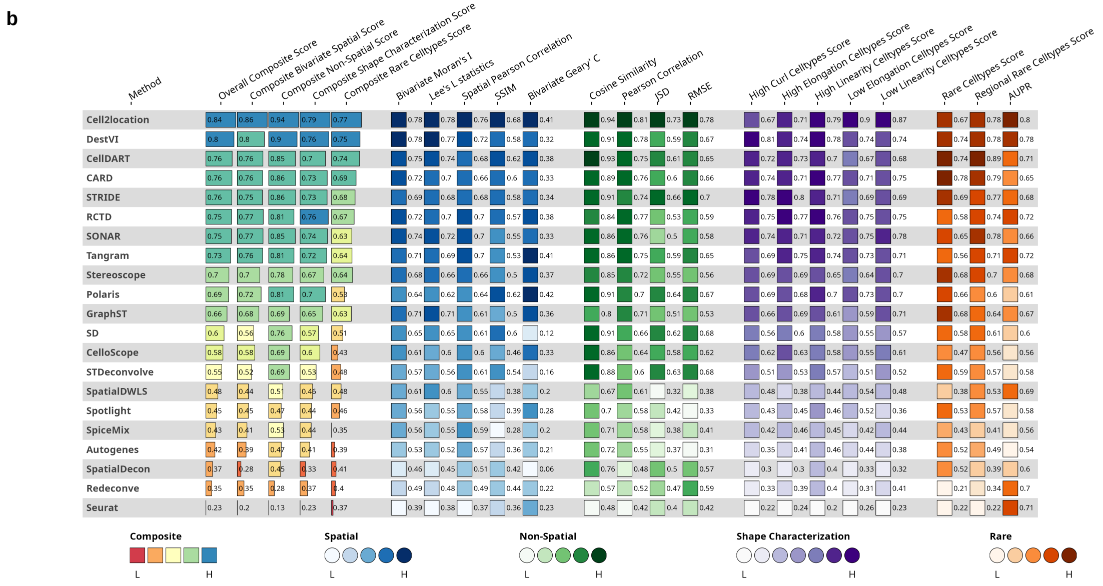


**Brain: No Penalty**

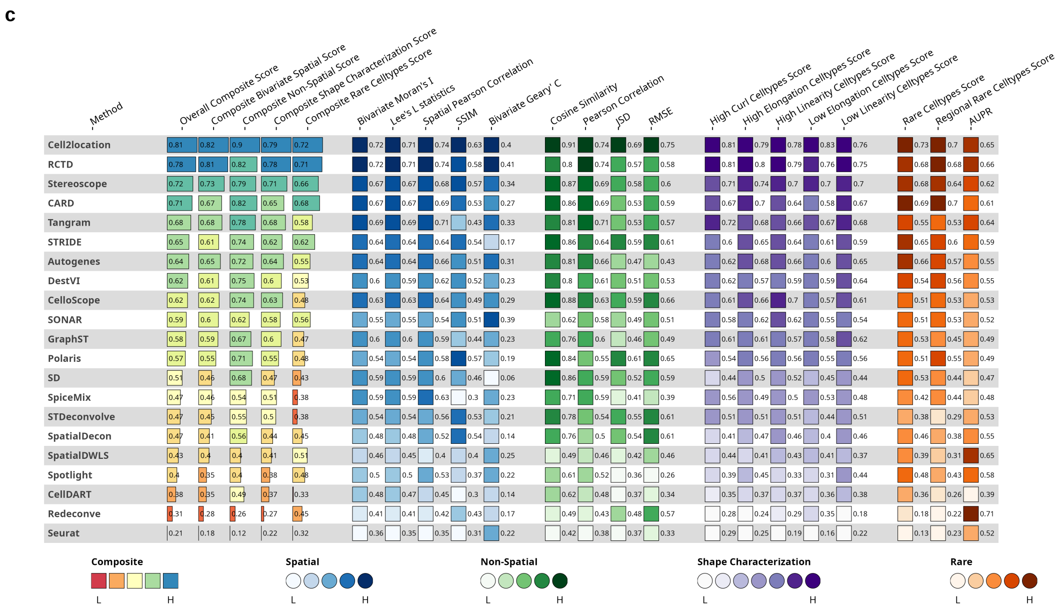


Cancer: Slope-based Penalty

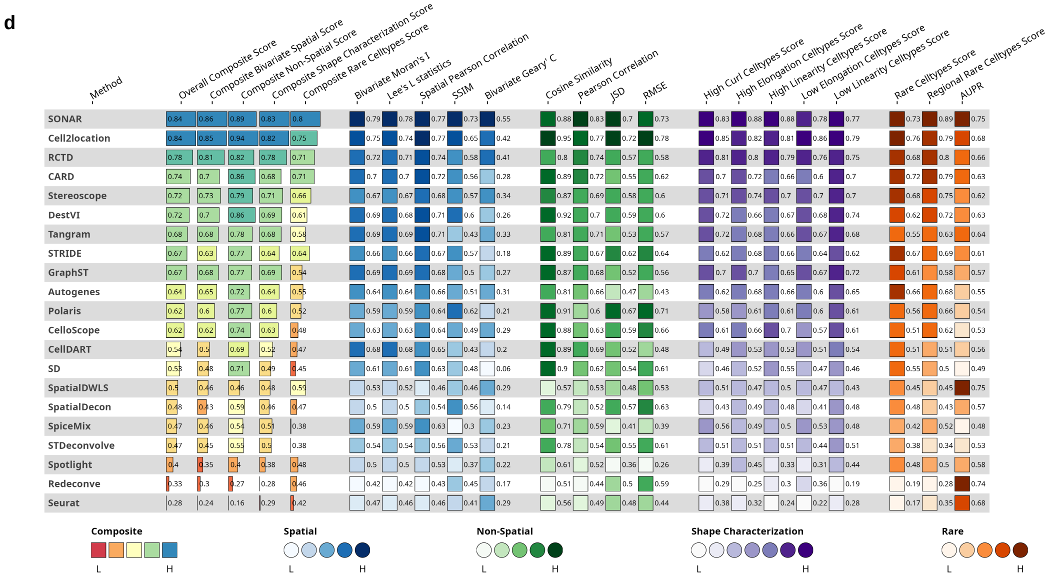


Cancer: No Penalty

**e**

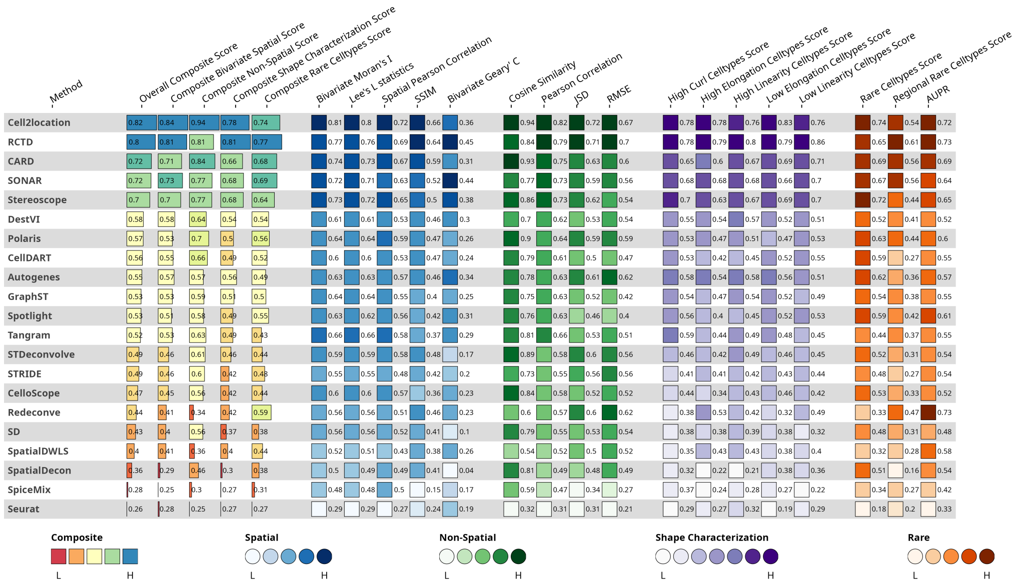


**Organs: Slope-based Penalty**

**f**

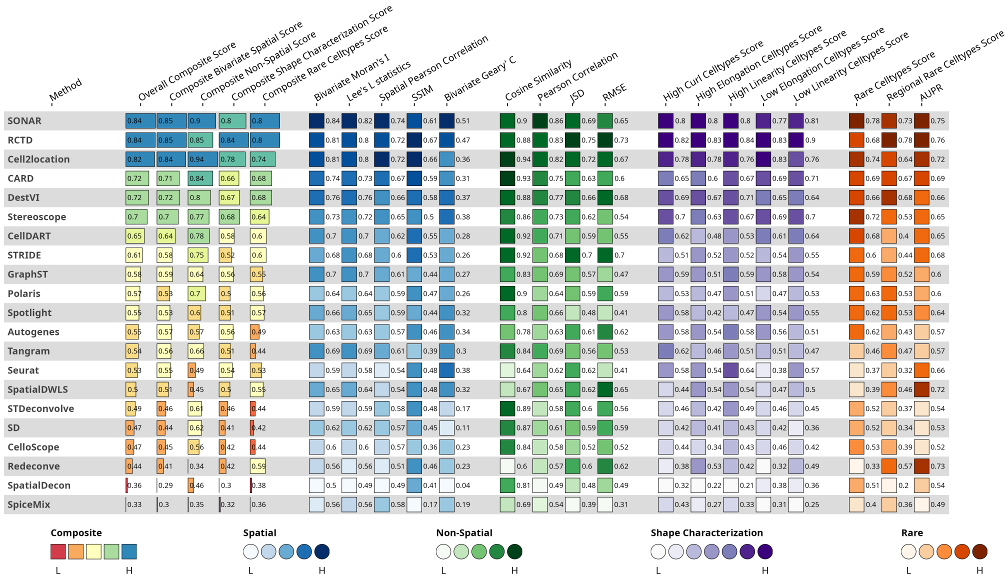


**Organs: No Penalty**

Supplementary Figure 5: Quantitative assessment of 21 deconvolution methods (Autogenes, CARD, Cell2location, CellDART, CelloScope, GraphST, Polaris, RCTD, Redeconve, Seurat, SONAR, SpatialDecon, SpatialDWLS, Spotlight, STDeconvolve, Stereoscope, STRIDE, Tangram, DestVI, SD and SpiceMix) comparing composite scores and corresponding individual metrics at the tissue level, aggregated across all datasets for each tissue type, i.e. Brain, Cancer and Organs. Heatmap visualization illustrates composite scores for a) brain datasets with slope-based penalty, b) brain datasets without penalty, c) cancer datasets with slope-based penalty, d) cancer datasets without penalty, e) organs datasets with slope-based penalty, and f) organs datasets without penalty.

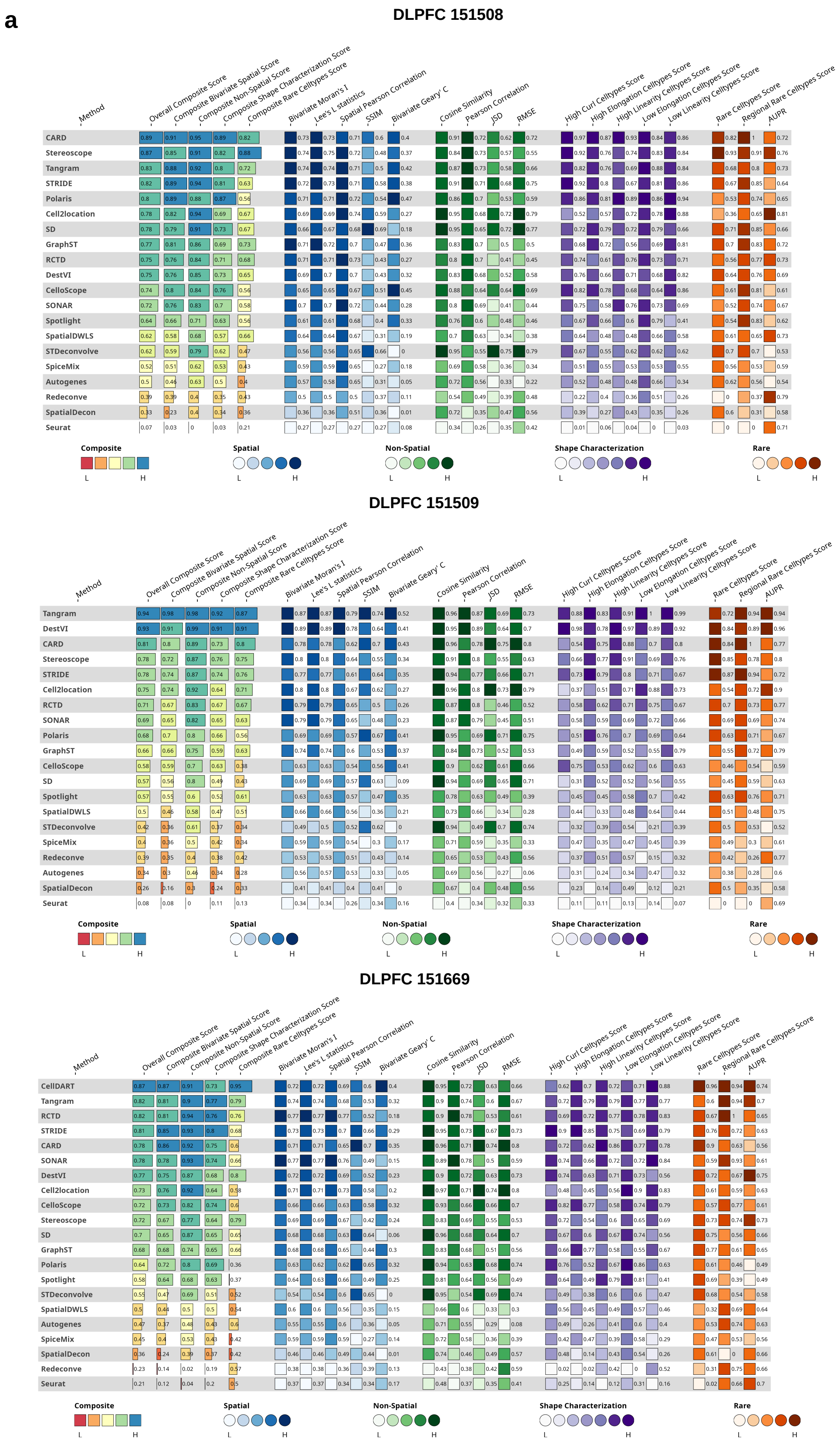
a
DLPFC 151508
DLPFC 151509
DLPFC 151669
Method
Overall Composite Score
Composite Bivariate Spatial Score
Composite Non-Spatial Score
Composite Shape Characterization Score
Composite Rare Celltypes Score
Bivariate Moran's I
Lee's L statistics
Spatial Pearson Correlation
SSIM
Bivariate Geary' C
Cosine Similarity
Pearson Correlation
JSD
RMSE
High Curl Celltypes Score
High Elongation Celltypes Score
High Linearity Celltypes Score
Low Elongation Celltypes Score
Low Linearity Celltypes Score
Rare Celltypes Score
Regional Rare Celltypes Score
AUPR
Composite
Spatial
Non-Spatial
Shape Characterization
Rare
L
H

#### DLPFC 151670

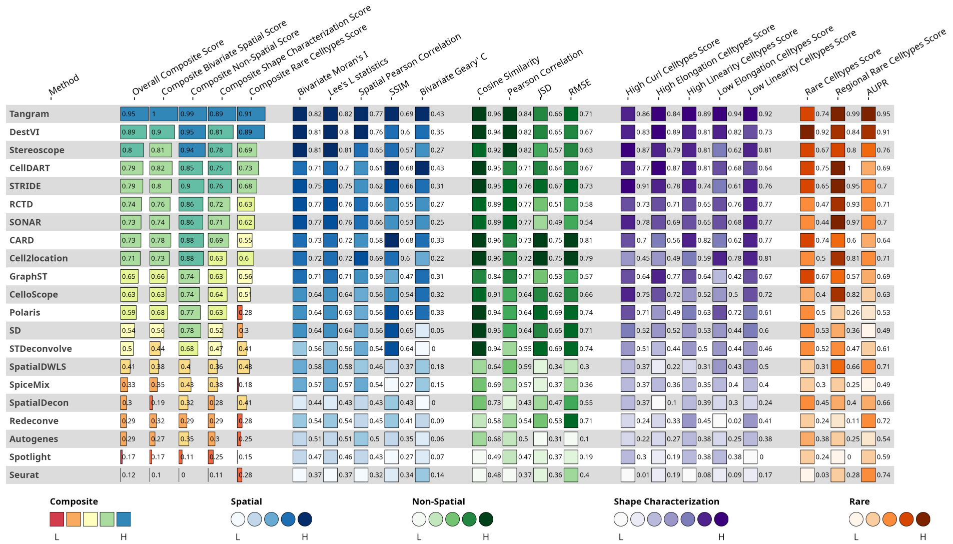


#### DLPFC 151673

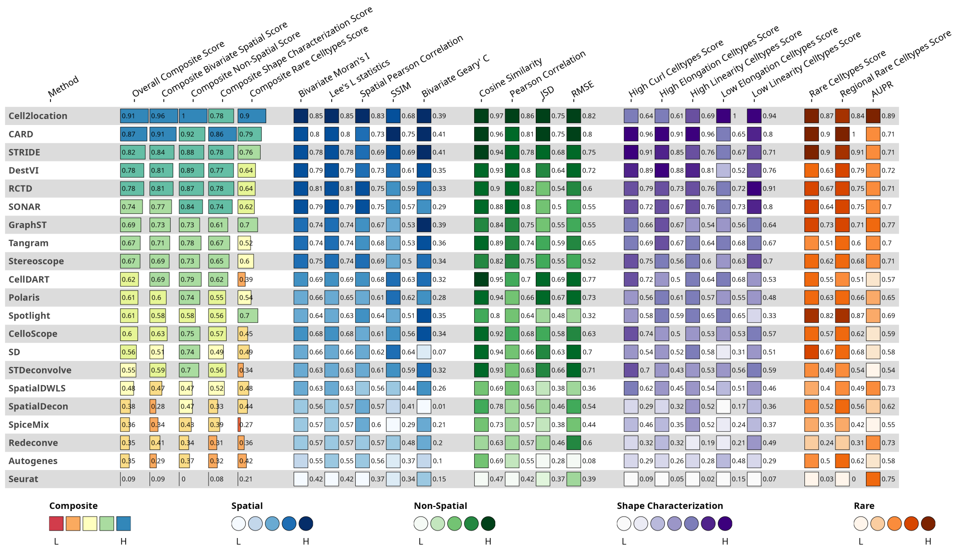


#### DLPFC 151674

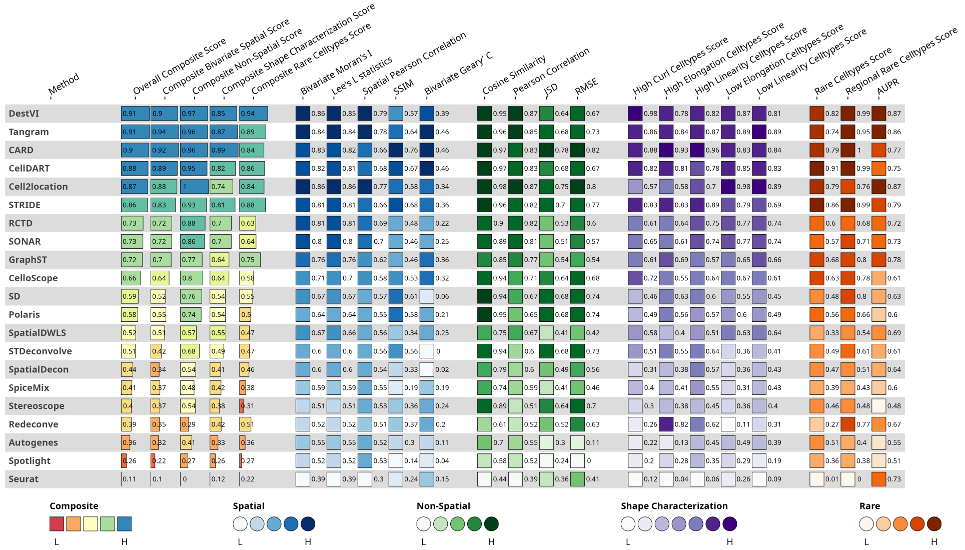

**b**

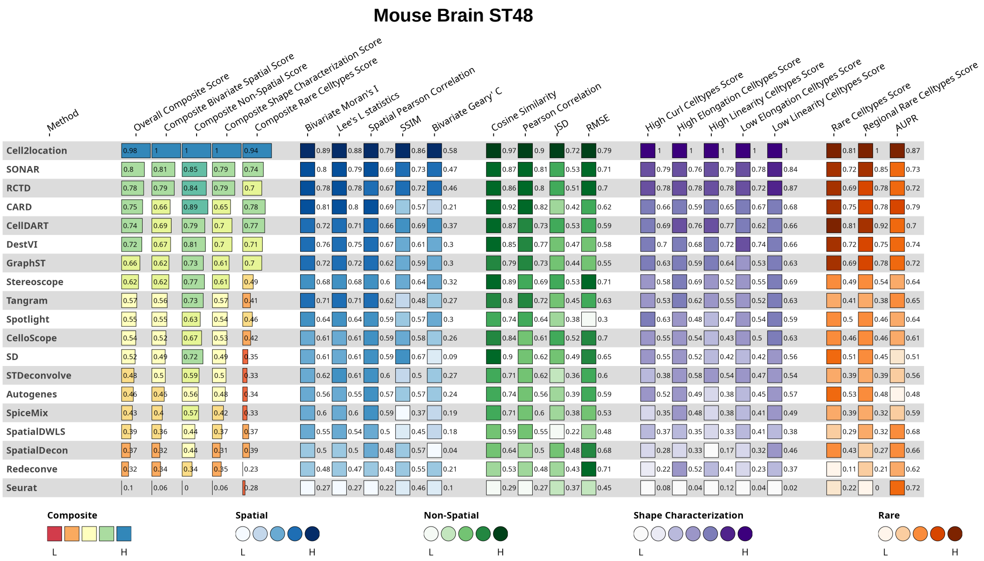
Mouse Brain ST48
Method
Overall Composite Score
Composite Bivariate Spatial Score
Composite Non-Spatial Score
Composite Shape Characterization Score
Composite Rare Celltypes Score
Bivariate Moran's I
Lee's L statistics
Spatial Pearson Correlation
SSIM
Bivariate Geary' C
Cosine Similarity
Pearson Correlation
JSD
RMSE
High Curl Celltypes Score
High Elongation Celltypes Score
High Linearity Celltypes Score
Low Elongation Celltypes Score
Low Linearity Celltypes Score
Rare Celltypes Score
Regional Rare Celltypes Score
AUPR
Cell2location
SONAR
RCTD
CARD
CellDART
DestVI
GraphST
Stereoscope
Tangram
Spotlight
CelloScope
SD
STDeconvolve
Autogenes
SpiceMix
SpatialDWLS
SpatialDecon
Redeconve
Seurat
Composite
Spatial
Non-Spatial
Shape Characterization
Rare
L
H

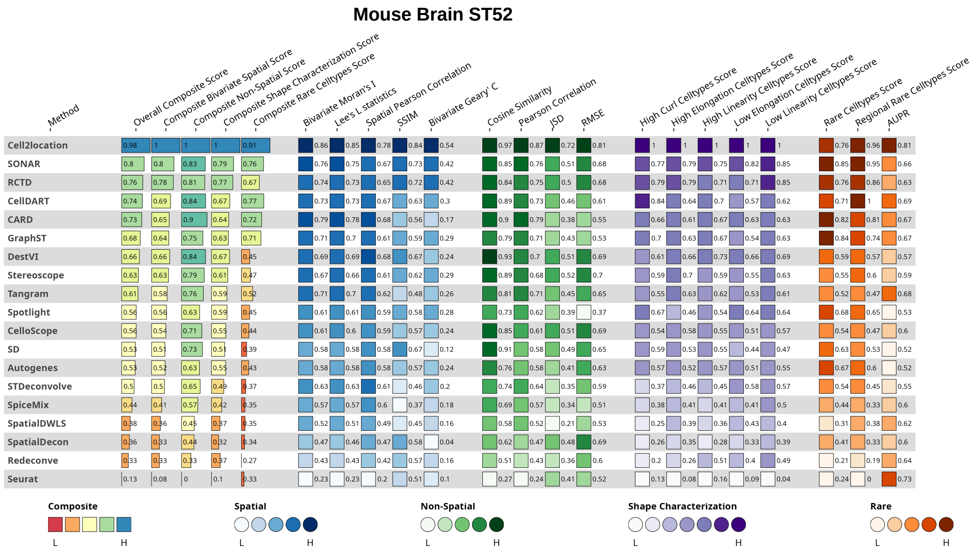
Mouse Brain ST52
Method
Overall Composite Score
Composite Bivariate Spatial Score
Composite Non-Spatial Score
Composite Shape Characterization Score
Composite Rare Celltypes Score
Bivariate Moran's I
Lee's L statistics
Spatial Pearson Correlation
SSIM
Bivariate Geary' C
Cosine Similarity
Pearson Correlation
JSD
RMSE
High Curl Celltypes Score
High Elongation Celltypes Score
High Linearity Celltypes Score
Low Elongation Celltypes Score
Low Linearity Celltypes Score
Rare Celltypes Score
Regional Rare Celltypes Score
AUPR
Cell2location
SONAR
RCTD
CellDART
CARD
GraphST
DestVI
Stereoscope
Tangram
Spotlight
CelloScope
SD
Autogenes
STDeconvolve
SpiceMix
SpatialDWLS
SpatialDecon
Redeconve
Seurat
Composite
Spatial
Non-Spatial
Shape Characterization
Rare
L
H

**c**

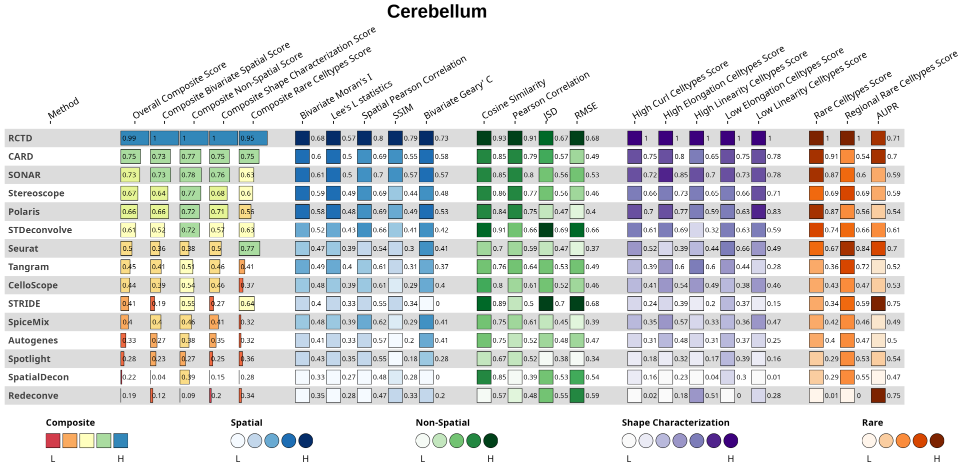
Cerebellum
Method
Overall Composite Score
Composite Bivariate Spatial Score
Composite Non-Spatial Score
Composite Shape Characterization Score
Composite Rare Celltypes Score
Bivariate Moran's I
Lee's L statistics
Spatial Pearson Correlation
SSIM
Bivariate Geary' C
Cosine Similarity
Pearson Correlation
JSD
RMSE
High Curl Celltypes Score
High Elongation Celltypes Score
High Linearity Celltypes Score
Low Elongation Celltypes Score
Low Linearity Celltypes Score
Rare Celltypes Score
Regional Rare Celltypes Score
AUPR
RCTD
CARD
SONAR
Stereoscope
Polaris
STDeconvolve
Seurat
Tangram
CelloScope
STRIDE
SpiceMix
Autogenes
Spotlight
SpatialDecon
Redeconve
Composite
Spatial
Non-Spatial
Shape Characterization
Rare
L
H

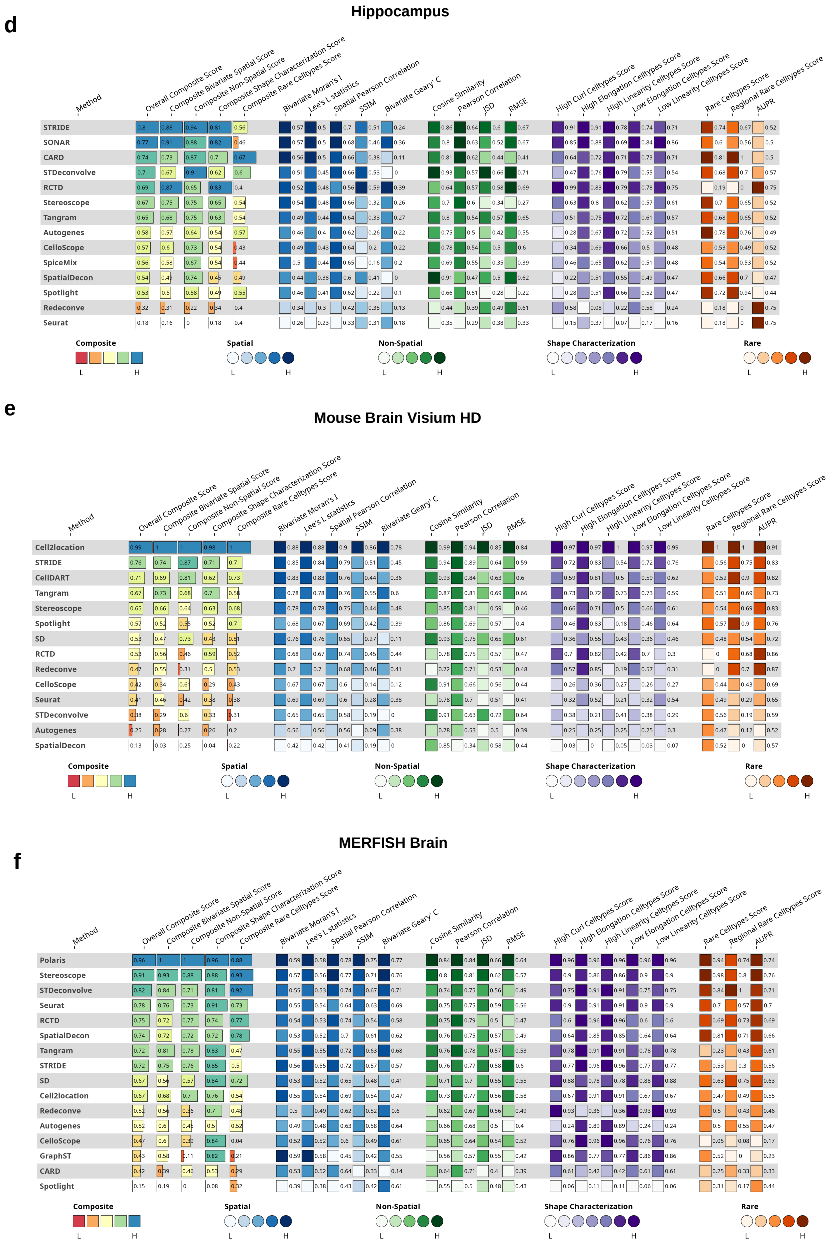


Supplementary Figure 6: Quantitative assessment of 21 deconvolution methods (Autogenes, CARD, Cell2location, CellDART, CelloScope, GraphST, Polaris, RCTD, Redeconve, Seurat, SONAR, SpatialDecon, SpatialDWLS, Spotlight, STDeconvolve, Stereoscope, STRIDE, Tangram, DestVI, SD and SpiceMix) comparing composite scores and corresponding individual metrics at the dataset level. Heatmap visualization of composite scores and corresponding individual metrics for a) DLPFC - 151508, 151509, 151669, 151670, 151673, 151674, b) Mouse Brain - ST8059048 (ST48), ST8059052 (ST52), c) Cerebellum, d) Hippocampus, e) Mouse brain Visium HD and f) Merfish Brain datasets.

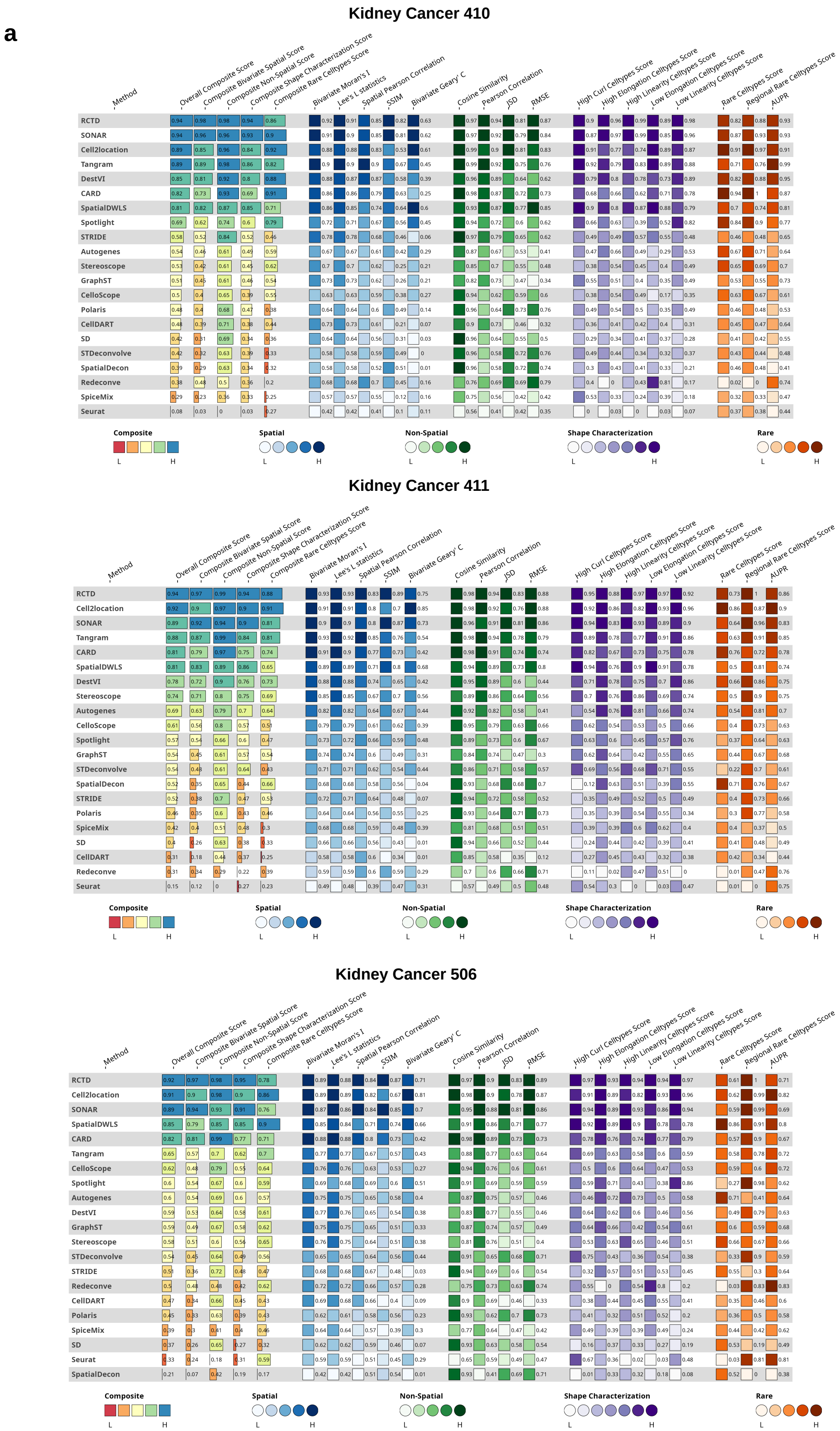
a
Kidney Cancer 410
Method
Overall Composite Score
Composite Bivariate Spatial Score
Composite Non-Spatial Score
Composite Shape Characterization Score
Composite Rare Celltypes Score
Bivariate Moran's I
Lee's L statistics
Spatial Pearson Correlation
SSIM
Bivariate Geary' C
Cosine Similarity
Pearson Correlation
JSD
RMSE
High Curl Celltypes Score
High Elongation Celltypes Score
High Linearity Celltypes Score
Low Elongation Celltypes Score
Low Linearity Celltypes Score
Rare Celltypes Score
Regional Rare Celltypes Score
AUPR
RCTD
SONAR
Cell2location
Tangram
DestVI
CARD
SpatialDWLS
Spotlight
STRIDE
Autogenes
Stereoscope
GraphST
CelloScope
Polaris
CellDART
SD
STDeconvolve
SpatialDecon
Redeconve
SpiceMix
Seurat
Composite
Spatial
Non-Spatial
Shape Characterization
Rare
L
H
Kidney Cancer 411
RCTD
Cell2location
SONAR
Tangram
CARD
SpatialDWLS
DestVI
Stereoscope
Autogenes
CelloScope
Spotlight
GraphST
STDeconvolve
SpatialDecon
STRIDE
Polaris
SpiceMix
SD
CellDART
Redeconve
Seurat
Kidney Cancer 506
RCTD
Cell2location
SONAR
SpatialDWLS
CARD
Tangram
CelloScope
Spotlight
Autogenes
DestVI
GraphST
Stereoscope
STDeconvolve
STRIDE
Redeconve
CellDART
Polaris
SpiceMix
SD
Seurat
SpatialDecon

b

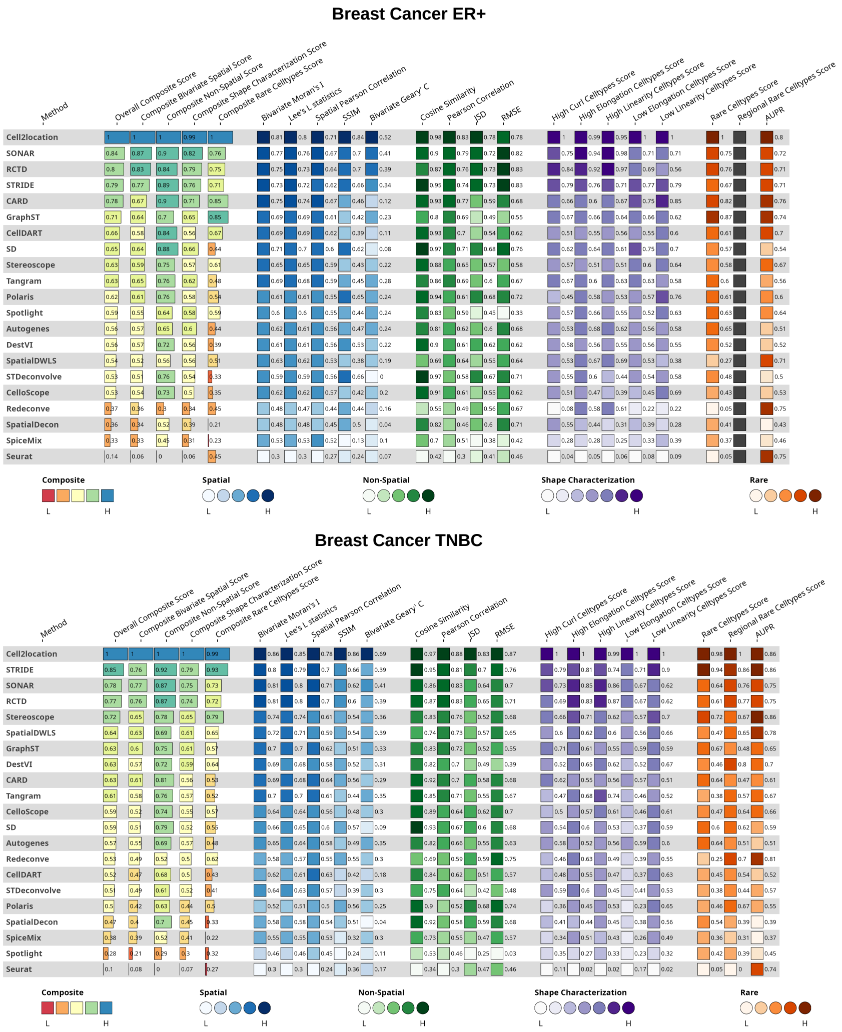
Breast Cancer ER+
Method
Overall Composite Score
Composite Bivariate Spatial Score
Composite Non-Spatial Score
Composite Shape Characterization Score
Composite Rare Celltypes Score
Bivariate Moran's I
Lee's L statistics
Spatial Pearson Correlation
SSIM
Bivariate Geary' C
Cosine Similarity
Pearson Correlation
JSD
RMSE
High Curl Celltypes Score
High Elongation Celltypes Score
High Linearity Celltypes Score
Low Elongation Celltypes Score
Low Linearity Celltypes Score
Rare Celltypes Score
Regional Rare Celltypes Score
AUPR
Cell2location
SONAR
RCTD
STRIDE
CARD
GraphST
CellDART
SD
Stereoscope
Tangram
Polaris
Spotlight
Autogenes
DestVI
SpatialDWLS
STDeconvolve
CelloScope
Redeconve
SpatialDecon
SpiceMix
Seurat
Composite
Spatial
Non-Spatial
Shape Characterization
Rare
L
H
Breast Cancer TNBC
Cell2location
STRIDE
SONAR
RCTD
Stereoscope
SpatialDWLS
GraphST
DestVI
CARD
Tangram
CelloScope
SD
Autogenes
Redeconve
CellDART
STDeconvolve
Polaris
SpatialDecon
SpiceMix
Spotlight
Seurat

c

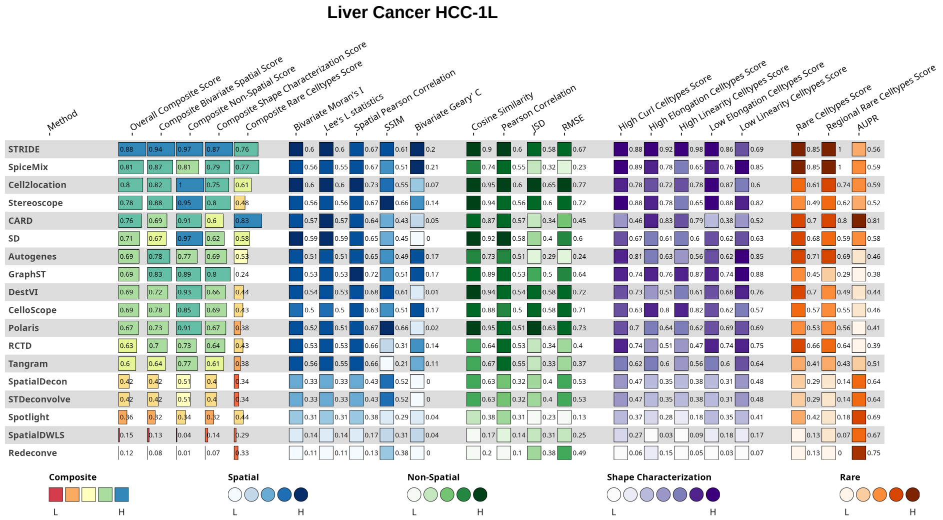
Liver Cancer HCC-1L
Method
Overall Composite Score
Composite Bivariate Spatial Score
Composite Non-Spatial Score
Composite Shape Characterization Score
Composite Rare Celltypes Score
Bivariate Moran's I
Lee's L statistics
Spatial Pearson Correlation
SSIM
Bivariate Geary' C
Cosine Similarity
Pearson Correlation
JSD
RMSE
High Curl Celltypes Score
High Elongation Celltypes Score
High Linearity Celltypes Score
Low Elongation Celltypes Score
Low Linearity Celltypes Score
Rare Celltypes Score
Regional Rare Celltypes Score
AUPR
STRIDE
SpiceMix
Cell2location
Stereoscope
CARD
SD
Autogenes
GraphST
DestVI
CelloScope
Polaris
RCTD
Tangram
SpatialDecon
STDeconvolve
Spotlight
SpatialDWLS
Redeconve
Composite
Spatial
Non-Spatial
Shape Characterization
Rare
L
H

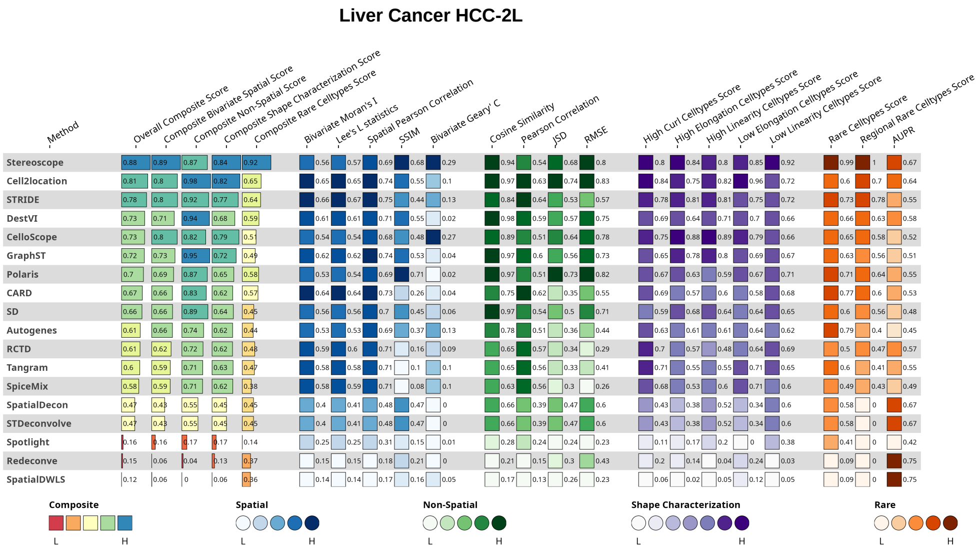
Liver Cancer HCC-2L
Method
Overall Composite Score
Composite Bivariate Spatial Score
Composite Non-Spatial Score
Composite Shape Characterization Score
Composite Rare Celltypes Score
Bivariate Moran's I
Lee's L statistics
Spatial Pearson Correlation
SSIM
Bivariate Geary' C
Cosine Similarity
Pearson Correlation
JSD
RMSE
High Curl Celltypes Score
High Elongation Celltypes Score
High Linearity Celltypes Score
Low Elongation Celltypes Score
Low Linearity Celltypes Score
Rare Celltypes Score
Regional Rare Celltypes Score
AUPR
Stereoscope
Cell2location
STRIDE
DestVI
CelloScope
GraphST
Polaris
CARD
SD
Autogenes
RCTD
Tangram
SpiceMix
SpatialDecon
STDeconvolve
Spotlight
Redeconve
SpatialDWLS
Composite
Spatial
Non-Spatial
Shape Characterization
Rare
L
H

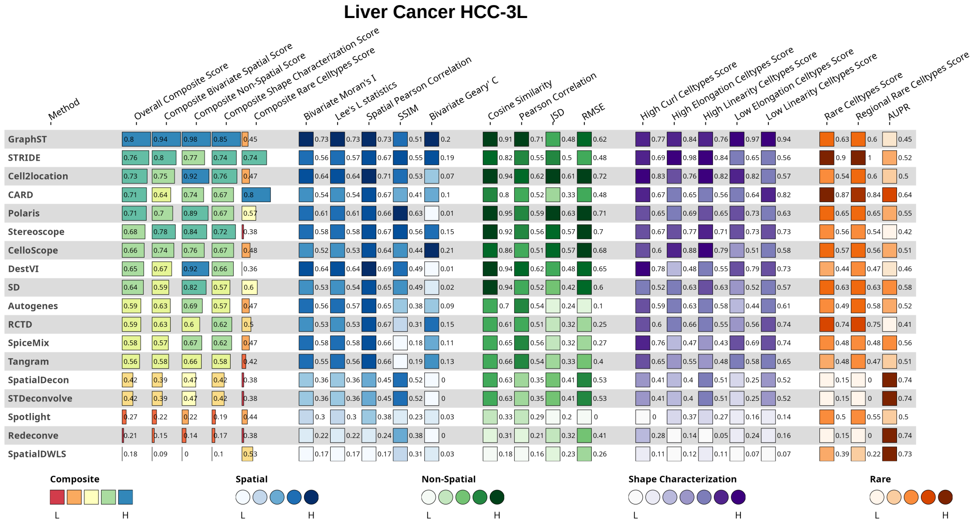
Liver Cancer HCC-3L
Method
Overall Composite Score
Composite Bivariate Spatial Score
Composite Non-Spatial Score
Composite Shape Characterization Score
Composite Rare Celltypes Score
Bivariate Moran's I
Lee's L statistics
Spatial Pearson Correlation
SSIM
Bivariate Geary' C
Cosine Similarity
Pearson Correlation
JSD
RMSE
High Curl Celltypes Score
High Elongation Celltypes Score
High Linearity Celltypes Score
Low Elongation Celltypes Score
Low Linearity Celltypes Score
Rare Celltypes Score
Regional Rare Celltypes Score
AUPR
GraphST
STRIDE
Cell2location
CARD
Polaris
Stereoscope
CelloScope
DestVI
SD
Autogenes
RCTD
SpiceMix
Tangram
SpatialDecon
STDeconvolve
Spotlight
Redeconve
SpatialDWLS
Composite
Spatial
Non-Spatial
Shape Characterization
Rare
L
H

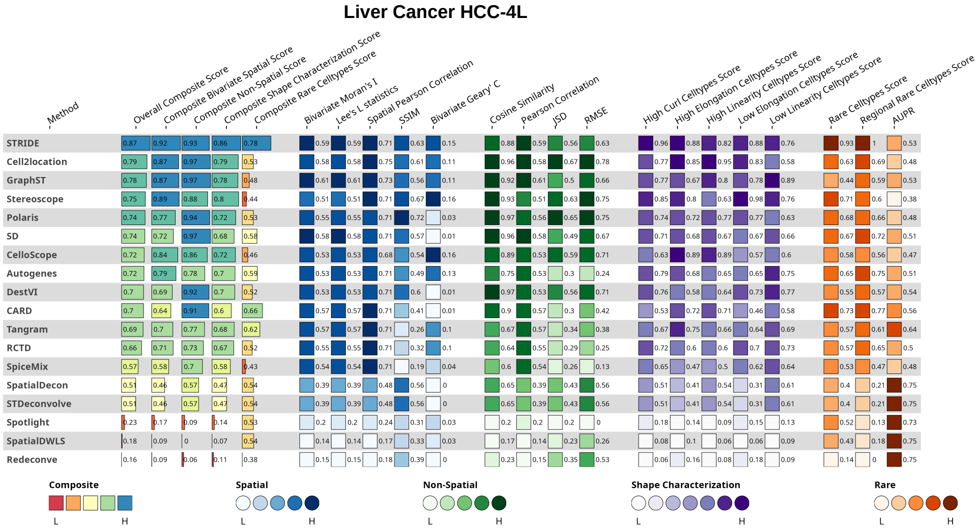
Liver Cancer HCC-4L
Method
Overall Composite Score
Composite Bivariate Spatial Score
Composite Non-Spatial Score
Composite Shape Characterization Score
Composite Rare Celltypes Score
Bivariate Moran's I
Lee's L statistics
Spatial Pearson Correlation
SSIM
Bivariate Geary' C
Cosine Similarity
Pearson Correlation
JSD
RMSE
High Curl Celltypes Score
High Elongation Celltypes Score
High Linearity Celltypes Score
Low Elongation Celltypes Score
Low Linearity Celltypes Score
Rare Celltypes Score
Regional Rare Celltypes Score
AUPR
STRIDE
Cell2location
GraphST
Stereoscope
Polaris
SD
CelloScope
Autogenes
DestVI
CARD
Tangram
RCTD
SpiceMix
SpatialDecon
STDeconvolve
Spotlight
SpatialDWLS
Redeconve
Composite
Spatial
Non-Spatial
Shape Characterization
Rare
L
H

d

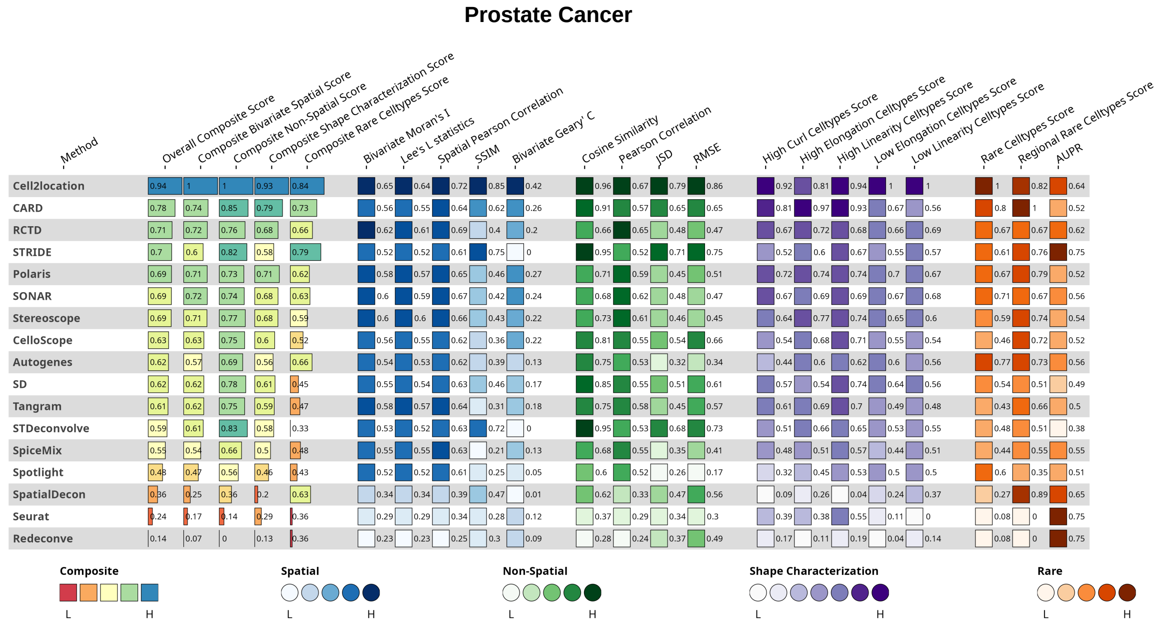
Prostate Cancer
Method
Overall Composite Score
Composite Bivariate Spatial Score
Composite Non-Spatial Score
Composite Shape Characterization Score
Composite Rare Celltypes Score
Bivariate Moran's I
Lee's L statistics
Spatial Pearson Correlation
SSIM
Bivariate Geary' C
Cosine Similarity
Pearson Correlation
JSD
RMSE
High Curl Celltypes Score
High Elongation Celltypes Score
High Linearity Celltypes Score
Low Elongation Celltypes Score
Low Linearity Celltypes Score
Rare Celltypes Score
Regional Rare Celltypes Score
AUPR
Cell2location
CARD
RCTD
STRIDE
Polaris
SONAR
Stereoscope
CelloScope
Autogenes
SD
Tangram
STDeconvolve
SpiceMix
Spotlight
SpatialDecon
Seurat
Redeconve
Composite
Spatial
Non-Spatial
Shape Characterization
Rare
L
H

e

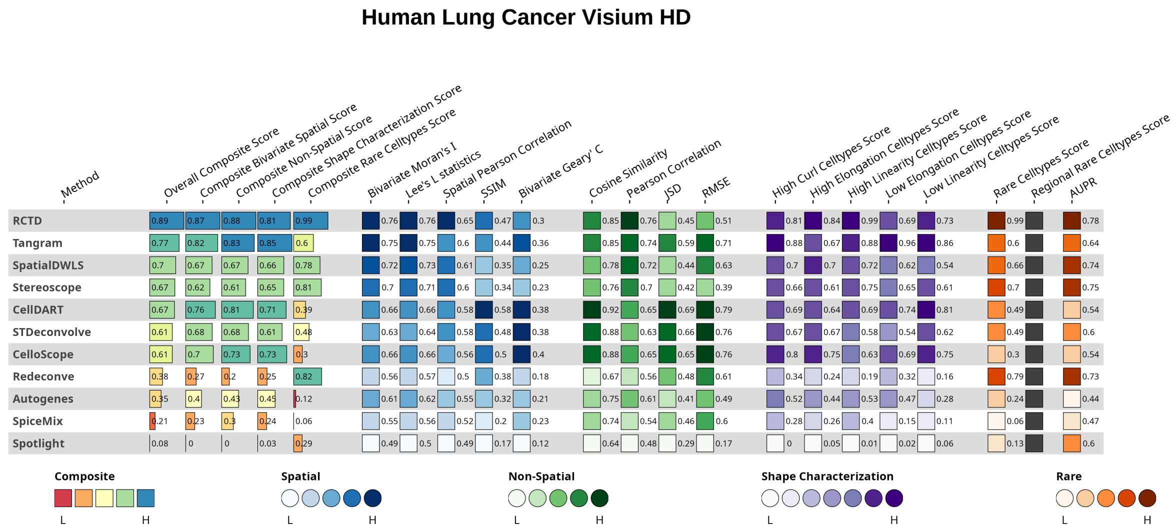
Human Lung Cancer Visium HD
Method
Overall Composite Score
Composite Bivariate Spatial Score
Composite Non-Spatial Score
Composite Shape Characterization Score
Composite Rare Celltypes Score
Bivariate Moran's I
Lee's L statistics
Spatial Pearson Correlation
SSIM
Bivariate Geary' C
Cosine Similarity
Pearson Correlation
JSD
RMSE
High Curl Celltypes Score
High Elongation Celltypes Score
High Linearity Celltypes Score
Low Elongation Celltypes Score
Low Linearity Celltypes Score
Rare Celltypes Score
Regional Rare Celltypes Score
AUPR
RCTD
Tangram
SpatialDWLS
Stereoscope
CellDART
STDeconvolve
CelloScope
Redeconve
Autogenes
SpiceMix
Spotlight
Composite
Spatial
Non-Spatial
Shape Characterization
Rare
L
H

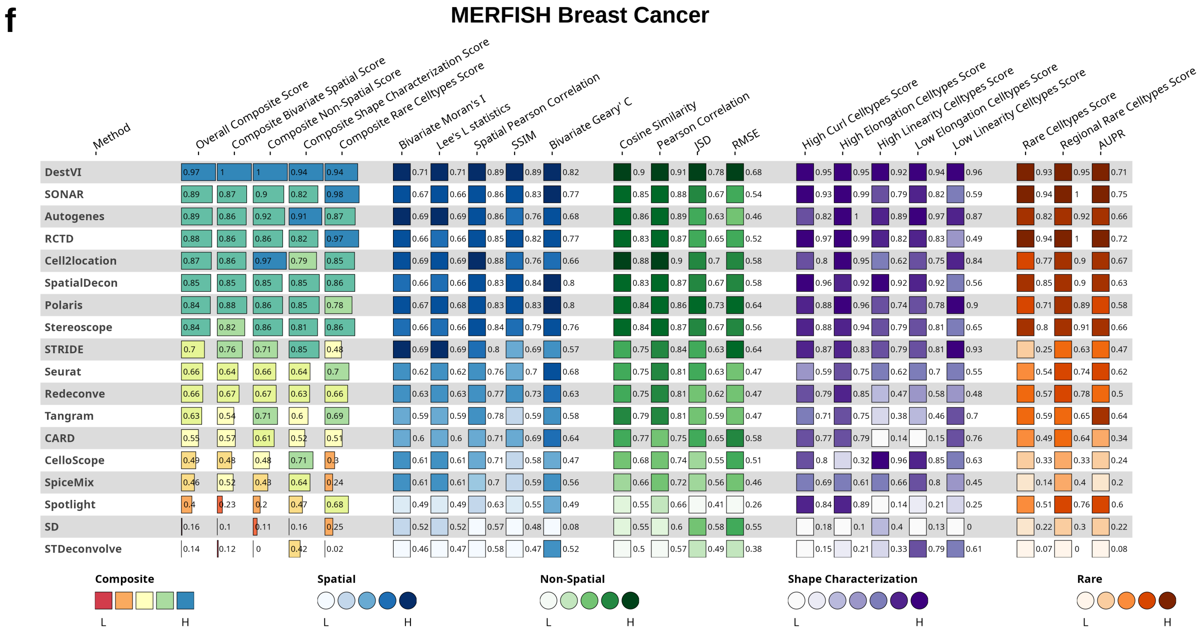


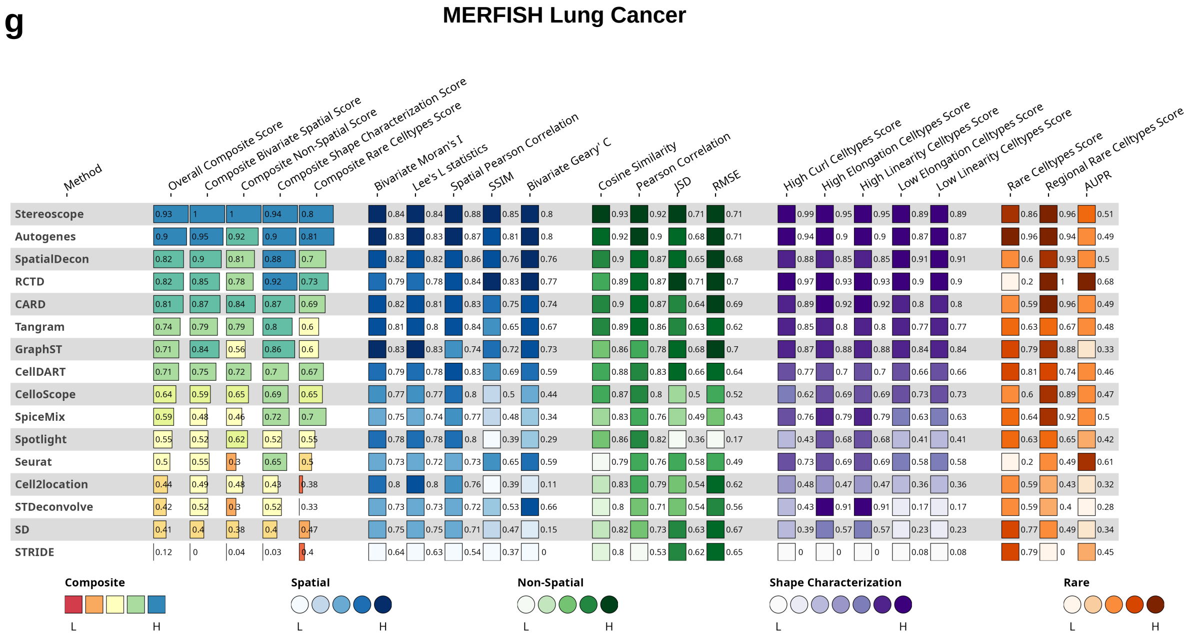


Supplementary Figure 7: Quantitative assessment of 21 deconvolution methods (Autogenes, CARD, Cell2location, CellDART, CelloScope, GraphST, Polaris, RCTD, Redeconve, Seurat, SONAR, SpatialDecon, SpatialDWLS, Spotlight, STDeconvolve, Stereoscope, STRIDE, Tangram, DestVI, SD and SpiceMix) comparing composite scores and corresponding individual metrics at the dataset level. Heatmap visualization of composite scores and corresponding individual metrics for a) Kidney Cancer - 410, 411, 506, b) Breast Cancer - ER+, TNBC, c) Liver Cancer - HCC-1L, HCC-2L, HCC-3L, HCC-4L, d) Prostate Cancer, e) Human Lung Cancer Visium HD, f) Merfish Breast cancer, and g) Merfish Lung Cancer datasets.

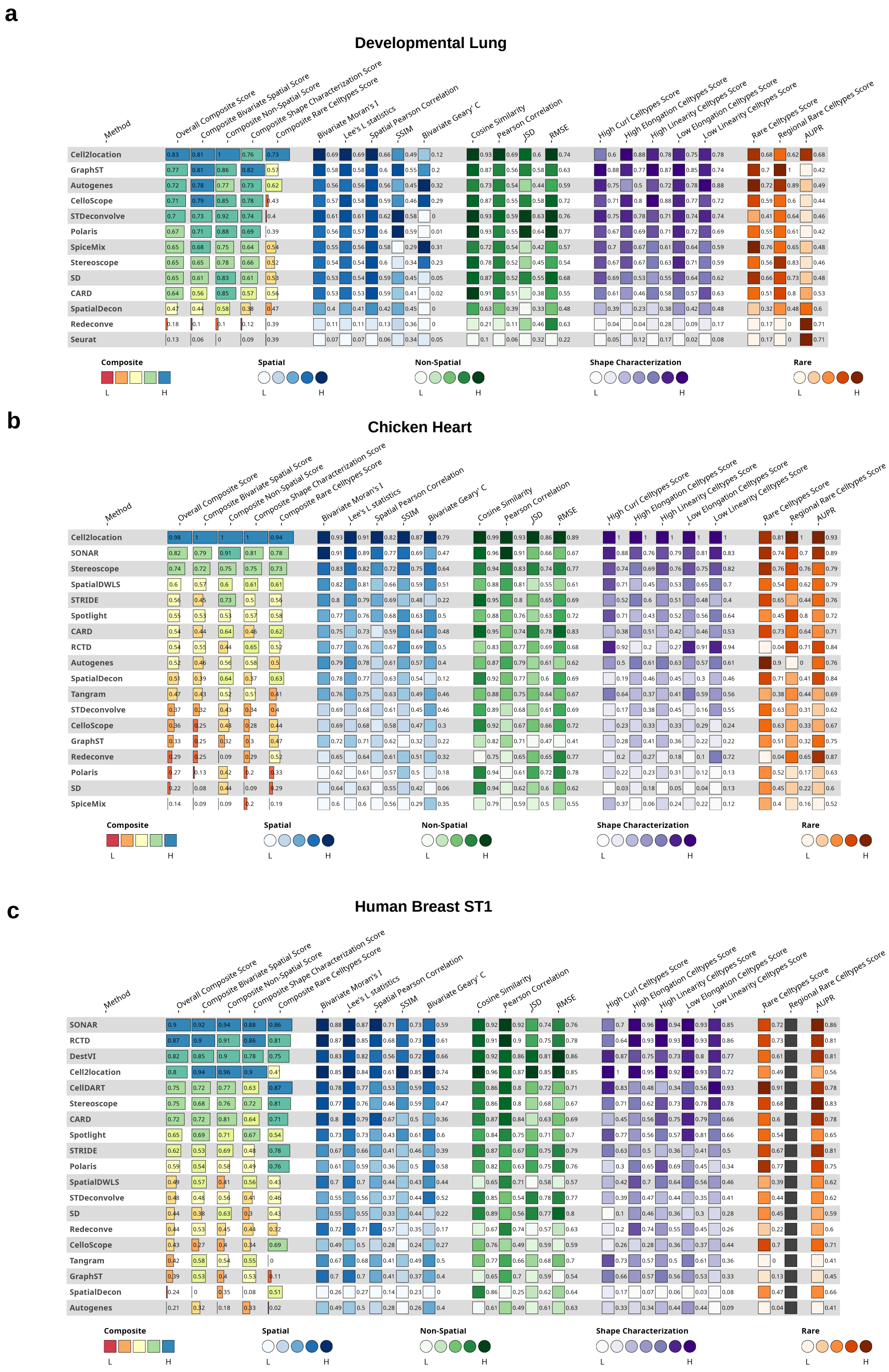
a
Developmental Lung
b
Chicken Heart
c
Human Breast ST1
Method
Overall Composite Score
Composite Bivariate Spatial Score
Composite Non-Spatial Score
Composite Shape Characterization Score
Composite Rare Celltypes Score
Bivariate Moran's I
Lee's L statistics
Spatial Pearson Correlation
SSIM
Bivariate Geary' C
Cosine Similarity
Pearson Correlation
JSD
RMSE
High Curl Celltypes Score
High Elongation Celltypes Score
High Linearity Celltypes Score
Low Elongation Celltypes Score
Low Linearity Celltypes Score
Rare Celltypes Score
Regional Rare Celltypes Score
AUPR
Composite
Spatial
Non-Spatial
Shape Characterization
Rare
L
H

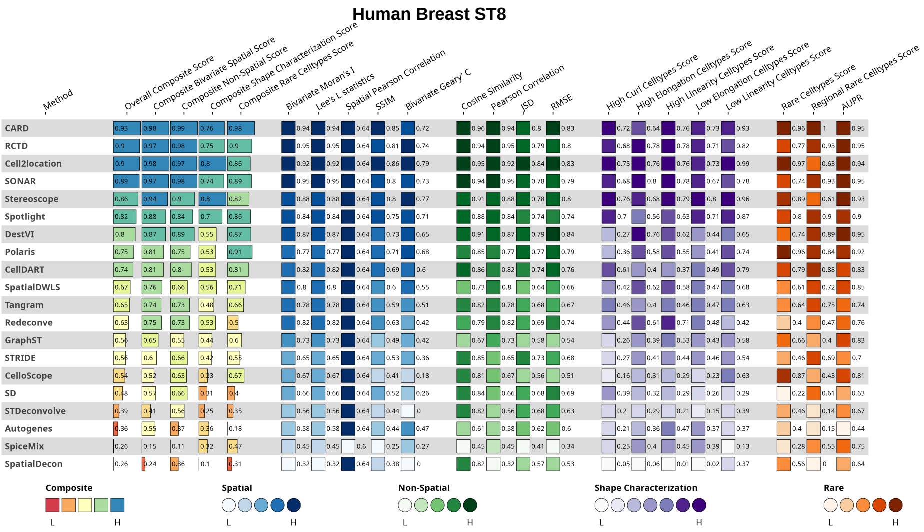
Human Breast ST8
Method
Overall Composite Score
Composite Bivariate Spatial Score
Composite Non-Spatial Score
Composite Shape Characterization Score
Composite Rare Celltypes Score
Bivariate Moran's I
Lee's L statistics
Spatial Pearson Correlation
SSIM
Bivariate Geary' C
Cosine Similarity
Pearson Correlation
JSD
RMSE
High Curl Celltypes Score
High Elongation Celltypes Score
High Linearity Celltypes Score
Low Elongation Celltypes Score
Low Linearity Celltypes Score
Rare Celltypes Score
Regional Rare Celltypes Score
AUPR
CARD
RCTD
Cell2location
SONAR
Stereoscope
Spotlight
DestVI
Polaris
CellDART
SpatialDWLS
Tangram
Redeconve
GraphST
STRIDE
CelloScope
SD
STDeconvolve
Autogenes
SpiceMix
SpatialDecon
Composite
Spatial
Non-Spatial
Shape Characterization
Rare
L
H

d

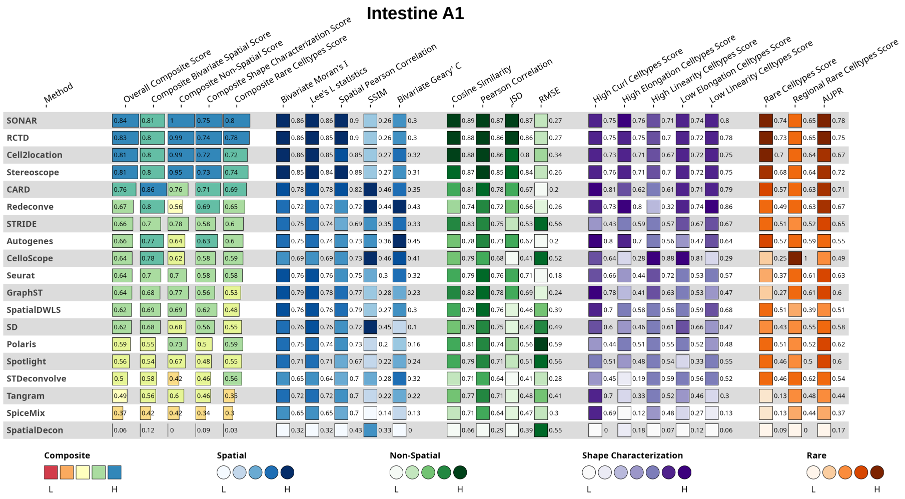
Intestine A1
Method
SONAR
RCTD
Cell2location
Stereoscope
CARD
Redeconve
STRIDE
Autogenes
CelloScope
Seurat
GraphST
SpatialDWLS
SD
Polaris
Spotlight
STDeconvolve
Tangram
SpiceMix
SpatialDecon
Composite
Spatial
Non-Spatial
Shape Characterization
Rare
L
H

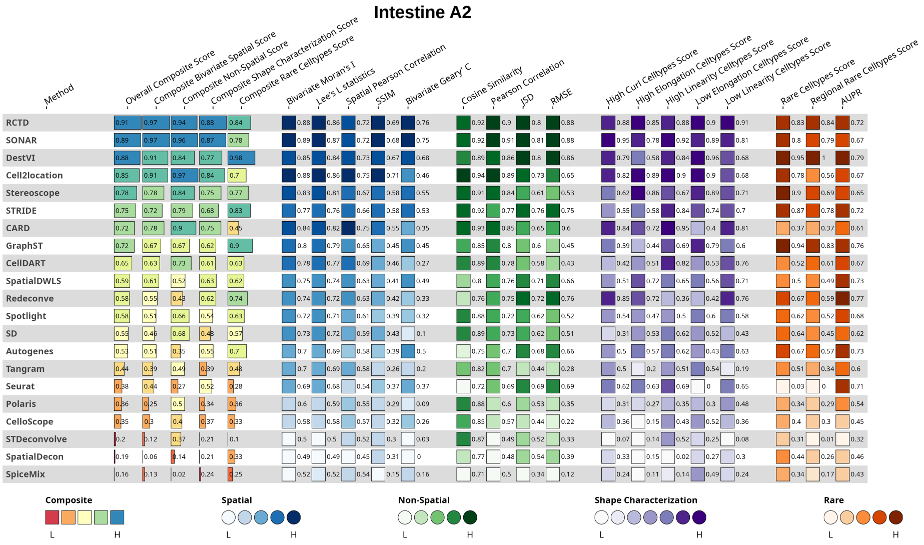
Intestine A2
Method
RCTD
SONAR
DestVI
Cell2location
Stereoscope
STRIDE
CARD
GraphST
CellDART
SpatialDWLS
Redeconve
Spotlight
SD
Autogenes
Tangram
Seurat
Polaris
CelloScope
STDeconvolve
SpatialDecon
SpiceMix
Composite
Spatial
Non-Spatial
Shape Characterization
Rare
L
H

e

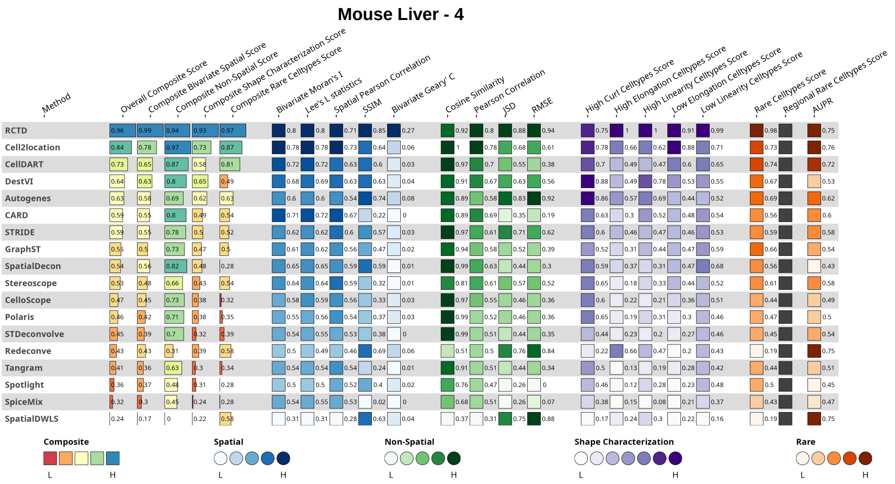
Mouse Liver - 4
Method
Overall Composite Score
Composite Bivariate Spatial Score
Composite Non-Spatial Score
Composite Shape Characterization Score
Composite Rare Celltypes Score
Bivariate Moran's I
Lee's L statistics
Spatial Pearson Correlation
SSIM
Bivariate Geary' C
Cosine Similarity
Pearson Correlation
JSD
RMSE
High Curl Celltypes Score
High Elongation Celltypes Score
High Linearity Celltypes Score
Low Elongation Celltypes Score
Low Linearity Celltypes Score
Rare Celltypes Score
Regional Rare Celltypes Score
AUPR
RCTD
Cell2location
CellDART
DestVI
Autogenes
CARD
STRIDE
GraphST
SpatialDecon
Stereoscope
CelloScope
Polaris
STDeconvolve
Redeconve
Tangram
Spotlight
SpiceMix
SpatialDWLS
Composite
Spatial
Non-Spatial
Shape Characterization
Rare
L
H

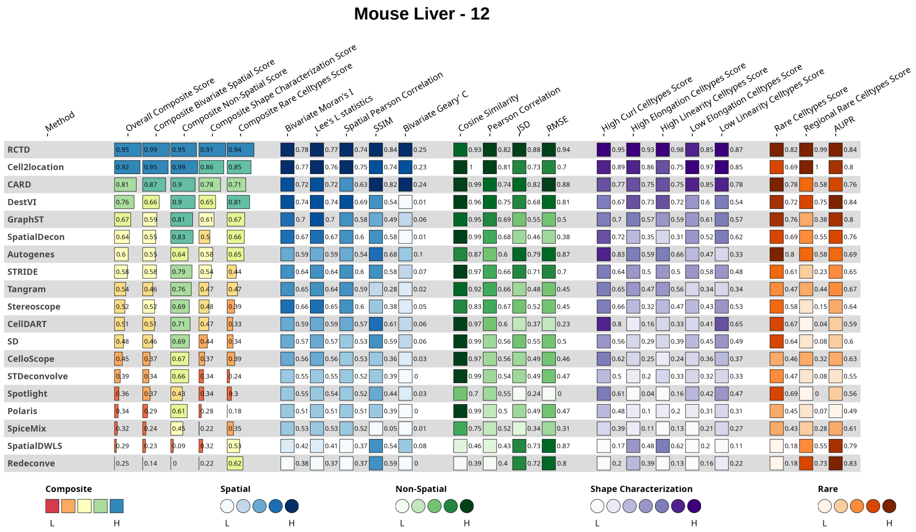
Mouse Liver - 12
Method
RCTD
Cell2location
CARD
DestVI
GraphST
SpatialDecon
Autogenes
STRIDE
Tangram
Stereoscope
CellDART
SD
CelloScope
STDeconvolve
Spotlight
Polaris
SpiceMix
SpatialDWLS
Redeconve
Composite
Spatial
Non-Spatial
Shape Characterization
Rare
L
H

f

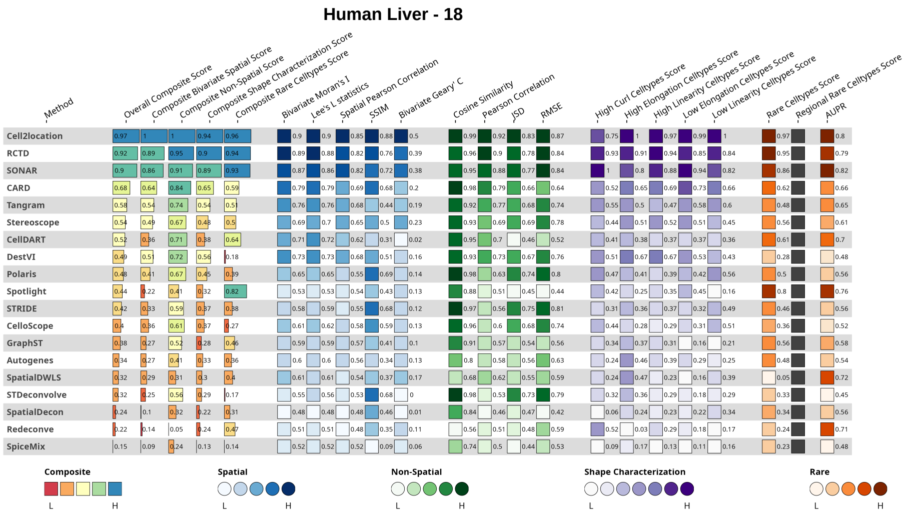
Human Liver - 18
Method
Cell2location
RCTD
SONAR
CARD
Tangram
Stereoscope
CellDART
DestVI
Polaris
Spotlight
STRIDE
CelloScope
GraphST
Autogenes
SpatialDWLS
STDeconvolve
SpatialDecon
Redeconve
SpiceMix
Composite
Spatial
Non-Spatial
Shape Characterization
Rare
L
H

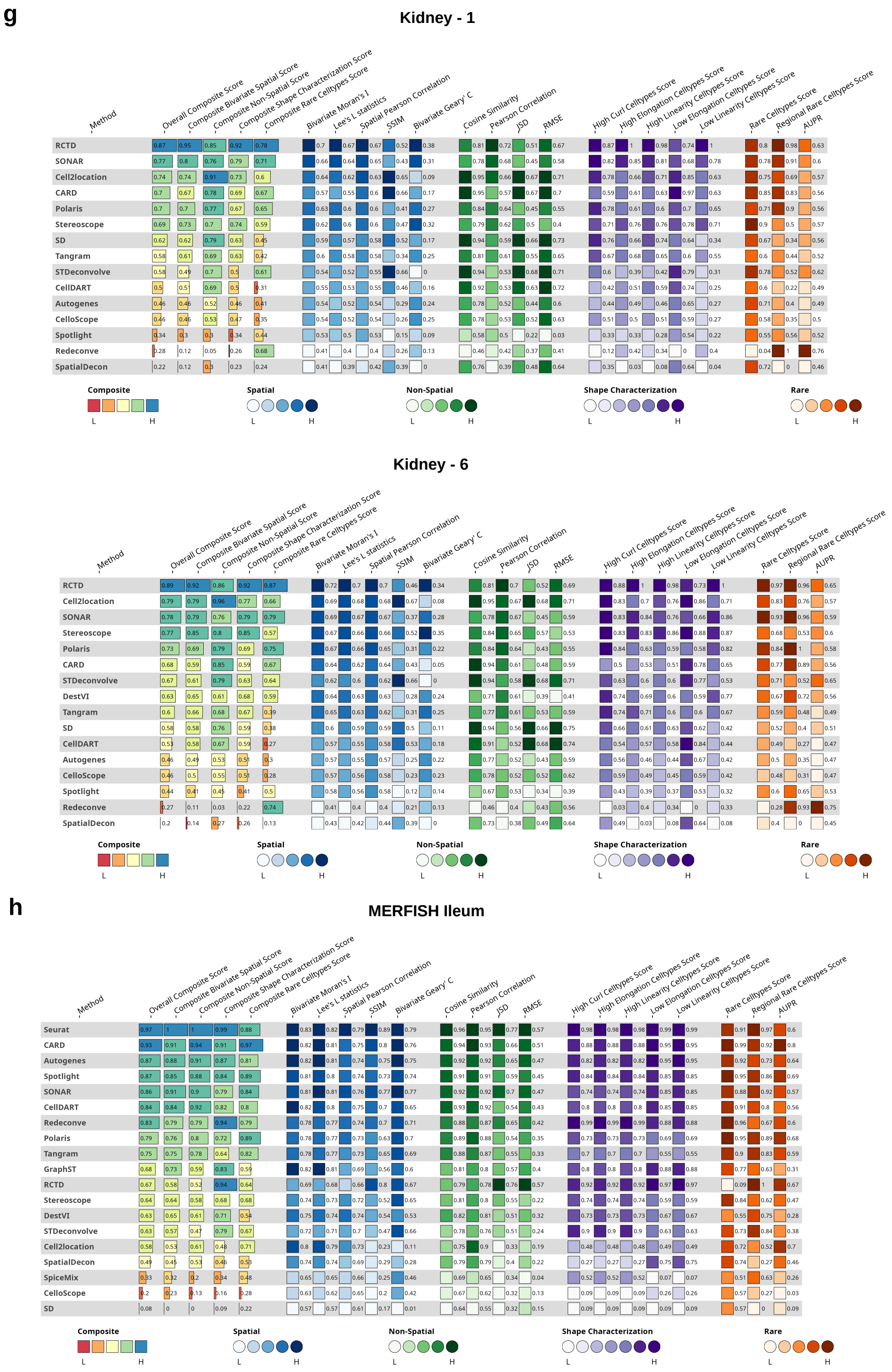
g
Kidney - 1
Method
Overall Composite Score
Composite Bivariate Spatial Score
Composite Non-Spatial Score
Composite Shape Characterization Score
Composite Rare Celltypes Score
Bivariate Moran's I
Lee's L statistics
Spatial Pearson Correlation
SSIM
Bivariate Geary' C
Cosine Similarity
Pearson Correlation
JSD
RMSE
High Curl Celltypes Score
High Elongation Celltypes Score
High Linearity Celltypes Score
Low Elongation Celltypes Score
Low Linearity Celltypes Score
Rare Celltypes Score
Regional Rare Celltypes Score
AUPR
RCTD
SONAR
Cell2location
CARD
Polaris
Stereoscope
SD
Tangram
STDeconvolve
CellDART
Autogenes
CelloScope
Spotlight
Redeconve
SpatialDecon
Composite
Spatial
Non-Spatial
Shape Characterization
Rare
L
H
Kidney - 6
RCTD
Cell2location
SONAR
Stereoscope
Polaris
CARD
STDeconvolve
DestVI
Tangram
SD
CellDART
Autogenes
CelloScope
Spotlight
Redeconve
SpatialDecon
h
MERFISH Ileum
Seurat
CARD
Autogenes
Spotlight
SONAR
CellDART
Redeconve
Polaris
Tangram
GraphST
RCTD
Stereoscope
DestVI
STDeconvolve
Cell2location
SpatialDecon
SpiceMix
CelloScope
SD

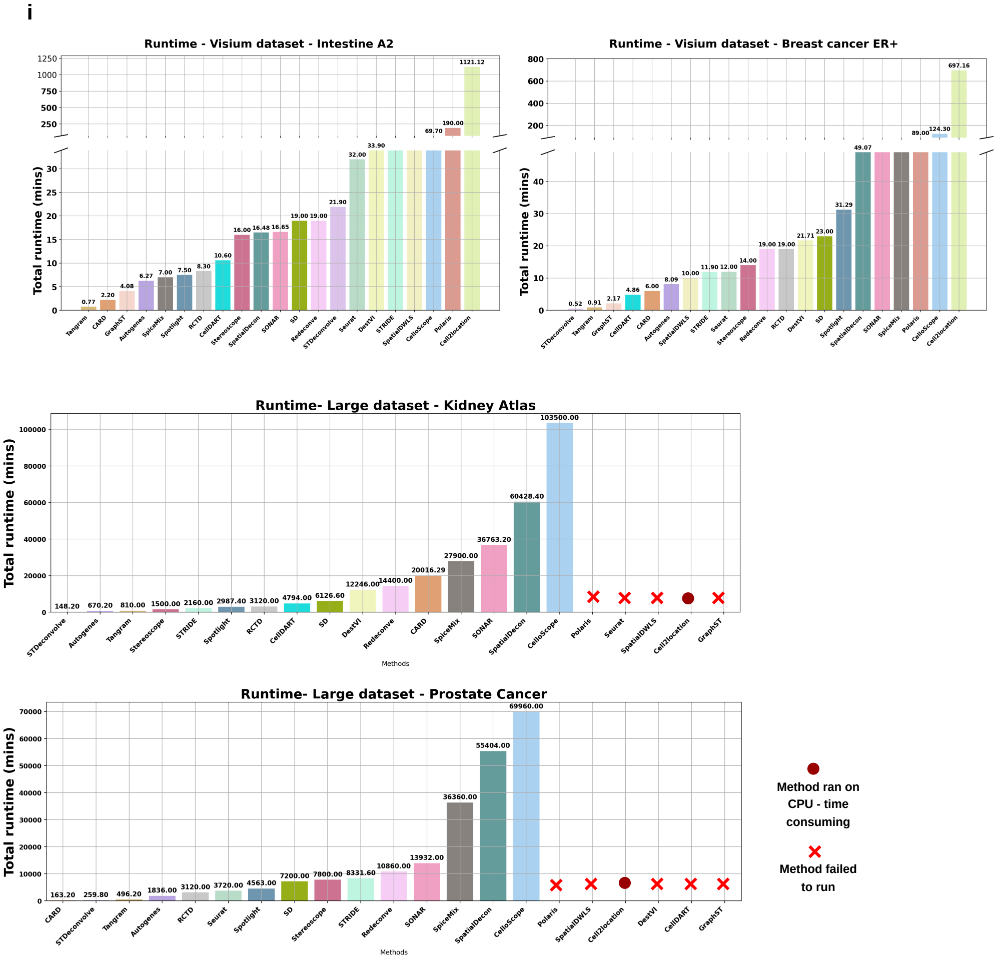


Supplementary Figure 8: Quantitative assessment of 21 deconvolution methods (Autogenes, CARD, Cell2location, CellDART, CelloScope, GraphST, Polaris, RCTD, Redeconve, Seurat, SONAR, SpatialDecon, SpatialDWLS, Spotlight, STDeconvolve, Stereoscope, STRIDE, Tangram, DestVI, SD and SpiceMix) comparing composite scores and corresponding individual metrics at the dataset level. Heatmap visualization of composite scores and corresponding individual metrics for a) Developmental Lung, b) Chicken Heart, c) Human Breast - ST_1, ST_8, d) Intestine A1, A2, e) Mouse Liver - 4, 12, f) Human Liver, g) Human Kidney 200903_1, 200903_6 and h) Merfish Ileum datasets. i) Comparison of methods in terms of total run time for Visium and SlideseqV2 datasets.

**a**

| Sno | Visium Datasets | Cell2 location | RCTD | SONAR | CARD | STRIDE | Tangram | Stereo scope | DestVI | GraphST | Cell DART | SpiceMix | Auto genes |
|---|---|---|---|---|---|---|---|---|---|---|---|---|---|
| 1 | DLPFC 151508 | | | | I | | III | II | | | | | |
| 2 | DLPFC 151509 | | | | III | | I | | II | | | | |
| 3 | DLPFC 151669 | | III | | | | II | | | | I | | |
| 4 | DLPFC 151670 | | | | | | I | III | II | | | | |
| 5 | DLPFC 151673 | I | | | II | III | | | | | | | |
| 6 | DLPFC 151674 | | | | III | | II | | I | | | | |
| 7 | Mouse Brain ST 48 | I | II | III | | | | | | | | | |
| 8 | Mouse Brain ST 52 | I | II | III | | | | | | | | | |
| 9 | Kidney Cancer 410 | III | I | II | | | | | | | | | |
| 10 | Kidney Cancer 411 | II | I | III | | | | | | | | | |
| 11 | Kidney Cancer 506 | II | I | III | | | | | | | | | |
| 12 | Breast Cancer ER+ | I | III | II | | | | | | | | | |
| 13 | Breast Cancer TNBC | I | | III | | II | | | | | | | |
| 14 | Liver Cancer 1L | III | | | | I | | | | | | II | |
| 15 | Liver Cancer 2L | II | | | | III | | I | | | | | |
| 16 | Liver Cancer 3L | III | | | | II | | | | I | | | |
| 17 | Liver Cancer 4L | II | | | | I | | | | III | | | |
| 18 | Developmental Lung | I | | | | | | | | II | | | III |
| 19 | Chicken Heart | I | | II | | | | III | | | | | |
| 20 | Breast Atlas ST1 | | II | I | | | | | III | | | | |
| 21 | Breast Atlas ST8 | III | II | | I | | | | | | | | |
| 22 | Intestine A1 | III | II | I | | | | | | | | | |
| 23 | Intestine A2 | | I | II | | | | | III | | | | |
| 24 | Liver Atlas Mouse - 4 | II | I | | | | | | | | III | | |
| 25 | Liver Atlas Mouse - 12 | II | I | | III | | | | | | | | |
| 26 | Liver Atlas Human | I | II | III | | | | | | | | | |

Visium datasets

**b**

| Sno | Large Datasets | RCTD | SONAR | Cell2 location | CARD | STRIDE | CellDART | Tangram | Spatial DWLS |
|---|---|---|---|---|---|---|---|---|---|
| 1 | Cerebellum | I | III | | II | | | | |
| 2 | Hippocampus | | II | | III | I | | | |
| 3 | Mouse Brain Visium HD | | | I | | II | III | | |
| 4 | Prostate Cancer | III | | I | II | | | | |
| 5 | Lung Cancer Visium HD | I | | | | | | II | III |
| 6 | Kidney Atlas 1 | I | II | III | | | | | |
| 7 | Kidney Atlas 6 | I | III | II | | | | | |

Large datasets

**c**

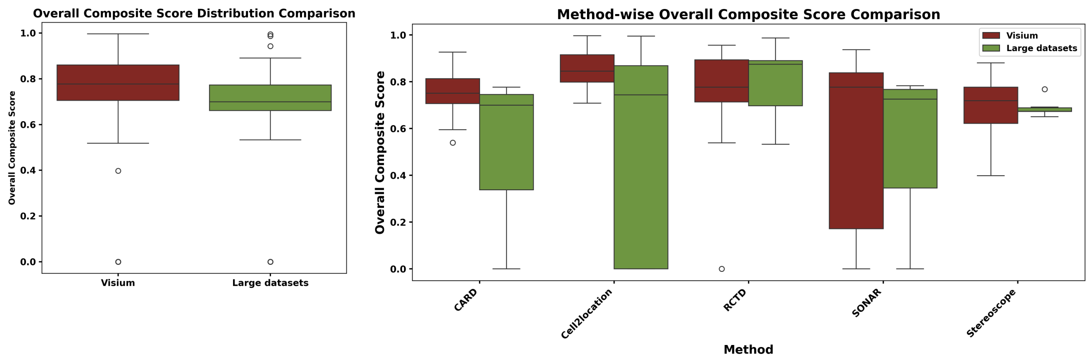

d

| Sno | Simulation Strategy 2 datasets | Auto genes | Stereo scope | SONAR | DestVI | CARD | Polaris | SpatialDecon | Seurat | STDeconvolve |
|---|---|---|---|---|---|---|---|---|---|---|
| 1 | MERFISH Brain | | II | | | | I | | | III |
| 2 | MERFISH Breast Cancer | III | | II | I | | | | | |
| 3 | MERFISH Lung Cancer | II | I | | | | | III | | |
| 4 | MERFISH Ileum | III | | | | II | | | I | |

Simulation Strategy 2 datasets

Supplementary Figure 9: a-b) Summary tables showing datasets (rows) and deconvolution methods (columns) that ranked among the top three performers classified by technology type: (a) Visium datasets and (b) Large datasets. c) Boxplot comparison of composite scores between Visium datasets (n = 26) and large-scale datasets (n = 7), showing the overall composite score (left) and method-wise composite scores (right). d) Summary tables showing datasets (rows) and deconvolution methods (columns) that ranked among the top three performers for Simulation Strategy 2 datasets.

**a**

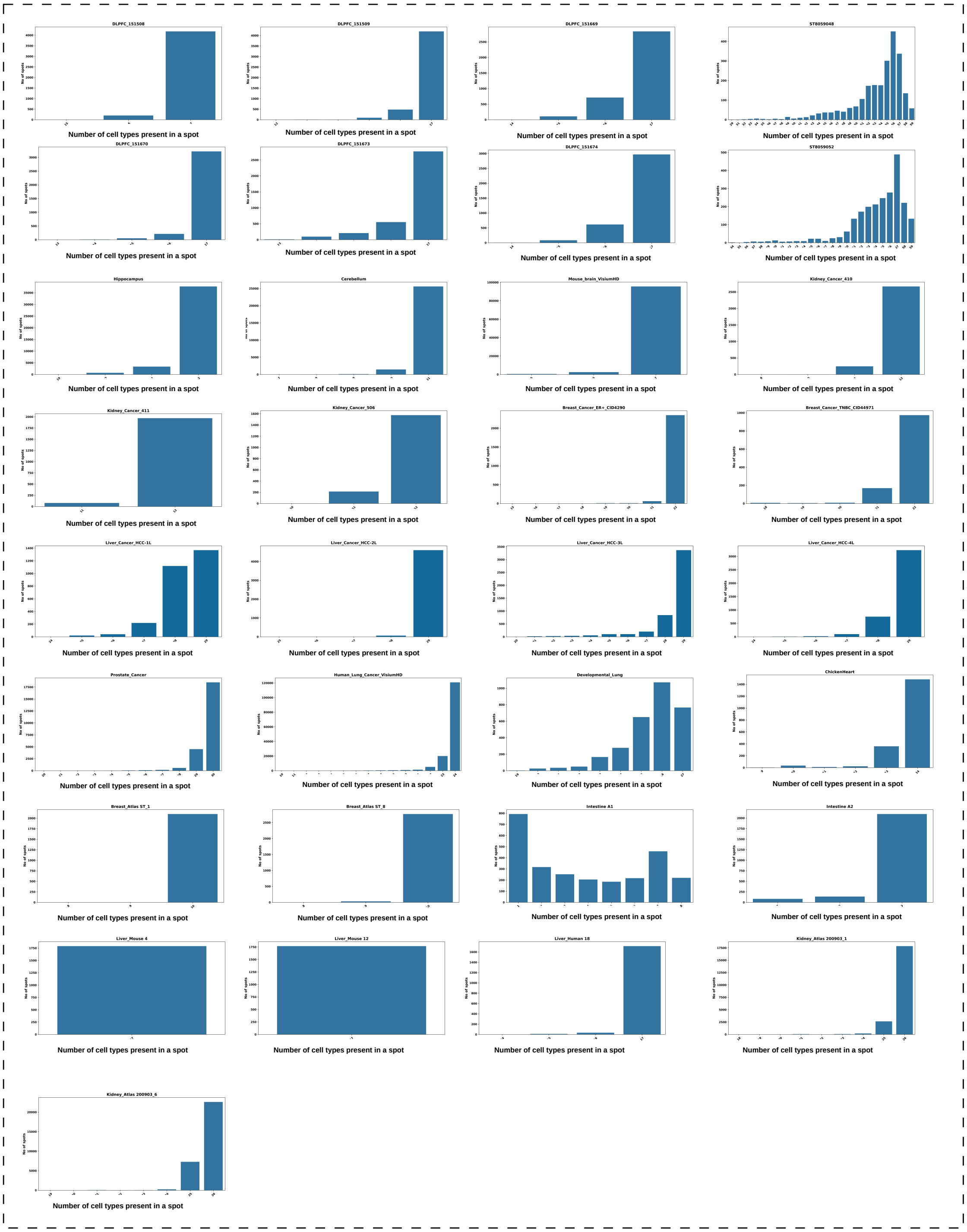

DLPFC_151508
DLPFC_151509
DLPFC_151669
ST8059048
DLPFC_151670
DLPFC_151673
DLPFC_151674
ST8059052
Hippocampus
Cerebellum
Mouse_brain_VisiumHD
Kidney_Cancer_410
Kidney_Cancer_411
Kidney_Cancer_506
Breast_Cancer_ER+_CID4290
Breast_Cancer_TNBC_CID44971
Liver_Cancer_HCC-1L
Liver_Cancer_HCC-2L
Liver_Cancer_HCC-3L
Liver_Cancer_HCC-4L
Prostate_Cancer
Human_Lung_Cancer_VisiumHD
Developmental_Lung
ChickenHeart
Breast_Atlas ST_1
Breast_Atlas ST_8
Intestine A1
Intestine A2
Liver_Mouse 4
Liver_Mouse 12
Liver_Human 18
Kidney_Atlas 200903_1
Kidney_Atlas 200903_6
No of spots
Number of cell types present in a spot

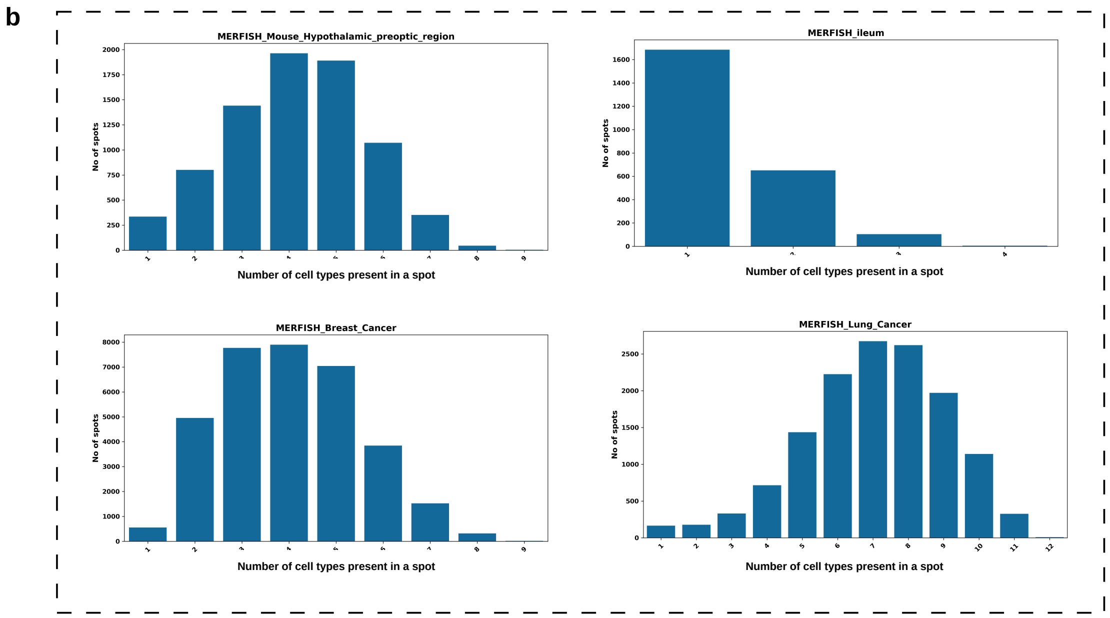


Supplementary Figure 10: Histogram showing the distribution of the number of spots and the number of cell types present in the spot for a) simulated datasets generated using simulation strategy 1 for sequencing-based datasets (i.e. 151508, 151509, 151669, 151670, 151673 and 151674, Mouse brain ST48, Mouse brain ST52, Hippocampus, Cerebellum, Mouse brain Visium HD dataset, Kidney Cancer 410, Kidney Cancer 411, Kidney Cancer 506, Breast Cancer ER+, Breast Cancer TNBC, Liver Cancer HCC-1L, HCC-2L, HCC-3L, HCC-4L, Prostate Cancer, Human Lung Cancer Visium HD, Developmental Lung, Chicken Heart, Human Breast ST_1, Human Breast ST_8, Intestine A1, Intestine A2, Liver Mouse - 4, Liver Mouse - 12, Liver Human - 18, Human Kidney - 200903_1 and Human Kidney - 200903_6) and b) simulated datasets generated using simulation strategy 2 for Imaging-based datasets (i.e. Merfish Breast Cancer, Merfish Ileum, Merfish Lung Cancer, Merfish Brain)

a

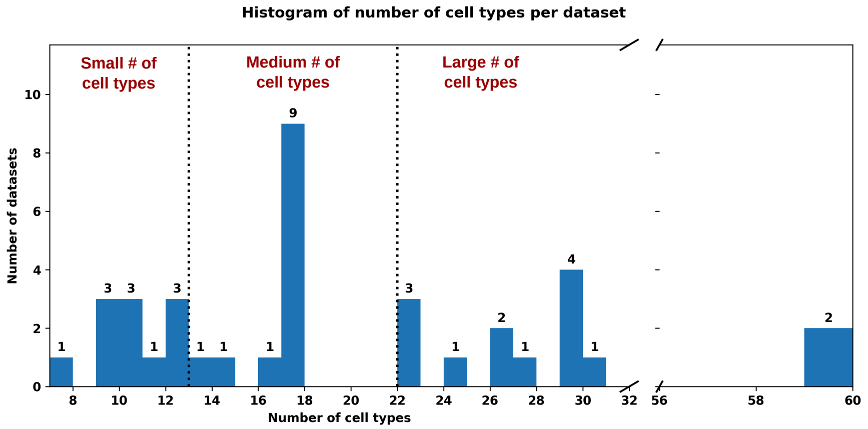


b

| Sno | Datasets | # cell types | Cell2 location | RCTD | SONAR | CARD | STRIDE | Stereo scope | DestVI | Polaris | Cell DART | STDecon volve | Auto genes |
|---|---|---|---|---|---|---|---|---|---|---|---|---|---|
| 1 | Kidney Cancer 410 | 12 | III | I | II | | | | | | | | |
| 2 | Kidney Cancer 411 | 12 | II | I | III | | | | | | | | |
| 3 | Kidney Cancer 506 | 12 | II | I | III | | | | | | | | |
| 4 | Breast Atlas ST1 | 10 | | II | I | | | | III | | | | |
| 5 | Breast Atlas ST8 | 10 | III | II | | I | | | | | | | |
| 6 | Intestine A1 | 9 | III | II | I | | | | | | | | |
| 7 | Intestine A2 | 9 | | I | II | | | | III | | | | |
| 8 | Cerebellum | 11 | | I | III | II | | | | | | | |
| 9 | Mouse Brain Visium HD | 7 | I | | | | II | | | | III | | |
| 10 | MERFISH Brain | 10 | | | | | | II | | I | | III | |
| 11 | MERFISH Breast Cancer | 9 | | | II | | | | I | | | | III |

Datasets with small number of cell types

c

| Sno | Datasets | # cell types | Cell2 location | RCTD | SONAR | CARD | STRIDE | Tang ram | Stereo scope | DestVI | Seurat | Cell DART | Spatial Decon | Auto genes |
|---|---|---|---|---|---|---|---|---|---|---|---|---|---|---|
| 1 | DLPFC 151508 | 17 | | | | I | | III | II | | | | | |
| 2 | DLPFC 151509 | 17 | | | | III | | I | | II | | | | |
| 3 | DLPFC 151669 | 17 | | III | | | | II | | | | I | | |
| 4 | DLPFC 151670 | 17 | | | | | | I | III | II | | | | |
| 5 | DLPFC 151673 | 17 | I | | | II | III | | | | | | | |
| 6 | DLPFC 151674 | 17 | | | | III | | II | | I | | | | |
| 7 | Chicken Heart | 14 | I | | II | | | | III | | | | | |
| 8 | Liver Atlas Mouse - 4 | 17 | II | I | | | | | | | | III | | |
| 9 | Liver Atlas Mouse - 12 | 17 | II | I | | III | | | | | | | | |
| 10 | Liver Atlas Human | 17 | I | II | III | | | | | | | | | |
| 11 | MERFISH Lung Cancer | 13 | | | | | | | I | | | | III | II |
| 12 | MERFISH Ileum | 16 | | | | II | | | | | I | | | III |

Datasets with medium number of cell types

**d**

| Sno | Datasets | # celltypes | Cell2 location | RCTD | SONAR | CARD | STRIDE | Tan gram | Stereo scope | GraphST | Spatial DWLS | Spice Mix | Auto genes |
|---|---|---|---|---|---|---|---|---|---|---|---|---|---|
| 1 | Mouse Brain ST 48 | 59 | I | II | III | | | | | | | | |
| 2 | Mouse Brain ST 52 | 59 | I | II | III | | | | | | | | |
| 3 | Breast Cancer ER+ | 22 | I | III | II | | | | | | | | |
| 4 | Breast Cancer TNBC | 22 | I | | III | | II | | | | | | |
| 5 | Liver Cancer 1L | 29 | III | | | | I | | | | | II | |
| 6 | Liver Cancer 2L | 29 | II | | | | III | | I | | | | |
| 7 | Liver Cancer 3L | 29 | III | | | | II | | | I | | | |
| 8 | Liver Cancer 4L | 29 | II | | | | I | | | III | | | |
| 9 | Developmental Lung | 27 | I | | | | | | | II | | | III |
| 10 | Hippocampus | 22 | | | II | III | I | | | | | | |
| 11 | Prostate Cancer | 30 | I | III | | II | | | | | | | |
| 12 | Lung Cancer Visium | 24 | | I | | | | II | | | III | | |
| 13 | Kidney Atlas 1 | 26 | III | I | II | | | | | | | | |
| 14 | Kidney Atlas 6 | 26 | II | I | III | | | | | | | | |

Datasets with large number of cell types

**e**

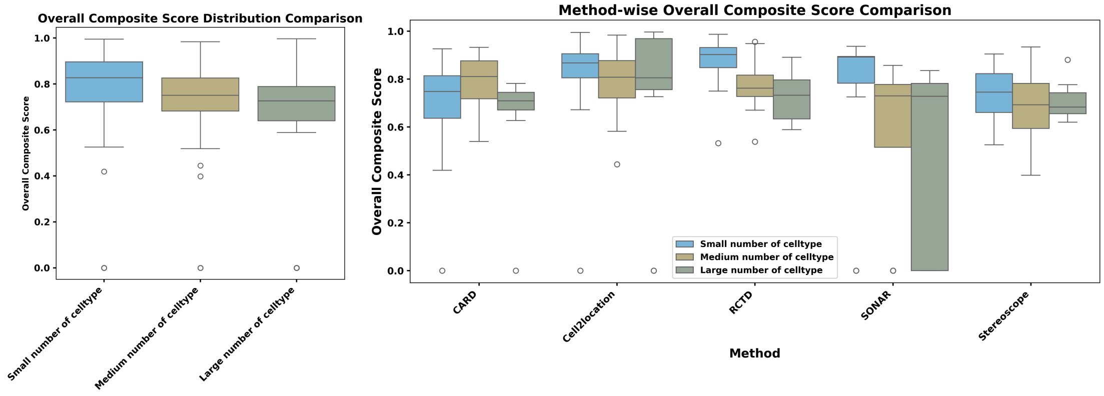


Supplementary Figure 11: (a) Histogram illustrating the distribution of cell types across all datasets. b-d) Summary tables showing datasets (rows) and deconvolution methods (columns) that ranked among the top three performers classified by different groups of cell-type distribution, b) Small number of cell types, c) Medium number of cell types, d) Large number of cell types. e) Boxplot comparison of composite scores between dataset groups based on the number of cell types: Small(0-12 cell types, n = 11), Medium(13-21 cell types, n=12) and Large($\geq$ 22, n = 14), showing the overall composite score (left) and method-wise composite scores (right).

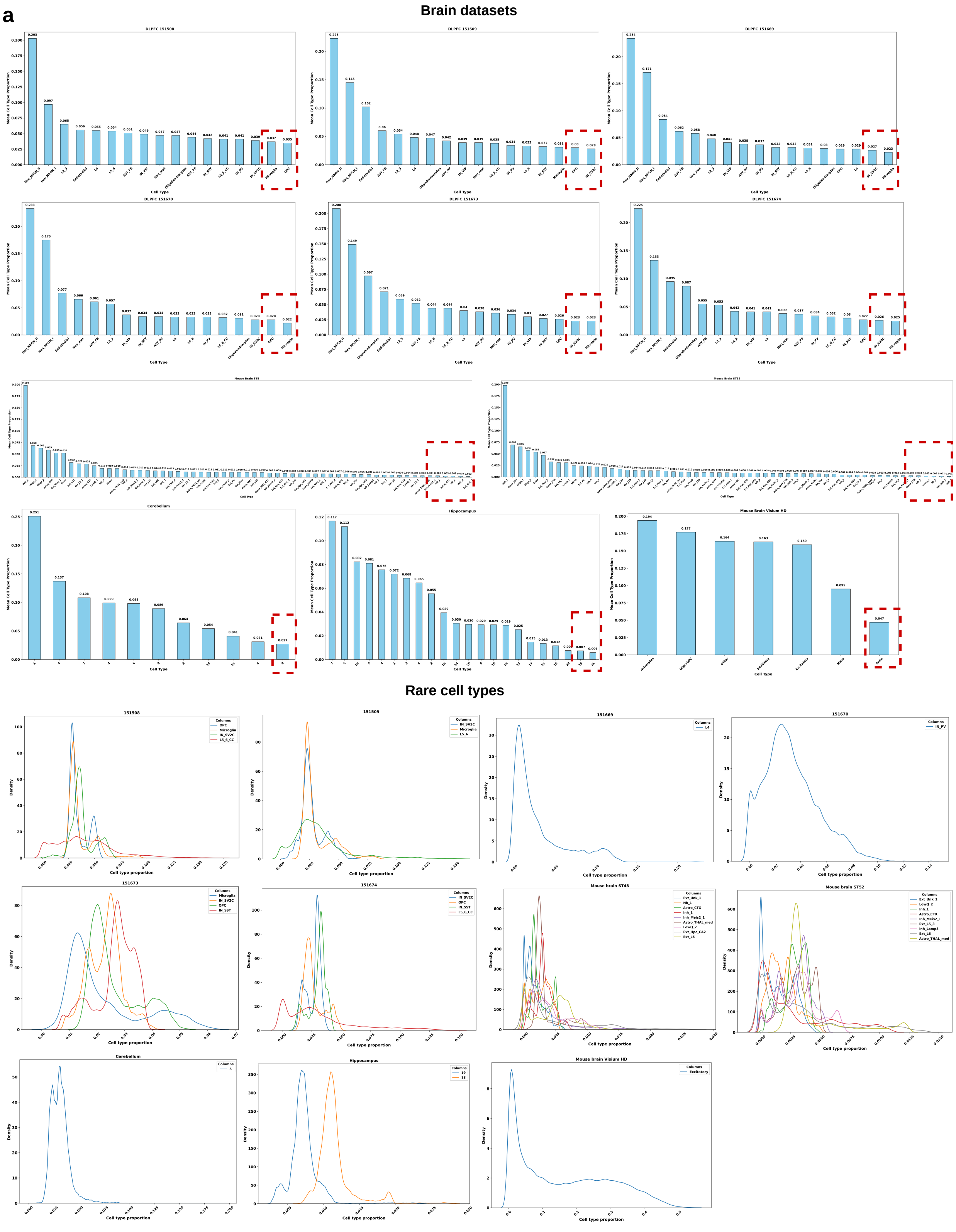


#### Regionally Rare cell types

**b**

#### Cancer datasets

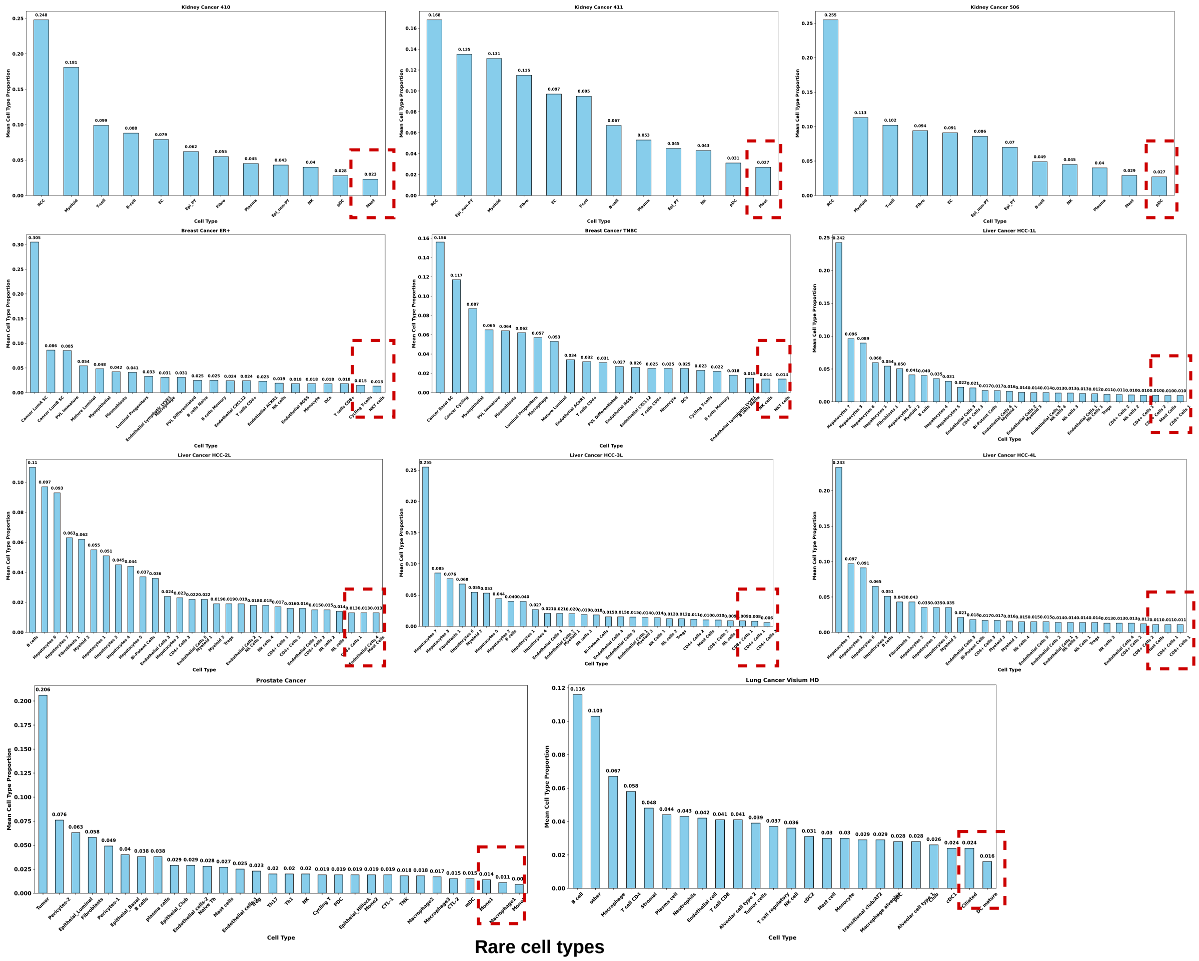


#### Rare cell types

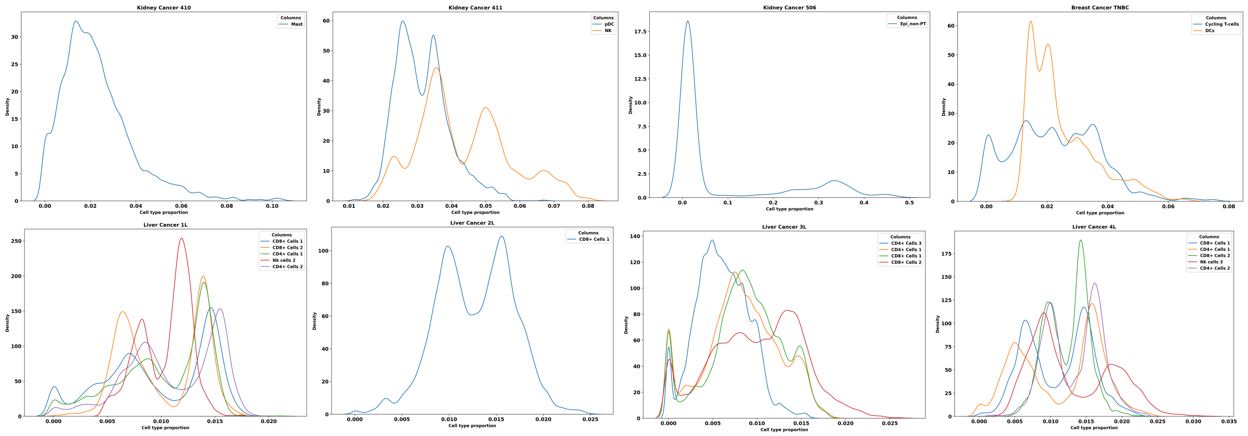


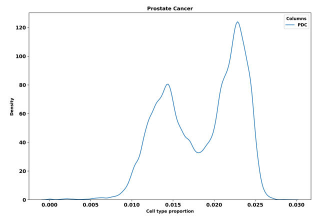


#### Regionally Rare cell types

#### Organs datasets

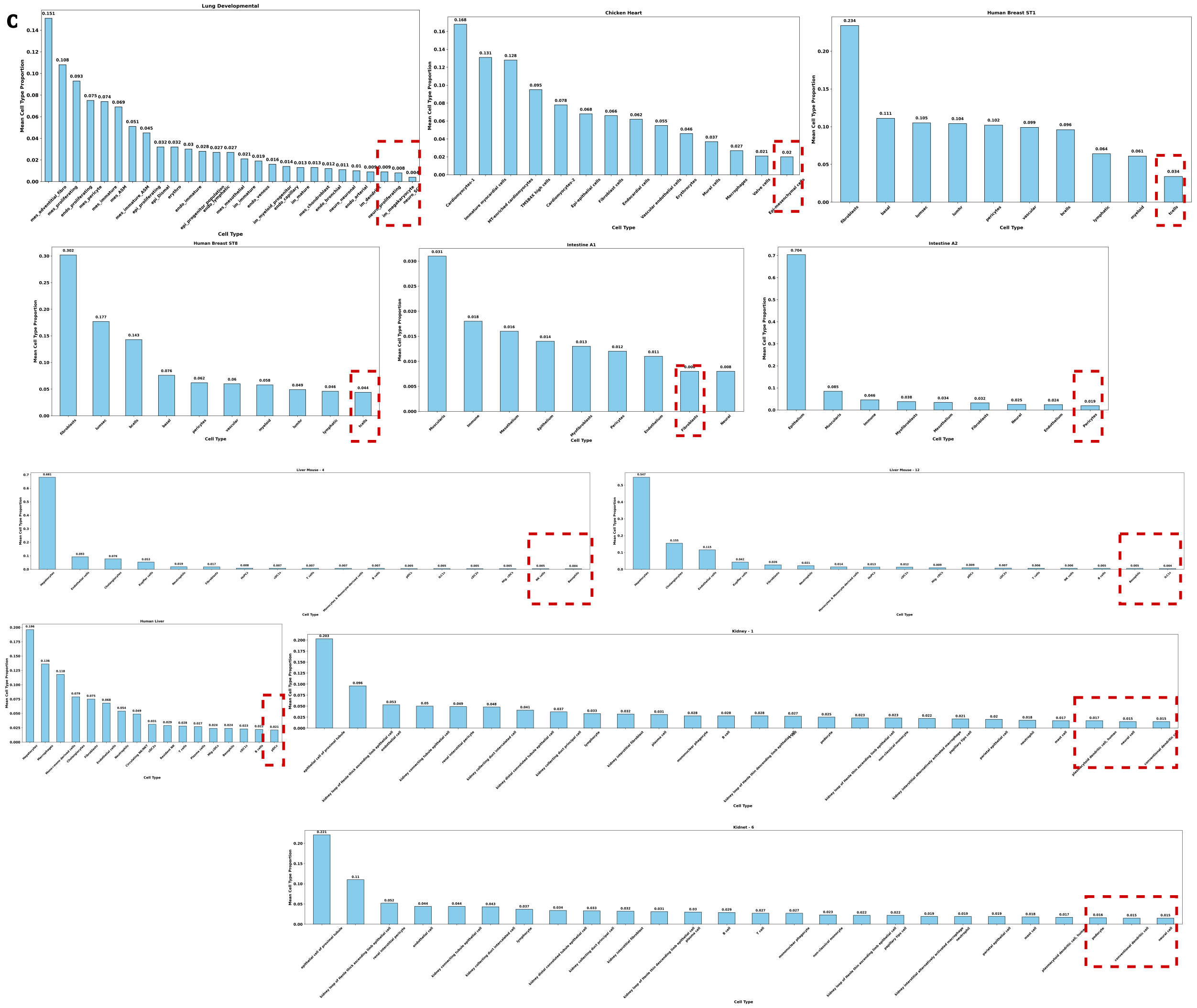


#### Rare cell types

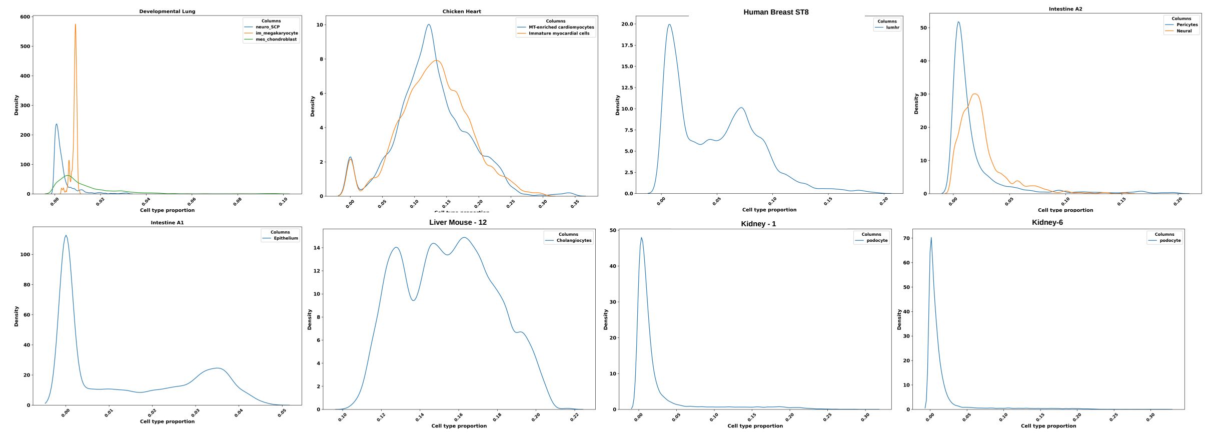


#### Regionally Rare cell types

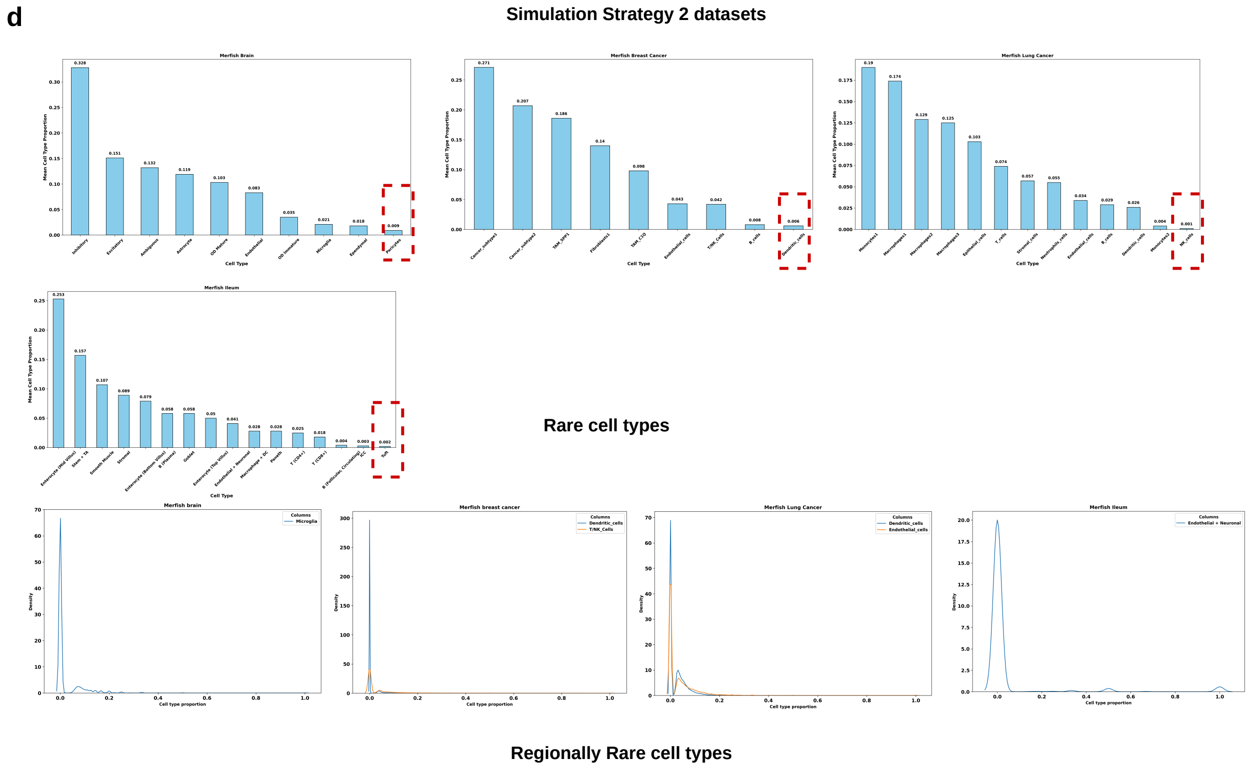


Supplementary Figure 12: Histogram of mean cell type proportions across all spots(top) and Density plot of distribution of cell type proportions across all spots for regionally rare cell types (bottom) for a) Brain datasets (i.e. 10X Visium DLPFC 151508, DLPFC 151509, DLPFC 151669, DLPFC 151670, DLPFC 151673, DLPFC 151674, 10X Visium Mouse Brain ST8059048(ST48), Mouse Brain ST8059052 (ST52), SlideseqV2 Cerebellum and SlideseqV2 Hippocampus, b) Cancer datasets (i.e. 10X Visium Kidney Cancer 410, Kidney Cancer 411, Kidney Cancer 506, 10X Visium Breast Cancer ER+ CID4290 (ER+), Breast Cancer TNBC CID44971(TNBC), 10X Visium Liver Cancer HCC-1L (1L), Liver Cancer HCC-2L (2L), Liver Cancer HCC-3L (3L), Liver Cancer HCC-4L (4L), SlideseqV2 Prostate Cancer and Visium HD Human Lung Cancer), c) Organs datasets (i.e. 10X Visium Developmental Lung, 10X Visium Chicken Heart, 10X Visium Intestine A1, Intestine A2, 10X Visium Human Breast $ST\_1$, Human Breast $ST\_8$, 10X Visium Mouse Liver 4, Mouse Liver - 12, Human Liver - 18, SlideseqV2 Kidney 200903_1 (1) and Kidney 200903_6) and d) Simulation strategy 2 datasets (MERFISH brain, MERFISH Breast Cancer, MERFISH Lung Cancer and MERFISH ileum).

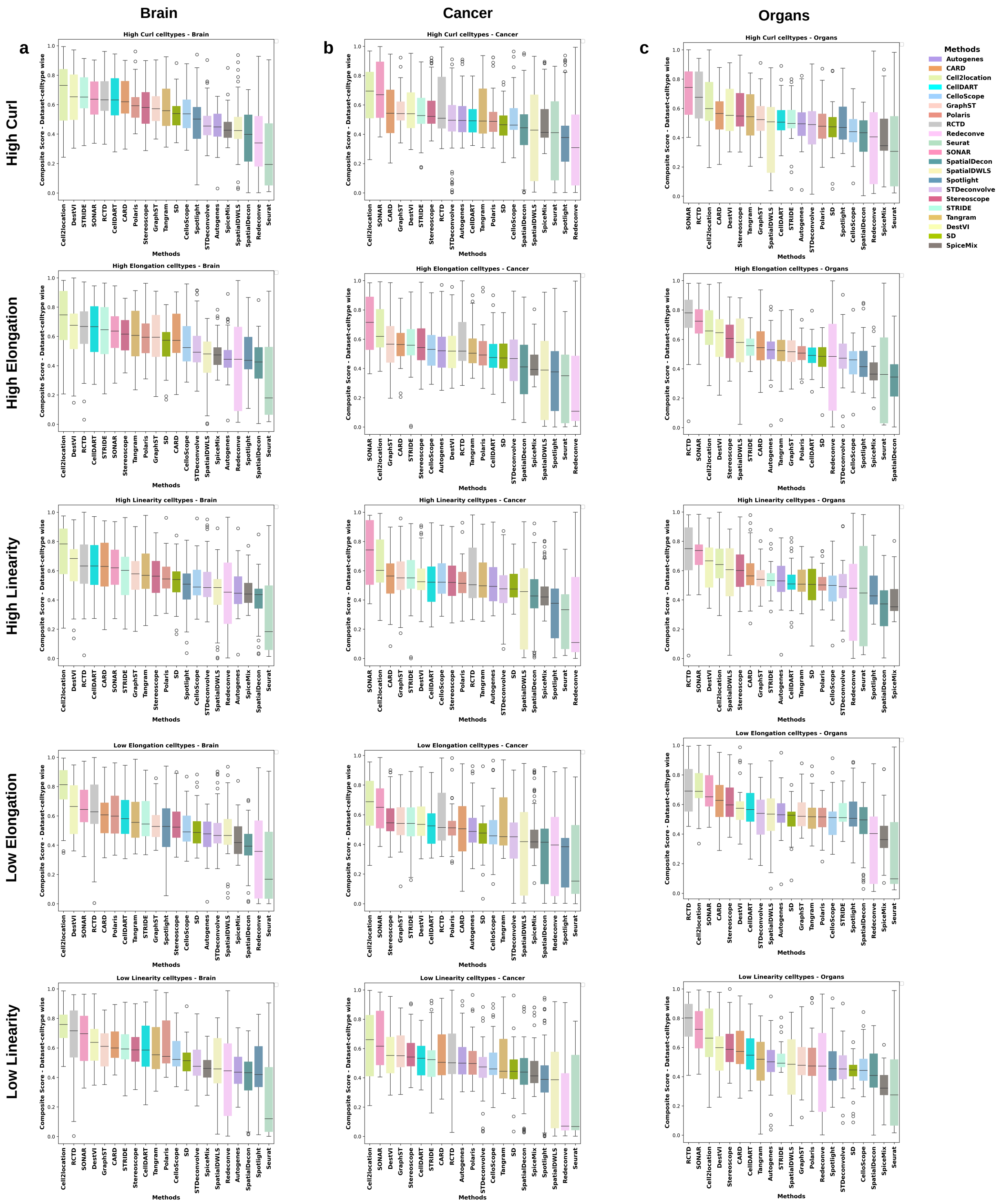
Brain
Cancer
Organs
a
b
c
High Curl
High Elongation
High Linearity
Low Elongation
Low Linearity
Methods
Autogenes
CARD
Cell2location
CellDART
CelloScope
GraphST
Polaris
RCTD
Redeconve
Seurat
SONAR
SpatialDecon
SpatialDWLS
Spotlight
STDeconvolve
Stereoscope
STRIDE
Tangram
DestVI
SD
SpiceMix

Supplementary Figure 13: Quantitative assessment of 21 deconvolution methods (Autogenes, CARD, Cell2location, CellDART, CelloScope, GraphST, Polaris, RCTD, Redeconve, Seurat, SONAR, SpatialDecon, SpatialDWLS, Spotlight, STDeconvolve, Stereoscope, STRIDE, Tangram, DestVI, SD and SpiceMix) comparing Bivariate Composite scores at a celltype level for cell types identified within shape characterization categories, including High Curl, High Elongation, High Linearity, Low Elongation, Low linearity for a) Brain datasets (i.e. 10X Visium DLPFC 151508, DLPFC 151509, DLPFC 151669, DLPFC 151670, DLPFC 151673, DLPFC 151674, 10X Visium Mouse Brain ST8059048(ST48), Mouse Brain ST8059052 (ST52), SlideseqV2 Cerebellum, SlideseqV2 Hippocampus and Merfish brain), b) Cancer datasets (i.e. 10X Visium Kidney Cancer 410, Kidney Cancer 411, Kidney Cancer 506, 10X Visium Breast Cancer ER+ CID4290 (ER+), Breast Cancer TNBC CID44971(TNBC), 10X Visium Liver Cancer HCC-1L (1L), Liver Cancer HCC-2L (2L), Liver Cancer HCC-3L (3L), Liver Cancer HCC-4L (4L), SlideseqV2 Prostate Cancer, Visium HD Human Lung Cancer, Merfish Breast Cancer and Merfish Lung Cancer), c) Organs datasets(i.e. 10X Visium Developmental Lung, 10X Visium Chicken Heart, 10X Visium Intestine A1, Intestine A2, 10X Visium Human Breast $ST\_1$, Human Breast $ST\_8$, 10X Visium Mouse Liver 4, Mouse Liver - 12, Human Liver - 18, SlideseqV2 Kidney 200903_1 (1), Kidney 200903_6 (6) and Merfish ileum)

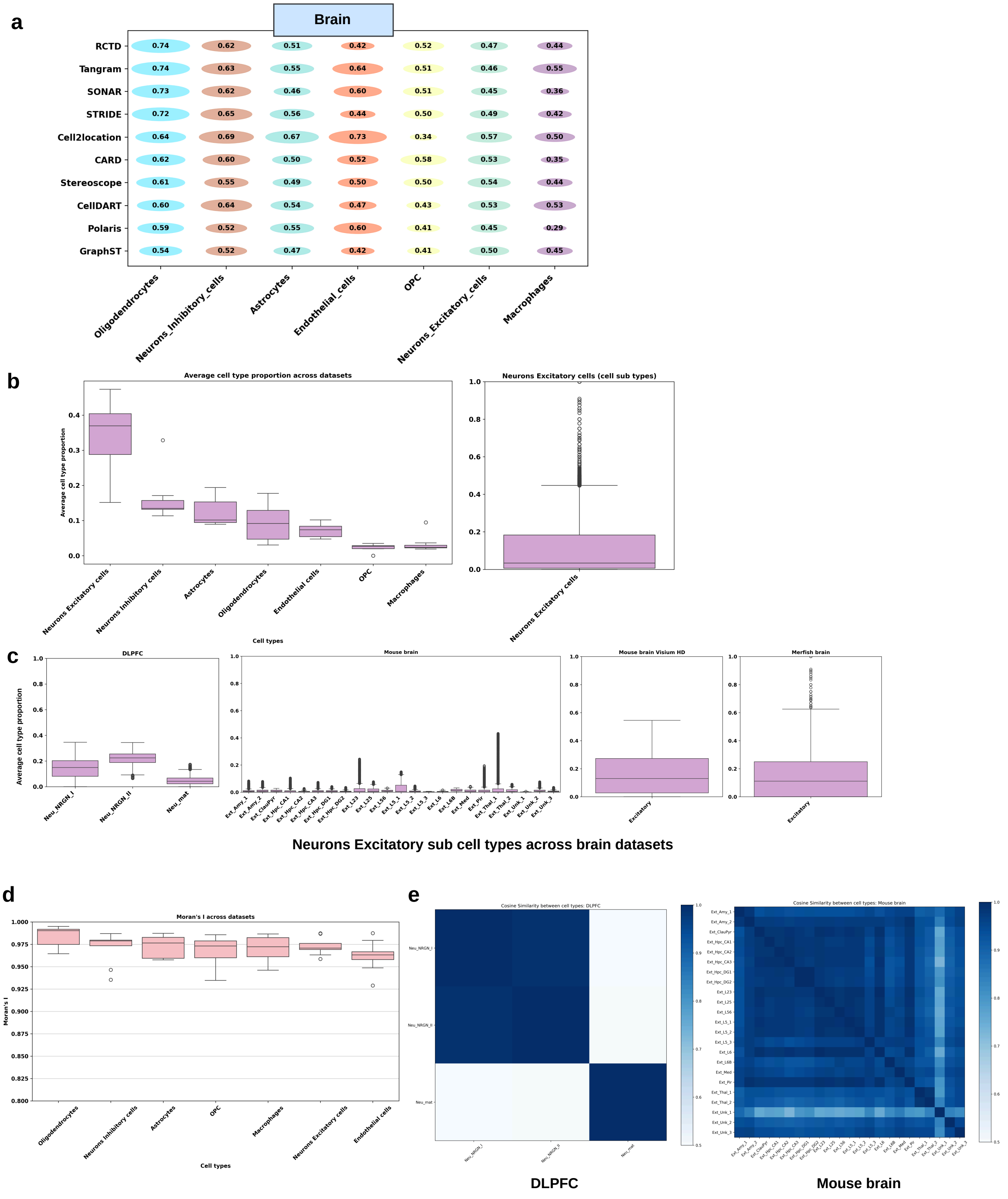
a
Brain
RCTD
Tangram
SONAR
STRIDE
Cell2location
CARD
Stereoscope
CellDART
Polaris
GraphST
Oligodendrocytes
Neurons_Inhibitory_cells
Astrocytes
Endothelial_cells
OPC
Neurons_Excitatory_cells
Macrophages
0.74 0.62 0.51 0.42 0.52 0.47 0.44
0.74 0.63 0.55 0.64 0.51 0.46 0.55
0.73 0.62 0.46 0.60 0.51 0.45 0.36
0.72 0.65 0.56 0.44 0.50 0.49 0.42
0.64 0.69 0.67 0.73 0.34 0.57 0.50
0.62 0.60 0.50 0.52 0.58 0.53 0.35
0.61 0.55 0.49 0.50 0.50 0.54 0.44
0.60 0.64 0.54 0.47 0.43 0.53 0.53
0.59 0.52 0.55 0.60 0.41 0.45 0.29
0.54 0.52 0.47 0.42 0.41 0.50 0.45
b
Average cell type proportion across datasets
Average cell type proportion
Cell types
Neurons Excitatory cells
Neurons Inhibitory cells
Astrocytes
Oligodendrocytes
Endothelial cells
OPC
Macrophages
Neurons Excitatory cells (cell sub types)
c
DLPFC
Mouse brain
Mouse brain Visium HD
Merfish brain
Average cell type proportion
Neu_NRGN_I
Neu_NRGN_II
Neu_mat
Excitatory
Neurons Excitatory sub cell types across brain datasets
d
Moran's I across datasets
Moran's I
Cell types
e
Cosine Similarity between cell types: DLPFC
Cosine Similarity between cell types: Mouse brain
DLPFC
Mouse brain

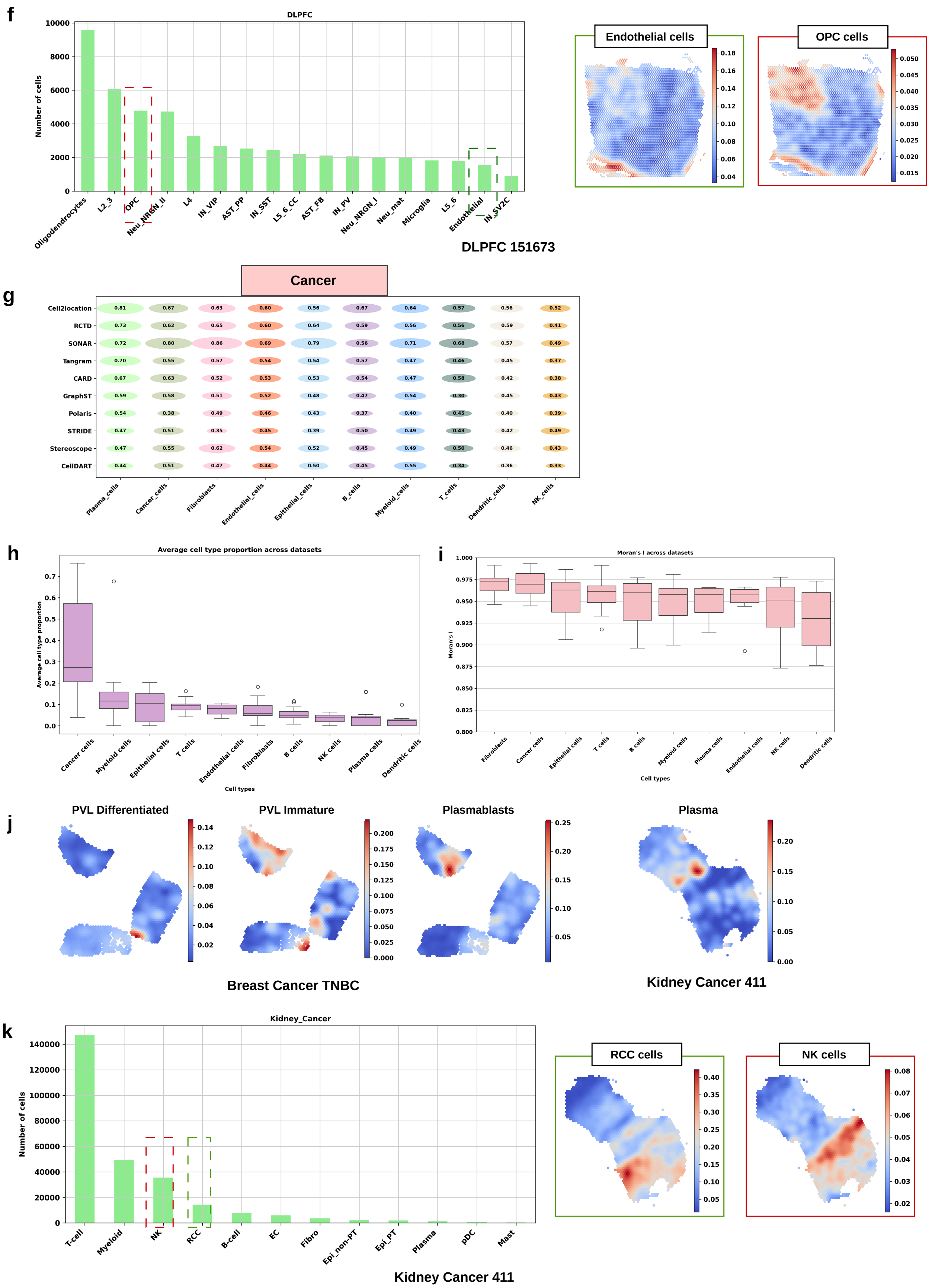
f
DLPFC
Number of cells
Oligodendrocytes
L2_3
OPC
Neu_NRGN_II
L4
IN_VIP
AST_PP
IN_SST
L5_6_CC
AST_FB
IN_PV
Neu_NRGN_I
Neu_mat
Microglia
L5_6
Endothelial
IN_SV2C
Endothelial cells
OPC cells
DLPFC 151673
g
Cancer
Cell2location 0.81 0.67 0.63 0.60 0.56 0.67 0.64 0.57 0.56 0.52
RCTD 0.73 0.62 0.65 0.60 0.64 0.59 0.56 0.56 0.59 0.41
SONAR 0.72 0.80 0.86 0.69 0.79 0.56 0.71 0.68 0.57 0.49
Tangram 0.70 0.55 0.57 0.54 0.54 0.57 0.47 0.46 0.45 0.37
CARD 0.67 0.63 0.52 0.53 0.53 0.54 0.47 0.58 0.42 0.38
GraphST 0.59 0.58 0.51 0.52 0.48 0.47 0.54 0.30 0.45 0.43
Polaris 0.54 0.38 0.49 0.46 0.43 0.37 0.40 0.45 0.40 0.39
STRIDE 0.47 0.51 0.35 0.45 0.39 0.50 0.49 0.43 0.42 0.49
Stereoscope 0.47 0.55 0.62 0.54 0.52 0.45 0.49 0.50 0.46 0.43
CellDART 0.44 0.51 0.47 0.44 0.50 0.45 0.55 0.34 0.36 0.33
Plasma_cells
Cancer_cells
Fibroblasts
Endothelial_cells
Epithelial_cells
B_cells
Myeloid_cells
T_cells
Dendritic_cells
NK_cells
h
Average cell type proportion across datasets
Average cell type proportion
Cancer cells
Myeloid cells
Epithelial cells
T cells
Endothelial cells
Fibroblasts
B cells
NK cells
Plasma cells
Dendritic cells
Cell types
i
Moran's I across datasets
Moran's I
Fibroblasts
Cancer cells
Epithelial cells
T cells
B cells
Myeloid cells
Plasma cells
Endothelial cells
NK cells
Dendritic cells
Cell types
j
PVL Differentiated
PVL Immature
Plasmablasts
Plasma
Breast Cancer TNBC
Kidney Cancer 411
k
Kidney_Cancer
Number of cells
T-cell
Myeloid
NK
RCC
B-cell
EC
Fibro
Epi_non-PT
Epi_PT
Plasma
pDC
Mast
RCC cells
NK cells
Kidney Cancer 411

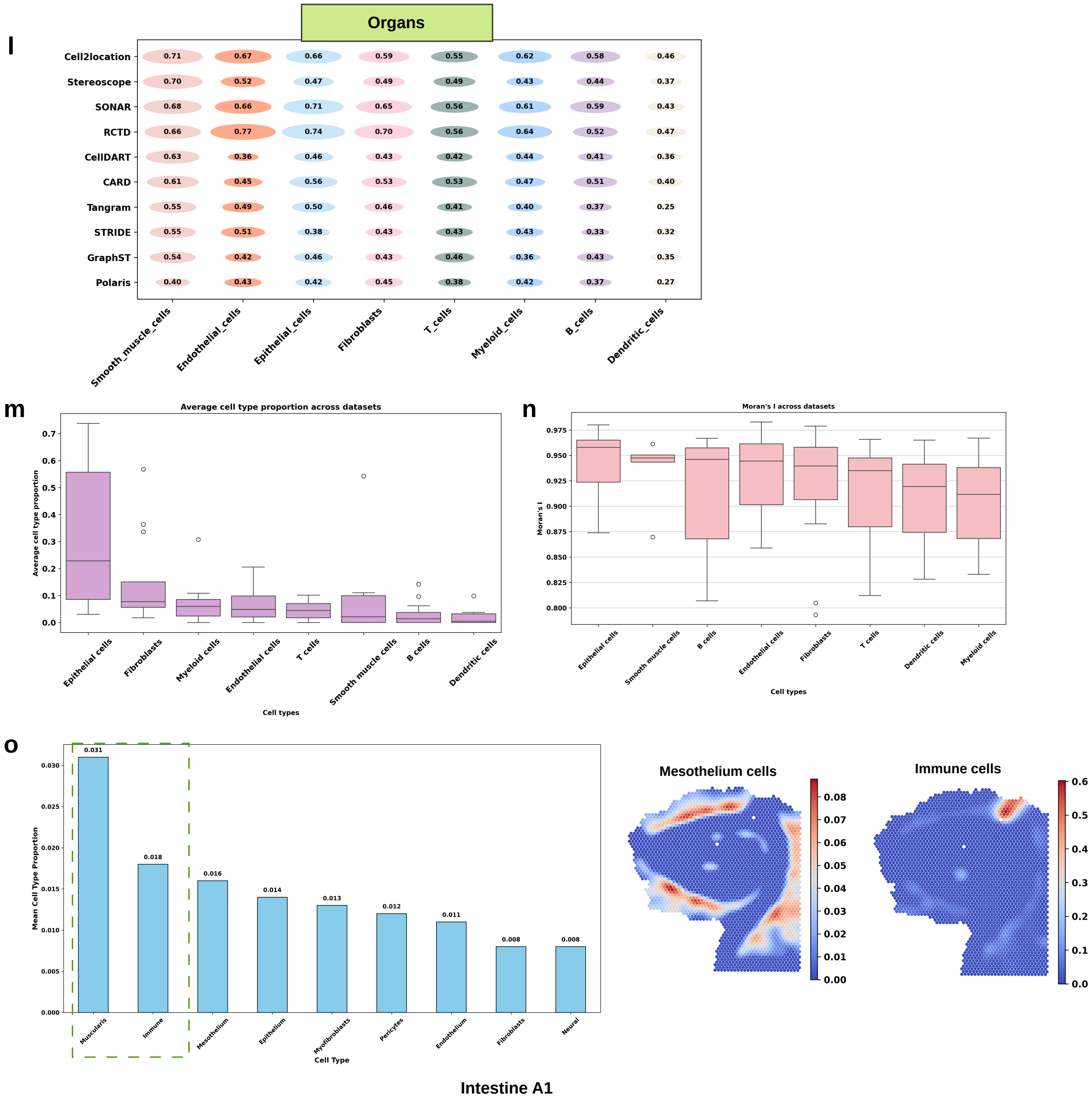
l
Organs
Cell2location 0.71 0.67 0.66 0.59 0.55 0.62 0.58 0.46
Stereoscope 0.70 0.52 0.47 0.49 0.49 0.43 0.44 0.37
SONAR 0.68 0.66 0.71 0.65 0.56 0.61 0.59 0.43
RCTD 0.66 0.77 0.74 0.70 0.56 0.64 0.52 0.47
CellDART 0.63 0.36 0.46 0.43 0.42 0.44 0.41 0.36
CARD 0.61 0.45 0.56 0.53 0.53 0.47 0.51 0.40
Tangram 0.55 0.49 0.50 0.46 0.41 0.40 0.37 0.25
STRIDE 0.55 0.51 0.38 0.43 0.43 0.43 0.33 0.32
GraphST 0.54 0.42 0.46 0.43 0.46 0.36 0.43 0.35
Polaris 0.40 0.43 0.42 0.45 0.38 0.42 0.37 0.27
Smooth_muscle_cells
Endothelial_cells
Epithelial_cells
Fibroblasts
T_cells
Myeloid_cells
B_cells
Dendritic_cells
m
Average cell type proportion across datasets
Average cell type proportion
Epithelial cells
Fibroblasts
Myeloid cells
Endothelial cells
T cells
Smooth muscle cells
B cells
Dendritic cells
Cell types
n
Moran's I across datasets
Moran's I
Epithelial cells
Smooth muscle cells
B cells
Endothelial cells
Fibroblasts
T cells
Dendritic cells
Myeloid cells
Cell types
o
Mean Cell Type Proportion
0.031
0.018
0.016
0.014
0.013
0.012
0.011
0.008
0.008
Muscularis
Immune
Mesothelium
Epithelium
Myofibroblasts
Pericytes
Endothelium
Fibroblasts
Neural
Cell Type
Mesothelium cells
Immune cells
Intestine A1

Supplementary Figure 14: a) Comparison of composite scores across major cell types-Astrocytes, Endothelial cells, Macrophages, Excitatory neurons, Inhibitory neurons, Oligodendrocyte progenitor cells (OPCs), and Oligodendrocytes, for top-performing methods on Brain datasets. b) Boxplot comparison showing (left) the average cell-type proportion across all brain datasets and (right) the average cell-type proportion for excitatory neuron subtypes across brain datasets. c) Boxplot comparison showing the average cell-type proportion of individual excitatory neuron subtypes for each brain dataset. d) Boxplot comparison showing Moran's I statistics across all brain datasets. e) Heatmap of cosine similarity of gene expression profiles between different cell subtypes of excitatory neurons. f) Example analysis on the DLPFC 151673 dataset, including a comparison of cell counts from the corresponding single-cell reference and spatial pattern plots showing the predicted proportions of Endothelial and OPC cell types. g) Comparison of composite scores across major cell types—B cells, Cancer cells, Dendritic cells, Endothelial cells, Epithelial cells, Fibroblasts, Myeloid cells, NK cells, Plasma cells, and T cells, for top-performing methods on Cancer datasets. h) Boxplot comparison showing the average cell-type proportion, and i) Moran's I statistics across all cancer datasets. j) Spatial pattern plots of plasma cells for cancer datasets, Breast Atlas TNBC and Kidney Cancer 411. k) Example analysis on the Kidney Cancer 411 dataset, including a comparison of cell counts from the single-cell reference and spatial pattern plots highlighting RCC and NK cell type proportions. l) Comparison of composite scores across major cell types—B cells, Dendritic cells, Endothelial cells, Epithelial cells, Fibroblasts, Myeloid cells, Smooth muscle cells, and T cells, for Organ datasets. m) Boxplot comparison showing the average cell-type proportion, and n) Moran's I statistics across all organs datasets. o)Example analysis on the Intestine A1 dataset, including a comparison of mean cell-type proportions across all spots and spatial pattern plots highlighting Mesothelium and Immune cell types.

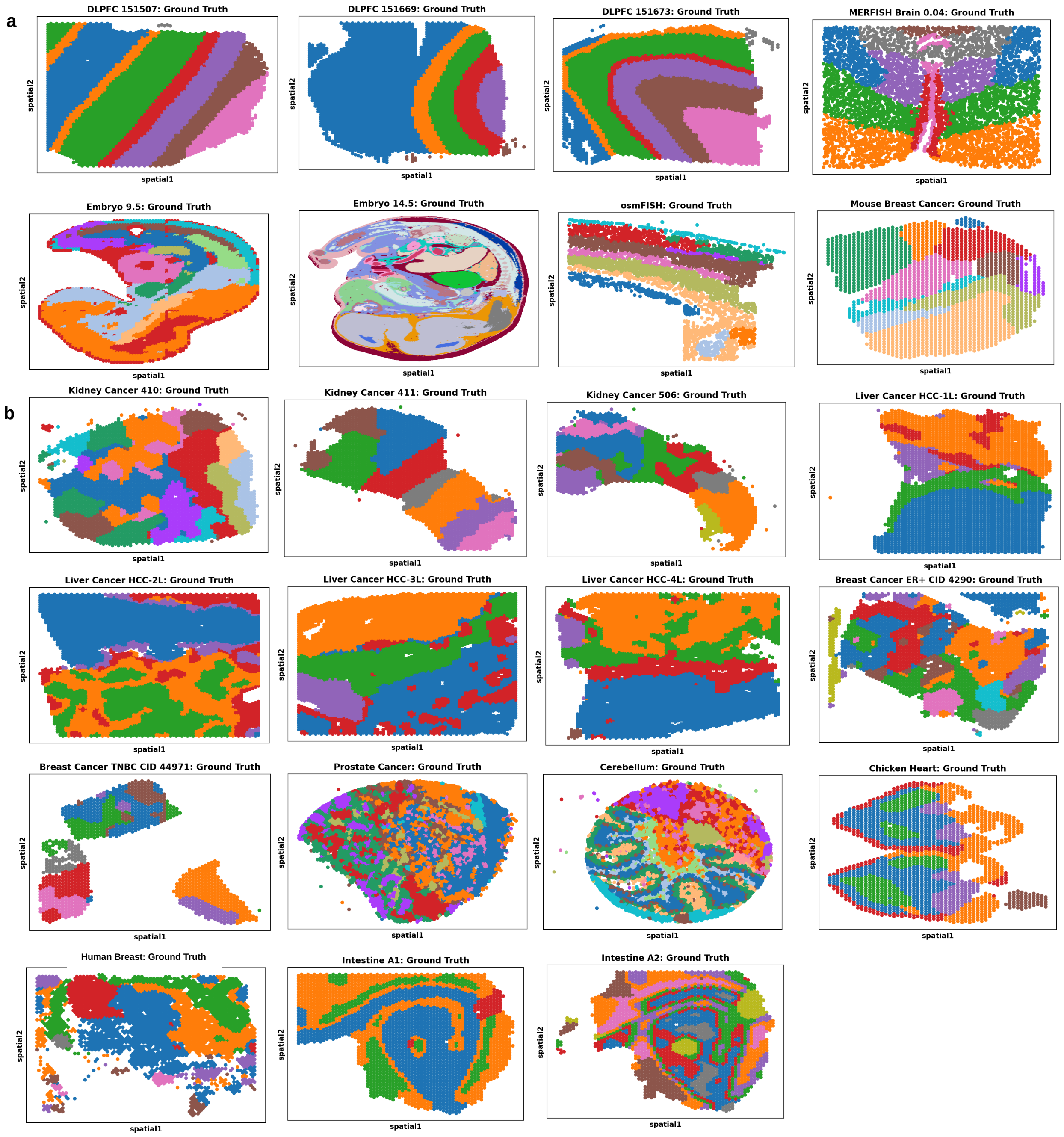


Supplementary Figure 15: a) Variation in the different spatial patterns present in a) real datasets: DLPFC (151507, 151669, 151673), Merfish Mouse brain (0.04), Embryo(9.5, 14.5), osmFISH Mouse brain, Mouse Breast Cancer, and simulated datasets: Kidney Cancer(410, 411, 506), Liver Cancer(HCC-1L, HCC-2L, HCC-3L, HCC-4L), Breast Cancer (ER+, TNBC), Prostate Cancer, Cerebellum, Chicken Heart, Human Breast (ST8), Intestine (A1, A2).

| Sno | Methods | DLPFC | Mouse Embryo-9.5 | Mouse Embryo-14.5 | Mouse breast cancer | MERFISH brain | Mouse cortex-osmFISH | Kidney Cancer | Breast Cancer | Liver Cancer | Breast Atlas ST8 | Intestine | Chicken Heart | Prostate Cancer | Cere-bellum |
|---|---|---|---|---|---|---|---|---|---|---|---|---|---|---|---|
| 1 | Banksy | | | | | | | | | | | | | | |
| 2 | BASS | | | | | | | | | | | | | | |
| 3 | BayesSpace | | | | | | | | | | | | | | |
| 4 | BayesCafe | | | | | | | | | | | | | | |
| 5 | CCST | | | | | | | | | | | | | | |
| 6 | DeepST | | | | | | | | | | | | | | |
| 7 | DRSC | | | | | | | | | | | | | | |
| 8 | Giotto | | | | | | | | | | | | | | |
| 9 | GraphST | | | | | | | | | | | | | | |
| 10 | IRIS | | | | | | | | | | | | | | |
| 11 | iSCMEB | | | | | | | | | | | | | | |
| 12 | PRECAST | | | | | | | | | | | | | | |
| 13 | PROST | | | | | | | | | | | | | | |
| 14 | SCANIT | | | | | | | | | | | | | | |
| 15 | SpaceFlow | | | | | | | | | | | | | | |
| 16 | SpaSRL | | | | | | | | | | | | | | |
| 17 | SpatialPCA | | | | | | | | | | | | | | |
| 18 | STAGATE | | | | | | | | | | | | | | |

Supplementary Figure 16: Summary table of spatial domain detection methods (rows) included in our benchmarking study and datasets (columns). Red boxes represent cases where a method failed to run on a given dataset.

a

| Sno | Method | Language | Input | # Top 3 | # Top 3 Visium | # Top 3 Image based datasets | # Top 3 Large datasets | # datasets failed to run | Time |
|---|---|---|---|---|---|---|---|---|---|
| 1 | PROST | Python | # clusters | 15 | 12 | 1 | 2 | 3 | M |
| 2 | GraphST | Python | # clusters | 15 | 14 | 1 | 0 | 3 | L |
| 3 | CCST | Python | # clusters | 12 | 12 | 0 | 0 | 2 | H |
| 4 | SCANIT | Python | resolution | 11 | 9 | 0 | 2 | 1 | L |
| 5 | IRIS | Python | # clusters | 10 | 10 | 0 | 0 | 6 | L |
| 6 | SpaceFlow | Python | resolution | 9 | 2 | 4 | 3 | 0 | L |
| 7 | BASS | R | # clusters | 9 | 4 | 5 | 0 | 2 | M |
| 8 | Banksy | Python/R | resolution | 8 | 1 | 7 | 0 | 2 | L |
| 9 | SpatialPCA | R | # clusters | 5 | 4 | 1 | 0 | 7 | H |
| 10 | DeepST | Python | # clusters | 5 | 5 | 0 | 0 | 3 | L |
| 11 | ISC.MEB | R | # clusters | 3 | 2 | 1 | 0 | 1 | L |
| 12 | BayesSpace | R | # clusters | 3 | 2 | 0 | 1 | 0 | M |
| 13 | SpaSRL | Python | resolution | 1 | 0 | 1 | 0 | 2 | M |
| 14 | PRECAST | R | # clusters | 1 | 0 | 0 | 1 | 1 | L |
| 15 | BayesCafe | R | # clusters | 1 | 1 | 0 | 0 | 4 | H |
| 16 | STAGATE | Python | # clusters | 0 | 0 | 0 | 0 | 1 | L |
| 17 | Giotto | R | resolution | 0 | 0 | 0 | 0 | 2 | L |
| 18 | DRSC | R | # clusters | 0 | 0 | 0 | 0 | 2 | L |

b

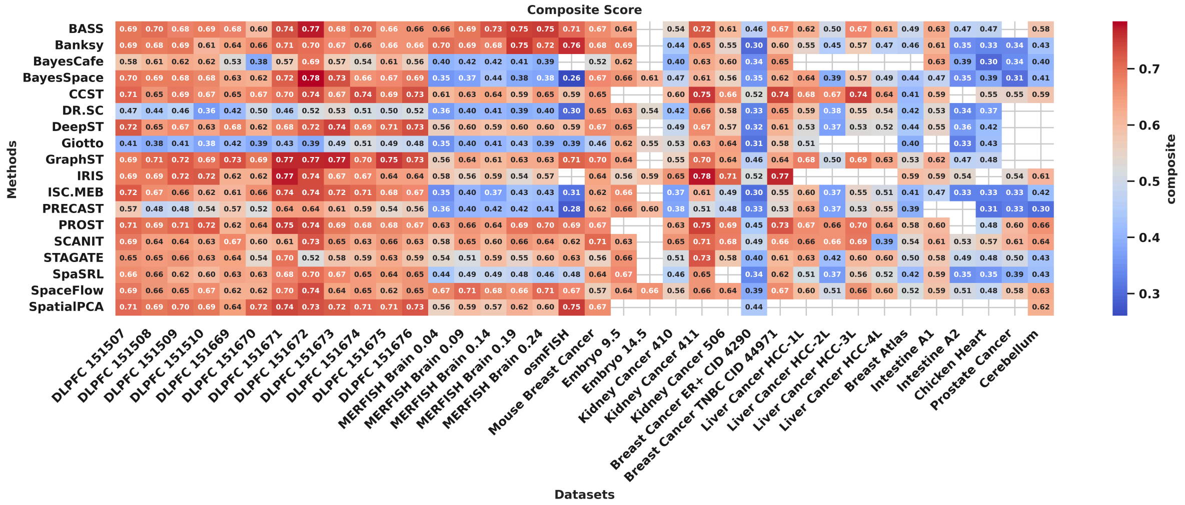


c

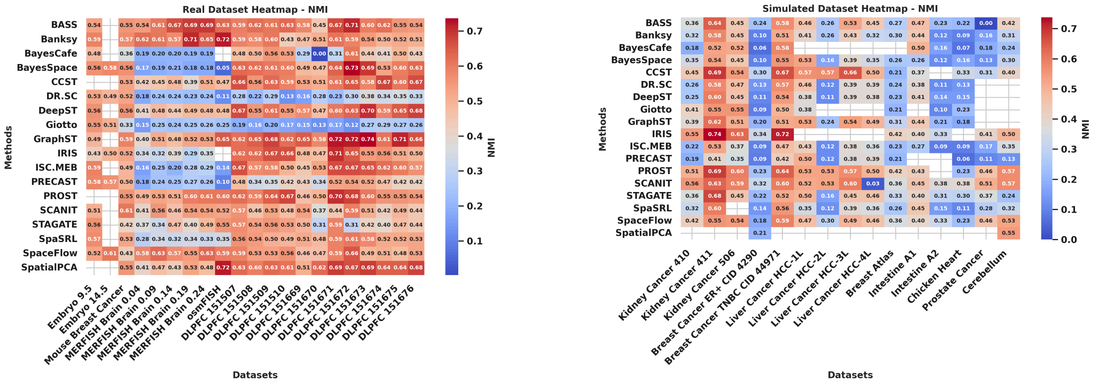

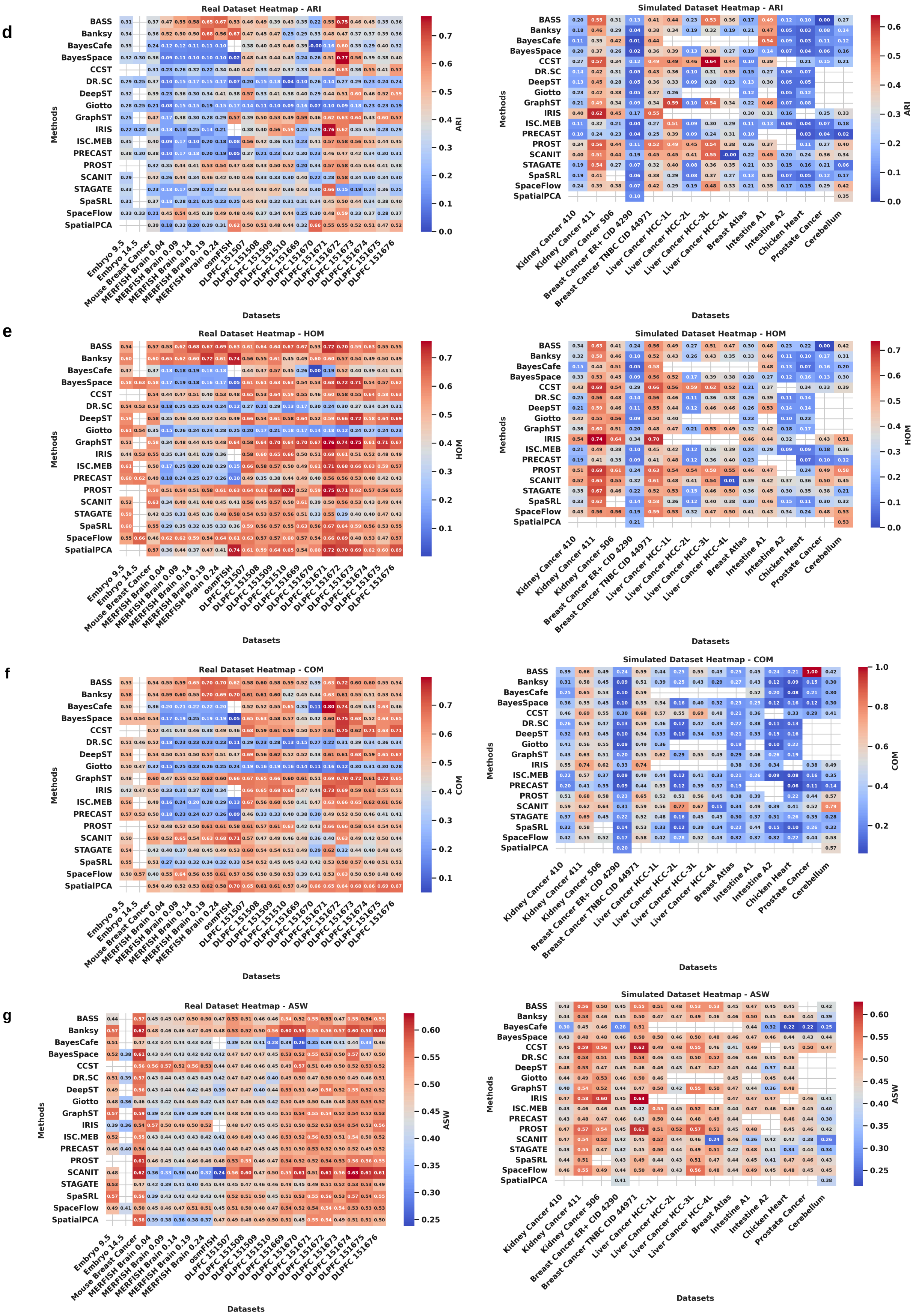
d
Real Dataset Heatmap - ARI
Simulated Dataset Heatmap - ARI
e
Real Dataset Heatmap - HOM
Simulated Dataset Heatmap - HOM
f
Real Dataset Heatmap - COM
Simulated Dataset Heatmap - COM
g
Real Dataset Heatmap - ASW
Simulated Dataset Heatmap - ASW
Methods
Datasets
BASS
Banksy
BayesCafe
BayesSpace
CCST
DR.SC
DeepST
Giotto
GraphST
IRIS
ISC.MEB
PRECAST
PROST
SCANIT
STAGATE
SpaSRL
SpaceFlow
SpatialPCA
Embryo 9.5
Embryo 14.5
Mouse Breast Cancer
MERFISH Brain 0.04
MERFISH Brain 0.09
MERFISH Brain 0.14
MERFISH Brain 0.19
MERFISH Brain 0.24
osmFISH
DLPFC 151507
DLPFC 151508
DLPFC 151509
DLPFC 151510
DLPFC 151669
DLPFC 151670
DLPFC 151671
DLPFC 151672
DLPFC 151673
DLPFC 151674
DLPFC 151675
DLPFC 151676
Kidney Cancer 410
Kidney Cancer 411
Kidney Cancer 506
Breast Cancer ER+ CID 4290
Breast Cancer TNBC CID 44971
Liver Cancer HCC-1L
Liver Cancer HCC-2L
Liver Cancer HCC-3L
Liver Cancer HCC-4L
Breast Atlas
Intestine A1
Intestine A2
Chicken Heart
Prostate Cancer
Cerebellum
ARI
HOM
COM
ASW

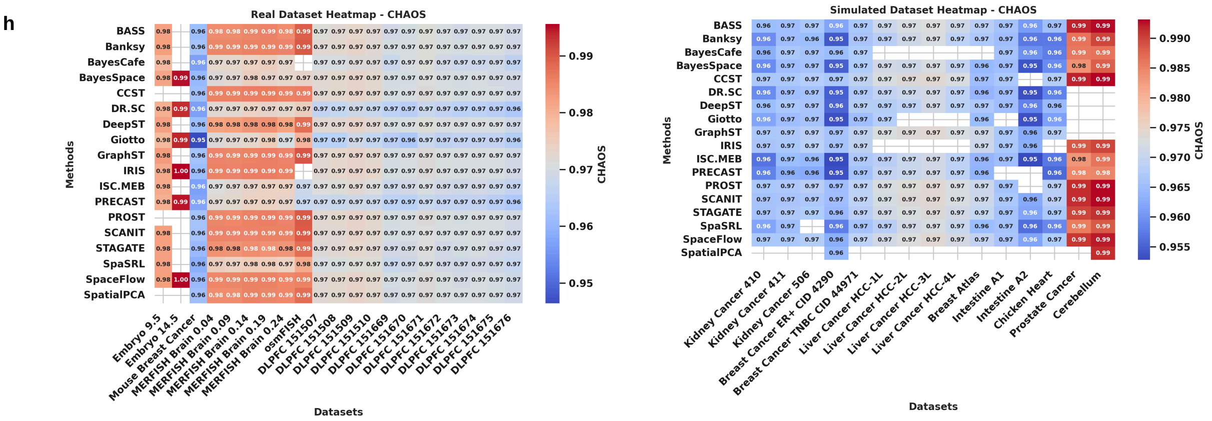


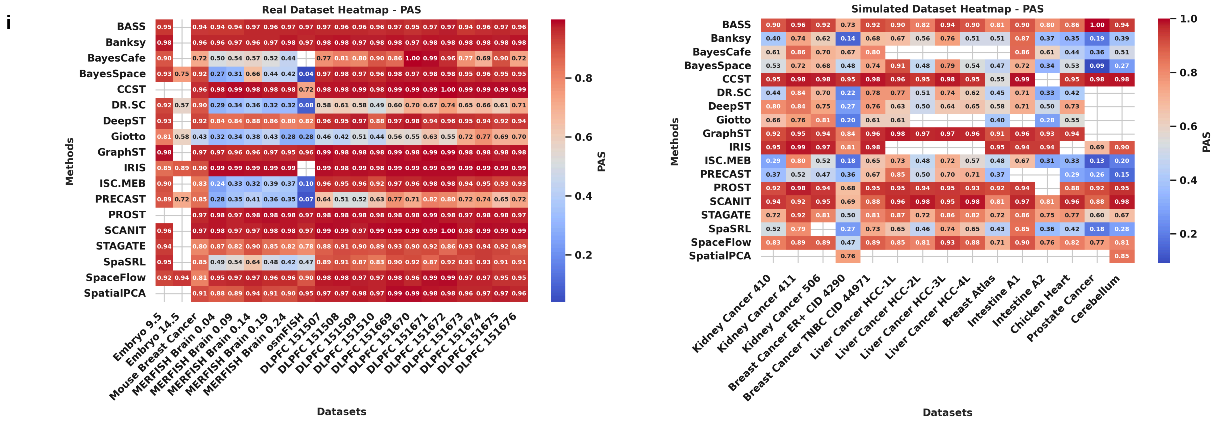


Supplementary Figure 17: a) Summary table of Usability performance metrics for 18 domain detection methods (BASS, Banksy, BayesCafe, BayesSpace, CCST, DR.SC, DeepST, Giotto, GraphST, iSC.MEB, IRIS, PRECAST, PROST, SCANIT, STAGATE, SpaSRL, SpaceFlow and SpatialPCA). A light green box indicates top-performing method. Quantitative assessment of the 18 domain detection methods across 35 datasets comparing b) overall composite scores, c) NMI, d) ARI, e) HOM, f) COM, g) ASW, h) CHAOS and i) PAS for real (left) and simulated (right) datasets.

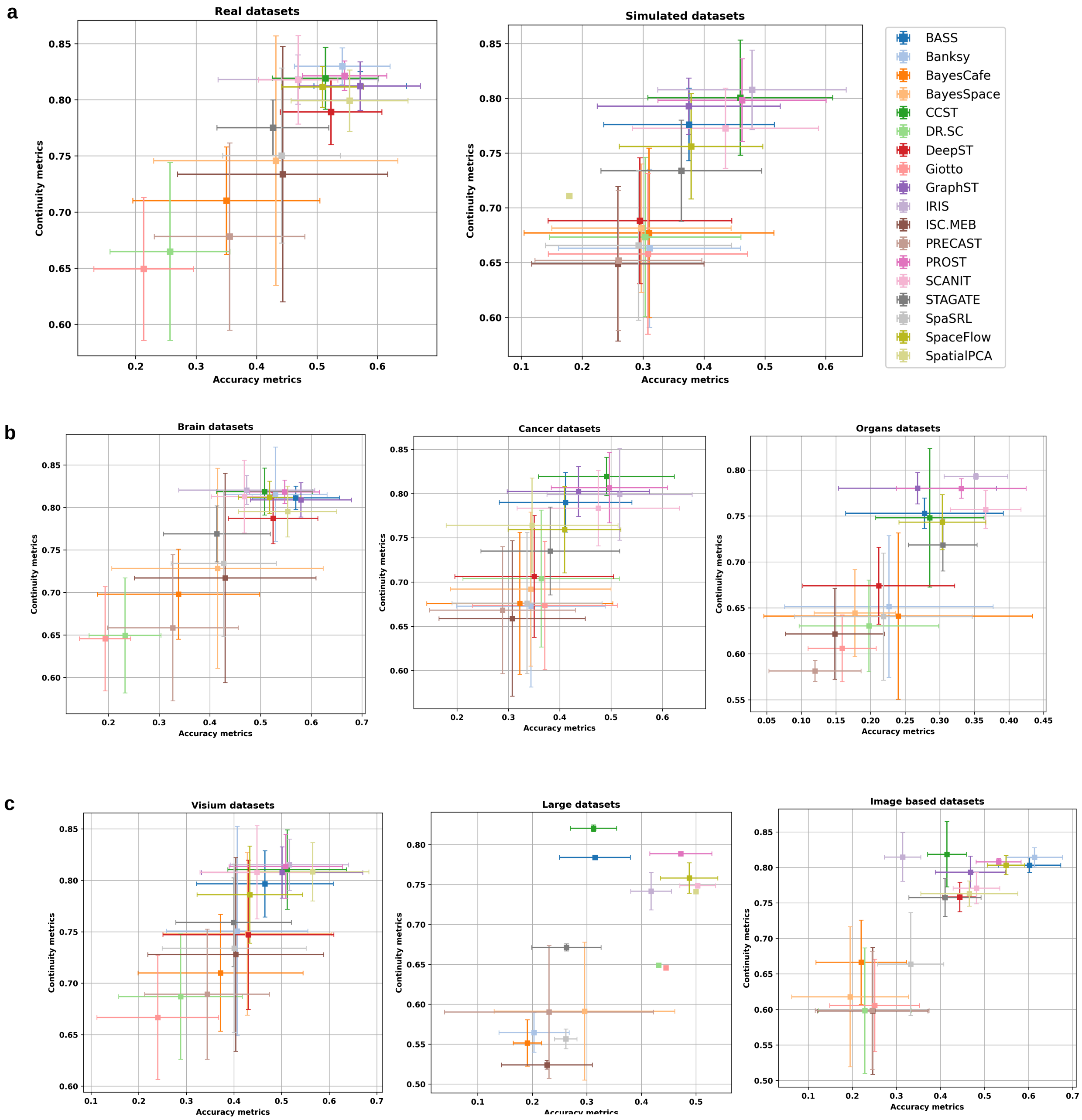


Supplementary Figure 18: Quantitative assessment of 18 domain detection methods (BASS, Banksy, BayesCafe, BayesSpace, CCST, DR.SC, DeepST, Giotto, GraphST, iSC.MEB, IRIS, PRECAST, PROST, SCANIT, STAGATE, SpaSRL, SpaceFlow and SpatialPCA) comparing composite scores. Comparison of Composite continuity score and Composite accuracy score for a) 21 real datasets and 15 simulated datasets across, b) Brain, Cancer and Organs datasets, c) Visium, Large datasets and Imaging-based datasets. Data are presented as mean values ± standard deviation.

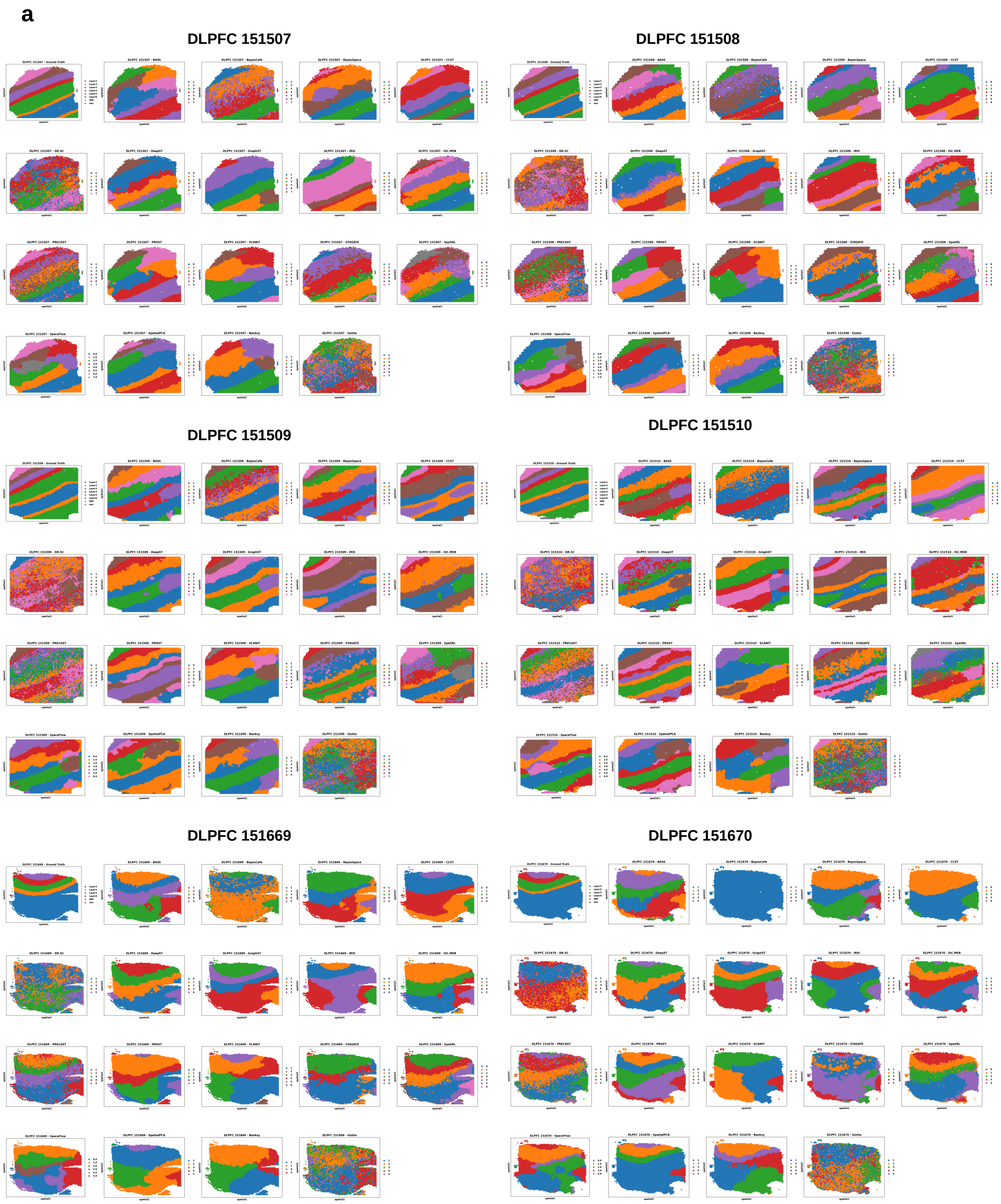
a
DLPFC 151507
DLPFC 151508
DLPFC 151509
DLPFC 151510
DLPFC 151669
DLPFC 151670

#### DLPFC 151671

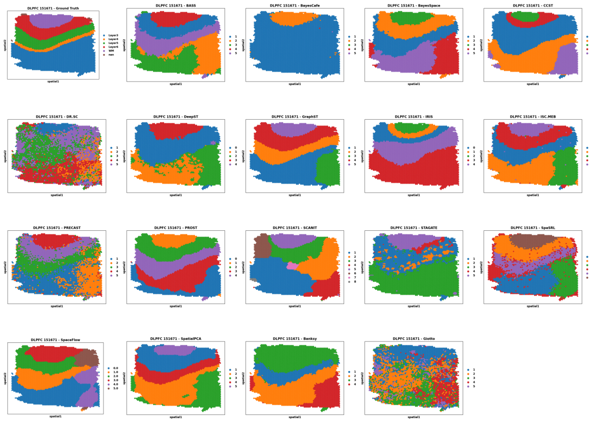

#### DLPFC 151672

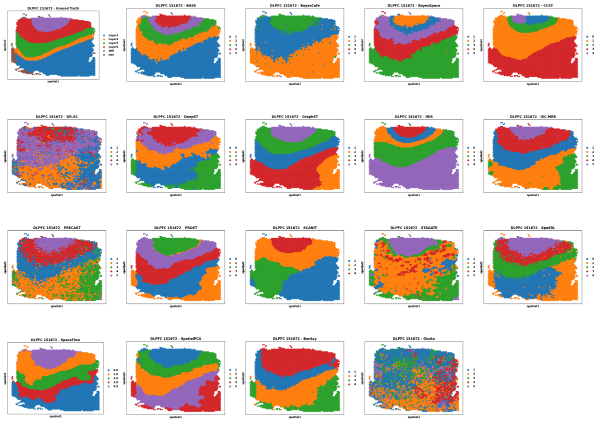

#### DLPFC 151673

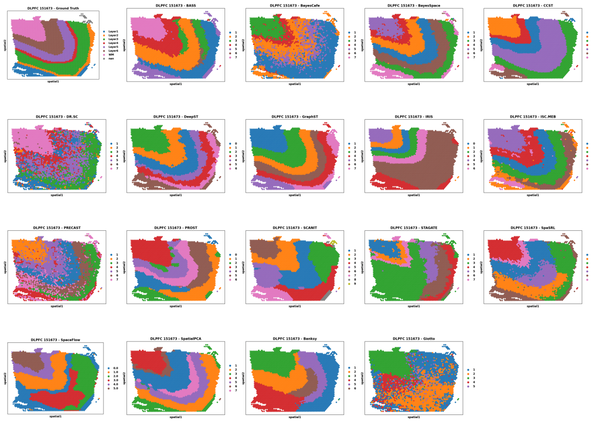

#### DLPFC 151674

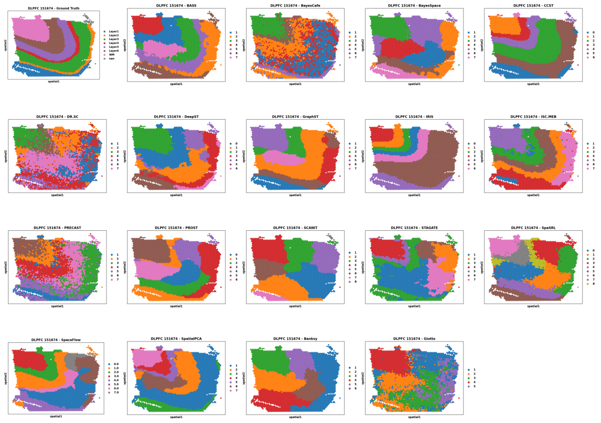

#### DLPFC 151675

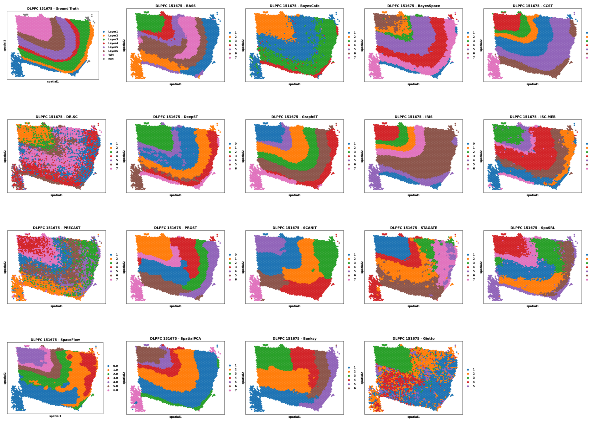

#### DLPFC 151676

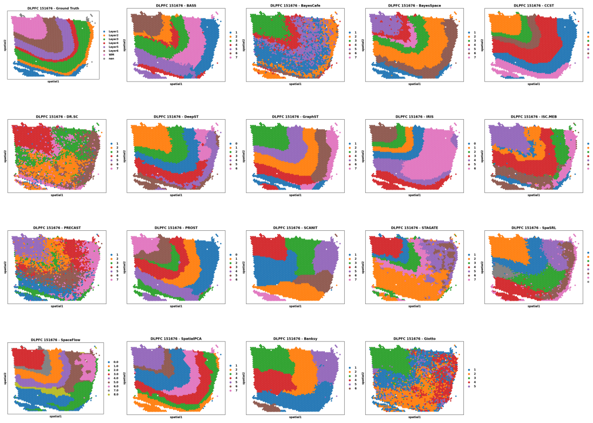

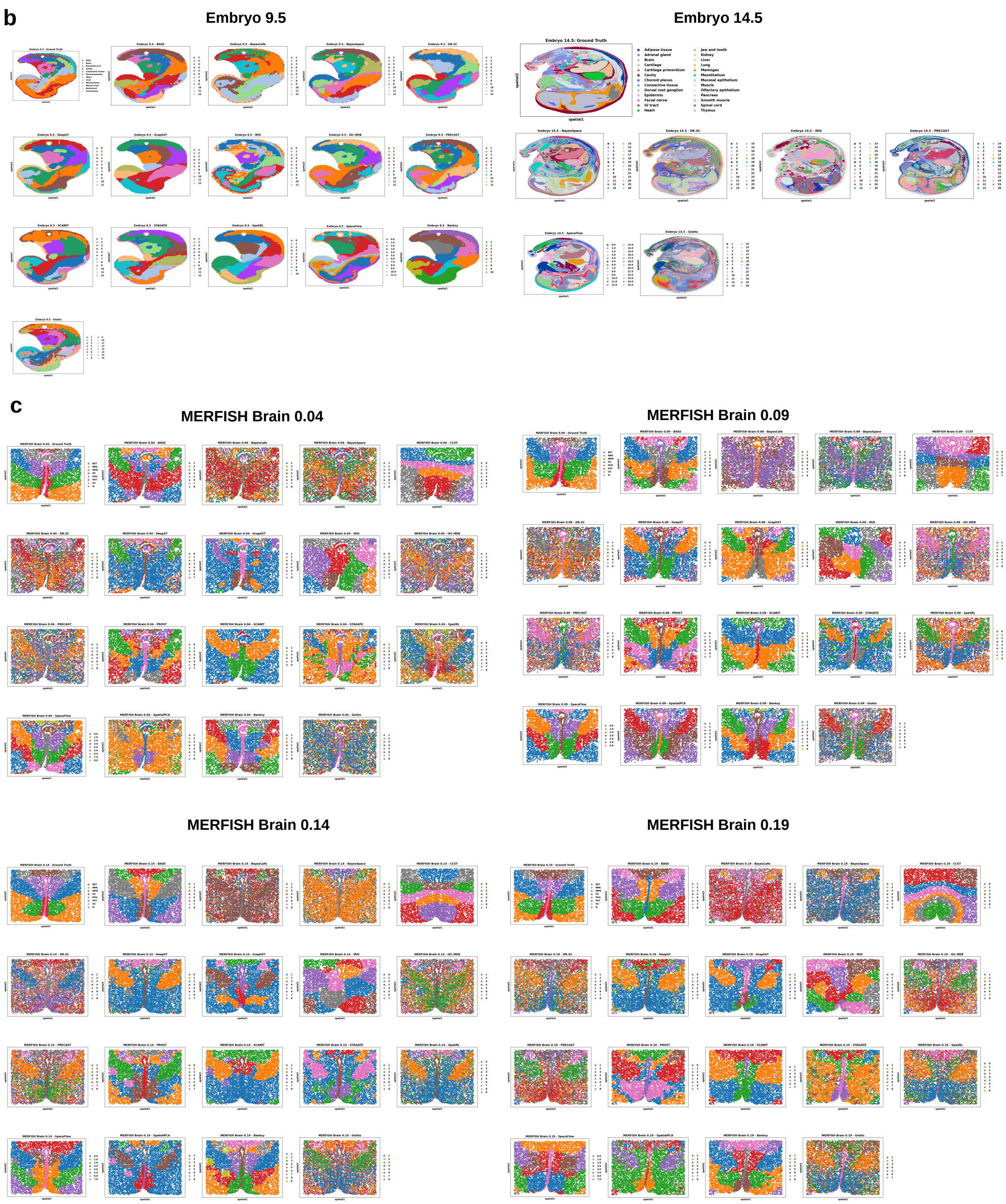
b
Embryo 9.5
Embryo 14.5
c
MERFISH Brain 0.04
MERFISH Brain 0.09
MERFISH Brain 0.14
MERFISH Brain 0.19

#### MERFISH Brain 0.24

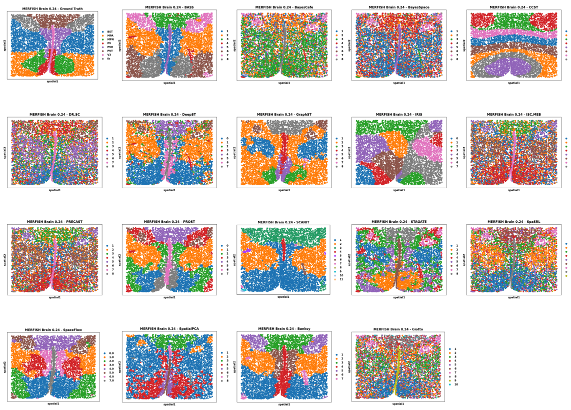

**d** 

#### Mouse Cortex

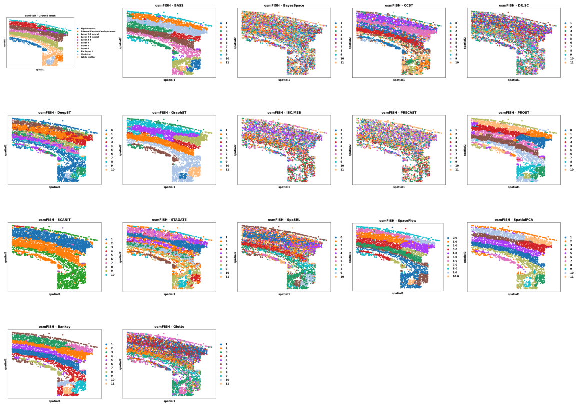

**e** 

#### Mouse Breast Cancer

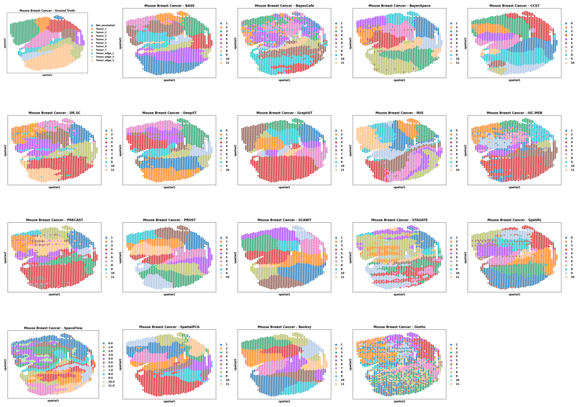

**f** 

#### Kidney Cancer 410

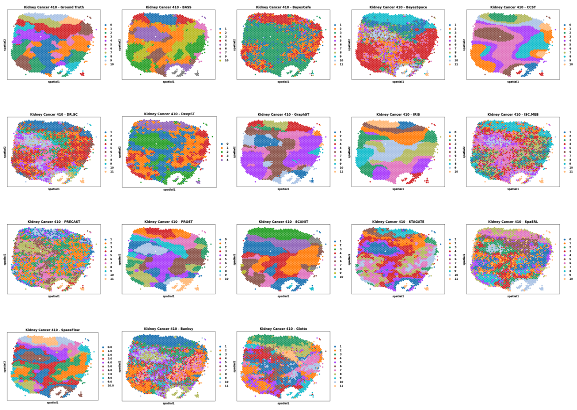

#### Kidney Cancer 411

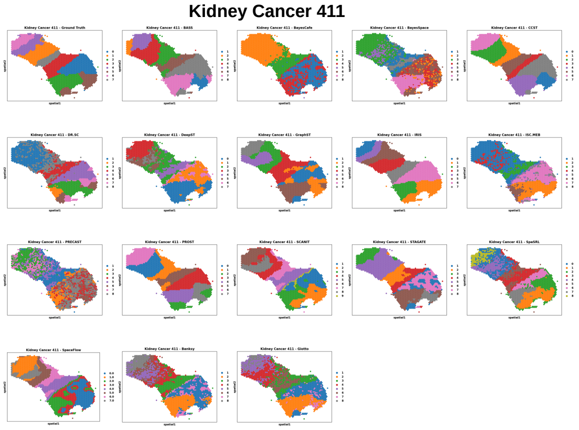

#### Kidney Cancer 506

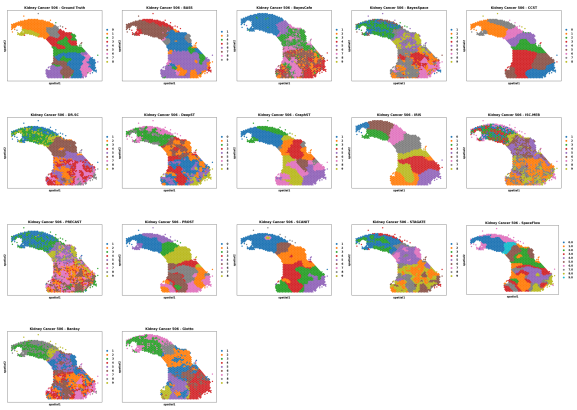

g

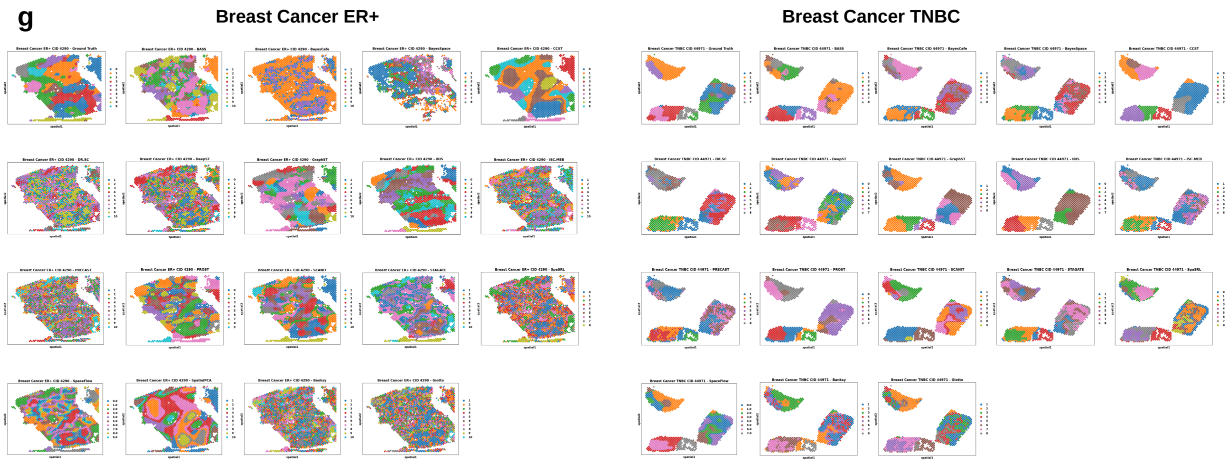
Breast Cancer ER+
Breast Cancer TNBC

h

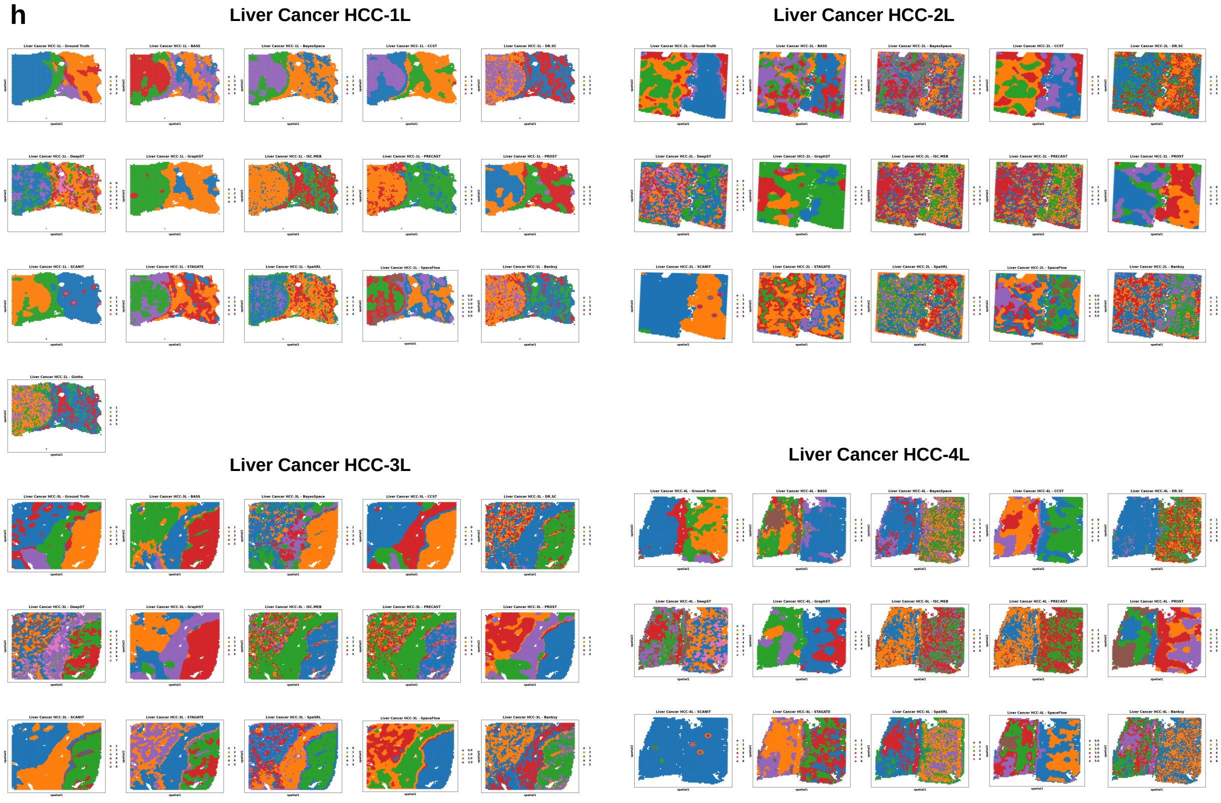
Liver Cancer HCC-1L
Liver Cancer HCC-2L
Liver Cancer HCC-3L
Liver Cancer HCC-4L

**i** **Breast Atlas**

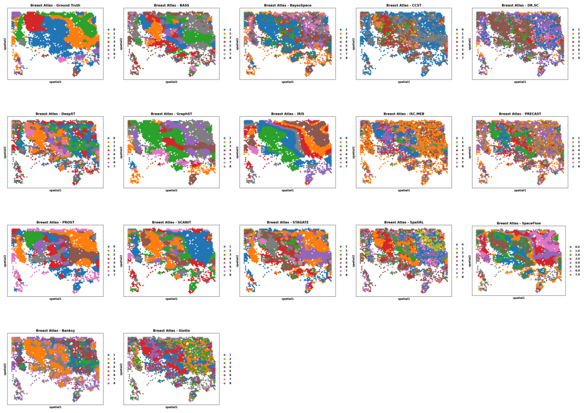

**j** **Chicken Heart**

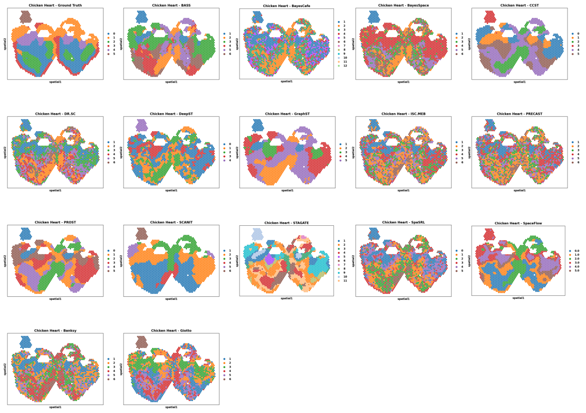

**k** **Intestine A1**

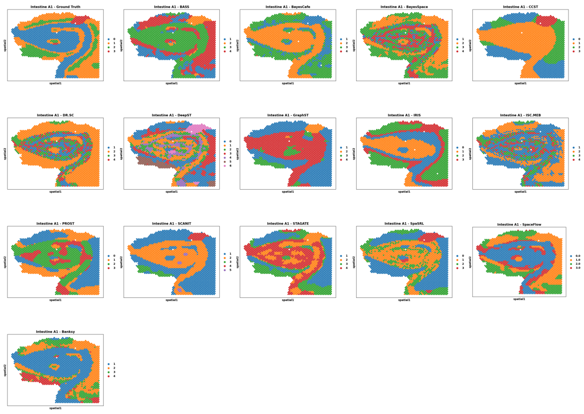

**Intestine A2**

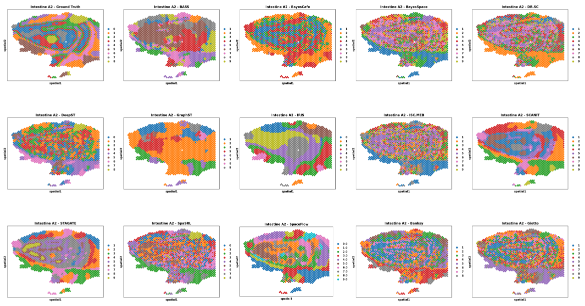

**l** **Prostate Cancer**

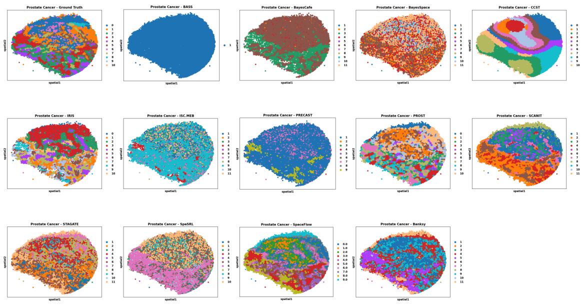

**m** **Cerebellum**

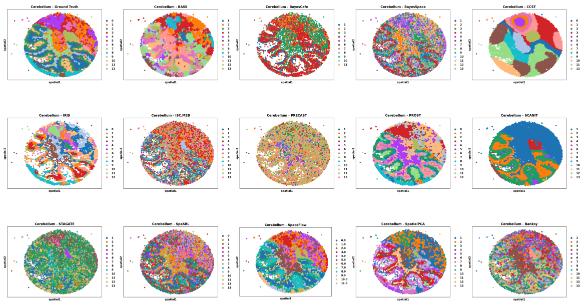

Supplementary Figure 19: Qualitative assessment of spatial domains identified by domain detection methods (BASS, Banksy, BayesCafe, BayesSpace, CCST, DR.SC, DeepST, Giotto, GraphST, iSC.MEB, IRIS, PRECAST, PROST, SCANIT, STAGATE, SpaSRL, SpaceFlow and SpatialPCA) compared to ground truth annotations for 21 real and 15 simulated datasets. Visualization of spatial domains identified by different methods in comparison to ground truth domain for real datasets: a) DLPFC-151507,151508, 151509, 151510, 151669, 151670, 151671, 151672, 151673, 151674, 151675, 151676, b) Embryo - 9.5, 14.5, c) Merfish Mouse brain - 0.04, 0.08, 0.14, 0.19, 0.24, d) osmFISH Mouse brain, e) Mouse Breast Cancer, and simulated datasets: f) Kidney Cancer - 410, 411, 506, g) Breast Cancer - ER+, TNBC, h) Liver Cancer - HCC-1L, HCC-2L, HCC-3L, HCC-4L, i) Human Breast ST8, j) Chicken Heart, k) Intestine A1, A2, l) Prostate Cancer and m) Cerebellum datasets

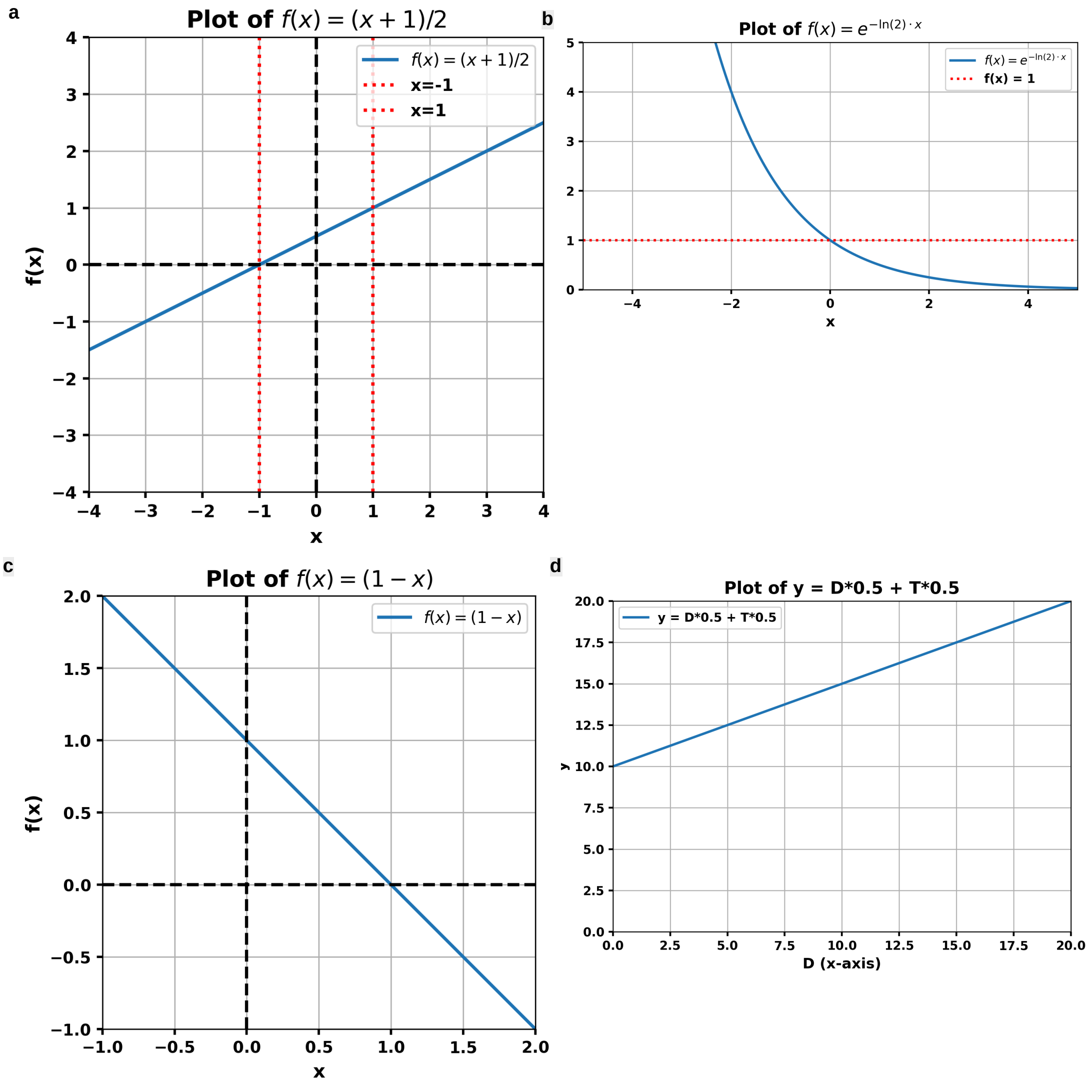


Supplementary Figure 20: Transformation functions were applied to map each metric into the [0, 1] range, ensuring a consistent "higher-is-better" interpretation. The visualizations include: (a) transformation functions for metrics originally ranging in [-1, 1]—including Bivariate Moran's I, Lee's L Statistic, Spatial Pearson Correlation, SSIM, Cosine Similarity, and Pearson Correlation; b) transformation for metrics with range$[0, \infty]$ such as Bivariate Geary's C, c) evaluation metrics bounded in range [0, 1] with "lower-is-better" interpretation such as RMSE and JSD; and d) the formulation used to apply penalties when computing composite scores.

### Supplementary Tables

Supplementary Table 1: Description of Spatial Deconvolution methods

| Sno | Methods | Principle | Github (https://github.com/) | Link |
|---|---|---|---|---|
| 1 | AutoGeneS (1) | Linear model based | theislab/AutoGeneS | Link |
| 2 | CARD (2) | Bayesian model based | YMa-lab/CARD | Link |
| 3 | Cell2location (3) | Bayesian model based | BayraktarLab/cell2location | Link |
| 4 | CellDART (4) | Deep learning based | mexchy1000/CellDART | Link |
| 5 | CelloScope (5) | Bayesian model based | szczurek-lab/Celloscope | Link |
| 6 | DestVI (6) | Deep learning based | scverse/scvi-tools | Link |
| 7 | GraphST (7) | Deep learning based | JinmiaoChenLab/GraphST | Link |
| 8 | Polaris (8) | Deep learning based | JiawenChenn/POLARIS | Link |
| 9 | RCTD (9) | Linear Model based | dmcable/spacexr | Link |
| 10 | Redeconve (10) | Linear Model based | ZxZhou4150/Redeconve | Link |
| 11 | SD (11) | Graph based | leihouyeung/SD2 | Link |
| 12 | Seurat V3 (12) | Anchor based | satijalab/seurat | Link |
| 13 | SONAR (13) | Bayesian model based | lzygenomics/SONAR | Link |
| 14 | SpatialDecon (14) | Bayesian model based | Nanostring-Biostats/SpatialDecon | Link |
| 15 | spatialDWLS (15) | NMF based | RubD/Giotto | Link |
| 16 | SpiceMix (16) | w/o scRNA reference | ma-compbio/SpiceMix | Link |
| 17 | Spotlight (17) | NMF based | MarcElosua/SPOTlight | Link |
| 18 | STdeconvolve (18) | w/o scRNA reference | JEFworks-Lab/STdeconvolve | Link |
| 19 | Stereoscope (19) | Linear Model based | almaan/stereoscope | Link |
| 20 | STRIDE (20) | NMF based | DongqingSun96/STRIDE | Link |
| 21 | Tangram (21) | Deep learning based | broadinstitute/Tangram | Link |

Supplementary Table 2: Description of the 37 datasets used for spatial deconvolution benchmarking.

| Sno | Dataset | Slice ID | Tissue Type | Technology | # of cell types | ST dimensions (spots*genes) | Single-cell Reference |
|---|---|---|---|---|---|---|---|
| 1 | **DLPFC** (22) | 151508 | Brain | 10X Visium | 17 | 4382*11403 | Velmeshev et al. (23) |
| 2 | | 151509 | Brain | 10X Visium | 17 | 4787*11403 | |
| 3 | | 151669 | Brain | 10X Visium | 17 | 3660*11403 | |
| 4 | | 151670 | Brain | 10X Visium | 17 | 3497*11403 | |
| 5 | | 151673 | Brain | 10X Visium | 17 | 3639*11403 | |
| 6 | | 151674 | Brain | 10X Visium | 17 | 3673*11403 | |
| 7 | **Mouse Brain** (3) | ST48 | Brain | 10X Visium | 59 | 2328*12152 | Kleshchevnikov et al. (3) |
| 8 | | ST52 | Brain | 10X Visium | 59 | 2328*12152 | |
| 9 | **Hippocampus** (24) | | Brain | SlideseqV2 | 22 | 41798*11702 | Stickels et al. (24) |
| 10 | **Cerebellum** (24) | | Brain | SlideseqV2 | 11 | 27196*11808 | Stickels et al. (24) |
| 11 | **Mouse Brain** (25) | | Brain | 10X Visium HD | 7 | 98569*10130 | Kleshchevnikov et al. (3) |
| 12 | **Kidney Cancer** (26) | 410 | Cancer | 10X Visium | 12 | 2917*9648 | Li et al. (26) |
| 13 | | 411 | Cancer | 10X Visium | 12 | 2048*9648 | |
| 14 | | 506 | Cancer | 10X Visium | 12 | 1790*9648 | |
| 15 | **Breast Cancer** (27) | ER+ | Cancer | 10X Visium | 22 | 2432*11854 | Wu et al. (27) |
| 16 | | TNBC | Cancer | 10X Visium | 22 | 1162*12353 | |
| 17 | **Liver Cancer HCC**(28) | 1L | Cancer | 10X Visium | 29 | 2765*7093 | Wu et al. (29) |
| 18 | | 2L | Cancer | 10X Visium | 29 | 4670*7093 | |
| 19 | | 3L | Cancer | 10X Visium | 29 | 4757*7093 | |
| 20 | | 4L | Cancer | 10X Visium | 29 | 4113*7093 | |
| 21 | **Prostate Cancer** (30) | | Cancer | SlideseqV2 | 30 | 23945*24728 | Hirz et al. (30) |
| 22 | **Lung Cancer** (25) | | Cancer | 10X Visium HD | 24 | 150589*17797 | Salcher et al. (31) |
| 23 | **Breast Atlas** (32) | ST_1 | Organs | 10X Visium | 10 | 2103*32664 | Kumar et al. (32) |
| 24 | | ST_8 | Organs | 10X Visium | 10 | 2801*32664 | |
| 25 | **Intestine** (33) | A1 | Organs | 10X Visium | 9 | 2647*33514 | Fawkner-Corbett et al. (33) |
| 26 | | A2 | Organs | 10X Visium | 9 | 2315*13541 | |
| 27 | **Chicken Heart** (34) | | Organs | 10X Visium | 14 | 1915*3688 | Mantri et al. (34) |
| 28 | **Dev. Lung** (35) | | Organs | 10X Visium | 27 | 3044*20374 | Sountoulidis et al. (35) |
| 29 | **Liver Atlas Mouse** (36) | 4 | Organs | 10X Visium | 17 | 1790*31017 | Guilliams et al. (36) |
| 30 | | 12 | Organs | 10X Visium | 17 | 1768*31017 | |
| 31 | **Liver Atlas Human** (36) | 18 | Organs | 10X Visiu | 17 | 1759*32643 | Guilliams et al. (36) |
| 32 | **Kidney Atlas** (37) | 1 | Organs | SlideseqV2 | 26 | 20852*16584 | Lake et al. (37) |
| 33 | | 6 | Organs | SlideseqV2 | 26 | 30296*16799 | |
| 34 | **Mouse brain** (38) | | Image-based | MERFISH | 10 | 7907*27998 | Moffitt et al. (38) |
| 35 | **Ileum** (39) | | Image-based | MERFISH | 16 | 2445*242 | |
| 36 | **Breast cancer**(40) | | Image-based | MERFISH | 9 | 33932*500 | |
| 37 | **Lung cancer**(40) | | Image-based | MERFISH | 13 | 13792*488 | |

Supplementary Table 3: Summary table of rare and regional cell types identified for all the datasets

| Sno | Datasets | Tissue | % | Rare cell types | Regional rare cell types |
|---|---|---|---|---|---|
| 1 | **Kidney_Cancer_410** | Cancer | 50 | Mast | Mast |
| 2 | **Kidney_Cancer_411** | Cancer | 50 | Mast | pDC, NK |
| 3 | **Kidney_Cancer_506** | Cancer | 50 | pDC | Epi_non-PT |
| 4 | **Liver_Cancer_1L** | Cancer | 20 | CD8+ Cells 1, Mast Cells, CD8+ Cells 2 | CD8+ Cells 1, CD8+ Cells 2, CD4+ Cells 1, Nk cells 2, CD4+ Cells 2 |
| 5 | **Liver_Cancer_2L** | Cancer | 20 | CD8+ Cells 1, Endothelial Cells 4, Mast Cells | CD8+ Cells 1 |
| 6 | **Liver_Cancer_3L** | Cancer | 20 | CD4+ Cells 3, CD4+ Cells 1, CD8+ Cells 1 | CD4+ Cells 3, CD4+ Cells 1, CD8+ Cells 1, CD8+ Cells 2 |
| 7 | **Liver_Cancer_4L** | Cancer | 20 | CD8+ Cells 1, Mast Cells, CD4+ Cells 1 | CD8+ Cells 1, CD4+ Cells 1, CD8+ Cells 2, Nk cells 3, CD4+ Cells 2 |
| 8 | **Prostate_Cancer** | Cancer | 30 | Mono3, Macrophage1, Mono1 | PDC |
| 9 | **Breast_Cancer_TNBC_CID44971** | Cancer | 30 | NK cells, NKT cells | Cycling T-cells, DCs |
| 10 | **Breast_Cancer_ER+_CID4290** | Cancer | | NKT cells, Cycling T-cells | NA |
| 11 | **Lung_Cancer_Visium_HD** | Cancer | | DC mature, Ciliated | NA |
| 12 | **Developmental_Lung** | Organs | 30 | neuro_SCP, im_megakaryocyte, neuro_proliferating | neuro_SCP, im_megakaryocyte, mes_chondroblast |
| 13 | **Breast_Atlas_ST1** | Organs | | tcells | NA |
| 14 | **Breast_Atlas_ST8** | Organs | 30 | tcells | lumhr |
| 15 | **Intestine_A1** | Organs | 70 | Fibroblasts | Epithelium |
| 16 | **Intestine_A2** | Organs | 30 | Pericytes | Pericytes, Neural |
| 17 | **Chicken_Heart** | Organs | 90 | Epi-mesenchymal cells | MT-enriched cardiomyocytes, Immature myocardial cells |
| 18 | **Liver Atlas Human - 18** | Organs | | pDCs, B cells | NA |
| 19 | **Liver Atlas Mouse - 4** | Organs | | Basophils, NK cells | NA |
| 20 | **Liver Atlas Mouse - 12** | Organs | 100 | ILC1s, Basophils | Cholangiocytes |
| 21 | **Kidney Atlas - 1** | Organs | 50 | neural cell, conventional dendritic cell, plasmacytoid dendritic cell, human | podocyte |
| 22 | **Kidney Atlas - 6** | Organs | 30 | neural cell, conventional dendritic cell, podocyte | podocyte |
| 23 | **DLPFC_151508** | Brain | 30 | OPC, Microglia | OPC, Microglia, IN_SV2C, L5_6_CC, |
| 24 | **DLPFC_151509** | Brain | 30 | IN_SV2C, OPC | IN_SV2C, Microglia, L5_6, |
| 25 | **DLPFC_151669** | Brain | 30 | Microglia, IN_SV2C | L4 |
| 26 | **DLPFC_151670** | Brain | 40 | Microglia, IN_SV2C | IN_PV |
| 27 | **DLPFC_151673** | Brain | 30 | Microglia, IN_SV2C | Microglia, IN_SV2C, OPC, IN_SST, |
| 28 | **DLPFC_151674** | Brain | 30 | Microglia, IN_SV2C | IN_SV2C, OPC, IN_SST, L5_6_CC |
| 29 | **Mouse Brain - ST48** | Brain | 20 | Ext_Unk_1, Unk_2, Nb_1, Astro_CTX, Inh_1, Inh_Meis2_1 | Ext_Unk_1, Nb_1, Astro_CTX, Inh_1, Inh_Meis2_1, Astro_THAL_med, LowQ_2, Ext_Hpc_CA2, Ext_L6 |
| 30 | **Mouse Brain - ST52** | Brain | 20 | Unk_2, Ext_Unk_1, Nb_1, LowQ_2, Inh_1, Astro_CTX | Ext_Unk_1, LowQ_2, Inh_1, Astro_CTX, Inh_Meis2_1, Ext_L5_3, Inh_Lamp5, Ext_L6, Astro_THAL_med |
| 31 | **Hippocampus** | Brain | 30 | 21, 19 | 18, 19 |
| 32 | **Cerebellum** | Brain | 30 | 9 | 5 |
| 33 | **Mouse_brain_VisiumHD** | Brain | 40 | Endo | Excitatory |
| 34 | **Merfish-brain** | Image_based | 30 | Pericytes | Microglia |
| 35 | **Merfish-ileum** | Image_based | 45 | Tuft | Endothelial + Neuronal |
| 36 | **Merfish_Lung_Cancer** | Image_based | 30 | NK_cells | Dendritic_cells, Endothelial_cells |
| 37 | **Merfish_Breast_Cancer** | Image_based | 30 | Dendritic_cells | T/NK_Cells, Dendritic_cells |

Supplementary Table 4: Summary table of cell types identified with high curl scores for all datasets

| **Sno** | Datasets | **Tissue** | **High curl cell types** |
|---|---|---|---|
| 1 | **Kidney_Cancer** | Cancer | Epi_PT, Epi_non-PT, Plasma |
| 2 | **Liver_Cancer** | Cancer | B cells, Bi-Potent Cells, Endothelial Cells 1, Hepatocytes 3, Mast Cells, Myeloid 2, Nk cells 2 |
| 3 | **Prostate_Cancer** | Cancer | Pericytes-2, TNK, Th1, Th17, Treg, Tumor, mDC, plasma cells |
| 4 | **Breast_Cancer ER+** | Cancer | Endothelial CXCL12, Luminal Progenitors, Mature Luminal, Monocyte, NK cells, NKT cells |
| | **Breast_Cancer TNBC** | Cancer | Endothelial CXCL12, Luminal Progenitors, Mature Luminal, Monocyte, NK cells, NKT cells |
| 5 | **Lung_Cancer_Visium_HD** | Cancer | Endothelial cell, Tumor cells, Plasma cell, Stromal, T cell CD8, T cell regulatory |
| 6 | **Developmental_Lung** | Organs | endo_bronchial, im_immature, im_myeloid_progenitor, mes_chondroblast, mes_immature_ASM, mes_mesothelial, neuro_SCP |
| 7 | **Breast_Atlas** | Organs | pericytes, vascular |
| 8 | **Intestine** | Organs | Immune, Mesothelium |
| 9 | **Chicken_Heart** | Organs | Endocardial cells, Epi-epithelial cells, Erythrocytes, Macrophages |
| 10 | **Liver Atlas Human** | Organs | Circulating NK/NKT, pDCs, Mono+mono derived cells, Fibroblasts |
| 11 | **Liver Atlas Mouse** | Organs | B cells, NK cells, ILC1s, Kupffer cells |
| 12 | **Kidney Atlas - 1** | Organs | kidney loop of Henle thin ascending limb epithelial cell, papillary tips cell, parietal epithelial cell, plasma cell, plasmacytoid dendritic cell, human, renal interstitial pericyte |
| 13 | **DLPFC** | Brain | IN_PV, Neu_mat, OPC, Oligodendrocytes |
| 14 | **Mouse Brain** | Brain | Astro_AMY_CTX, Ext_Amy_2, Ext_L5_3, Ext_L6, Ext_Med, Ext_Pir, Ext_Unk_1, Inh_1, Inh_Lamp5, Inh_Meis2_1, Inh_Sst, Inh_Vip, Micro, Nb_1, Unk_2 |
| 15 | **Hippocampus** | Brain | 11, 5, 6, 7, 8, 9 |
| 16 | **Cerebellum** | Brain | 3, 8, ,9 |
| 17 | **Mouse_brain_VisiumHD** | Brain | Astrocytes, Micro |
| 18 | **Merfish-brain** | Image-based | Inhibitory |
| 19 | **Merfish-ileum** | Image-based | Smooth Muscle |
| 20 | **Merfish_Lung_Cancer** | Image-based | Macrophages1, T_cells |
| 21 | **Merfish_Breast_Cancer** | Image-based | Fibroblasts1, TAM_C1Q |

Supplementary Table 5: Summary table of cell types identified with high and low elongation scores for all datasets

| **Sno** | Datasets | **Tissue** | **High elongation cell types** | **Low elongation cell types** |
|---|---|---|---|---|
| 1 | **Kidney_Cancer** | Cancer | Epi_PT,Mast,pDC | B-cell, Myeloid, Plasma |
| 2 | **Liver_Cancer** | Cancer | CD4+ Cells 1,CD4+ Cells 2,CD8+ Cells 2, Hepatocytes 1,Hepatocytes 4,Hepatocytes 5,Nk cells 4 | B cells, CD4+ Cells 3, Endothelial Cells 1, Fibroblasts 1, Hepatocytes 3, Myeloid 1, Myeloid 2 |
| 3 | **Prostate_Cancer** | Cancer | B cells,CTL-2,Epitheial_Club, Epitheial_Hillock,PDC,Th17,Treg,Tumor | Endothelial cells-1, Epitheial_Luminal, Macrophage1, Mono2, Mono3, NK, Pericytes-1, mDC |
| 4 | **Breast_Cancer ER+** | Cancer | DCs,Endothelial ACKR1, Endothelial Lymphatic LYVE1,Macrophage,Monocyte | Cancer LumB SC, Mature Luminal, NK cells, NKT cells, T cells CD8+ |
| | **Breast_Cancer TNBC** | Cancer | Cancer Cycling,DCs,Endothelial ACKR1, Endothelial Lymphatic LYVE1,Macrophage,Monocyte | Cancer Basal SC, Mature Luminal, NK cells, NKT cells, T cells CD8+ |
| 5 | **Lung_Cancer_Visium_HD** | Cancer | Club,DC mature,T cell CD4,cDC1,cDC2,other | Endothelial cell, Neutrophils, Plasma cell, T cell CD8, Tumor cells, B cell |
| 6 | **Developmental_Lung** | Organs | endo_arterial,endo_immature,epi_Dismal, epi_proliferating,mes_immature,mes_pericyte, mes_proliferating | endo_bronchial, endo_venous, im_immature, im_mature, im_myeloid_progenitor, mes_adventitial_fibro, mes_immature_ASM |
| 7 | **Breast_Atlas** | Organs | bcells, myeloid | lymphatic, vascular |
| 8 | **Intestine** | Organs | Endothelium, Fibroblasts | Muscularis, Pericytes |
| 9 | **Chicken_Heart** | Organs | Cardiomyocytes-1,Cardiomyocytes-2, Mural cells,Valve cells | Endocardial cells, Fibroblast cells, MT-enriched cardiomyocytes, Macrophages |
| 10 | **Liver Atlas Human** | Organs | B cells,Cholangiocytes,Mig.cDCs,T cells | Fibroblasts, Resident NK, cDC1s, pDCs |
| 11 | **Liver Atlas Mouse** | Organs | Cholangiocytes,Hepatocytes,HsPCs,Neutrophils | Basophils, Endothelial cells, NK cells, cDC2s |
| 12 | **Kidney Atlas - 1** | Organs | T cell,endothelial cell,epithelial cell of proximal tubule, kidney collecting duct intercalated cell, kidney collecting duct principal cell, kidney connecting tubule epithelial cell | B cell, conventional dendritic cell, kidney interstitial alternatively activated macrophage, lymphocyte, mononuclear phagocyte, plasmacytoid dendritic cell, human |
| 13 | **DLPFC** | Brain | L5_6, L5_6_CC, Neu_NRGN_II, OPC | AST_PP, Endothelial, IN_SST, Oligodendrocytes |
| 14 | **Mouse Brain** | Brain | Astro_AMY,Astro_CTX,Astro_STR,Ext_ClauPyr, Ext_Hpc_CA1,Ext_L23,Ext_L25,Ext_L56,Ext_L5_1, Ext_L5_2,Ext_L6B,Ext_Unk_2,Inh_Pvalb,LowQ_2,Unk_1 | Astro_THAL_hab, Endo, Ext_Hpc_DG1, Ext_Unk_1, Inh_1, Inh_2, Inh_6, Inh_Lamp5, Inh_Meis2_1, Inh_Sst, Inh_Vip, LowQ_1, OPC_2, Oligo_1, Oligo_2 |
| 15 | **Hippocampus** | Brain | 2,20,21,3,4,9 | 11, 12, 17, 19, 22, 7 |
| 16 | **Cerebellum** | Brain | 10, 2, 2007 | 11, 5, ,9 |
| 17 | **Mouse_brain_VisiumHD** | Brain | Excitatory, Inhibitory | Astrocytes, Micro |
| 18 | **Merfish-brain** | Image-based | OD Mature | Inhibitory |
| 19 | **Merfish-ileum** | Image-based | Smooth Muscle | Enterocyte (Mid Villus) |
| 20 | **Merfish_Lung_Cancer** | Image-based | Stromal_cells, T_cells | Neutrophils_cells, Endothelial_cells |
| 21 | **Merfish_Breast_Cancer** | Image-based | Endothelial_cells, T/NK_Cells | TAM_SPP1, Fibroblasts1 |

#### Supplementary Table 6: Summary table of cell types identified with high and low linearity scores for all datasets

| Sno | Datasets | Tissue | High linearity cell types | Low linearity cell types |
|---|---|---|---|---|
| 1 | **Kidney_Cancer** | Cancer | Myeloid,T-cell,pDC | EC,Epi_PT,Mast |
| 2 | **Liver_Cancer** | Cancer | CD8+ Cells 2,Endothelial Cells 4,Hepatocytes 1, Hepatocytes 4,Hepatocytes 5,Hepatocytes 7,Tregs | B cells,Bi-Potent Cells,CD4+ Cells 3, Endothelial Cells 5,Fibroblasts 1,Mast Cells,Nk cells 3 |
| 3 | **Prostate_Cancer** | Cancer | Mono3,Naive Th,Pericytes-1,TNK,Th1,Th17,Treg,Tumor | Endothelial cells-1,Epitheial_Luminal,Macrophage2, Macrophage3,Mono1,Mono2,NK,plasma cells |
| 4 | **Breast_Cancer ER+** | Cancer | Cancer LumA SC,DCs,Endothelial ACKR1, Endothelial Lymphatic LYVE1,Macrophage,PVL Immature | B cells Naive,Cancer LumB SC,Cycling T-cells, NK cells,T cells CD4+,T cells CD8+ |
| | **Breast_Cancer TNBC** | Cancer | DCs,Endothelial ACKR1,Endothelial Lymphatic LYVE1, Macrophage,PVL Immature | B cells Naive,Cycling T-cells,NK cells, T cells CD4+,T cells CD8+ |
| 5 | **Lung_Cancer_Visium_HD** | Cancer | DC mature,Endothelial cell,T cell CD4, T cell CD8,T cell regulatory,cDC1 | Alveolar cell type 2,B cell,Club,Neutrophils, Tumor cells,cDC2 |
| 6 | **Developmental_Lung** | Organs | endo_venous,epi_progenitor_population, im_dendritic,im_mature, mes_adventitial_fibro,mes_immature,mes_proliferating | endo_capillary,endo_lymphatic,erythro,mes_chondroblast, mes_immature_ASM,neuro_SCP,neuro_neuronal |
| 7 | **Breast_Atlas** | Organs | bcells,lymphatic | tcells,vascular |
| 8 | **Intestine** | Organs | Epithelium,Myofibroblasts | Fibroblasts,Neural |
| 9 | **Chicken_Heart** | Organs | Cardiomyocytes-2,Erythrocytes,Mural cells,Valve cells | Epi-epithelial cells,Fibroblast cells, MT-enriched cardiomyocytes,TMSB4X high cells |
| 10 | **Liver Atlas Human** | Organs | Cholangiocytes,Mig.cDCs,T cells,pDCs | Endothelial cells,Macrophages, Neutrophils,Plasma cells |
| 11 | **Liver Atlas Mouse** | Organs | Fibroblasts,Hepatocytes,HsPCs,Kupffer cells | Basophils,Mig. cDCs, Monocytes & Monocyte-derived cells,cDC1s |
| 12 | **Kidney Atlas - 1** | Organs | T cell,endothelial cell,epithelial cell of proximal tubule, kidney collecting duct intercalated cell, kidney connecting tubule epithelial cell, kidney interstitial fibroblast | kidney distal convoluted tubule epithelial cell, kidney loop of Henle thick ascending limb epithelial cell, kidney loop of Henle thin ascending limb epithelial cell,mast cell, podocyte,renal interstitial pericyte |
| 13 | **DLPFC** | Brain | IN_SV2C,L2_3,Neu_NRGN_II,OPC | AST_FB,IN_PV,L4,Neu_NRGN_I |
| 14 | **Mouse Brain** | Brain | Ext_Hpc_DG1,Ext_L23,Ext_L5_1,Ext_L6B, Ext_Thal_1,Ext_Thal_2,Ext_Unk_1,Inh_1,Inh_Lamp5, Inh_Meis2_1,Inh_Meis2_3,Inh_Pvalb,Inh_Sst,Unk_1,Unk_2 | Astro_HYPO,Astro_THAL_hab,Astro_THAL_lat, Astro_WM,Ext_L56,Ext_Unk_3,Inh_3,Inh_4,Inh_5, Inh_6,Inh_Meis2_4,LowQ_1,OPC_1,OPC_2,Oligo_1 |
| 15 | **Hippocampus** | Brain | 10,11,15,20,3,9 | 12,14,17,18,19,7 |
| 16 | **Cerebellum** | Brain | 3, 7, 2008 | 4,6,9 |
| 17 | **Mouse_brain_VisiumHD** | Brain | Micro, Other | Endo,Inhibitory |
| 18 | **Merfish-brain** | Image-based | OD Mature | Inhibitory |
| 19 | **Merfish-ileum** | Image-based | Smooth Muscle | Enterocyte (Mid Villus) |
| 20 | **Merfish_Lung_Cancer** | Image-based | Stromal_cells, T_cells | Endothelial_cells, Neutrophils_cells |
| 21 | **Merfish_Breast_Cancer** | Image-based | TAM_C1Q, TAM_SPP1 | Cancer_subtype2, Fibroblasts1 |

Supplementary Table 7: Summary table of cell types identified across all datasets, categorized by major brain cell type classes. The table organizes identified cell types under key brain cell type categories: Astrocytes, Endothelial Cells, Macrophages, Excitatory Neurons, Inhibitory Neurons, Oligodendrocyte Progenitor Cells (OPCs), and Oligodendrocytes.

| **Dataset** | **DLPFC** | **Mouse_brain** | **Mouse_brain_VisiumHD** |
|---|---|---|---|
| Astrocytes | AST_FB, AST_PP | Astro_AMY, Astro_AMY_CTX, Astro_CTX, Astro_HPC, Astro_HYPO, Astro_STR, Astro_THAL_hab, Astro_THAL_lat, Astro_THAL_med, Astro_WM | Astrocytes |
| Endothelial_cells | Endothelial | Endo | Endo |
| Macrophages | Microglia | Micro | Micro |
| Neurons Excitatory | Neu_NRGN_I, Neu_NRGN_II, Neu_mat | Ext_Amy_1, Ext_Amy_2, Ext_ClauPyr, Ext_Hpc_CA1, Ext_Hpc_CA2, Ext_Hpc_CA3, Ext_Hpc_DG1, Ext_Hpc_DG2, Ext_L23, Ext_L25, Ext_L56, Ext_L5_1, Ext_L5_2, Ext_L5_3, Ext_L6, Ext_L6B, Ext_Med, Ext_Pir, Ext_Thal_1, Ext_Thal_2, Ext_Unk_1, Ext_Unk_2, Ext_Unk_3 | Excitatory |
| Neurons Inhibitory | IN_PV, IN_SST, IN_SV2C, IN_VIP | Nb_1, Nb_2, Inh_1, Inh_2, Inh_3, Inh_4, Inh_5, Inh_6, Inh_Lamp5, Inh_Meis2_1, Inh_Meis2_2, Inh_Meis2_3, Inh_Meis2_4, Inh_Pvalb, Inh_Sst, Inh_Vip | Inhibitory |
| Oligodendrocytes Progenitor cells (OPCs) | OPC | OPC_1, OPC_2 | |
| Oligodendrocytes | Oligodendrocytes | Oligo_1, Oligo_2 | Oligo_OPC |

Supplementary Table 8: Summary table of cell types identified across all datasets, categorized by major cancer tissue cell type classes. The table organizes identified cell types under key cancer cell type categories: B cells, Endothelial cells, Epithelial cells, Fibroblasts, Myeloid cells, NK cells, Plasma cells, T cells, Cancer cells and Dendritic cells.

| **Dataset** | **Kidney_Cancer** | **Breast_Cancer_TNBC** | **Breast_Cancer_ER+** | **Liver_Cancer** | **Prostate_Cancer** | **Lung_cancer_VisiumHD** |
|---|---|---|---|---|---|---|
| B_cells | B_cell | B_cells_Memory, B_cells_Naive | B_cells_Memory, B_cells_Naive | B_cells | B_cells | B_cell |
| Endothelial_cells | EC | Endothelial_ACKR1, Endothelial_CXCL12, Endothelial_Lymphatic_LYVE1, Endothelial_RGS5 | Endothelial_ACKR1, Endothelial_CXCL12, Endothelial_Lymphatic_LYVE1, Endothelial_RGS5 | Endothelial_Cells_1, Endothelial_Cells_2, Endothelial_Cells_3, Endothelial_Cells_4, Endothelial_Cells_5 | Endothelial_cells_1, Endothelial_cells_2 | Endothelial_cell |
| Epithelial_cells | Epi_non_PT, Epi_PT | Myoepithelial, Luminal_Progenitors, Mature_Luminal | Myoepithelial, Luminal_Progenitors, Mature_Luminal | Bi_Potent_Cells | Epitheial_Basal, Epitheial_Club, Epitheial_Hillock, Epitheial_Luminal | Alveolar_cell_type_1, Alveolar_cell_type_2, Ciliated, Club, transitional_club_AT2 |
| Fibroblasts | Fibro | | | Fibroblasts_1 | Fibroblasts, Pericytes_1, Pericytes_2 | Stromal |
| Myeloid_cells | Myeloid, Mast | Macrophage, Monocyte | Macrophage, Monocyte | Mast_Cells, Myeloid_1, Myeloid_2, Myeloid_3 | Macrophage1, Macrophage2, Macrophage3, Mast_cells, Mono1, Mono2, Mono3 | Macrophage, Macrophage_alveolar, Mast_cell, Monocyte, Neutrophils |
| NK_cells | NK | NK_cells | NK_cells | Nk_Cells_1, Nk_cells_2, Nk_cells_3, Nk_cells_4 | NK | NK_cell |
| Plasma_cells | Plasma | PVL_Differentiated, PVL_Immature, Plasmablasts | PVL_Differentiated, PVL_Immature, Plasmablasts | | plasma_cells | Plasma_cell |
| T_cells | T_cell | NKT_cells, T_cells_CD4_, T_cells_CD8_, Cycling_T_cells | NKT_cells, T_cells_CD4_, T_cells_CD8_, Cycling_T_cells | CD4__Cells_1, CD4__Cells_2, CD4__Cells_3, CD8__Cells_1, CD8__Cells_2, Tregs | CTL_1, CTL_2, Cycling_T, Naive_Th, Treg, TNK, Th1, Th17 | T_cell_CD4, T_cell_CD8, T_cell_regulatory |
| Cancer_cells | RCC | Cancer_Basal_SC, Cancer_Cycling | Cancer_LumA_SC, Cancer_LumB_SC | Hepatocytes_1, Hepatocytes_2, Hepatocytes_3, Hepatocytes_4, Hepatocytes_5, Hepatocytes_6, Hepatocytes_7 | Tumor | Tumor_cells |
| Dendritic_cells | pDC | DCs | DCs | | PDC, mDC | DC_mature, cDC1, cDC2, pDC |

Supplementary Table 9: Summary table of cell types identified across all datasets, categorized by major organs tissue cell type classes. The table organizes identified cell types under key organs cell type categories: B cells, Endothelial cells, Epithelial cells, Fibroblasts, Myeloid cells, NK cells, Plasma cells, T cells, Dendritic cells, Smooth muscle cells and Neural cells.

| **Dataset** | **Breast_Atlas** | **Intestine** | **Liver_Atlas_Mouse** | **Liver_Atlas_Human** | **Kidney_Atlas** | **Developmental_Lung** | **ChickenHeart** |
|---|---|---|---|---|---|---|---|
| B_cells | bcells | | B_cells | B_cells | B_cell | | |
| Endothelial_cells | | Endothelium | Endothelial_cells | Endothelial_cells | endothelial_cell | endo_arterial, endo_bronchial, endo_capillary, endo_immature, endo_lymphatic, endo_proliferating, endo_venous | Endocardial_cells, Vascular_endothelial_cells |
| Epithelial_cells | lumhr, lumsec, basal | Epithelium, Mesothelium | Cholangiocytes | Cholangiocytes | epithelial_cell_of_proximal_tubule, kidney_collecting_duct_intercalated_cell, kidney_connecting_tubule_epithelial_cell, kidney_distal_convoluted_tubule_epithelial_cell, kidney_loop_of_Henle_thick_ascending_limb_epithelial_cell, kidney_loop_of_Henle_thin_ascending_limb_epithelial_cell, kidney_loop_of_Henle_thin_descending_limb_epithelial_cell, parietal_epithelial_cell, papillary_tips_cell, podocyte | epi_Dismal, epi_progenitor_population, epi_proliferating | Epi_epithelial_cells, Epi_mesenchymal_cells |
| Fibroblasts | fibroblasts, pericytes | Fibroblasts, Myofibroblasts | Fibroblasts | Fibroblasts | kidney_interstitial_fibroblast, renal_interstitial_pericyte | mes_ASM, mes_adventitial_fibro, mes_chondroblast, mes_immature, mes_immature_ASM, mes_mesothelial, mes_pericyte, mes_proliferating | Fibroblast_cells |
| Myeloid_cells | myeloid | | Kupffer_cells, Monocytes___Monocyte_derived_cells, Neutrophils | Macrophages, Mono_mono_derived_cells, Neutrophils | mast_cell, kidney_interstitial_alternatively_activated_macrophage, non_classical_monocyte, mononuclear_phagocyte, neutrophil | im_myeloid_progenitor | Macrophages |
| NK_cells | | | NK_Cells | Resident_NK | | | |
| Plasma_cells | | | | Plasma_cells | plasma_cell | | |
| T_cells | tcells, lymphatic | Immune | T_cells, Basophils, ILC1s | T_cells, Circulating_NK_NKT, Basophils | T_cell, lymphocyte | im_immature, im_mature, im_megakaryocyte | |
| Dendritic_cells | | | Mig__cDCs, cDC1s, cDC2s, pDCs | Mig__cDCs, cDC1s, cDC2s, pDCs | conventional_dendritic_cell, plasmacytoid_dendritic_cell__human | im_dendritic | |
| Smooth_muscle_cells | vascular | Pericytes, Muscularis | | | | | Cardiomyocytes_1, Cardiomyocytes_2, Immature_myocardial_cells, MT_enriched_cardiomyocytes, Mural_cells |
| Neural cells | | Neural | | | neural_cell | neuro_SCP, neuro_neuronal, neuro_proliferating | |

Supplementary Table 10: Description of Spatial Domain Detection methods

| Sno | Methods | Principle | Github (https://github.com/) | Link |
|---|---|---|---|---|
| 1 | Banksy (41) | Spatial kernel based | prabhakarlab/Banksy | Link |
| 2 | BASS (42) | Bayesian model | zhengli09/BASS | Link |
| 3 | BayesSpace (43) | Bayesian model | edward130603/BayesSpace | Link |
| 4 | BayesCafe (44) | Bayesian model | huimin230/BayesCafe | Link |
| 5 | CCST (45) | GCN + Contrastive Learning | xiaoyeye/CCST | Link |
| 6 | DeepST (46) | GCN + Contrastive Learning | JiangBioLab/DeepST | Link |
| 7 | DRSC (47) | HMRF-based | feiyoung/DR.SC | Link |
| 8 | Giotto (48) | HMRF-based | RubD/Giotto_site | Link |
| 9 | GraphST (7) | GCN + Contrastive Learning | JinmiaoChenLab/GraphST | Link |
| 10 | IRIS (49) | Reference based | YMa-lab/IRIS | Link |
| 11 | iSCMEB (50) | HMRF-based | XiaoZhangryy/iSC.MEB | Link |
| 12 | PRECAST (51) | Bayesian model | feiyoung/PRECAST | Link |
| 13 | PROST (52) | Graph Attention Networks | Tang-Lab-super/PROST | Link |
| 14 | SCANIT (53) | Graph Deep Learning | zcang/SCAN-IT | Link |
| 15 | SpaceFlow (54) | GCN + Contrastive Learning | hongleir/SpaceFlow | Link |
| 16 | SpaSRL (55) | Graph Deep Learning | zccqq/SpaSRL | Link |
| 17 | SpatialPCA (56) | Graph Deep Learning | shangll123/SpatialPCA | Link |
| 18 | STAGATE (57) | Graph Attention Networks | zhanglabtools/STAGATE | Link |

Supplementary Table 11: Description of the datasets used for spatial domain detection benchmarking

| **Sno** | **Data** | **SliceID** | **Type** | **Tissue Type** | **Technology** | **# of domains** | **ST dimensions** | **Single-cell Reference** |
|---|---|---|---|---|---|---|---|---|
| 1 | **DLPFC** (22) | 151507 | Real | Brain | 10X Visium | 7 | 4226*11403 | Velmeshev et al. (23) |
| 2 | | 151508 | Real | Brain | 10X Visium | 7 | 4384*11403 | |
| 3 | | 151509 | Real | Brain | 10X Visium | 7 | 4789*11403 | |
| 4 | | 151510 | Real | Brain | 10X Visium | 7 | 4634*11403 | |
| 5 | | 151669 | Real | Brain | 10X Visium | 5 | 3661*11403 | |
| 6 | | 151670 | Real | Brain | 10X Visium | 5 | 3498*11403 | |
| 7 | | 151671 | Real | Brain | 10X Visium | 5 | 4110*11403 | |
| 8 | | 151672 | Real | Brain | 10X Visium | 5 | 4015*11403 | |
| 9 | | 151673 | Real | Brain | 10X Visium | 7 | 3639*11403 | |
| 10 | | 151674 | Real | Brain | 10X Visium | 7 | 3673*11403 | |
| 11 | | 151675 | Real | Brain | 10X Visium | 7 | 3592*11403 | |
| 12 | | 151676 | Real | Brain | 10X Visium | 7 | 3460*11403 | |
| 13 | **Embryo** (58) | 9.5 | Real | Whole | Stereoseq | 12 | 5913*25568 | Farrell et al. (59) |
| 14 | | 14.5 | Real | Whole | Stereoseq | 28 | 102519*28463 | |
| 15 | **Mouse breast cancer** (7) | | Real | Cancer | Merfish | 8 | 1950*32285 | Yeo et al. (60) |
| 16 | **Mouse brain** (38) | 0.4 | Real | Brain | Merfish | 8 | 5488*155 | Kleshchevnikov et al. (3) |
| 17 | | 0.9 | Real | Brain | Merfish | 8 | 5926*155 | |
| 18 | | 1.4 | Real | Brain | Merfish | 8 | 5557*155 | |
| 19 | | 1.9 | Real | Brain | Merfish | 8 | 5803 *155 | |
| 20 | | 0.24 | Real | Brain | Merfish | 8 | 5543*155 | |
| 21 | **Mouse cortex** (61)) | | Real | Brain | Osmfish | 11 | 4839*33 | Zeisel et al. (62) |
| 22 | **Kidney Cancer** (26) | 410 | Simulated | Cancer | 10X Visium | 11 | 2917*9648 | Li et al. (26) |
| 23 | | 411 | Simulated | Cancer | 10X Visium | 8 | 2048*9648 | |
| 24 | | 506 | Simulated | Cancer | 10X Visium | 9 | 1790*9648 | |
| 25 | **Breast Cancer** (27) | ER+ | Simulated | Cancer | 10X Visium | 10 | 2432*11854 | Wu et al. (27) |
| 26 | | TNBC | Simulated | Cancer | 10X Visium | 8 | 1162*12353 | |
| 27 | **Liver Cancer HCC** (28) | 1L | Simulated | Cancer | 10X Visium | 5 | 2765*7093 | Wu et al. (29) |
| 28 | | 2L | Simulated | Cancer | 10X Visium | 5 | 4670*7093 | |
| 29 | | 3L | Simulated | Cancer | 10X Visium | 5 | 4757*7093 | |
| 30 | | 4L | Simulated | Cancer | 10X Visium | 6 | 4113*7093 | |
| 31 | **Breast Atlas** (32) | ST 1 | Simulated | Organs | 10X Visium | 8 | 2103*32664 | Kumar et al. (32) |
| 32 | **Intestine** (33) | A1 | Simulated | Organs | 10X Visium | 4 | 2647*33514 | Fawkner-Corbett et al. (33) |
| 33 | | A2 | Simulated | Organs | 10X Visium | 9 | 2315*13541 | |
| 34 | **Chicken Heart** (33) | D10 | Simulated | Organs | 10X Visium | 6 | 1915*3688 | Mantri et al. (34) |
| 35 | **Prostate Cancer** (30) | | Simulated | Cancer | SlideseqV2 | 11 | 23945*24728 | Hirz et al. (30) |
| 36 | **Cerebellum** (24) | | Simulated | Brain | SlideseqV2 | 13 | 27196*11808 | |

Supplementary Table 12: Details of hyperparamters used for computation of bivariate metrics (l) and simulation of datasets(r)

| Sno | Dataset name | l values | r values |
|---|---|---|---|
| 1 | DLPFC 151508 | 1.2 | 150 |
| 2 | DLPFC 151509 | 1.2 | 150 |
| 3 | DLPFC 151669 | 1.2 | 150 |
| 4 | DLPFC 151670 | 1.2 | 150 |
| 5 | DLPFC 151673 | 1.2 | 150 |
| 6 | DLPFC 151674 | 1.2 | 150 |
| 7 | Mouse Brain ST48 | 1.2 | 250 |
| 8 | Mouse Brain ST52 | 1.2 | 250 |
| 9 | Cerebellum | 50 | 50 |
| 10 | Hippocampus | 50 | 40 |
| 11 | Mouse brain Visium HD | 5 | 100 |
| 12 | Kidney Cancer 410 | 1.2 | 150 |
| 13 | Kidney Cancer 411 | 1.2 | 150 |
| 14 | Kidney Cancer 506 | 1.2 | 150 |
| 15 | Breast Cancer ER+ | 1.2 | 150 |
| 16 | Breast Cancer TNBC | 1.2 | 150 |
| 17 | Liver Cancer HCC-1L | 1.2 | 250 |
| 18 | Liver Cancer HCC-2L | 1.2 | 250 |
| 19 | Liver Cancer HCC-3L | 1.2 | 250 |
| 20 | Liver Cancer HCC-4L | 1.2 | 250 |
| 21 | Prostate Cancer | 50 | 40 |
| 22 | Lung Cancer Visium HD | 5 | 100 |
| 23 | Developmental Lung | 1.2 | 600 |
| 24 | Chicken Heart | 1.2 | 300 |
| 25 | Breast Atlas ST1 | 50 | 100 |
| 26 | Breast Atlas ST8 | 400 | 500 |
| 27 | Intestine A1 | 1.2 | 100 |
| 28 | Intestine A2 | 1.2 | 2 |
| 29 | Liver Altas Mouse - 4 | 1.2 | 50 |
| 30 | Liver Altas Mouse - 12 | 1.2 | 250 |
| 31 | Liver Altas Human - 18 | 1.2 | 250 |
| 32 | Kidney Atlas 1 | 80 | 50 |
| 33 | Kidney Atlas 6 | 80 | 50 |
| 34 | Merfish Brain | 70 | NA |
| 35 | Merfish Ileum | 100 | NA |
| 36 | Merfish Breast Cancer | 70 | NA |
| 37 | Merfish Lung Cancer | 150 | NA |

### Supplementary Note

#### Supplementary Note 1 - Graph attention autoencoder: simulation of cell type proportion matrix

##### Encoder

The graph attention encoder takes as input the cell type proportions for each spot and generates spot embeddings by aggregating information from neighboring spots through an attention-based mechanism. Let $x_i$ denote the cell type proportion vector for spot $i$, $\mathscr{N}_i$ the set of neighbors of spot $i$, and $L$ the number of layers in the encoder. The initial embedding is denoted as $h_i^{(0)} = x_i$. For each subsequent layer $k \in \{1, 2, ..L-1\}$, the embedding is computed as:

$$h_i^{(k)} = \sigma\big(W^{(k)}.\ AGGREGATE\big(h_j^{(k-1)} : j \in \mathscr{N}_i \cup \{i\}\big)\big) \tag{1}$$

where $W^{(k)} \in \mathbb{R}^{d^{(k)} \times d^{(k-1)}}$ is the learnable weight matrix at layer $k$, $d^{(k)}$ denotes the hidden dimensionality at layer $k$, $\sigma$ is a non-linear activation function (e.g., RELU or ELU). The $AGGREGATE$ operation combines the embeddings of spot $i$ and its neighbors, and may be instantiated as a mean, sum, or concatenation.

To capture the similarity between neighboring spots, we employed a self-attention mechanism (63; 64; 65) implemented as a single-layer feed-forward neural network with parameters shared across all nodes. Specifically, at encoder layer $k$, the relevance (attention coefficient) of neighboring node $j$ to node $i$ is computed as:

$$e_{ij}^{(k)} = \sigma_s\left(\mathbf{a}_1^{(k)\top} W^{(k)} h_i^{(k-1)} + \mathbf{a}_2^{(k)\top} W^{(k)} h_j^{(k-1)}\right), \qquad \sigma_s(x) = \frac{1}{1+e^{-x}} \tag{2}$$

where $\mathbf{a}_1^{(k)}, \mathbf{a}_2^{(k)} \in \mathbb{R}^{d^{(k)}}$ are trainable attention parameter vectors at layer $k$, and $\sigma_s$ denotes the sigmoid activation. To make the raw coefficients of all neighbors of node $i$ comparable, they are normalized via a softmax:

$$\alpha_{ij}^{(k)} = \frac{\exp\left(e_{ij}^{(k)}\right)}{\sum\limits_{l \in \mathscr{N}_i} \exp\left(e_{il}^{(k)}\right)} \tag{3}$$

where $\alpha_{ij}^{(k)}$ denotes the normalized attention score of neighbor $j$ with respect to node $i$ at layer $k$. The updated node representation at layer $k$ ($1 \leq k < L$) is then obtained by taking an attention-weighted sum of the transformed neighbor embeddings:

$$h_i^{(k)} = \sigma\left(\sum_{j \in \mathcal{N}_i} \alpha_{ij}^{(k)} W^{(k)} h_j^{(k-1)}\right) \tag{4}$$

The final encoder layer $L$ forgoes the attention mechanism and applies a linear projection followed by a nonlinear activation directly:

$$h_i^{(L)} = \sigma\Big(W^{(L)} h_i^{(L-1)}\Big) \tag{5}$$

The graph attention mechanism at layer $k$ can be expressed compactly in matrix form. Let $H^{(k)} \in \mathbb{R}^{d^{(k)} \times n}$ denote the matrix of spot embeddings at layer $k$, and let $A \in \{0, 1\}^{n \times n}$ be the binary adjacency matrix encoding the spatial neighbourhood graph. The attention matrix $C^{(k)} \in \mathbb{R}^{n \times n}$ is defined entry-wise as:

$$C_{ij}^{(k)} = \begin{cases} \alpha_{ij}^{(k)} & \text{if } A_{ij} = 1, \\ 0 & \text{otherwise,} \end{cases} \tag{6}$$

where $\alpha_{ij}^{(k)}$ is the normalized attention score. In matrix form, $C^{(k)}$ is computed as:

$$C^{(k)} = \text{Softmax}\Big(A \odot \big(\mathbf{a}_1^{(k)\top} W^{(k)} H^{(k-1)}\big) + A \odot \big(\mathbf{a}_2^{(k)\top} W^{(k)} H^{(k-1)}\big)^\top\Big) \tag{7}$$

where $\odot$ denotes the element-wise product, and the softmax is applied row-wise so that each row of $C^{(k)}$ sums to one. The node-embedding matrix at layer $k$ and layer $L$ are then updated as:

$$H^{(k)} = \text{ELU}\Big(W^{(k)} H^{(k-1)} C^{(k)}\Big); \quad H^{(L)} = \text{ELU}\Big(W^{(L)} H^{(L-1)}\Big) \tag{8}$$

**Decoder**

The graph attention decoder mirrors the encoder architecture. Its attention matrices, weight matrices, and input representations are tied to their encoder counterparts: $\hat{C}^{(k)} = C^{(k)}$, $\hat{W}^{(k)} = W^{(k)}$, and $\hat{H}^{(L)} = H^{(L)}$, where $H^{(L)}$ is the latent representation produced by the final encoder layer and serves as the decoder's input. The $k$th layer of the decoder reconstructs the embeddings of layer $k-1$ as:

$$\hat{H}^{(k-1)} = \mathrm{ELU}\Big(\hat{W}^{(k)}\hat{H}^{(k)}\hat{C}^{(k)}\Big) \tag{9}$$

The final decoder layer, which produces the final reconstruction, forgoes attention aggregation and applies a direct linear transformation:

$$\hat{H}^{(0)} = \mathrm{ELU}\Big(\hat{W}^{(1)}\hat{H}^{(1)}\Big) \tag{10}$$

**Inner product decoder**

In addition to the graph attention decoder, we incorporate an inner-product decoder (66) to reconstruct the spatial neighbourhood graph. Concretely, the probability that spots $i$ and $j$ are neighbours is estimated from the inner product of their latent representations:

$$p(A_{ij} = 1 \mid z_i, z_j) = \sigma\Big(z_i^\top z_j\Big), \qquad \sigma(x) = \frac{1}{1+e^{-x}} \tag{11}$$

Assuming conditional independence across all pairs, the full likelihood of the adjacency matrix $A$ given the latent code $Z$ factorizes as:

$$p(A \mid Z) = \prod_{i=1}^{n}\prod_{j=1}^{n} p(A_{ij} \mid z_i, z_j) \tag{12}$$

In matrix form, the reconstructed adjacency matrix is given as:

$$\bar{A} = \sigma\Big(ZZ^\top\Big) \tag{13}$$

**Loss functions**

The graph autoencoder is trained by jointly minimizing three loss functions described below.

**Reconstruction loss.** The first term penalizes the discrepancy between the original cell-type proportion matrix $x$ and its reconstruction $\bar{x}$, measured by the root mean squared error (RMSE):

$$\mathscr{L}_{\text{recon}} = \sqrt{\sum_{i=1}^{n}\sum_{k=1}^{K}(x_{ik} - \bar{x}_{ik})^2} \tag{14}$$

where $n$ is the number of spots and $K$ is the number of cell types.

**Graph reconstruction loss.** The second term is the binary cross-entropy (BCE) between the ground-truth adjacency matrix $A = [a_{ij}]_{i,j=1}^{n}$ and the predicted adjacency matrix $\bar{A} = [\bar{a}_{ij}]_{i,j=1}^{n}$:

$$\mathscr{L}_{\text{BCE}} = -\frac{1}{n^2}\sum_{i=1}^{n}\sum_{j=1}^{n}\Big[a_{ij}\log\bar{a}_{ij} + (1 - a_{ij})\log(1 - \bar{a}_{ij})\Big] \tag{15}$$

**Weight-decay regularization.** To prevent overfitting and discourage the model from over-optimizing any single loss component, we add an $\ell_2$ regularization term over all trainable parameters $\{W_i\}$:

$$\mathscr{L}_W = \frac{1}{2}\sum_i W_i^2 \tag{16}$$

**Total loss.** The full training objective is the sum of the three terms:

$$\mathscr{L}_{\text{total}} = \mathscr{L}_{\text{recon}} + \mathscr{L}_{\text{BCE}} + \mathscr{L}_W \tag{17}$$

The remaining settings and the training parameters needed to train the model are described below.

- SynthST consists of two fully connected feed-forward neural networks, one for the encoder and one for the decoder, each comprising 4 layers: an input layer, two hidden layers and an output layer. Additionally, two graph attention layers are integrated into the encoder, one at each hidden layer, enabling the model to incorporate spatial neighbourhood information during encoding. For training the model, we adopted the ADAM optimizer (67). Training is performed in a single-batch regime, where the full dataset, comprising the cell-type proportion matrix $x$ together with its corresponding adjacency matrix $A \in \{0,1\}^{n\times n}$, is provided as one batch per iteration. To promote training stability and mitigate the exploding-gradient problem, global norm gradient clipping is applied at each update step.

- Construction of adjacency matrix, $A$: The binary adjacency matrix $A \in \{0,1\}^{n\times n}$ is constructed from the pairwise Euclidean distances between spots computed using their spatial coordinates. For each pair of spots $(i, j)$, an edge is placed if and only if their distance falls within a user-defined radius threshold $r$:

$$A_{ij} = \begin{cases} 1 & \text{if } \|coord_i - coord_j\|_2 \le r, \\ 0 & \text{otherwise,} \end{cases} \tag{18}$$

  where $coord_i, coord_j \in \mathbb{R}^2$ denote the spatial coordinates of spots $i$ and $j$, respectively. The threshold $r$ is selected empirically such that each spot has between 6 and 15 neighbours on average. Summary statistics of the resulting neighbourhood sizes across all datasets are provided in Supplementary Table 12.

#### Supplementary Note 2 - Simulation of gene expression dataset

Let $X \in \mathbb{R}^{n\times G}$ denote the observed real spatial gene expression matrix, $\bar{X} \in \mathbb{R}^{n\times G}$ denote the simulated spatial gene expression matrix corresponding to the simulated cell type proportion matrix $\bar{x}^{CTP} \in \mathbb{R}^{n\times K}$ obtained from module 1, where $n$, $K$, and $G$ denote the number of spots, cell types, and genes, respectively. We utilized Cell2location (3) to estimate three quantities: the cell-type gene signature matrix $J \in \mathbb{R}^{K\times G}$, the cell-type abundance matrix $\boldsymbol{W} \in \mathbb{R}^{n\times K}$, and the gene-level sensitivity parameter $m_g$. Let $U_s^X = \sum_{g=1}^{G} X_{sg}$ denote the total UMI count for spot $s$ in the real data.

**Step 1: Estimating per-spot cell counts.** The total number of cells at each spot $s$ is estimated from the abundance matrix $\boldsymbol{W}$ as:

$$n_s = \sum_{k=1}^{K} \boldsymbol{W}_{sk}, \qquad \forall\, s \in \{1, \ldots, n\}. \tag{19}$$

The resulting values are rescaled to lie within a platform-appropriate range: 5–15 cells per spot for 10x Visium data, reflecting typical spot occupancy, and 1–3 cells per spot for higher-resolution platforms such as Slide-seq V2 or 10X Visium HD. The number of cells contributed by cell type $k$ at spot $s$ is then computed as:

$$n_{sk} = \lceil \bar{x}_{sk}^{CTP} \cdot n_s \rceil \tag{20}$$

**Step 2: Generating the simulated gene expression matrix.** The simulated expression $\bar{X}$ is obtained by aggregating the per-cell-type signature profiles, weighted by the estimated cell counts, and scaled by the gene sensitivity parameter:

$$\bar{X}_{sg} = \left( \sum_{k=1}^{K} n_{sk} \, J_{kg} \right) m_g \tag{21}$$

**Step 3: Transferring spatial UMI count distribution via Gaussian Processes.** To ensure that $\bar{X}$ reproduces the global spatial structure of UMI counts present in $X$, we model the target UMI counts for $\bar{X}$ using a Gaussian Process (GP) defined over the observed data:

$$f(X) \sim \mathscr{GP}(\, m(X), \, \Sigma) \tag{22}$$

The mean and covariance functions are defined spot-wise as:

$$m(X_s) = U_s^X \tag{23}$$

$$k(X_{s_1}, X_{s_2}) = \Sigma_{s_1 s_2} = \exp\left( - \frac{d_{s_1 s_2}^2}{\lambda \, \tilde{\sigma}^2} \right) \tag{24}$$

where $d_{s_1 s_2}$ is the Euclidean distance between spots $s_1$ and $s_2$, $\tilde{\sigma}$ is the median pairwise distance across all spots, and $\lambda > 0$ is a length-scale hyperparameter. Target UMI counts $U_s^{\bar{X}}$ are sampled from this GP and used to rescale $\bar{X}$ column-wise:

$$\bar{X}_{sg} \leftarrow \bar{X}_{sg} \cdot \frac{U_s^{\bar{X}}}{\sum_{g=1}^{G} \bar{X}_{sg}} \tag{25}$$

This rescaling preserves the relative gene expression proportions within each spot while aligning the total UMI count of $\bar{X}$ with the spatial pattern learned from $X$.

#### Supplementary Note 3 - Realistic Reconstruction of Spatial Patterns in Simulated Spatial Transcriptomics data

To evaluate the performance of SynthST, we evaluated the performance against some representative datasets spanning tissue categories: brain (DLPFC 151673 and Mouse Brain ST48), cancer (Kidney Cancer 410 and Breast Cancer TNBC), and organ (Breast Atlas ST8 and Intestine A1). For qualitative assessment, we generated spatial pattern plots of cell-type distributions in the simulated ground-truth cell-type proportion (CTP) matrix and compared them with the corresponding CTP matrices used to train SynthST. Specifically, we assessed whether the dominant spatial patterns consistently observed across the majority of training CTP matrices were preserved in the simulated ground-truth CTP matrix. We further validated the simulated spatial patterns by examining marker gene expression profiles in the original spatial gene expression datasets.

In DLPFC 151673, we observed that the major spatial patterns present in the four input cell-type distributions were preserved in the simulated cell-type distributions, including patterns for $L2\ 3$, $Neu\ NRGN\ II$, and oligodendrocytes. Notably, these patterns were retained even when only a subset of methods captured them accurately, for example, Stereoscope and Tangram for L4, and RCTD and Cell2location for $Neu\ NRGN\ 1$. The reconstructed spatial patterns were further validated using marker gene expression from the original dataset Supplementary Figure 21a. A similar trend was observed in the Mouse Brain ST48 dataset, where spatial patterns for $ExtL2/3$ and Ext Thal1 were preserved in the simulated distributions. The simulation also retained patterns captured by only a subset of methods, such as $Astro\ CTX$, $Inh\ 4$, and $Oligo\ 2$. All simulated spatial patterns were validated against marker gene expression profiles from the original dataset Supplementary Figure 21b. For the Kidney Cancer 410 dataset, Supplementary Figure 21c shows spatial pattern comparisons for EC, $Epi\ PT$, Myeloid, RCC, T cells, and pDC. The simulation preserved patterns when they were consistently captured by different subsets of input methods—for example, Stereoscope and Tangram for *Epi PT*; RCTD and Cell2location for Myeloid and RCC; and RCTD, Stereoscope, and Cell2location for T cells. Interestingly, for pDCs, distinct components of the spatial pattern were captured by different methods (RCTD and Cell2location for one region; Stereoscope and Tangram for another). The simulation successfully integrated these complementary patterns, and the resulting distribution was validated using the marker gene IRF7. In the Breast Cancer TNBC and Liver Cancer HCC-3L datasets, the simulation effectively captured the dominant spatial patterns across all evaluated cell types. Supplementary Figure 21d presents spatial pattern plots for DCs, endothelial cells, macrophages, NK cells, and myoepithelial cells in breast cancer, as well as bi-potent cells, CD8+ cells, hepatocytes 3, Tregs, and NK cells in liver cancer. We further conducted qualitative assessments on organ datasets, which exhibit more complex intrinsic spatial organization. In the Chicken Heart dataset, spatial patterns for Cardiomyocytes-1, Epi-epithelial cells, Epi-mesenchymal cells, fibroblasts, immature myocardial cells, and TMSB4X-high cells were preserved, with the simulation capturing dominant patterns across different subsets of input distributions Supplementary Figure 21e. Similarly, Supplementary Figure 21f-g illustrate preserved spatial patterns in the Breast Atlas dataset (basal cells, B cells, fibroblasts, lumHR, lumSEC, lymphatic cells, and T cells) and the Intestine A1 dataset (endothelium, immune cells, muscularis, and pericytes).

Lastly, we evaluated the spatial domains inferred by SynthST in the simulated datasets by validating domain-specific patterns using marker gene expression. In the Kidney Cancer 411 dataset, spatial patterns in domains 0 and 5 were confirmed by the T-cell marker CD3D. The spatial pattern of domain 7 aligned with the expression of ACRA2, a fibroblast marker; domain 3 corresponded to PLVAP expression (endothelial marker); and domain 2 was validated by PT–GPX3 expression, an epithelial marker. Similarly, for the Kidney Cancer 506 dataset, marker genes for endothelial cells, T cells, epithelial cells, and myeloid cells showed clear correspondence with distinct spatial domains. We observed consistent validation across additional cancer datasets, including liver and breast cancer samples, where major cell-type marker genes aligned well with SynthST-identified domains. In organ datasets, the spatial patterns were more intricate: in the chicken heart dataset, domains enriched for cardiomyocytes and fibroblasts were accurately recovered; in the breast atlas dataset, luminal secretory cells, myeloid cells, fibroblasts, and basal cells corresponded to welldefined spatial domains. In the intestine dataset, SynthST not only reproduced the characteristic coiled architecture of the tissue but also accurately identified regions enriched for immune cells Supplementary Figure 23.

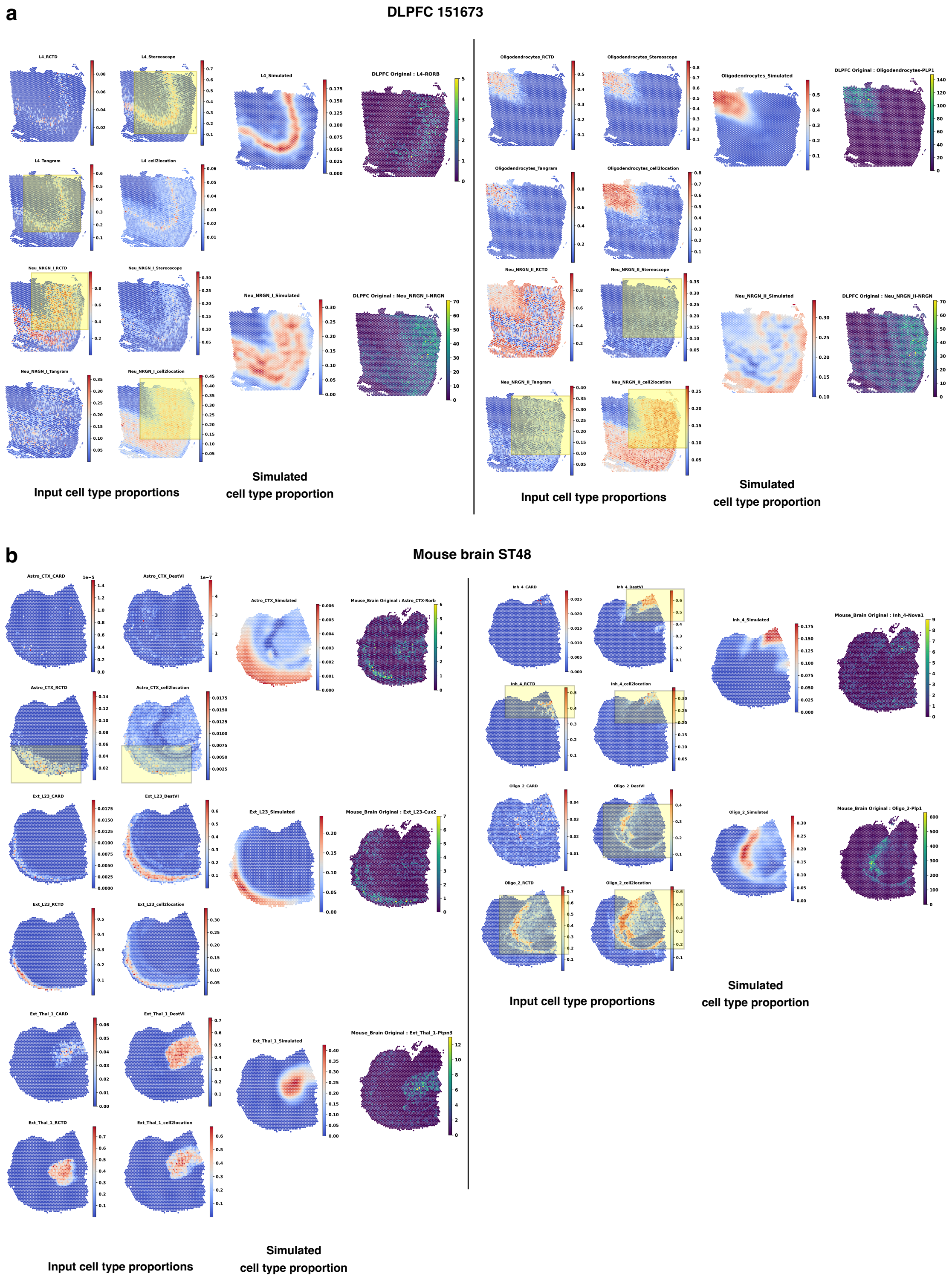

c

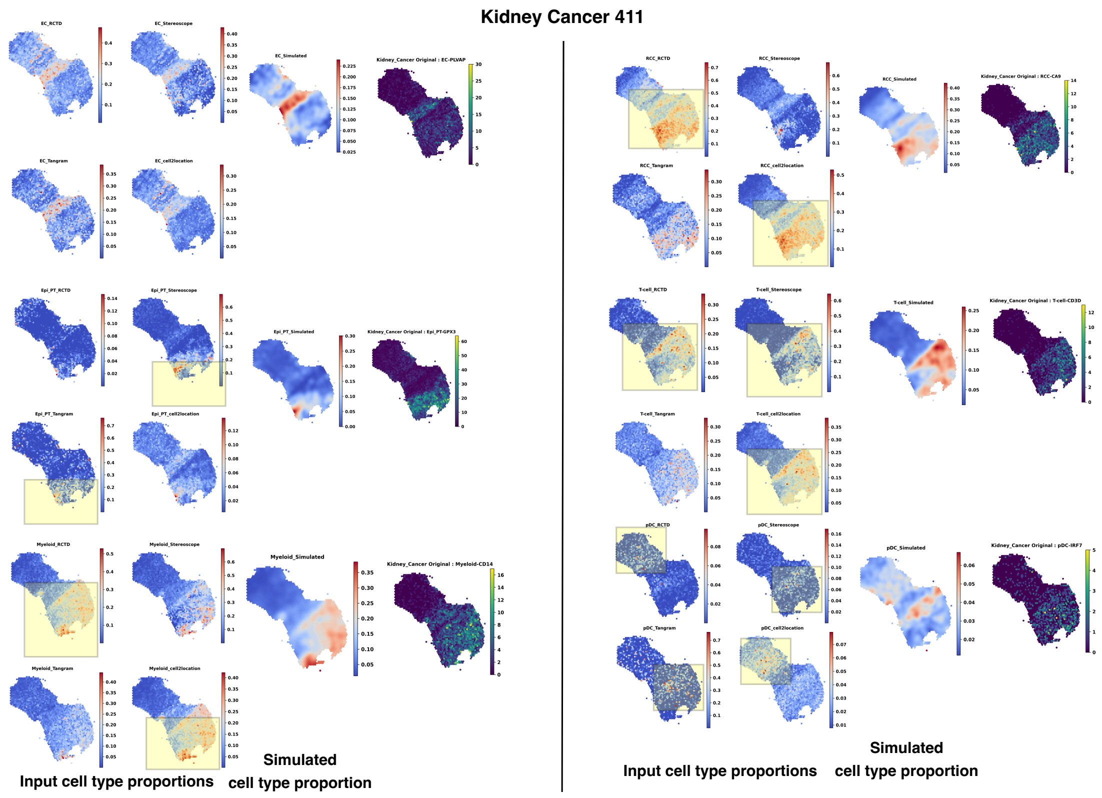
Kidney Cancer 411
Input cell type proportions
Simulated
cell type proportion
Input cell type proportions
Simulated
cell type proportion

d

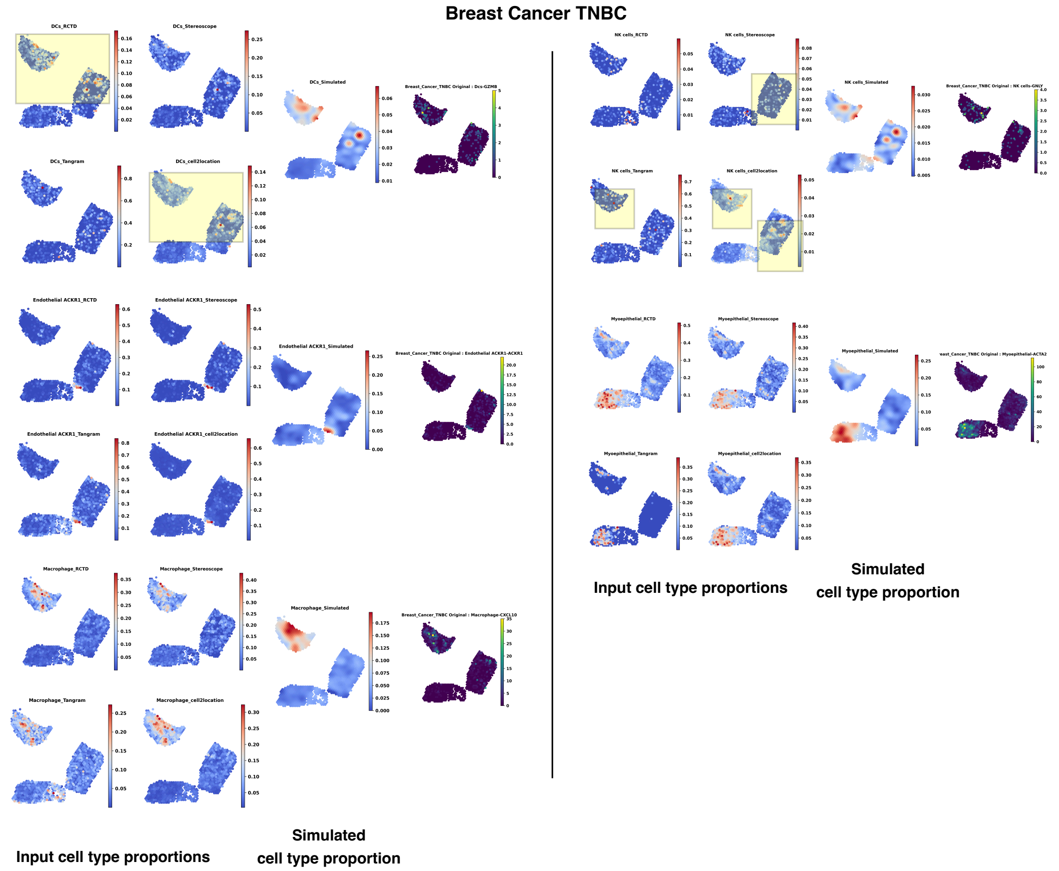
Breast Cancer TNBC
Input cell type proportions
Simulated
cell type proportion
Input cell type proportions
Simulated
cell type proportion

e

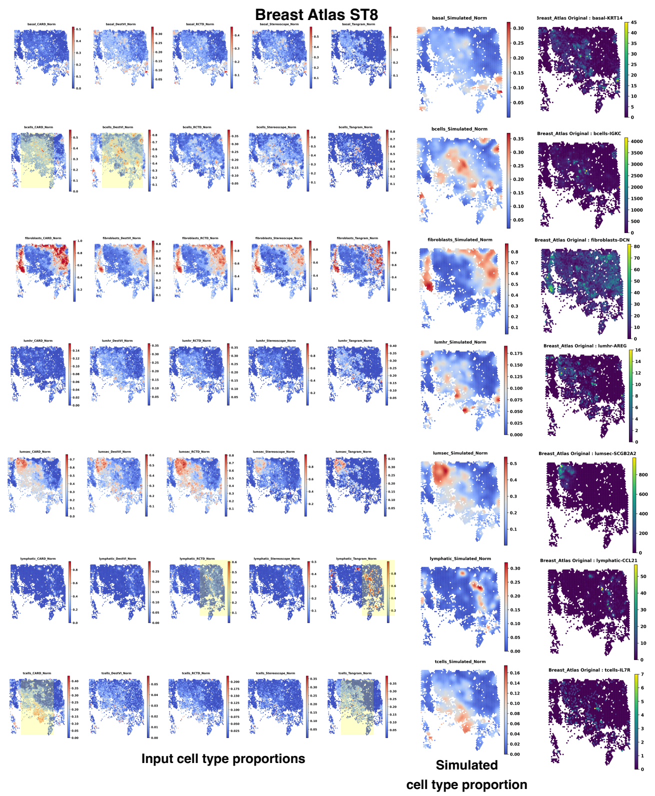
Breast Atlas ST8
basal_Simulated_Norm
bcells_Simulated_Norm
fibroblasts_Simulated_Norm
lumhr_Simulated_Norm
lumsec_Simulated_Norm
lymphatic_Simulated_Norm
tcells_Simulated_Norm
Breast_Atlas Original : bcells-IGKC
Breast_Atlas Original : fibroblasts-DCN
Breast_Atlas Original : lumhr-AREG
Breast_Atlas Original : lumsec-SCGB2A2
Breast_Atlas Original : lymphatic-CCL21
Breast_Atlas Original : tcells-IL7R
Input cell type proportions
Simulated
cell type proportion

f

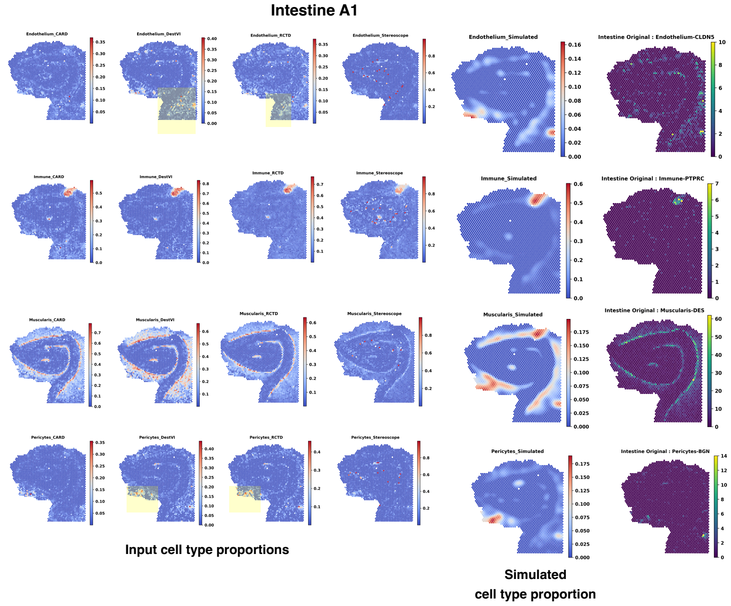
Intestine A1
Endothelium_CARD
Endothelium_DestVI
Endothelium_RCTD
Endothelium_Stereoscope
Endothelium_Simulated
Intestine Original : Endothelium-CLDN5
Immune_CARD
Immune_DestVI
Immune_RCTD
Immune_Stereoscope
Immune_Simulated
Intestine Original : Immune-PTPRC
Muscularis_CARD
Muscularis_DestVI
Muscularis_RCTD
Muscularis_Stereoscope
Muscularis_Simulated
Intestine Original : Muscularis-DES
Pericytes_CARD
Pericytes_DestVI
Pericytes_RCTD
Pericytes_Stereoscope
Pericytes_Simulated
Intestine Original : Pericytes-BGN
Input cell type proportions
Simulated
cell type proportion

Supplementary Figure 21: Qualitative assessment of cell-type proportion matrices used during training SynthST(left), the simulated ground-truth cell-type proportion (CTP) matrix generated during inference (center), and validation of simulated spatial patterns using marker gene expression profiles from the original spatial gene expression data (right), across representative datasets from brain, cancer, and organ tissues. Representative spatial pattern visualizations are shown for (a) DLPFC 151673, (b) Mouse Brain ST48, (c) Kidney Cancer 411, (d) Breast Cancer TNBC, (e) Human Breast ST8, and (f) Intestine datasets

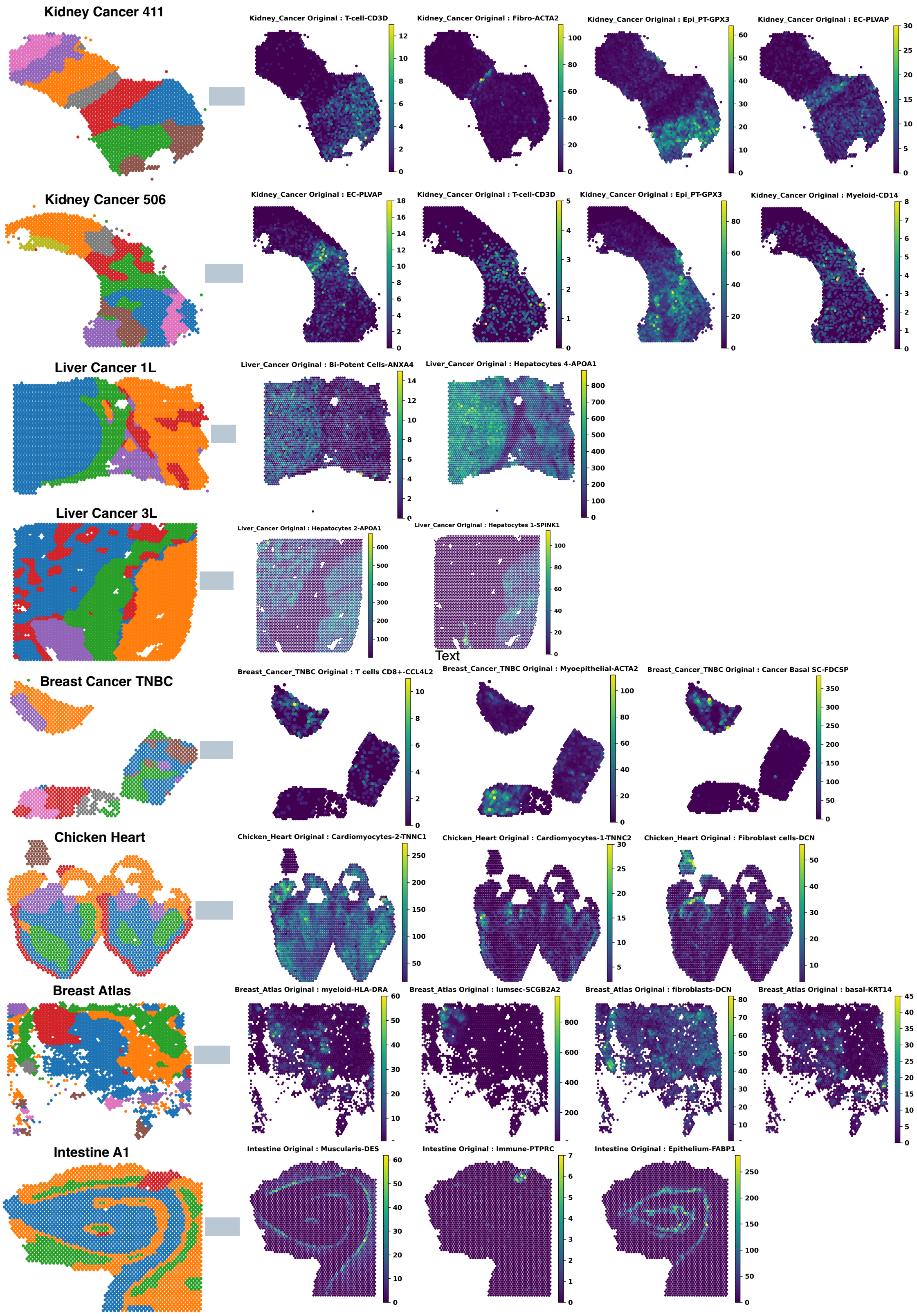


Supplementary Figure 22: Simulated spatial domains for the synthetic spatial gene expression datasets, along with validation using marker gene expression for selected major cell types across representative datasets from brain, cancer, and organ tissues. From top to bottom: Kidney Cancer 411, Kidney Cancer 506, Liver Cancer HCC-1L, Liver Cancer HCC-3L, Breast Cancer TNBC, Chicken Heart, Human Breast, and Intestine A1.
